## Supplementary Files 1-19 for "Mutational sources of *trans-*regulatory variation affecting gene expression in *Saccharomyces cerevisiae*": SupplementaryFile12.R

```
##############################################################################################
################################## Supplementary File 12 #####################################
##############################################################################################
#                R scripts used for the analysis of flow cytometry data.                     #
##############################################################################################

# 1 - R functions used for filtering out artifacts in flow data and to transform                  ===> Line 35
#     fluorescence levels into mRNA levels.
# 2 - Cleaning flow data using functions defined in 1.                                            ===> Line 460
# 3 - Correction for technical variation between samples and calculation of                       ===> Line 546
#     median fluorescence level for each genotype (Figure 1).
# 4 - Hierarchical permutation test to assess statistical significance of                         ===> Line 765
#     fluorescence changes (Figure 1).
# 5 - Plotting expression data shown on Figure 1.                                                 ===> Line 856
# 6 - Correction for technical variation between samples and calculation of                       ===> Line 1203
#     median mRNA level for each genotype (Figure 2 & 6).
# 7 - Permutation tests used to compare the expression of each single-site mutant                 ===> Line 1501
#     to the expression of the progenitor strain.
# 8 - Expression changes caused by linked mutations (Figure 2E).                                  ===> Line 1758
# 9 - Relationship between expression changes in single-site mutants and signed G-values          ===> Line 1828
#     of corresponding mutations (Figure 2F).
# 10- Permutation tests used to compare the expression of each single-site mutant                 ===> Line 2005             
#     to the expression of the EMS mutant carrying the same mutation.             
# 11- Relationship between expression levels of single-site mutants and EMS mutants (Figure 2G).  ===> Line 2227
# 12- Factors contributing to expression differences observed between EMS and                     ===> Line 2287
#     single-site mutants (Figure 2 - figure supplement 5).
# 13- Distribution of expression levels observed among RAP1 mutants (Figure 5E).                  ===> Line 2465
# 14- Distribution of expression levels observed among GCR1 mutants (Figure 5F).                  ===> Line 2531
# 15- Fitness of GCR1 mutants in glucose.                                                         ===> Line 2589
# 16- Relationship between fitness and YFP expression in GCR1 mutants (Figure 5G).                ===> Line 3034
# 17- Effects of mutations in purine biosynthesis genes on YFP expression levels                  ===> Line 3079
#     driven by different promoters(Figure 2 - figure supplement 4).

########################################################################################################################
# 1 - R functions used for filtering out artifacts in flow data and to transform fluorescence levels into mRNA levels. #
########################################################################################################################

### a) ROTATION OF FCS DATA ###

ROT <- function(x,Rotation){
  Result <- Rotation%*%x
  return(Result)
}

#--------------------------------------------------------------------------------------------------------------------------------#

### b) HARD GATE CALIBRATION ###

GATE.CALIB <- function(x,logFSC.A.MIN,logFSC.A.MAX,logFSC.H.MIN,logFSC.H.MAX,FSC.A_FSC.H.MIN,FSC.A_FSC.H.MAX,Width.MIN,Width.MAX) {
  
  if (missing(logFSC.A.MIN)) {logFSC.A.MIN <- 5.1}
  if (missing(logFSC.A.MAX)) {logFSC.A.MAX <- 6.1}
  if (missing(logFSC.H.MIN)) {logFSC.H.MIN <- 5.2}
  if (missing(logFSC.H.MAX)) {logFSC.H.MAX <- 6.8}
  if (missing(FSC.A_FSC.H.MIN)) {FSC.A_FSC.H.MIN <- 0.88}
  if (missing(FSC.A_FSC.H.MAX)) {FSC.A_FSC.H.MAX <- 0.94}
  if (missing(Width.MIN)) {Width.MIN <- 30}
  if (missing(Width.MAX)) {Width.MAX <- 80}
  
  Merge.Frame <- read.FCS(x,transformation=FALSE,alter.names=TRUE)
  
  ##Log transformation of data##

  logTrans <- logTransform(transformationId="log10-transformation",logbase=10,r=1,d=1)
  Merge.Frame <- transform(Merge.Frame,`logFSC.A`=logTrans(`FSC.A`))
  Merge.Frame <- transform(Merge.Frame,`logFSC.H`=logTrans(`FSC.H`))
  Merge.Frame <- transform(Merge.Frame,`logFL1.A`=logTrans(`FL1.A`))
  Merge.Frame <- transform(Merge.Frame,`logFL1.H`=logTrans(`FL1.H`))
  
  ##Calculate phenotypes of interest##

  Data.Fluo <- as.data.frame(exprs(Merge.Frame))
  Data.Fluo[Data.Fluo == 0] <- 1
  
  Phenotype3 <- Data.Fluo[,"logFL1.A"]/Data.Fluo[,"logFSC.A"]
  Phenotype3 <- as.matrix(Phenotype3)
  colnames(Phenotype3) <- "FL1/FSC"
  Merge.Frame <- cbind2(Merge.Frame, Phenotype3)
  
  Phenotype4 <- (Data.Fluo[,"logFSC.A"])/(Data.Fluo[,"logFSC.H"])
  Phenotype4 <- as.matrix(Phenotype4)
  colnames(Phenotype4) <- "FSC.A/FSC.H"
  Merge.Frame <- cbind2(Merge.Frame, Phenotype4)		
  
  PlotAll <- as.data.frame(exprs(Merge.Frame))
  
  
  ###Quick plot to set gates###

  # #Quick plot to set gates
  # quartz(height=14,width=14)
  par(mfrow=c(2,2))
  
  plot(PlotAll[,"Width"],PlotAll[,"logFSC.A"],pch=20,cex=0.5,col="#00000022",xlim=c(20,100),ylim=c(4,7))
  abline(v=Width.MIN)
  abline(v=Width.MAX)
  abline(h=logFSC.A.MIN)
  abline(h=logFSC.A.MAX)
  
  plot(PlotAll[,"logFSC.A"],PlotAll[,"logFL1.A"],pch=20,cex=0.5,col="#00000022",xlim=c(4,7),ylim=c(1.5,6.5))
  abline(v=logFSC.A.MIN)
  abline(v=logFSC.A.MAX)
  
  plot(PlotAll[,"logFSC.H"],PlotAll[,"logFSC.A"],pch=20,cex=0.4,col="#00000022",xlim=c(3,8),ylim=c(4,7))
  abline(v=logFSC.H.MIN)
  abline(v=logFSC.H.MAX)
  abline(h=logFSC.A.MIN)
  abline(h=logFSC.A.MAX)
  
  plot(PlotAll[,"logFSC.A"],PlotAll[,"FSC.A/FSC.H"],pch=20,cex=0.4,col="#00000022",xlim=c(3,8))
  abline(v=logFSC.A.MIN)
  abline(v=logFSC.A.MAX)
  abline(h=FSC.A_FSC.H.MIN)
  abline(h=FSC.A_FSC.H.MAX)
  
  
  
  OUTPUT <- c(logFSC.A.MIN,logFSC.A.MAX,logFSC.H.MIN,logFSC.H.MAX,FSC.A_FSC.H.MIN,FSC.A_FSC.H.MAX,Width.MIN,Width.MAX)
  names(OUTPUT) <- c("logFSC.A.MIN","logFSC.A.MAX","logFSC.H.MIN","logFSC.H.MAX","FSC.A_FSC.H.MIN","FSC.A_FSC.H.MAX","Width.MIN","Width.MAX")
  
  return(OUTPUT)
}


#--------------------------------------------------------------------------------------------------------------------------------#

### c) CLEANING FCS DATA ###

CLEANING <- function(x,GATES) {
  
  Merge.Frame <- read.FCS(x["FILENAMES"],transformation=FALSE,alter.names=TRUE)
  
  OUTPUT <- data.frame(matrix(nrow=1,ncol=length(c("COUNTS.INITIAL",	"COUNTS.GATES",	"COUNTS.SINGLES",	"COUNTS.FINAL", "FSC.KURTOSIS",	"WIDTH",	"FSC.MEDIAN.INITIAL",	"FSC.MAD.INITIAL",	"FL1.MEDIAN.INITIAL",	"FL1.MAD.INITIAL",	"YFP.MEDIAN.INITIAL",	"YFP.MAD.INITIAL", "YFP.SD.INITIAL",	"INTERCEPT.INITIAL",	"SLOPE.INITIAL", "THETA", "YFP.MEDIAN.ROT","YFP.MAD.ROT","YFP.SD.ROT",	"FSC.MEDIAN.FINAL",	"FSC.MAD.FINAL",	"YFP.MEDIAN.FINAL",	"YFP.MAD.FINAL", "YFP.SD.FINAL","log.YFP.MEDIAN","log.YFP.MAD","log.YFP.SD")
  )))
  colnames(OUTPUT) <- c("COUNTS.INITIAL",	"COUNTS.GATES",	"COUNTS.SINGLES",	"COUNTS.FINAL", "FSC.KURTOSIS",	"WIDTH",	"FSC.MEDIAN.INITIAL",	"FSC.MAD.INITIAL",	"FL1.MEDIAN.INITIAL",	"FL1.MAD.INITIAL",	"YFP.MEDIAN.INITIAL",	"YFP.MAD.INITIAL", "YFP.SD.INITIAL",	"INTERCEPT.INITIAL",	"SLOPE.INITIAL", "THETA", "YFP.MEDIAN.ROT","YFP.MAD.ROT","YFP.SD.ROT",	"FSC.MEDIAN.FINAL",	"FSC.MAD.FINAL",	"YFP.MEDIAN.FINAL",	"YFP.MAD.FINAL", "YFP.SD.FINAL","log.YFP.MEDIAN","log.YFP.MAD","log.YFP.SD")
  
  OUTPUT["COUNTS.INITIAL"] <- nrow(exprs(Merge.Frame))
  
  if (OUTPUT["COUNTS.INITIAL"] > 1500 & x["SKIP"] == "NO")	{
    
    ##Log transformation of data##

    Start.exp <- exprs(Merge.Frame)
    Start.exp[,"FL1.A"] <- Start.exp[,"FL1.A"] + 10
    
    Merge.Frame <- new("flowFrame",Start.exp)
    
    Merge.Frame <- transform(Merge.Frame,`logFSC.A`=logTrans(`FSC.A`))
    Merge.Frame <- transform(Merge.Frame,`logFSC.H`=logTrans(`FSC.H`))
    Merge.Frame <- transform(Merge.Frame,`logFL1.A`=logTrans(`FL1.A`))
    Merge.Frame <- transform(Merge.Frame,`logFL1.H`=logTrans(`FL1.H`))
    
    ##Calculate phenotypes of interest##

    Data.Fluo <- as.data.frame(exprs(Merge.Frame))
    Data.Fluo[Data.Fluo == 0] <- 1
    
    Phenotype3 <- Data.Fluo[,"logFL1.A"]/Data.Fluo[,"logFSC.A"]
    Phenotype3 <- as.matrix(Phenotype3)
    colnames(Phenotype3) <- "YFP.INITIAL"
    Merge.Frame <- cbind2(Merge.Frame, Phenotype3)
    
    Phenotype4 <- (Data.Fluo[,"logFSC.A"])/(Data.Fluo[,"logFSC.H"])
    Phenotype4 <- as.matrix(Phenotype4)
    colnames(Phenotype4) <- "FSC.A/FSC.H"
    Merge.Frame <- cbind2(Merge.Frame, Phenotype4)		
    
    PlotAll <- as.data.frame(exprs(Merge.Frame))	
    
    ###Hard Gates###

    rectGate <- rectangleGate(filterId="Noise Removal","logFSC.A"=c(GATES["logFSC.A.MIN"],GATES["logFSC.A.MAX"]), "logFSC.H"=c(GATES["logFSC.H.MIN"],GATES["logFSC.H.MAX"]), "FSC.A/FSC.H"=c(GATES["FSC.A_FSC.H.MIN"],GATES["FSC.A_FSC.H.MAX"]),"Width"=c(GATES["Width.MIN"],GATES["Width.MAX"]),"YFP.INITIAL"=c(min(PlotAll[,"YFP.INITIAL"]),max(PlotAll[,"YFP.INITIAL"])),"FL1.A"=c(11,max(PlotAll[,"FL1.A"])))
    
    Hard.Gates <- Subset(Merge.Frame, rectGate)
    Hard.Gates.exp <- exprs(Hard.Gates)
    
    OUTPUT["COUNTS.GATES"] <- nrow(Hard.Gates.exp)
    
    ###Doublet Hard Gates### 

    Doublet.Model <- PCAgrid(cbind(Hard.Gates.exp[,"logFSC.A"],Hard.Gates.exp[,"FSC.A/FSC.H"]),k=2,method="sd",scores=TRUE,center="median")
    
    Scores <- Doublet.Model$scores
    
    Distri <- normalmixEM2comp(Scores[,2],sigsqrd=c(0.0022,0.0068)^2,mu=c(-0.0013,0.0086),lambda=c(0.56,0.44))
    
    Lambda <- Distri$lambda
    Mu <- Distri$mu
    Sigma <- Distri$sigma 
    
    Order <- c(which(Mu == min(Mu)),which(Mu == max(Mu)))
    
    Lambda <- Lambda[Order]
    Mu <- Mu[Order]
    Sigma <- Sigma[Order] 
    
    #Good cluster
    f <- function(x) dnorm(x,m=Mu[1],sd=Sigma[1])*Lambda[1]-dnorm(x,m=Mu[2],sd=Sigma[2])*Lambda[2]
    Threshold <- try(uniroot(f,interval=c(Mu[1],Mu[2]+Sigma[2]))$root)
    
    # if (Threshold > 0.1)
    # {
    # Threshold <- 0.075
    # }
    
    #Remove big cells based on FSC.A/FSC.H
    Position <- which(Scores[,2] < Threshold)
    Doublet.Gates.exp <- Hard.Gates.exp[Position,]
    Doublet.Gates <- new("flowFrame",Doublet.Gates.exp)
    
    #Remove cells with extreme FSC.A 
    DOUBLETS <- Doublet.Gates.exp[,"logFSC.A"]
    
    MEDIAN <- median(DOUBLETS)
    MAD <- mad(DOUBLETS)
    LOW <- MEDIAN - 2*MAD
    HIGH <- MEDIAN + 2*MAD
    
    NEW.MEDIAN <- MEDIAN
    OLD.MEDIAN <- median(DOUBLETS[which(DOUBLETS > LOW & DOUBLETS < HIGH)])
    
    while (abs(NEW.MEDIAN-OLD.MEDIAN) > 0.001)
    {
      NEW.DOUBLETS <- DOUBLETS[which(DOUBLETS > LOW & DOUBLETS < HIGH)]
      OLD.MEDIAN <- NEW.MEDIAN
      NEW.MEDIAN <- median(NEW.DOUBLETS)
      NEW.MAD <- mad(NEW.DOUBLETS)
      
      LOW <- NEW.MEDIAN - 2*NEW.MAD
      HIGH <- NEW.MEDIAN + 2*NEW.MAD
    }
    
    rectGate <- rectangleGate(filterId="Outliers logFSC.A","logFSC.A"=c(LOW,HIGH))
    
    Final.Doublets <- Subset(Doublet.Gates, rectGate)
    
    OUTPUT["FSC.KURTOSIS"] <- kurtosis(DOUBLETS)
    
    ###Clustering single cells###

    Doublet.filter <- flowClust(Final.Doublets,varNames=c("logFSC.H","logFSC.A"),K=1,B=50,min.count=1000,nu.est=2,trans=0,seed=10,z.cutoff=0,level=0.9,tol=1e-4)
    Well.pop <- split(Final.Doublets,Doublet.filter,population=list(sc1=1))
    Well.Doublet <- Well.pop$sc1
    
    Doublets.exp <- as.data.frame(exprs(Well.Doublet))
    
    OUTPUT["COUNTS.SINGLES"] <- nrow(Doublets.exp)
    
    # plot(Hard.Gates.exp[,"logFSC.A"],Hard.Gates.exp[,"logFSC.H"],pch=20,col="#00000066")
    # points(Doublets.exp[,"logFSC.A"],Doublets.exp[,"logFSC.H"],pch=20,col="#FF000099")
    

    ###Fluo filter###

    #REMOVE OUTLIERS FL1/FSC
    FLUO <- Doublets.exp[,"YFP.INITIAL"]
    
    MEDIAN <- median(FLUO)
    MAD <- mad(FLUO)
    LOW <- MEDIAN - 4*MAD
    HIGH <- MEDIAN + 4*MAD
    
    NEW.MEDIAN <- MEDIAN
    OLD.MEDIAN <- median(FLUO[which(FLUO > LOW & FLUO < HIGH)])
    
    while (abs(NEW.MEDIAN-OLD.MEDIAN) > 0.001)
    {
      NEW.FLUO <- FLUO[which(FLUO > LOW & FLUO < HIGH)]
      OLD.MEDIAN <- NEW.MEDIAN
      NEW.MEDIAN <- median(NEW.FLUO)
      NEW.MAD <- mad(NEW.FLUO)
      
      LOW <- NEW.MEDIAN - 4*NEW.MAD
      HIGH <- NEW.MEDIAN + 4*NEW.MAD
    }
    
    rectGate <- rectangleGate(filterId="Outliers FL1/FSC Removal","YFP.INITIAL"=c(LOW,HIGH))
    
    Hard.Fluo <- Subset(Well.Doublet, rectGate)
    
    Gate.Fluo <- flowClust(Hard.Fluo, varNames=c("logFSC.A","logFL1.A"),K=1,B=50,min.count=1000,nu.est=1,trans=0,z.cutoff=0.5,seed=10,tol=1e-5,nu=1.5,level=0.98)
    Well.pop <- split(Hard.Fluo,Gate.Fluo,population=list(sc1=1))
    Well.Fluo <- Well.pop$sc1
    Fluo.exp <- as.data.frame(exprs(Well.Fluo))
    
    #SAVE DATA
    OUTPUT["COUNTS.FINAL"] <- nrow(Fluo.exp)
    OUTPUT["WIDTH"] <- median(Fluo.exp[,"Width"])
    OUTPUT["FSC.MEDIAN.INITIAL"] <- median(Fluo.exp[,"logFSC.A"])
    OUTPUT["FSC.MAD.INITIAL"] <- mad(Fluo.exp[,"logFSC.A"])
    OUTPUT["FL1.MEDIAN.INITIAL"] <- median(Fluo.exp[,"logFL1.A"])
    OUTPUT["FL1.MAD.INITIAL"] <- mad(Fluo.exp[,"logFL1.A"])	
    OUTPUT["YFP.MEDIAN.INITIAL"] <- median(Fluo.exp[,"YFP.INITIAL"])
    OUTPUT["YFP.MAD.INITIAL"] <- mad(Fluo.exp[,"YFP.INITIAL"])
    OUTPUT["YFP.SD.INITIAL"] <- sd(Fluo.exp[,"YFP.INITIAL"])
    
    # plot(Doublets.exp$logFSC.A,Doublets.exp$logFL1.A,pch=20,col="#00000066")
    # points(Fluo.exp$logFSC.A,Fluo.exp$logFL1.A,pch=20,col="#FF000099")		
    
    ###Remove correlation between logFSC.A and FL1.A/FSC.A###

    #1-Define orthogonal regression
    Intercept <- c()
    Slope <- c()
    Theta <- c()
    
    Fluo.Model <- PCAgrid(cbind(Fluo.exp[,"logFSC.A"],Fluo.exp[,"YFP.INITIAL"]),k=2,method="sd",scores=FALSE,center="median")
    
    #2-Center of rotation
    x.center <- Fluo.Model$center[1]
    y.center <- Fluo.Model$center[2]
    
    #3-Initial Intercept and Slope
    Slope[1] <- Fluo.Model$loadings[2,1] / Fluo.Model$loadings[1,1]
    Intercept[1] <- Fluo.Model$center[2] - Slope[1]*Fluo.Model$center[1]
    
    #4-Calculate angle of rotation
    a <- c(x.center-0,y.center-Intercept[1]) #Vector from Intercept to Centroid
    b <- c(x.center-0,y.center-y.center) #Vector with slope 0 through Centroid
    
    Theta[1] <- acos(sum(a*b)/(sqrt(sum(a*a))*sqrt(sum(b*b)))) #Angle between 2 vectors
    
    if (Slope[1] < 0)
    {
      Theta[1] <- -Theta[1]
    }		
    
    #5-Define rotation matrix
    Rotation <- matrix(c(cos(Theta[1]),-sin(Theta[1]),sin(Theta[1]),cos(Theta[1])),ncol=2,nrow=2)
    
    #6-Transform Data
    Coord <- t(as.matrix(Fluo.exp[,c("logFSC.A","YFP.INITIAL")]))
    
    Coord[1,] <- Coord[1,] - x.center
    Coord[2,] <- Coord[2,] - y.center
    
    Result <- ROT(x=Coord,Rotation=Rotation)
    
    Result[1,] <- Result[1,] + x.center
    Result[2,] <- Result[2,] + y.center
    
    #7-Keep record of rotated values
    Fluo.exp[,"FSC.FINAL"] <- Result[1,]
    Fluo.exp[,"YFP.ROT"] <- Result[2,]
    

    ###Apply correction for linear relation between fluorescence and mRNA levels###

    REF <- 0.905811693
    NEG <- 0.519116913
    
    for (j in 1:nrow(Fluo.exp))
    {
      Fluo.exp[j,"YFP.FINAL"] <- (exp((Fluo.exp[j,"YFP.ROT"]-0.905274742)*log(10)/0.294448097) - 0.05) * (REF - NEG) + NEG
    }
    
    OUTPUT["INTERCEPT.INITIAL"] <- Intercept[1]
    OUTPUT["SLOPE.INITIAL"] <- Slope[1]
    OUTPUT["THETA"] <- Theta[1]
    OUTPUT["YFP.MEDIAN.ROT"] <- median(Fluo.exp[,"YFP.ROT"])
    OUTPUT["YFP.MAD.ROT"] <- mad(Fluo.exp[,"YFP.ROT"])
    OUTPUT["YFP.SD.ROT"] <- sd(Fluo.exp[,"YFP.ROT"])
    OUTPUT["FSC.MEDIAN.FINAL"] <- median(Fluo.exp[,"FSC.FINAL"])
    OUTPUT["FSC.MAD.FINAL"] <- mad(Fluo.exp[,"FSC.FINAL"])	
    OUTPUT["YFP.MEDIAN.FINAL"] <- median(Fluo.exp[,"YFP.FINAL"])
    OUTPUT["YFP.MAD.FINAL"] <- mad(Fluo.exp[,"YFP.FINAL"])	
    OUTPUT["YFP.SD.FINAL"] <- sd(Fluo.exp[,"YFP.FINAL"])	
    OUTPUT["log.YFP.MEDIAN"] <- median(log(Fluo.exp[,"YFP.FINAL"]))
    OUTPUT["log.YFP.MAD"] <- mad(log(Fluo.exp[,"YFP.FINAL"]))	
    OUTPUT["log.YFP.SD"] <- sd(log(Fluo.exp[,"YFP.FINAL"]))	
    
    x["COUNTER"]
    print(x["COUNTER"])
    
    #Save clean data with rotation correction
    Data <- Fluo.exp[,c("Width","Time","logFSC.A","logFSC.H","logFL1.A","logFL1.H","YFP.INITIAL","YFP.ROT","FSC.FINAL","YFP.FINAL")]
    write.table(Data,file=paste("CLEAN.DATA/","Day",x["DAY"],"_Rep",x["REP.UNIQUE"],"_Plate",x["PLATE"],"_Well",x["POSITION"],".txt",sep=""),row.names=FALSE,col.names=TRUE,sep="\t",quote=FALSE)
    
    return(OUTPUT)	
    
  } else {
    OUTPUT[] <- NA
    OUTPUT["COUNTS.INITIAL"] <- nrow(exprs(Merge.Frame))
    
    Data <- as.data.frame(matrix(data=NA,ncol=10))
    colnames(Data) <- c("Width","Time","logFSC.A","logFSC.H","logFL1.A","logFL1.H","YFP.INITIAL","YFP.ROT","FSC.FINAL","YFP.FINAL")
    write.table(Data,file=paste("CLEAN.DATA/","Day",x["DAY"],"_Rep",x["REP.UNIQUE"],"_Plate",x["PLATE"],"_Well",x["POSITION"],".txt",sep=""),row.names=FALSE,col.names=TRUE,sep="\t",quote=FALSE)
    
    return(OUTPUT)
  }
  
}


#--------------------------------------------------------------------------------------------------------------------------------#

### d) QUALITY CONTROL PLOTS ###

FCS.PLOT <- function(x) {
  
  INITIAL.DATA <- read.FCS(x["INITIAL"],transformation=FALSE,alter.names=TRUE)
  CLEAN.DATA <-  read.table(x["CLEAN"],header=TRUE,as.is=TRUE)
  
  ###Log transformation of data###

  Merge.Frame <- transform(INITIAL.DATA,`logFSC.A`=logTrans(`FSC.A`))
  Merge.Frame <- transform(Merge.Frame,`logFSC.H`=logTrans(`FSC.H`))
  Merge.Frame <- transform(Merge.Frame,`logFL1.A`=logTrans(`FL1.A`))
  Merge.Frame <- transform(Merge.Frame,`logFL1.H`=logTrans(`FL1.H`))
  
  ###Calculate phenotypes of interest###

  Data.Fluo <- as.data.frame(exprs(Merge.Frame))
  Data.Fluo[Data.Fluo == 0] <- 1
  
  Phenotype3 <- Data.Fluo[,"logFL1.A"]/Data.Fluo[,"logFSC.A"]
  Phenotype3 <- as.matrix(Phenotype3)
  colnames(Phenotype3) <- "YFP.INITIAL"
  Merge.Frame <- cbind2(Merge.Frame, Phenotype3)
  
  Phenotype4 <- (Data.Fluo[,"logFSC.A"])/(Data.Fluo[,"logFSC.H"])
  Phenotype4 <- as.matrix(Phenotype4)
  colnames(Phenotype4) <- "FSC.A/FSC.H"
  Merge.Frame <- cbind2(Merge.Frame, Phenotype4)		
  
  PLOT.ALL <- as.data.frame(exprs(Merge.Frame))
  
  NAME <- paste("P",x["PLATE"],"R",x["REP"],"_",x["POSITION"],"_",x["STRAIN"],"_",x["CLASS"],sep="")
  
  
  pdf(paste("CLEANING.PLOTS/",NAME,".pdf",sep=""))
  
  par(mfrow=c(2,2))
  
  plot(PLOT.ALL$logFSC.A,PLOT.ALL$logFSC.H,pch=20,cex=0.5,col="#00000033",xlim=c(4.3,6.5),ylim=c(5,7),xlab="logFSC.A",ylab="logFSC.H",main=NAME)
  points(CLEAN.DATA$logFSC.A,CLEAN.DATA$logFSC.H,pch=20,cex=0.5,col="#FF000066",main=NAME)
  
  plot(PLOT.ALL$logFSC.A,PLOT.ALL$logFL1.A,pch=20,cex=0.5,col="#00000033",xlim=c(4.3,6.5),ylim=c(1.5,6.5),xlab="logFSC.A",ylab="logFL1.A",main=NAME)
  points(CLEAN.DATA$logFSC.A,CLEAN.DATA$logFL1.A,pch=20,cex=0.5,col="#FF000066",main=NAME)
  
  plot(CLEAN.DATA$logFSC.A,CLEAN.DATA$YFP.INITIAL,pch=20,cex=0.5,col="#FF0000AA",xlab="logFSC.A",ylab="FL1/FSC",main=NAME)
  abline(lm(CLEAN.DATA$YFP.INITIAL~CLEAN.DATA$logFSC.A),lty=2,col="red",lwd=2.5)
  points(CLEAN.DATA$FSC.FINAL,CLEAN.DATA$YFP.FINAL,pch=20,cex=0.5,col="#00FF00AA",main=NAME)
  abline(lm(CLEAN.DATA$YFP.FINAL~CLEAN.DATA$FSC.FINAL),lty=2,col="green",lwd=2.5)
  
  hist(CLEAN.DATA$YFP.FINAL,breaks=50,main=NAME,xlab="FL1/FSC")
  
  dev.off()	
  
}

#--------------------------------------------------------------------------------------------------------------------------------#

### e) log10 transformation of flow data ###
logTrans <- logTransform(transformationId="log10-transformation",logbase=10,r=1,d=1)


########################################################
# 2 - Cleaning flow data using functions defined in 1. #
########################################################

#Clear memory
rm(list=ls())
options(warn=-1)

### a) LOADING LIBRARIES ###

library(flowCore)
library(flowClust)
library(flowViz)
library(plotrix)
library(nlme)
library(MethComp)
library(outliers)
library(pcaPP)

library(reshape2)
library(MASS)
library(ggplot2)
library(Hmisc)
library(fBasics)
library(lawstat)
library(fitdistrplus)
library(mixtools)
library(vioplot)
library(gplots)
library(RColorBrewer)
library(calibrate)

box <- graphics::box


### b) LOADING FUNCTIONS ###

#File containing R functions defined in section 2.
source("Path.To.Functions/Cleaning.Functions.R")


### c) CLEAN DATA ###

#Set working directory
parent.dir <- "/Path.to.input.file"
setwd(parent.dir)

#Load the template file containing a table with the description of each sample (strain, conditions, plate position).
#A different template file was used for each independent flow cytometry experiment. All template files can be found in SupplementaryFile15.zip.
SETUP <- read.table("Flow.Template/Flow.Template.X.txt",header=TRUE,as.is=TRUE)

#Load paths of FCS files. Warning: The order of FCS files in their folder should be the same as the order of samples in "Flow.Template.X.txt"
FILENAMES <- list.files("..",pattern=".fcs",recursive=TRUE,include.dirs=TRUE,full.names=TRUE)
SETUP[,"FILENAMES"] <- FILENAMES
SETUP[,"COUNTER"] <- c(1:nrow(SETUP))

#Analyze Glucose data.
CUR <- SETUP[1:384,]

#Determine Hard Gates
GATES <- GATE.CALIB(FILENAMES[1])
GATES[2] <- 6.2
GATES[1] <- 5.2
CUR[,"SKIP"] <- "NO"

#Clean Data
Output <- apply(CUR,1,CLEANING,GATES=GATES)


OUTPUT <- as.data.frame(Output[[1]])
OUTPUT[1:nrow(SETUP),] <- NA

for (i in 1:nrow(CUR))
{
  if (is.null(Output[[i]]))
  {} else {
    OUTPUT[i,] <- Output[[i]]
  }
}

CLEAN <- cbind.data.frame(SETUP[,1:(ncol(SETUP)-2)],OUTPUT)

write.table(CLEAN,"Clean.Data.txt",row.names=FALSE,sep="\t")


##########################################################################################################################
# 3 - Correction for technical variation between samples and calculation of median fluorescence level for each genotype. #
##########################################################################################################################

###This script was used to compute the expression values (fluorescence) shown on Figure 1.

###a-LOADING LIBRARIES###
library(flowCore)
library(flowClust)
library(flowViz)
library(pcaPP)
library(mixtools)
library(plyr)
library(robustlmm)

box <- graphics::box

options(warn=-1)

###b-Quality Control and Corrections###
parent.dir <- "/Path.to.input.file"
setwd(parent.dir)

DATA.TYPE <- c(
  rep("factor",18),rep("integer",4),rep("numeric",17))

###Read in data sets
DATA <- read.table("Clean.Data.txt",header=TRUE,colClasses=DATA.TYPE)

###Focus on EMS mutants
TRANS.DATA <- subset(DATA, PLATE %in% c(3:7))

###Remove samples with less than 1000 events
TRANS.DATA <- subset(TRANS.DATA,TRANS.DATA$COUNTS.FINAL >= 1000)

TRANS.DATA <- droplevels(TRANS.DATA)

###Seperate controls and remove FSC outliers
TRANS.CONTROL <- subset(TRANS.DATA,TRANS.DATA$CLASS == "CTRL")

###1-CORRECT FOR FSC MEDIAN###
FSC.MEDIAN.CORRECT <- rlm(TRANS.CONTROL$FSC.MEDIAN.FINAL ~ TRANS.CONTROL$FLOW.RUN)
COEF.FLOW.RUN   <- c(0,coefficients(FSC.MEDIAN.CORRECT)[grep("FLOW.RUN",names(coefficients(FSC.MEDIAN.CORRECT)))])

I <- 1:nrow(TRANS.DATA)
OUT <- numeric(length(I))
for(i in I) {
  FLOW.RUN <- which(levels(TRANS.DATA$FLOW.RUN) == TRANS.DATA$FLOW.RUN[i])
  OUT[i] <- COEF.FLOW.RUN[FLOW.RUN]
}
TRANS.DATA[,"FSC.MEDIAN.CORRECT"] <- TRANS.DATA$FSC.MEDIAN.FINAL - OUT

TRANS.CONTROL <- subset(TRANS.DATA,TRANS.DATA$CLASS == "CTRL")

# plot(TRANS.CONTROL$FSC.MEDIAN.CORRECT ~ TRANS.CONTROL$FLOW.RUN)
# plot(TRANS.CONTROL$FSC.MEDIAN.CORRECT ~ TRANS.CONTROL$ROW)
# plot(TRANS.CONTROL$FSC.MEDIAN.CORRECT ~ TRANS.CONTROL$COL)
# plot(TRANS.CONTROL$FSC.MEDIAN.CORRECT ~ TRANS.CONTROL$POSITION)
# plot(TRANS.CONTROL$FSC.MEDIAN.CORRECT ~ TRANS.CONTROL$PLATE)
# plot(TRANS.CONTROL$FSC.MEDIAN.CORRECT ~ TRANS.CONTROL$REP)

TRANS.CONTROL <- subset(TRANS.DATA,TRANS.DATA$CLASS == "CTRL")
LOW <- median(TRANS.CONTROL$FSC.MEDIAN.CORRECT) - 4*mad(TRANS.CONTROL$FSC.MEDIAN.CORRECT)
HIGH <- median(TRANS.CONTROL$FSC.MEDIAN.CORRECT) + 4*mad(TRANS.CONTROL$FSC.MEDIAN.CORRECT)
TRANS.CONTROL <- subset(TRANS.CONTROL, FSC.MEDIAN.CORRECT >= LOW & FSC.MEDIAN.CORRECT <= HIGH)

###2-CORRECT FOR YFP MEDIAN###
YFP.MEDIAN.CORRECT <- rlmer(YFP.MEDIAN.FINAL ~ REP + ROW + (1|FLOW.RUN:ROW),data=TRANS.CONTROL)
TRANS.DATA[,"YFP.MEDIAN.CORRECT"] <- TRANS.DATA$YFP.MEDIAN.FINAL - predict(YFP.MEDIAN.CORRECT, TRANS.DATA) + mean(TRANS.CONTROL$YFP.MEDIAN.FINAL)
TRANS.CONTROL <- subset(TRANS.DATA,TRANS.DATA$CLASS == "CTRL")
LOW <- median(TRANS.CONTROL$FSC.MEDIAN.CORRECT) - 4*mad(TRANS.CONTROL$FSC.MEDIAN.CORRECT)
HIGH <- median(TRANS.CONTROL$FSC.MEDIAN.CORRECT) + 4*mad(TRANS.CONTROL$FSC.MEDIAN.CORRECT)
TRANS.CONTROL <- subset(TRANS.CONTROL, FSC.MEDIAN.CORRECT >= LOW & FSC.MEDIAN.CORRECT <= HIGH)

#plot(TRANS.CONTROL$YFP.MEDIAN.FINAL ~ TRANS.CONTROL$FLOW.RUN)
#plot(TRANS.CONTROL$YFP.MEDIAN.CORRECT ~ TRANS.CONTROL$FLOW.RUN)
#plot(TRANS.CONTROL$YFP.MEDIAN.CORRECT ~ TRANS.CONTROL$ROW)
#plot(TRANS.CONTROL$YFP.MEDIAN.CORRECT ~ TRANS.CONTROL$COL)
#plot(TRANS.CONTROL$YFP.MEDIAN.CORRECT ~ TRANS.CONTROL$POSITION)
#plot(TRANS.CONTROL$YFP.MEDIAN.CORRECT ~ TRANS.CONTROL$PLATE)
#plot(TRANS.CONTROL$YFP.MEDIAN.CORRECT ~ TRANS.CONTROL$REP)

###2-CORRECT FOR YFP MAD###
YFP.MAD.CORRECT <- rlmer(YFP.MAD.FINAL ~ REP + ROW + (1|FLOW.RUN:ROW),data=TRANS.CONTROL)
TRANS.DATA[,"YFP.MAD.CORRECT"] <- TRANS.DATA$YFP.MAD.FINAL - predict(YFP.MAD.CORRECT, TRANS.DATA) + mean(TRANS.CONTROL$YFP.MAD.FINAL)
TRANS.CONTROL <- subset(TRANS.DATA,TRANS.DATA$CLASS == "CTRL")
LOW <- median(TRANS.CONTROL$FSC.MEDIAN.CORRECT) - 4*mad(TRANS.CONTROL$FSC.MEDIAN.CORRECT)
HIGH <- median(TRANS.CONTROL$FSC.MEDIAN.CORRECT) + 4*mad(TRANS.CONTROL$FSC.MEDIAN.CORRECT)
TRANS.CONTROL <- subset(TRANS.CONTROL, FSC.MEDIAN.CORRECT >= LOW & FSC.MEDIAN.CORRECT <= HIGH)

#plot(TRANS.CONTROL$YFP.MAD.FINAL ~ TRANS.CONTROL$FLOW.RUN)
#plot(TRANS.CONTROL$YFP.MAD.CORRECT ~ TRANS.CONTROL$FLOW.RUN)
#plot(TRANS.CONTROL$YFP.MAD.CORRECT ~ TRANS.CONTROL$ROW)
#plot(TRANS.CONTROL$YFP.MAD.CORRECT ~ TRANS.CONTROL$COL)
#plot(TRANS.CONTROL$YFP.MAD.CORRECT ~ TRANS.CONTROL$POSITION)
#plot(TRANS.CONTROL$YFP.MAD.CORRECT ~ TRANS.CONTROL$PLATE)
#plot(TRANS.CONTROL$YFP.MAD.CORRECT ~ TRANS.CONTROL$REP)

write.table(TRANS.DATA,"DATA.CORRECT.txt",row.names=FALSE,sep="\t",quote=FALSE)


###c- Calculate fluorescence of each sample relative to reference.###

TRANS.DATA <- read.table("DATA.CORRECT.txt",header=TRUE)

WT.1 <- subset(TRANS.DATA, CLASS == "SHAM")
NEG <- subset(TRANS.DATA, CLASS == "NEG")
MUT <- subset(TRANS.DATA, CLASS == "EMS")

LOW <- median(WT.1$FSC.MEDIAN.CORRECT) - 4*mad(WT.1$FSC.MEDIAN.CORRECT)
HIGH <- median(WT.1$FSC.MEDIAN.CORRECT) + 4*mad(WT.1$FSC.MEDIAN.CORRECT)

WT.1 <- subset(WT.1, FSC.MEDIAN.CORRECT < HIGH & FSC.MEDIAN.CORRECT > LOW)

LOW <- median(NEG$FSC.MEDIAN.CORRECT) - 4*mad(NEG$FSC.MEDIAN.CORRECT)
HIGH <- median(NEG$FSC.MEDIAN.CORRECT) + 4*mad(NEG$FSC.MEDIAN.CORRECT)

NEG <- subset(NEG, FSC.MEDIAN.CORRECT < HIGH & FSC.MEDIAN.CORRECT > LOW)

TRANS.DATA[,"YFP.MEAN"] <- (TRANS.DATA$YFP.MEDIAN.CORRECT - median(NEG$YFP.MEDIAN.CORRECT))/median((WT.1$YFP.MEDIAN.CORRECT - median(NEG$YFP.MEDIAN.CORRECT)))
TRANS.DATA[,"YFP.SD"]   <- TRANS.DATA$YFP.MAD.CORRECT/median(WT.1$YFP.MAD.CORRECT)
TRANS.DATA[,"YFP.NOISE"]<- (TRANS.DATA$YFP.MAD.CORRECT/(TRANS.DATA$YFP.MEDIAN.CORRECT - median(NEG$YFP.MEDIAN.CORRECT)))/median((WT.1$YFP.MAD.CORRECT/(WT.1$YFP.MEDIAN.CORRECT - median(NEG$YFP.MEDIAN.CORRECT))))
TRANS.DATA[,"FSC.MEAN"]<- TRANS.DATA$FSC.MEDIAN.CORRECT/median(WT.1$FSC.MEDIAN.CORRECT)

#Write processed data to file
write.table(TRANS.DATA,"TRANS.TEMP.txt",sep="\t",quote=FALSE,row.names=FALSE)


###d- REMOVE OUTLIERS###

TRANS.DATA <- read.table("TRANS.TEMP.txt",header=TRUE)

#BASED ON FSC
for (i in 1:nrow(TRANS.DATA))
{
  CUR <- subset(TRANS.DATA, PLATE == TRANS.DATA[i,"PLATE"] & POSITION == TRANS.DATA[i,"POSITION"])
  
  LOW <- median(CUR$FSC.MEAN) - 6*mad(CUR$FSC.MEAN)
  HIGH <- median(CUR$FSC.MEAN) + 6*mad(CUR$FSC.MEAN)
  
  if (TRANS.DATA[i,"FSC.MEAN"] > LOW & TRANS.DATA[i,"FSC.MEAN"] < HIGH)
  {
    TRANS.DATA[i,"FSC.OUTLIER"] <- "NO"
  } else {
    TRANS.DATA[i,"FSC.OUTLIER"] <- "YES"
  }
}


#BASED ON YFP MEAN
for (i in 1:nrow(TRANS.DATA))
{
  CUR <- subset(TRANS.DATA, PLATE == TRANS.DATA[i,"PLATE"] & POSITION == TRANS.DATA[i,"POSITION"])
  
  LOW <- median(CUR$YFP.MEAN) - 6*mad(CUR$YFP.MEAN)
  HIGH <- median(CUR$YFP.MEAN) + 6*mad(CUR$YFP.MEAN)
  
  if (TRANS.DATA[i,"YFP.MEAN"] > LOW & TRANS.DATA[i,"YFP.MEAN"] < HIGH)
  {
    TRANS.DATA[i,"YFP.OUTLIER"] <- "NO"
  } else {
    TRANS.DATA[i,"YFP.OUTLIER"] <- "YES"
  }
}

#BASED ON YFP MAD
for (i in 1:nrow(TRANS.DATA))
{
  CUR <- subset(TRANS.DATA, PLATE == TRANS.DATA[i,"PLATE"] & POSITION == TRANS.DATA[i,"POSITION"])
  
  LOW <- median(CUR$YFP.MAD.CORRECT) - 6*mad(CUR$YFP.MAD.CORRECT)
  HIGH <- median(CUR$YFP.MAD.CORRECT) + 6*mad(CUR$YFP.MAD.CORRECT)
  
  if (TRANS.DATA[i,"YFP.MAD.CORRECT"] > LOW & TRANS.DATA[i,"YFP.MAD.CORRECT"] < HIGH)
  {
    TRANS.DATA[i,"YFP.MAD.OUTLIER"] <- "NO"
  } else {
    TRANS.DATA[i,"YFP.MAD.OUTLIER"] <- "YES"
  }
}

TRANS.DATA$COLLECTION <- as.character(TRANS.DATA$COLLECTION)

for (i in 1:nrow(TRANS.DATA))
{
  if (is.na(TRANS.DATA$COLLECTION[i]))
  {
    TRANS.DATA$COLLECTION[i] <- "NONE"
  }
}

TRANS.DATA$COLLECTION <- as.factor(TRANS.DATA$COLLECTION)

FILTER.DATA <- subset(TRANS.DATA, YFP.OUTLIER == "NO" & YFP.MAD.OUTLIER == "NO")

write.table(FILTER.DATA,"FILTER.DATA.txt",sep="\t",quote=FALSE,row.names=FALSE)


###e- Estimate strain effect for mean expression and expression noise.###

TRANS.DATA <- read.table("FILTER.DATA.txt",header=TRUE)

TRANS.MEDIAN <- aggregate(cbind(YFP.MEAN,YFP.NOISE,FSC.MEAN) ~ ID + STRAIN + COLLECTION + CLASS + EXPERIMENT, data=FILTER.DATA, FUN = median)
TRANS.SD <- aggregate(cbind(YFP.MEAN,YFP.NOISE,FSC.MEAN) ~ ID + STRAIN + COLLECTION + CLASS + EXPERIMENT, data=FILTER.DATA, FUN = sd)
TRANS.MAD <- aggregate(cbind(YFP.MEAN,YFP.NOISE,FSC.MEAN) ~ ID + STRAIN + COLLECTION + CLASS + EXPERIMENT, data=FILTER.DATA, FUN = mad)
TRANS.N <- aggregate(cbind(YFP.MEAN,YFP.NOISE,FSC.MEAN) ~ ID + STRAIN + COLLECTION + CLASS + EXPERIMENT, data=FILTER.DATA, FUN = length)

TRANS.MEDIAN <- TRANS.MEDIAN[which(TRANS.N[,"YFP.MEAN"] >= 3),]
TRANS.SD <- TRANS.SD[which(TRANS.N[,"YFP.MEAN"] >= 3),]
TRANS.MAD <- TRANS.MAD[which(TRANS.N[,"YFP.MEAN"] >= 3),]
TRANS.N <- TRANS.N[which(TRANS.N[,"YFP.MEAN"] >= 3),]

TRANS <- cbind.data.frame(TRANS.MEDIAN,TRANS.SD[,6:8],TRANS.MAD[,6:8],TRANS.N[,8])
colnames(TRANS) <- c("ID","STRAIN","COLLECTION","CLASS","EXPERIMENT","YFP.MEAN","YFP.NOISE","FSC.MEAN","YFP.MEAN.SD","YFP.NOISE.SD","FSC.MEAN.SD","YFP.MEAN.MAD","YFP.NOISE.MAD","FSC.MEAN.MAD","N")

write.table(TRANS,"SUMMARY.TRANS.txt",sep="\t",quote=FALSE,row.names=FALSE)


############################################################################################################
# 4 - Hierarchical permutation test to assess statistical significance of fluorescence changes (Figure 1). #
############################################################################################################

library(permute)
library(Hmisc)

parent.dir <- "/Path.to.input.file"
setwd(parent.dir)

#The input files are generated with the R script in section 3.
ALL.REP <- read.table("FILTER.DATA.txt",header=TRUE)
ALL.REP <- subset(ALL.REP, YFP.OUTLIER == "NO" & FSC.OUTLIER == "NO")

SUMMARY <- read.table("SUMMARY.DATA.txt",header=TRUE)
SUMMARY <- droplevels(subset(SUMMARY, STRAIN != "1139"))


for (i in 1:nrow(ALL.REP))
{
  N.ZERO <- 2 - nchar(as.character(ALL.REP[i,"PLATE"]))
  
  ALL.REP[i,"STRAIN"] <- paste("1P",rep(0,N.ZERO), ALL.REP[i,"PLATE"], ALL.REP[i,"POSITION"],sep="")
}

WT <- subset(ALL.REP, TREATMENT == "SHAM" & CLASS == "ALL")
LOW.FSC <- median(WT$FSC.MEDIAN.CORRECT) - 4*mad(WT$FSC.MEDIAN.CORRECT)
HIGH.FSC <- median(WT$FSC.MEDIAN.CORRECT) + 4*mad(WT$FSC.MEDIAN.CORRECT)
LOW.FLUO <- median(WT$YFP.MEDIAN.CORRECT) - 4*mad(WT$YFP.MEDIAN.CORRECT)
HIGH.FLUO <- median(WT$YFP.MEDIAN.CORRECT) + 4*mad(WT$YFP.MEDIAN.CORRECT)
WT <- subset(WT, FSC.MEDIAN.CORRECT > LOW.FSC & FSC.MEDIAN.CORRECT < HIGH.FSC & YFP.MEDIAN.CORRECT > LOW.FLUO & YFP.MEDIAN.CORRECT < HIGH.FLUO)

#MEDIAN

#Create list of WT phenotype grouped by strain

N.WT <- length(unique(WT$STRAIN))
WT.LIST <- vector('list',N.WT)
for (i in 1:N.WT)
{
  CUR.WT <- subset(WT, STRAIN == unique(WT$STRAIN)[i])
  WT.LIST[[i]] <- CUR.WT$YFP.MEDIAN.CORRECT 
}


#Create list of all strain phenotypes grouped by strain

SUMMARY <- SUMMARY[order(SUMMARY$STRAIN),]
ALL.REP <- droplevels(subset(ALL.REP,STRAIN %in% as.character(SUMMARY$STRAIN)))
ALL.REP <- ALL.REP[order(ALL.REP$STRAIN),]
N.STRAIN <- length(unique(ALL.REP$STRAIN))
STRAIN.LIST <- vector('list',N.STRAIN)
for (i in 1:N.STRAIN)
{
  CUR.STRAIN <- subset(ALL.REP, STRAIN == unique(ALL.REP$STRAIN)[i])
  STRAIN.LIST[[i]] <- CUR.STRAIN$YFP.MEDIAN.CORRECT
}

#Permutation function

SHUFFLE <- function(STRAIN.LIST = STRAIN.LIST[[i]],WT.LIST = WT.LIST) {
  SAMPLE <- c(STRAIN.LIST,WT.LIST[[sample(1:length(WT.LIST),1)]])
  OUTPUT <- sample(SAMPLE,length(SAMPLE))
  DIST.SIM <- abs(mean(OUTPUT[1:length(STRAIN.LIST)]) - mean(OUTPUT[(length(STRAIN.LIST)+1):length(SAMPLE)]))
  DIST.OBS <- abs(mean(SAMPLE[1:length(STRAIN.LIST)]) - mean(SAMPLE[(length(STRAIN.LIST)+1):length(SAMPLE)]))
  
  DIFF <- DIST.OBS - DIST.SIM
  
  return(DIFF)
}


N.PERM <- 10000

for (i in 1:length(STRAIN.LIST))
{
  DIFF <- sapply(1:N.PERM,FUN=function(x)	SHUFFLE(STRAIN.LIST = STRAIN.LIST[[i]],WT.LIST = WT.LIST))
  P.VAL <- (length(which(DIFF < 0)) + 1) / (length(which(DIFF < 0)) + length(which(DIFF > 0)) + 1) 	
  SUMMARY[i,"P.VAL.MEDIAN"] <- P.VAL
  print(round(i/length(STRAIN.LIST)*100,1))
}

write.table(SUMMARY,"TRANS.MUTANTS.SUMMARY.txt",sep="\t",quote=FALSE,row.names=FALSE)

#A different file name was used for the different experiments shown in Figure 1C, 1D and 1E
#write.table(SUMMARY,"Figure1C-D_SourceData3.txt",sep="\t",quote=FALSE,row.names=FALSE)
#write.table(SUMMARY,"Figure1D_SourceData4.txt",sep="\t",quote=FALSE,row.names=FALSE)
#write.table(SUMMARY,"Figure1E_SourceData5.txt",sep="\t",quote=FALSE,row.names=FALSE)


###################################################
# 5 - Plotting expression data shown on Figure 1. #
###################################################

#Clear memory
rm(list=ls())
options(warn=-1)

#Load packages
library(VariantAnnotation)
library(Deducer)
library(gtools)
library(zoo)
library(Hmisc)
library(ggplot2)
library(plotrix)

#Set directory
setwd("/Path.to.input.files")


### Figure 1D: First screen of random mutants from Metzger et al. 2015 ###

#Input files can be found in SupplementaryFile11.zip
PILOT.SECONDARY <- read.table("Figure1C-D_SourceData3.txt",header=TRUE)
BIG.SECONDARY <- read.table("Figure1D_SourceData4.txt",header=TRUE)
BIG.TERTIARY <- read.table("Figure1E_SourceData5.txt",header=TRUE)

ALL.MUT <- read.table("SourceData1.txt",header=TRUE)

SANGER <- as.character(unique(ALL.MUT[which(ALL.MUT$SEQ.RUN == "SANGER"),"STRAIN"]))
BSA.SEQ <- as.character(unique(ALL.MUT[which(ALL.MUT$SEQ.RUN != "SANGER"),"STRAIN"]))

for (i in 1:nrow(PILOT.SECONDARY))
{
  PILOT.SECONDARY[i,"STRAIN"] <- paste("1P",rep(0,2-nchar(PILOT.SECONDARY[i,"PLATE"])),PILOT.SECONDARY[i,"PLATE"],PILOT.SECONDARY[i,"POSITION"],sep="")
}

#Combine random mutants from pilot and big screens.
RANDOM.PILOT.SECONDARY <- subset(PILOT.SECONDARY, TREATMENT == "EMS" & CLASS == "ALL")
RANDOM.BIG.SECONDARY <- subset(BIG.SECONDARY, TREATMENT == "EMS" & TYPE == "RANDOM")

STRAIN <- c(as.character(RANDOM.PILOT.SECONDARY$STRAIN),as.character(RANDOM.BIG.SECONDARY$STRAIN))
YFP.LEVEL <- c(RANDOM.PILOT.SECONDARY$YFP.MEAN.RELATIVE,RANDOM.BIG.SECONDARY$YFP.MEAN)
YFP.SD <- c(RANDOM.PILOT.SECONDARY$YFP.MEAN.RELATIVE.SD,RANDOM.BIG.SECONDARY$YFP.MEAN.SD)
YFP.N <- c(RANDOM.PILOT.SECONDARY$N,RANDOM.BIG.SECONDARY$N)
P.VALUE <- c(RANDOM.PILOT.SECONDARY$P.VAL.RELATIVE,RANDOM.BIG.SECONDARY$P.VAL.MEAN)

RANDOM.SECONDARY <- data.frame(STRAIN,YFP.LEVEL,YFP.SD,YFP.N,P.VALUE)

#Subset of EMS mutants selected for mapping or Sanger sequencing
for (i in 1:nrow(RANDOM.SECONDARY))
{
  if (RANDOM.SECONDARY[i,"P.VALUE"] < 0.05 & RANDOM.SECONDARY[i,"YFP.LEVEL"] > 1.01 & grepl("2P",RANDOM.SECONDARY[i,"STRAIN"]))
  {
    RANDOM.SECONDARY[i,"COLOR"] <- "blue"
  } else if (RANDOM.SECONDARY[i,"P.VALUE"] < 0.05 & RANDOM.SECONDARY[i,"YFP.LEVEL"] < 0.99 & grepl("2P",RANDOM.SECONDARY[i,"STRAIN"])) {
    RANDOM.SECONDARY[i,"COLOR"] <- "blue"
  } else if (grepl("1P",RANDOM.SECONDARY[i,"STRAIN"]) & RANDOM.SECONDARY[i,"STRAIN"] %in% BSA.SEQ) {
    RANDOM.SECONDARY[i,"COLOR"] <- "red"
  } else if (grepl("1P",RANDOM.SECONDARY[i,"STRAIN"]) & RANDOM.SECONDARY[i,"STRAIN"] %in% SANGER) {
    RANDOM.SECONDARY[i,"COLOR"] <- "green"
  } else {
    RANDOM.SECONDARY[i,"COLOR"] <- "gray"
  }
  if (RANDOM.SECONDARY[i,"P.VALUE"] < 1e-4)
  {
    RANDOM.SECONDARY[i,"P.VALUE"] <- 1e-4
  }
}

length(which(RANDOM.SECONDARY$YFP.LEVEL < 0.99 & RANDOM.SECONDARY$P.VALUE < 0.05))/nrow(RANDOM.SECONDARY)
length(which(RANDOM.SECONDARY$YFP.LEVEL > 1.01 & RANDOM.SECONDARY$P.VALUE < 0.05))/nrow(RANDOM.SECONDARY)

#write.table(RANDOM.SECONDARY,"Random.Secondary.Screen.txt",sep="\t",row.names=FALSE)

for (i in 1:nrow(RANDOM.SECONDARY))
{
  if (RANDOM.SECONDARY$COLOR[i] == "gray")
  {
    RANDOM.SECONDARY[i,"ORDER"] <- 1
  }
  if (RANDOM.SECONDARY$COLOR[i] == "blue")
  {
    RANDOM.SECONDARY[i,"ORDER"] <- 2
  }
  if (RANDOM.SECONDARY$COLOR[i] == "red")
  {
    RANDOM.SECONDARY[i,"ORDER"] <- 4
  }
  if (RANDOM.SECONDARY$COLOR[i] == "green")
  {
    RANDOM.SECONDARY[i,"ORDER"] <- 3
  }
}

RANDOM.SECONDARY <- RANDOM.SECONDARY[order(RANDOM.SECONDARY$ORDER),]

#Plot Figure 1D
pdf("Figure1D.pdf",width=6,height=6,useDingbats = F)
#windows(width=6,height=6)
plotCI(x=RANDOM.SECONDARY[,"YFP.LEVEL"],y=-log10(RANDOM.SECONDARY[,"P.VALUE"]),err="x",uiw=1.96*RANDOM.SECONDARY[,"YFP.SD"]/sqrt(RANDOM.SECONDARY[,"YFP.N"]),xlim=c(0.85,1.15),ylim=c(0,4.5),pch=19,xlab="Fluorescence Level",ylab="-log10(P)",col=RANDOM.SECONDARY$COLOR,sfrac=F,lwd=0.9,main="1st Screen")
abline(v=c(0.99,1.01),h=-log10(0.05),lty=2)
dev.off()

RANDOM.SECONDARY.INCREASE <- subset(RANDOM.SECONDARY, COLOR %in% c("red","green") & YFP.LEVEL > 1)
RANDOM.SECONDARY.DECREASE <- subset(RANDOM.SECONDARY, COLOR %in% c("red","green") & YFP.LEVEL < 1)


### Figure 1E: Second screen of random mutants from Metzger et al. 2015 ###

#Exclude mutants that were included in tertiary screen only because of their effect on noise.
MEAN.BIG.SECONDARY <- subset(RANDOM.BIG.SECONDARY, (P.VAL.MEAN < 0.05 & YFP.MEAN < 0.99) | (P.VAL.MEAN < 0.05 & YFP.MEAN > 1.01))
NOISE.BIG.SECONDARY <- subset(RANDOM.BIG.SECONDARY, YFP.NOISE < 0.95 | YFP.NOISE > 1.05 | P.VAL.NOISE < 0.1)

MEAN.STRAINS <- intersect(MEAN.BIG.SECONDARY$STRAIN, NOISE.BIG.SECONDARY$STRAIN)
NOISE.STRAINS <- as.character(NOISE.BIG.SECONDARY[which(NOISE.BIG.SECONDARY$STRAIN %nin% MEAN.STRAINS),"STRAIN"])

#Combine random mutants from pilot and big screens.
RANDOM.BIG.TERTIARY <- subset(BIG.TERTIARY, STRAIN %nin% NOISE.STRAINS)

STRAIN <- c(as.character(RANDOM.BIG.TERTIARY$STRAIN))
YFP.LEVEL <- c(RANDOM.BIG.TERTIARY$YFP.MEAN)
YFP.SD <- c(RANDOM.BIG.TERTIARY$YFP.MEAN.SD)
YFP.N <- c(RANDOM.BIG.TERTIARY$N)
P.VALUE <- c(RANDOM.BIG.TERTIARY$P.VAL.MEAN)

RANDOM.TERTIARY <- data.frame(STRAIN,YFP.LEVEL,YFP.SD,YFP.N,P.VALUE)

ALL.MUT <- read.table("SourceData1.txt",header=TRUE)

SANGER <- as.character(unique(ALL.MUT[which(ALL.MUT$SEQ.RUN == "SANGER"),"STRAIN"]))
BSA.SEQ <- as.character(unique(ALL.MUT[which(ALL.MUT$SEQ.RUN != "SANGER"),"STRAIN"]))

#Subset of EMS mutants selected for mapping or Sanger sequencing
for (i in 1:nrow(RANDOM.TERTIARY))
{
  RANDOM.TERTIARY[i,"COLOR"] <- "gray"
  
  if (RANDOM.TERTIARY[i,"STRAIN"] %in% SANGER)
  {
    RANDOM.TERTIARY[i,"COLOR"] <- "green"
  }
  
  if (RANDOM.TERTIARY[i,"STRAIN"] %in% BSA.SEQ)
  {
    RANDOM.TERTIARY[i,"COLOR"] <- "red"
  }
  if (RANDOM.TERTIARY[i,"P.VALUE"] < 1e-4)
  {
    RANDOM.TERTIARY[i,"P.VALUE"] <- 1e-4
  }
}

length(which(RANDOM.TERTIARY$YFP.LEVEL < 0.99 & RANDOM.TERTIARY$P.VALUE < 0.05))/nrow(RANDOM.TERTIARY)
length(which(RANDOM.TERTIARY$YFP.LEVEL > 1.01 & RANDOM.TERTIARY$P.VALUE < 0.05))/nrow(RANDOM.TERTIARY)

#write.table(RANDOM.TERTIARY,"Random.Tertiary.Screen.txt",sep="\t",row.names=FALSE)

for (i in 1:nrow(RANDOM.TERTIARY))
{
  if (RANDOM.TERTIARY$COLOR[i] == "gray")
  {
    RANDOM.TERTIARY[i,"ORDER"] <- 1
  }
  if (RANDOM.TERTIARY$COLOR[i] == "blue")
  {
    RANDOM.TERTIARY[i,"ORDER"] <- 2
  }
  if (RANDOM.TERTIARY$COLOR[i] == "red")
  {
    RANDOM.TERTIARY[i,"ORDER"] <- 4
  }
  if (RANDOM.TERTIARY$COLOR[i] == "green")
  {
    RANDOM.TERTIARY[i,"ORDER"] <- 3
  }
}

RANDOM.TERTIARY <- RANDOM.TERTIARY[order(RANDOM.TERTIARY$ORDER),]

#Plot Figure 1E
pdf("Figure1E.pdf",width=6,height=6,useDingbats = F)
#windows(width=6,height=6)
plotCI(x=RANDOM.TERTIARY[,"YFP.LEVEL"],y=-log10(RANDOM.TERTIARY[,"P.VALUE"]),err="x",uiw=1.96*RANDOM.TERTIARY[,"YFP.SD"]/sqrt(RANDOM.TERTIARY[,"YFP.N"]),xlim=c(0.85,1.15),ylim=c(0,4.5),pch=19,xlab="Fluorescence Level",ylab="-log10(P)",col=RANDOM.TERTIARY$COLOR,sfrac=F,lwd=0.9,main="2nd Screen")
abline(v=c(0.99,1.01),h=-log10(0.01),lty=2)
dev.off()

RANDOM.TERTIARY.INCREASE <- subset(RANDOM.TERTIARY, COLOR %in% c("red","green") & YFP.LEVEL > 1)
RANDOM.TERTIARY.DECREASE <- subset(RANDOM.TERTIARY, COLOR %in% c("red","green") & YFP.LEVEL < 1)


### Figure 1C: Mutants enriched for large effects from Metzger et al. 2015 ###

PILOT.SECONDARY <- read.table("Figure1C-D_SourceData3.txt",header=TRUE)

ALL.MUT <- read.table("SourceData1.txt",header=TRUE)

SANGER <- as.character(unique(ALL.MUT[which(ALL.MUT$SEQ.RUN == "SANGER"),"STRAIN"]))
BSA.SEQ <- as.character(unique(ALL.MUT[which(ALL.MUT$SEQ.RUN != "SANGER"),"STRAIN"]))

for (i in 1:nrow(PILOT.SECONDARY))
{
  PILOT.SECONDARY[i,"STRAIN"] <- paste("1P",rep(0,2-nchar(PILOT.SECONDARY[i,"PLATE"])),PILOT.SECONDARY[i,"PLATE"],PILOT.SECONDARY[i,"POSITION"],sep="")
}

#Combine random mutants from pilot and big screens.
TAIL.PILOT.SECONDARY <- subset(PILOT.SECONDARY, (TREATMENT == "EMS" & CLASS == "LOW") | (TREATMENT == "EMS" & CLASS == "HIGH"))

STRAIN <- c(as.character(TAIL.PILOT.SECONDARY$STRAIN))
YFP.LEVEL <- c(TAIL.PILOT.SECONDARY$YFP.MEAN.RELATIVE)
YFP.SD <- c(TAIL.PILOT.SECONDARY$YFP.MEAN.RELATIVE.SD)
YFP.N <- c(TAIL.PILOT.SECONDARY$N)
P.VALUE <- c(TAIL.PILOT.SECONDARY$P.VAL.RELATIVE)

TAIL.SECONDARY <- data.frame(STRAIN,YFP.LEVEL,YFP.SD,YFP.N,P.VALUE)

#Subset of EMS mutants selected for mapping or Sanger sequencing
for (i in 1:nrow(TAIL.SECONDARY))
{
  if (TAIL.SECONDARY[i,"STRAIN"] %in% BSA.SEQ)
  {
    TAIL.SECONDARY[i,"COLOR"] <- "red"
  } else if (TAIL.SECONDARY[i,"STRAIN"] %in% SANGER) {
    TAIL.SECONDARY[i,"COLOR"] <- "green"
  } else {
    TAIL.SECONDARY[i,"COLOR"] <- "gray"
  }
  if (TAIL.SECONDARY[i,"P.VALUE"] < 1e-4)
  {
    TAIL.SECONDARY[i,"P.VALUE"] <- 1e-4
  }
}

length(which(TAIL.SECONDARY$YFP.LEVEL < 0.99 & TAIL.SECONDARY$P.VALUE < 0.05))/nrow(TAIL.SECONDARY)
length(which(TAIL.SECONDARY$YFP.LEVEL > 1.01 & TAIL.SECONDARY$P.VALUE < 0.05))/nrow(TAIL.SECONDARY)

#write.table(TAIL.SECONDARY,"Tail.Secondary.Screen.txt",sep="\t",row.names=FALSE)

for (i in 1:nrow(TAIL.SECONDARY))
{
  if (TAIL.SECONDARY$COLOR[i] == "gray")
  {
    TAIL.SECONDARY[i,"ORDER"] <- 1
  }
  if (TAIL.SECONDARY$COLOR[i] == "blue")
  {
    TAIL.SECONDARY[i,"ORDER"] <- 2
  }
  if (TAIL.SECONDARY$COLOR[i] == "red")
  {
    TAIL.SECONDARY[i,"ORDER"] <- 4
  }
  if (TAIL.SECONDARY$COLOR[i] == "green")
  {
    TAIL.SECONDARY[i,"ORDER"] <- 3
  }
}

TAIL.SECONDARY <- TAIL.SECONDARY[order(TAIL.SECONDARY$ORDER),]

pdf("Figure1C.pdf",width=6,height=6,useDingbats = F)
#windows(width=6,height=6)
plotCI(x=TAIL.SECONDARY[,"YFP.LEVEL"],y=-log10(TAIL.SECONDARY[,"P.VALUE"]),err="x",uiw=1.96*TAIL.SECONDARY[,"YFP.SD"]/sqrt(TAIL.SECONDARY[,"YFP.N"]),xlim=c(0.85,1.15),ylim=c(0,4.5),pch=19,xlab="Fluorescence Level",ylab="-log10(P)",col=TAIL.SECONDARY$COLOR,sfrac=F,lwd=0.9,main="1st Screen")
abline(v=c(0.99,1.01),h=-log10(0.05),lty=2)
dev.off()

TAIL.SECONDARY.INCREASE <- subset(TAIL.SECONDARY, COLOR %in% c("red","green") & YFP.LEVEL > 1)
TAIL.SECONDARY.DECREASE <- subset(TAIL.SECONDARY, COLOR %in% c("red","green") & YFP.LEVEL < 1)


### Figure 1B: Mutants enriched for large effects from Gruber et al. 2012 ###

GRUBER.DATA <- read.table("Figure1B_SourceData2.txt",header=TRUE)

NEGATIVE <- subset(GRUBER.DATA, sample == "8Q2A10")
BSA.SEQ <- c("7Q4D5","8Q4H5","4Q2G6","2Q1H10","1Q4G11","1Q4H10")

TAIL.GRUBER <- subset(GRUBER.DATA, class == "hi.EMS" | class == "lo.EMS")

STRAIN <- c(as.character(TAIL.GRUBER$sample))
YFP.LEVEL <- rep(0,length(STRAIN))
YFP.SD <- rep(0,length(STRAIN))
YFP.N <- rep(1,length(STRAIN))
P.VALUE <- rep(0,length(STRAIN))

TAIL.GRUBER.SECONDARY <- data.frame(STRAIN,YFP.LEVEL,YFP.SD,YFP.N,P.VALUE)

#Subset of EMS mutants selected for mapping or Sanger sequencing
for (i in 1:nrow(TAIL.GRUBER.SECONDARY))
{
  TAIL.GRUBER.SECONDARY[i,"YFP.LEVEL"] <- (TAIL.GRUBER[i,"unk.RATIOmedian.in.range"] - NEGATIVE[1,"unk.RATIOmedian.in.range"]) / (TAIL.GRUBER[i,"ref.RATIOmedian.in.range"] - NEGATIVE[1,"unk.RATIOmedian.in.range"])
  TAIL.GRUBER.SECONDARY[i,"P.VALUE"] <- 2 * pnorm(-abs(TAIL.GRUBER[i,"Z.YFPmedians"]))
}

for (i in 1:nrow(TAIL.GRUBER.SECONDARY))
{
  if (TAIL.GRUBER.SECONDARY[i,"STRAIN"] %in% BSA.SEQ)
  {
    TAIL.GRUBER.SECONDARY[i,"COLOR"] <- "red"
  } else {
    TAIL.GRUBER.SECONDARY[i,"COLOR"] <- "gray"
  }
  if (TAIL.GRUBER.SECONDARY[i,"P.VALUE"] < 1e-40)
  {
    TAIL.GRUBER.SECONDARY[i,"P.VALUE"] <- 1e-40
  }
}

#write.table(TAIL.GRUBER.SECONDARY,"Tail.Gruber.Screen.txt",sep="\t",row.names=FALSE)

for (i in 1:nrow(TAIL.GRUBER.SECONDARY))
{
  if (TAIL.GRUBER.SECONDARY$COLOR[i] == "gray")
  {
    TAIL.GRUBER.SECONDARY[i,"ORDER"] <- 1
  }
  if (TAIL.GRUBER.SECONDARY$COLOR[i] == "blue")
  {
    TAIL.GRUBER.SECONDARY[i,"ORDER"] <- 2
  }
  if (TAIL.GRUBER.SECONDARY$COLOR[i] == "red")
  {
    TAIL.GRUBER.SECONDARY[i,"ORDER"] <- 4
  }
  if (TAIL.GRUBER.SECONDARY$COLOR[i] == "green")
  {
    TAIL.GRUBER.SECONDARY[i,"ORDER"] <- 3
  }
}

length(which(TAIL.GRUBER.SECONDARY$YFP.LEVEL < 0.99 & TAIL.GRUBER.SECONDARY$P.VALUE < 0.05))/nrow(TAIL.GRUBER.SECONDARY)
length(which(TAIL.GRUBER.SECONDARY$YFP.LEVEL > 1.01 & TAIL.GRUBER.SECONDARY$P.VALUE < 0.05))/nrow(TAIL.GRUBER.SECONDARY)

TAIL.GRUBER.SECONDARY <- TAIL.GRUBER.SECONDARY[order(TAIL.GRUBER.SECONDARY$ORDER),]

pdf("Figure1B.pdf",width=6,height=6,useDingbats = F)
#windows(width=6,height=6)
plotCI(x=TAIL.GRUBER.SECONDARY[,"YFP.LEVEL"],y=-log10(TAIL.GRUBER.SECONDARY[,"P.VALUE"]),err="x",uiw=1.96*TAIL.GRUBER.SECONDARY[,"YFP.SD"]/sqrt(TAIL.GRUBER.SECONDARY[,"YFP.N"]),xlim=c(0.75,1.25),ylim=c(0,45),pch=19,xlab="Fluorescence Level",ylab="-log10(P)",col=TAIL.GRUBER.SECONDARY$COLOR,sfrac=F,lwd=0.9,main="1st Screen")
abline(v=c(0.99,1.01),h=-log10(0.05),lty=2)
dev.off()

TAIL.GRUBER.INCREASE <- subset(TAIL.GRUBER.SECONDARY, COLOR %in% c("red","green") & YFP.LEVEL > 1)
TAIL.GRUBER.DECREASE <- subset(TAIL.GRUBER.SECONDARY, COLOR %in% c("red","green") & YFP.LEVEL < 1)


##################################################################################################################
# 6 - Correction for technical variation between samples and calculation of median mRNA level for each genotype. #
##################################################################################################################

###This script was used to compute the expression values shown on Figures 2, 3 and 6.

### a) LOADING LIBRARIES ###

#Clear memory
rm(list=ls())
options(warn=-1)

library(flowCore)
library(flowClust)
library(flowViz)
library(pcaPP)
library(mixtools)
library(plyr)
library(robustlmm)
library(plotrix)
library(MASS)

box <- graphics::box

### b) Load data generated in section 2. ###

parent.dir <- "/Path.to.input.file"
setwd(parent.dir)

DATA.TYPE <- c(
  rep("factor",22),rep("integer",4),rep("numeric",24))

###Read in data sets
DATA <- read.table("Clean.Data.txt",header=TRUE,colClasses=DATA.TYPE)

### c) Correction for plate effects. ###
REF <- 0.905811693
NEG <- 0.519116913

DATA[,"log.RNA.MEDIAN"] <- log(((DATA[,"YFP.MEDIAN.FINAL"] - NEG) / (REF - NEG)) + 0.05)
DATA[,"log.RNA.SD"] <- DATA[,"YFP.SD.FINAL"] / ((DATA[,"YFP.MEDIAN.FINAL"] - NEG) + (REF - NEG)*0.05)


DATA[,"RUN"] <- as.factor(DATA[,"RUN"])
DATA[,"ROW"] <- as.factor(DATA[,"ROW"])

TRANS.DATA <- DATA
TRANS.DATA <- droplevels(TRANS.DATA)

###Separate controls and remove FSC outliers
TRANS.CONTROL <- subset(TRANS.DATA, TRANS.DATA$ID.1 == "CTRL")


###1-CORRECT FOR FSC MEDIAN###
FSC.MEDIAN.CORRECT <- rlm(FSC.MEDIAN.FINAL ~ RUN, data = TRANS.CONTROL)
COEF.FLOW.RUN   <- c(0,coefficients(FSC.MEDIAN.CORRECT)[grep("RUN",names(coefficients(FSC.MEDIAN.CORRECT)))])

I <- 1:nrow(TRANS.DATA)
OUT <- numeric(length(I))
for(i in I) {
  RUN <- which(levels(TRANS.DATA$RUN) == TRANS.DATA$RUN[i])
  OUT[i] <- COEF.FLOW.RUN[RUN]
}
TRANS.DATA[,"FSC.MEDIAN.CORRECT"] <- TRANS.DATA$FSC.MEDIAN.FINAL - OUT

TRANS.CONTROL <- subset(TRANS.DATA,TRANS.DATA$ID.1 == "CTRL")

FSC.MEDIAN.CORRECT <- rlm(FSC.MEDIAN.CORRECT ~ ROW, data = TRANS.CONTROL)
COEF.ROW   <- c(0,coefficients(FSC.MEDIAN.CORRECT)[grep("ROW",names(coefficients(FSC.MEDIAN.CORRECT)))])

I <- 1:nrow(TRANS.DATA)
OUT <- numeric(length(I))
for(i in I) {
  ROW <- which(levels(TRANS.DATA$ROW) == TRANS.DATA$ROW[i])
  OUT[i] <- COEF.ROW[ROW]
}

TRANS.DATA[,"FSC.MEDIAN.CORRECT"] <- TRANS.DATA$FSC.MEDIAN.CORRECT - OUT

TRANS.CONTROL <- subset(TRANS.DATA,TRANS.DATA$ID.1 == "CTRL")

# plot(TRANS.CONTROL$FSC.MEDIAN.FINAL ~ TRANS.CONTROL$RUN)
# plot(TRANS.CONTROL$FSC.MEDIAN.CORRECT ~ TRANS.CONTROL$RUN)
# plot(TRANS.CONTROL$FSC.MEDIAN.CORRECT ~ TRANS.CONTROL$ROW)
# plot(TRANS.CONTROL$FSC.MEDIAN.CORRECT ~ TRANS.CONTROL$COL)
# plot(TRANS.CONTROL$FSC.MEDIAN.CORRECT ~ TRANS.CONTROL$POSITION)
# plot(TRANS.CONTROL$FSC.MEDIAN.CORRECT ~ TRANS.CONTROL$PLATE)
# plot(TRANS.CONTROL$FSC.MEDIAN.CORRECT ~ TRANS.CONTROL$REP)
# plot(TRANS.CONTROL$FSC.MEDIAN.CORRECT ~ TRANS.CONTROL$DAY)

LOW <- median(TRANS.CONTROL$FSC.MEDIAN.CORRECT) - 4*mad(TRANS.CONTROL$FSC.MEDIAN.CORRECT)
HIGH <- median(TRANS.CONTROL$FSC.MEDIAN.CORRECT) + 4*mad(TRANS.CONTROL$FSC.MEDIAN.CORRECT)
TRANS.CONTROL <- subset(TRANS.CONTROL, FSC.MEDIAN.CORRECT >= LOW & FSC.MEDIAN.CORRECT <= HIGH)

###2-CORRECT FOR YFP MEDIAN###
YFP.MEDIAN.CORRECT <- rlm(log.RNA.MEDIAN ~ 0 + RUN, data = TRANS.CONTROL)

COEF.FLOW.RUN   <- c(coefficients(YFP.MEDIAN.CORRECT)[grep("RUN",names(coefficients(YFP.MEDIAN.CORRECT)))])

I <- 1:nrow(TRANS.DATA)
OUT <- numeric(length(I))
for(i in I) {
  RUN <- which(levels(TRANS.DATA$RUN) == TRANS.DATA$RUN[i])
  OUT[i] <- COEF.FLOW.RUN[RUN]
}

TRANS.DATA[,"log.RNA.MEDIAN.CORRECT"] <- TRANS.DATA$log.RNA.MEDIAN - OUT + mean(TRANS.CONTROL$log.RNA.MEDIAN)

TRANS.CONTROL <- subset(TRANS.DATA,TRANS.DATA$ID.1 == "CTRL")
TRANS.CONTROL <- subset(TRANS.CONTROL, FSC.MEDIAN.CORRECT >= LOW & FSC.MEDIAN.CORRECT <= HIGH)

YFP.MEDIAN.CORRECT <- rlm(log.RNA.MEDIAN.CORRECT ~ 0 + ROW, data = TRANS.CONTROL)
COEF.ROW   <- c(coefficients(YFP.MEDIAN.CORRECT)[grep("ROW",names(coefficients(YFP.MEDIAN.CORRECT)))])

I <- 1:nrow(TRANS.DATA)
OUT <- numeric(length(I))
for(i in I) {
  ROW <- which(levels(TRANS.DATA$ROW) == TRANS.DATA$ROW[i])
  OUT[i] <- COEF.ROW[ROW]
}

TRANS.DATA[,"log.RNA.MEDIAN.CORRECT"] <- TRANS.DATA$log.RNA.MEDIAN.CORRECT - OUT + mean(TRANS.CONTROL$log.RNA.MEDIAN.CORRECT)
TRANS.DATA[,"YFP.MEDIAN.CORRECT"] <- (exp(TRANS.DATA[,"log.RNA.MEDIAN.CORRECT"]) - 0.05) * (REF - NEG) + NEG

TRANS.CONTROL <- subset(TRANS.DATA,TRANS.DATA$ID.1 == "CTRL")
TRANS.CONTROL <- subset(TRANS.CONTROL, FSC.MEDIAN.CORRECT >= LOW & FSC.MEDIAN.CORRECT <= HIGH)

# plot(TRANS.CONTROL$log.RNA.MEDIAN ~ TRANS.CONTROL$RUN)
# plot(TRANS.CONTROL$log.RNA.MEDIAN.CORRECT ~ TRANS.CONTROL$RUN)
# plot(TRANS.CONTROL$log.RNA.MEDIAN.CORRECT ~ TRANS.CONTROL$ROW)
# plot(TRANS.CONTROL$log.RNA.MEDIAN.CORRECT ~ TRANS.CONTROL$COL)
# plot(TRANS.CONTROL$log.RNA.MEDIAN.CORRECT ~ TRANS.CONTROL$POSITION)
# plot(TRANS.CONTROL$log.RNA.MEDIAN.CORRECT ~ TRANS.CONTROL$PLATE)
# plot(TRANS.CONTROL$log.RNA.MEDIAN.CORRECT ~ TRANS.CONTROL$REP)
# plot(TRANS.CONTROL$log.RNA.MEDIAN.CORRECT ~ TRANS.CONTROL$DAY)
# plot(TRANS.CONTROL$log.RNA.MEDIAN.CORRECT ~ TRANS.CONTROL$BLOCK)

# plot(TRANS.CONTROL$YFP.MEDIAN.FINAL ~ TRANS.CONTROL$RUN)
# plot(TRANS.CONTROL$YFP.MEDIAN.CORRECT ~ TRANS.CONTROL$RUN)
# plot(TRANS.CONTROL$YFP.MEDIAN.CORRECT ~ TRANS.CONTROL$ROW)
# plot(TRANS.CONTROL$YFP.MEDIAN.CORRECT ~ TRANS.CONTROL$COL)
# plot(TRANS.CONTROL$YFP.MEDIAN.CORRECT ~ TRANS.CONTROL$POSITION)
# plot(TRANS.CONTROL$YFP.MEDIAN.CORRECT ~ TRANS.CONTROL$PLATE)
# plot(TRANS.CONTROL$YFP.MEDIAN.CORRECT ~ TRANS.CONTROL$REP)
# plot(TRANS.CONTROL$YFP.MEDIAN.CORRECT ~ TRANS.CONTROL$DAY)
# plot(TRANS.CONTROL$YFP.MEDIAN.CORRECT ~ TRANS.CONTROL$BLOCK)


###3-CORRECT FOR YFP SD ON LOG SCALE###
YFP.MAD.CORRECT <- rlm(log.RNA.SD ~ 0 + RUN, data = TRANS.CONTROL)
COEF.FLOW.RUN   <- c(coefficients(YFP.MAD.CORRECT)[grep("RUN",names(coefficients(YFP.MAD.CORRECT)))])

I <- 1:nrow(TRANS.DATA)
OUT <- numeric(length(I))
for(i in I) {
  RUN <- which(levels(TRANS.DATA$RUN) == TRANS.DATA$RUN[i])
  OUT[i] <- COEF.FLOW.RUN[RUN]
}

TRANS.DATA[,"log.RNA.SD.CORRECT"] <- TRANS.DATA$log.RNA.SD - OUT + mean(TRANS.CONTROL$log.RNA.SD) 

TRANS.CONTROL <- subset(TRANS.DATA,TRANS.DATA$ID.1 == "CTRL")
TRANS.CONTROL <- subset(TRANS.CONTROL, FSC.MEDIAN.CORRECT >= LOW & FSC.MEDIAN.CORRECT <= HIGH)

YFP.MAD.CORRECT <- rlm(log.RNA.SD.CORRECT ~ 0 + ROW, data = TRANS.CONTROL)
COEF.ROW   <- c(coefficients(YFP.MAD.CORRECT)[grep("ROW",names(coefficients(YFP.MAD.CORRECT)))])

I <- 1:nrow(TRANS.DATA)
OUT <- numeric(length(I))
for(i in I) {
  ROW <- which(levels(TRANS.DATA$ROW) == TRANS.DATA$ROW[i])
  OUT[i] <- COEF.ROW[ROW]
}

TRANS.DATA[,"log.RNA.SD.CORRECT"] <- TRANS.DATA$log.RNA.SD.CORRECT - OUT + mean(TRANS.CONTROL$log.RNA.SD.CORRECT)
TRANS.DATA[,"YFP.SD.CORRECT"] <- TRANS.DATA[,"log.RNA.SD.CORRECT"] * ((TRANS.DATA[,"YFP.MEDIAN.CORRECT"] - NEG) + (REF - NEG)*0.05)

TRANS.CONTROL <- subset(TRANS.DATA,TRANS.DATA$ID.1 == "CTRL")
TRANS.CONTROL <- subset(TRANS.CONTROL, FSC.MEDIAN.CORRECT >= LOW & FSC.MEDIAN.CORRECT <= HIGH)


# plot(TRANS.CONTROL$log.RNA.SD ~ TRANS.CONTROL$RUN)
# plot(TRANS.CONTROL$log.RNA.SD.CORRECT ~ TRANS.CONTROL$RUN)
# plot(TRANS.CONTROL$log.RNA.SD.CORRECT ~ TRANS.CONTROL$ROW)
# plot(TRANS.CONTROL$log.RNA.SD.CORRECT ~ TRANS.CONTROL$COL)
# plot(TRANS.CONTROL$log.RNA.SD.CORRECT ~ TRANS.CONTROL$POSITION)
# plot(TRANS.CONTROL$log.RNA.SD.CORRECT ~ TRANS.CONTROL$PLATE)
# plot(TRANS.CONTROL$log.RNA.SD.CORRECT ~ TRANS.CONTROL$REP)
# plot(TRANS.CONTROL$log.RNA.SD.CORRECT ~ TRANS.CONTROL$DAY)


# plot(TRANS.CONTROL$YFP.SD.FINAL ~ TRANS.CONTROL$RUN)
# plot(TRANS.CONTROL$YFP.SD.CORRECT ~ TRANS.CONTROL$RUN)
# plot(TRANS.CONTROL$YFP.SD.CORRECT ~ TRANS.CONTROL$ROW)
# plot(TRANS.CONTROL$YFP.SD.CORRECT ~ TRANS.CONTROL$COL)
# plot(TRANS.CONTROL$YFP.SD.CORRECT ~ TRANS.CONTROL$POSITION)
# plot(TRANS.CONTROL$YFP.SD.CORRECT ~ TRANS.CONTROL$PLATE)
# plot(TRANS.CONTROL$YFP.SD.CORRECT ~ TRANS.CONTROL$REP)
# plot(TRANS.CONTROL$YFP.SD.CORRECT ~ TRANS.CONTROL$DA

write.table(TRANS.DATA,"DATA.CORRECT.txt",row.names=FALSE,sep="\t",quote=FALSE)


### d) Expression for each sample. ###

TRANS.DATA <- read.table("DATA.CORRECT.txt",header=TRUE)

TRANS.DATA <- subset(TRANS.DATA, COUNTS.FINAL > 500)

#Check for duplicate files
QC <- TRANS.DATA

for (i in 1:nrow(QC))
{
  QC[,"Duplicate"] <- length(which(QC[,"COUNTS.INITIAL"] == QC[i,"COUNTS.INITIAL"] & QC[,"COUNTS.FINAL"] == QC[i,"COUNTS.FINAL"]))
}

length(which(QC$Duplicate == 2))

#Remove samples with extremely high FSC MAD (indicating bacterial contamination)

# plot(FILTER.DATA$FSC.MAD.INITIAL,FILTER.DATA$FSC.MEDIAN.CORRECT,pch=20)
TRANS.DATA <- subset(TRANS.DATA, FSC.MAD.INITIAL < 0.21)

#Determine appropriate control and null
TRANS.DATA[,"REFERENCE"] <- 172
TRANS.DATA[,"NEGATIVE"] <- 185

#Calculate Expression Relative to WT
for (i in 1:nrow(TRANS.DATA))
{
  CUR.WT <- subset(TRANS.DATA, ID.1 == TRANS.DATA[i,"REFERENCE"] & EXPERIMENT == TRANS.DATA[i,"EXPERIMENT"])
  CUR.NEG <- subset(TRANS.DATA, ID.1 == TRANS.DATA[i,"NEGATIVE"] & EXPERIMENT == TRANS.DATA[i,"EXPERIMENT"])
  
  WT.MEAN.ADJUST <- mean(CUR.WT[,"YFP.MEDIAN.CORRECT"]- mean(CUR.NEG[,"YFP.MEDIAN.CORRECT"]))
  WT.SD.ADJUST <- mean(CUR.WT[,"YFP.SD.CORRECT"]- mean(CUR.NEG[,"YFP.SD.CORRECT"]))
  
  TRANS.DATA[i,"YFP.MEDIAN.ADJUST"] <- TRANS.DATA[i,"YFP.MEDIAN.CORRECT"] - mean(CUR.NEG[,"YFP.MEDIAN.CORRECT"])
  TRANS.DATA[i,"YFP.SD.ADJUST"] <- TRANS.DATA[i,"YFP.SD.CORRECT"] - mean(CUR.NEG[,"YFP.SD.CORRECT"])
  
  TRANS.DATA[i,"FSC.MEDIAN.RELATIVE"] <- TRANS.DATA[i,"FSC.MEDIAN.CORRECT"]/mean(CUR.WT[,"FSC.MEDIAN.CORRECT"])
  TRANS.DATA[i,"YFP.MEDIAN.RELATIVE"] <- TRANS.DATA[i,"YFP.MEDIAN.ADJUST"]/WT.MEAN.ADJUST
  TRANS.DATA[i,"YFP.SD.SCALED"] <- TRANS.DATA[i,"YFP.SD.ADJUST"]/WT.MEAN.ADJUST
  TRANS.DATA[i,"YFP.SD.RELATIVE"] <- TRANS.DATA[i,"YFP.SD.ADJUST"]/WT.SD.ADJUST
  
  TRANS.DATA[i,"YFP.CV"] <- TRANS.DATA[i,"YFP.SD.SCALED"]/TRANS.DATA[i,"YFP.MEDIAN.RELATIVE"]
  TRANS.DATA[i,"YFP.CV.RELATIVE"] <- TRANS.DATA[i,"YFP.SD.RELATIVE"]/TRANS.DATA[i,"YFP.MEDIAN.RELATIVE"]
}

write.table(TRANS.DATA,"ALL.DATA.txt",sep="\t",quote=FALSE,row.names=FALSE)


### e) Remove outliers. ###

#BASED ON YFP MEDIAN
for (i in 1:nrow(TRANS.DATA))
{
  CUR <- subset(TRANS.DATA, ID.1 == TRANS.DATA[i,"ID.1"] & CLONE == TRANS.DATA[i,"CLONE"] & EXPERIMENT == TRANS.DATA[i,"EXPERIMENT"])
  
  LOW <- median(CUR$YFP.MEDIAN.RELATIVE) - 2.56*mad(CUR$YFP.MEDIAN.RELATIVE)
  HIGH <- median(CUR$YFP.MEDIAN.RELATIVE) + 2.56*mad(CUR$YFP.MEDIAN.RELATIVE)
  
  if (TRANS.DATA[i,"YFP.MEDIAN.RELATIVE"] > LOW & TRANS.DATA[i,"YFP.MEDIAN.RELATIVE"] < HIGH)
  {
    TRANS.DATA[i,"YFP.OUTLIER"] <- "NO"
  } else {
    TRANS.DATA[i,"YFP.OUTLIER"] <- "YES"
  }
}

FILTER.DATA <- subset(TRANS.DATA, YFP.OUTLIER == "NO")

FILTER.DATA[,"REFERENCE"] <- as.factor(FILTER.DATA[,"REFERENCE"])
FILTER.DATA[,"NEGATIVE"] <- as.factor(FILTER.DATA[,"NEGATIVE"])

write.table(FILTER.DATA,"FILTER.DATA.txt",sep="\t",quote=FALSE,row.names=FALSE)


### f) Calculate median expression and noise for each strain. ###

#Replace Manually all NAs with NAN.
FILTER.DATA <- read.table("FILTER.DATA.txt",header=TRUE)

TRANS.MEDIAN <- aggregate(cbind(FSC.MEDIAN.RELATIVE, YFP.MEDIAN.ADJUST, YFP.MEDIAN.RELATIVE, YFP.SD.SCALED, YFP.CV, YFP.SD.RELATIVE, YFP.CV.RELATIVE) ~ ID.1 + STRAIN.1 + EXPERIMENT + TYPE  + MUTATION + CLONE + EMS.COLLECTION + EMS.MUTANT + GENE + LINKAGE.GROUP + CAUSATIVE + MATING.TYPE + REFERENCE + NEGATIVE, data=FILTER.DATA, FUN = median, na.rm=TRUE)
TRANS.SD <- aggregate(cbind(FSC.MEDIAN.RELATIVE, YFP.MEDIAN.ADJUST, YFP.MEDIAN.RELATIVE, YFP.SD.SCALED, YFP.CV, YFP.SD.RELATIVE, YFP.CV.RELATIVE) ~ ID.1 + STRAIN.1 + EXPERIMENT + TYPE  + MUTATION + CLONE + EMS.COLLECTION + EMS.MUTANT + GENE + LINKAGE.GROUP + CAUSATIVE + MATING.TYPE + REFERENCE + NEGATIVE, data=FILTER.DATA, FUN = sd, na.rm=TRUE)
TRANS.N <- aggregate(cbind(FSC.MEDIAN.RELATIVE, YFP.MEDIAN.ADJUST, YFP.MEDIAN.RELATIVE, YFP.SD.SCALED, YFP.CV, YFP.SD.RELATIVE, YFP.CV.RELATIVE) ~ ID.1 + STRAIN.1 + EXPERIMENT + TYPE  + MUTATION + CLONE + EMS.COLLECTION + EMS.MUTANT + GENE + LINKAGE.GROUP + CAUSATIVE + MATING.TYPE + REFERENCE + NEGATIVE, data=FILTER.DATA, FUN = length)

TRANS.MEDIAN <- TRANS.MEDIAN[which(TRANS.N[,"YFP.MEDIAN.RELATIVE"] >= 2),]
TRANS.SD <- TRANS.SD[which(TRANS.N[,"YFP.MEDIAN.RELATIVE"] >= 2),]
TRANS.N <- TRANS.N[which(TRANS.N[,"YFP.MEDIAN.RELATIVE"] >= 2),]

TRANS <- cbind.data.frame(TRANS.MEDIAN,TRANS.SD[,15:21],TRANS.N[,21])
colnames(TRANS) <- c("ID","STRAIN","EXPERIMENT","TYPE","MUTATION","CLONE","EMS.COLLECTION","EMS.MUTANT","GENE","LINKAGE.GROUP","CAUSATIVE","MATING.TYPE","REFERENCE","NEGATIVE","FSC.MEDIAN.RELATIVE.MEAN","YFP.MEDIAN.ADJUST.MEAN","YFP.MEDIAN.RELATIVE.MEAN","YFP.SD.SCALED.MEAN","YFP.CV.MEAN","YFP.SD.RELATIVE.MEAN","YFP.CV.RELATIVE.MEAN","FSC.MEDIAN.RELATIVE.SD","YFP.MEDIAN.ADJUST.SD","YFP.MEDIAN.RELATIVE.SD","YFP.SD.SCALED.SD","YFP.CV.SD","YFP.SD.RELATIVE.SD","YFP.CV.RELATIVE.SD","N")

write.table(TRANS,"SUMMARY.DATA.txt",sep="\t",quote=FALSE,row.names=FALSE)


###################################################################################
# 7 - Permutation tests used to compare the expression of each single-site mutant #                 
#     to the expression of the progenitor strain.                                 #
###################################################################################

#The input file for this script is the output of the previous section.
TRANS <- read.table("SUMMARY.DATA.txt",header=TRUE,colClasses=DATA.TYPE)

###A-Compare Median expression of each sample to WT###

#Permutation function
SHUFFLE <- function(STRAIN.LIST = STRAIN.LIST[[i]],WT.LIST = WT.LIST) {
  SAMPLE <- c(STRAIN.LIST,WT.LIST[[sample(1:length(WT.LIST),1)]])
  OUTPUT <- sample(SAMPLE,length(SAMPLE))
  DIST.SIM <- abs(mean(OUTPUT[1:length(STRAIN.LIST)]) - mean(OUTPUT[(length(STRAIN.LIST)+1):length(SAMPLE)]))
  DIST.OBS <- abs(mean(SAMPLE[1:length(STRAIN.LIST)]) - mean(SAMPLE[(length(STRAIN.LIST)+1):length(SAMPLE)]))
  
  DIFF <- DIST.OBS - DIST.SIM
  
  return(DIFF)
}


for (i in 1:nrow(TRANS))
{
  if (TRANS[i,"REFERENCE"] == 172)
  {
    #Create list of WT
    WT <- subset(FILTER.DATA, ID.1 == 172 & EXPERIMENT == TRANS[i,"EXPERIMENT"])
    N.WT <- length(unique(WT[,"POSITION"]))
    WT.LIST <- vector('list',N.WT)
    
    for (j in 1:N.WT)
    {
      CUR.WT <- subset(WT, POSITION == unique(WT[,"POSITION"])[j])
      WT.LIST[[j]] <- CUR.WT$YFP.MEDIAN.RELATIVE
    }
    
    #Create list of samples
    CUR.SAMPLE <- subset(FILTER.DATA, ID.1 == TRANS[i,"ID"] & CLONE == TRANS[i,"CLONE"] & EXPERIMENT == TRANS[i,"EXPERIMENT"])
    STRAIN.LIST <- vector("list",1)
    STRAIN.LIST[[1]] <- CUR.SAMPLE$YFP.MEDIAN.RELATIVE
    
    #Permutation test
    N.PERM <- 10000
    
    DIFF <- sapply(1:N.PERM,FUN=function(x)	SHUFFLE(STRAIN.LIST = STRAIN.LIST[[1]],WT.LIST = WT.LIST))
    P.VAL <- (length(which(DIFF < 0)) + 1) / (length(which(DIFF < 0)) + length(which(DIFF > 0)) + 1) 	
    TRANS[i,"P.MEDIAN.WT"] <- P.VAL 
  }
  
  if (TRANS[i,"REFERENCE"] == 170)
  {
    #Create list of WT
    WT <- subset(FILTER.DATA, ID.1 == 170 & EXPERIMENT == TRANS[i,"EXPERIMENT"])
    N.WT <- length(unique(WT[,"POSITION"]))
    WT.LIST <- vector('list',N.WT)
    
    for (j in 1:N.WT)
    {
      CUR.WT <- subset(WT, POSITION == unique(WT[,"POSITION"])[j])
      WT.LIST[[j]] <- CUR.WT$YFP.MEDIAN.RELATIVE
    }
    
    #Create list of samples
    CUR.SAMPLE <- subset(FILTER.DATA, ID.1 == TRANS[i,"ID"] & CLONE == TRANS[i,"CLONE"] & EXPERIMENT == TRANS[i,"EXPERIMENT"])
    STRAIN.LIST <- vector("list",1)
    STRAIN.LIST[[1]] <- CUR.SAMPLE$YFP.MEDIAN.RELATIVE
    
    #Permutation test
    N.PERM <- 10000
    
    DIFF <- sapply(1:N.PERM,FUN=function(x)	SHUFFLE(STRAIN.LIST = STRAIN.LIST[[1]],WT.LIST = WT.LIST))
    P.VAL <- (length(which(DIFF < 0)) + 1) / (length(which(DIFF < 0)) + length(which(DIFF > 0)) + 1) 	
    TRANS[i,"P.MEDIAN.WT"] <- P.VAL 
  }
  
  if (TRANS[i,"REFERENCE"] == 182)
  {
    #Create list of WT
    WT <- subset(FILTER.DATA, ID.1 == 182 & EXPERIMENT == TRANS[i,"EXPERIMENT"])
    N.WT <- length(unique(WT[,"POSITION"]))
    WT.LIST <- vector('list',N.WT)
    
    for (j in 1:N.WT)
    {
      CUR.WT <- subset(WT, POSITION == unique(WT[,"POSITION"])[j])
      WT.LIST[[j]] <- CUR.WT$YFP.MEDIAN.RELATIVE
    }
    
    #Create list of samples
    CUR.SAMPLE <- subset(FILTER.DATA, ID.1 == TRANS[i,"ID"] & CLONE == TRANS[i,"CLONE"] & EXPERIMENT == TRANS[i,"EXPERIMENT"])
    STRAIN.LIST <- vector("list",1)
    STRAIN.LIST[[1]] <- CUR.SAMPLE$YFP.MEDIAN.RELATIVE
    
    #Permutation test
    N.PERM <- 10000
    
    DIFF <- sapply(1:N.PERM,FUN=function(x)	SHUFFLE(STRAIN.LIST = STRAIN.LIST[[1]],WT.LIST = WT.LIST))
    P.VAL <- (length(which(DIFF < 0)) + 1) / (length(which(DIFF < 0)) + length(which(DIFF > 0)) + 1) 	
    TRANS[i,"P.MEDIAN.WT"] <- P.VAL 
  }
  
  print(round(i/nrow(TRANS)*100,1))
}

TRANS[,"P.MEDIAN.WT"] <- p.adjust(TRANS[,"P.MEDIAN.WT"], method="fdr")

###B-Compare CV expression of each sample to WT###
for (i in 1:nrow(TRANS))
{
  if (TRANS[i,"REFERENCE"] == 172)
  {
    #Create list of WT
    WT <- subset(FILTER.DATA, ID.1 == 172 & EXPERIMENT == TRANS[i,"EXPERIMENT"])
    N.WT <- length(unique(WT[,"POSITION"]))
    WT.LIST <- vector('list',N.WT)
    
    for (j in 1:N.WT)
    {
      CUR.WT <- subset(WT, POSITION == unique(WT[,"POSITION"])[j])
      WT.LIST[[j]] <- CUR.WT$YFP.CV.RELATIVE
    }
    
    #Create list of samples
    CUR.SAMPLE <- subset(FILTER.DATA, ID.1 == TRANS[i,"ID"] & CLONE == TRANS[i,"CLONE"] & EXPERIMENT == TRANS[i,"EXPERIMENT"])
    STRAIN.LIST <- vector("list",1)
    STRAIN.LIST[[1]] <- CUR.SAMPLE$YFP.CV.RELATIVE
    
    #Permutation test
    N.PERM <- 10000
    
    DIFF <- sapply(1:N.PERM,FUN=function(x)	SHUFFLE(STRAIN.LIST = STRAIN.LIST[[1]],WT.LIST = WT.LIST))
    P.VAL <- (length(which(DIFF < 0)) + 1) / (length(which(DIFF < 0)) + length(which(DIFF > 0)) + 1) 	
    TRANS[i,"P.CV.WT"] <- P.VAL 
  }
  
  if (TRANS[i,"REFERENCE"] == 170)
  {
    #Create list of WT
    WT <- subset(FILTER.DATA, ID.1 == 170 & EXPERIMENT == TRANS[i,"EXPERIMENT"])
    N.WT <- length(unique(WT[,"POSITION"]))
    WT.LIST <- vector('list',N.WT)
    
    for (j in 1:N.WT)
    {
      CUR.WT <- subset(WT, POSITION == unique(WT[,"POSITION"])[j])
      WT.LIST[[j]] <- CUR.WT$YFP.CV.RELATIVE
    }
    
    #Create list of samples
    CUR.SAMPLE <- subset(FILTER.DATA, ID.1 == TRANS[i,"ID"] & CLONE == TRANS[i,"CLONE"] & EXPERIMENT == TRANS[i,"EXPERIMENT"])
    STRAIN.LIST <- vector("list",1)
    STRAIN.LIST[[1]] <- CUR.SAMPLE$YFP.CV.RELATIVE
    
    #Permutation test
    N.PERM <- 10000
    
    DIFF <- sapply(1:N.PERM,FUN=function(x)	SHUFFLE(STRAIN.LIST = STRAIN.LIST[[1]],WT.LIST = WT.LIST))
    P.VAL <- (length(which(DIFF < 0)) + 1) / (length(which(DIFF < 0)) + length(which(DIFF > 0)) + 1) 	
    TRANS[i,"P.CV.WT"] <- P.VAL 
  }
  
  if (TRANS[i,"REFERENCE"] == 182)
  {
    #Create list of WT
    WT <- subset(FILTER.DATA, ID.1 == 182 & EXPERIMENT == TRANS[i,"EXPERIMENT"])
    N.WT <- length(unique(WT[,"POSITION"]))
    WT.LIST <- vector('list',N.WT)
    
    for (j in 1:N.WT)
    {
      CUR.WT <- subset(WT, POSITION == unique(WT[,"POSITION"])[j])
      WT.LIST[[j]] <- CUR.WT$YFP.CV.RELATIVE
    }
    
    #Create list of samples
    CUR.SAMPLE <- subset(FILTER.DATA, ID.1 == TRANS[i,"ID"] & CLONE == TRANS[i,"CLONE"] & EXPERIMENT == TRANS[i,"EXPERIMENT"])
    STRAIN.LIST <- vector("list",1)
    STRAIN.LIST[[1]] <- CUR.SAMPLE$YFP.CV.RELATIVE
    
    #Permutation test
    N.PERM <- 10000
    
    DIFF <- sapply(1:N.PERM,FUN=function(x)	SHUFFLE(STRAIN.LIST = STRAIN.LIST[[1]],WT.LIST = WT.LIST))
    P.VAL <- (length(which(DIFF < 0)) + 1) / (length(which(DIFF < 0)) + length(which(DIFF > 0)) + 1) 	
    TRANS[i,"P.CV.WT"] <- P.VAL 
  }
  
  print(round(i/nrow(TRANS)*100,1))
}

TRANS[,"P.CV.WT"] <- p.adjust(TRANS[,"P.CV.WT"], method="fdr")

###C-Compare Median expression for Experiments A and B.###
for (i in 1:nrow(TRANS))
{
  EXP.A <- subset(FILTER.DATA, ID.1 == TRANS[i,"ID"] & CLONE == TRANS[i,"CLONE"] & EXPERIMENT == "A")
  EXP.B <- subset(FILTER.DATA, ID.1 == TRANS[i,"ID"] & CLONE == TRANS[i,"CLONE"] & EXPERIMENT == "B")
  
  if (nrow(EXP.A) > 1 & nrow(EXP.B) > 1)
  {
    TRANS[i,"P.MEDIAN.EXP"] <- t.test(EXP.A[,"YFP.MEDIAN.RELATIVE"], EXP.B[,"YFP.MEDIAN.RELATIVE"])$p.value
  }
}

TRANS[,"P.MEDIAN.EXP"] <- p.adjust(TRANS[,"P.MEDIAN.EXP"], method="fdr")

###D-Compare CV expression for Experiments A and B.###
for (i in 1:nrow(TRANS))
{
  EXP.A <- subset(FILTER.DATA, ID.1 == TRANS[i,"ID"] & CLONE == TRANS[i,"CLONE"] & EXPERIMENT == "A")
  EXP.B <- subset(FILTER.DATA, ID.1 == TRANS[i,"ID"] & CLONE == TRANS[i,"CLONE"] & EXPERIMENT == "B")
  
  if (nrow(EXP.A) > 1 & nrow(EXP.B) > 1)
  {
    TRANS[i,"P.CV.EXP"] <- t.test(EXP.A[,"YFP.CV.RELATIVE"], EXP.B[,"YFP.CV.RELATIVE"])$p.value
  }
}

TRANS[,"P.CV.EXP"] <- p.adjust(TRANS[,"P.CV.EXP"], method="fdr")

###E-Compare Median Clone 1 with Median Clone 2.###
for (i in 1:nrow(TRANS))
{
  CLONE.1 <- subset(FILTER.DATA, MUTATION == TRANS[i,"MUTATION"] & CLONE == 1 & EXPERIMENT == TRANS[i,"EXPERIMENT"])
  CLONE.2 <- subset(FILTER.DATA, MUTATION == TRANS[i,"MUTATION"] & CLONE == 2 & EXPERIMENT == TRANS[i,"EXPERIMENT"])
  
  if (nrow(CLONE.1) > 1 & nrow(CLONE.2) > 1)
  {
    TRANS[i,"P.MEDIAN.CLONES"] <- t.test(CLONE.1[,"YFP.MEDIAN.RELATIVE"], CLONE.2[,"YFP.MEDIAN.RELATIVE"])$p.value
  }
}

TRANS[,"P.MEDIAN.CLONES"] <- p.adjust(TRANS[,"P.MEDIAN.CLONES"], method="fdr")

###F-Compare Median Expression of Single Site mutants to EMS mutants.###

###Using t-tests###
for (i in 1:nrow(TRANS))
{
  SAMPLE <- subset(FILTER.DATA, ID.1 == TRANS[i,"ID"] & CLONE == TRANS[i,"CLONE"] & EXPERIMENT == TRANS[i,"EXPERIMENT"])
  EMS <- subset(FILTER.DATA, TYPE == "EMS.MUTANT" & EMS.MUTANT == TRANS[i,"EMS.MUTANT"] & EXPERIMENT == TRANS[i,"EXPERIMENT"])
  
  if (nrow(SAMPLE) > 1 & nrow(EMS) > 1 & TRANS[i,"TYPE"] != "EMS.MUTANT" & TRANS[i,"EMS.MUTANT"] != "NAN")
  {
    TRANS[i,"P.MEDIAN.EMS"] <- t.test(SAMPLE[,"YFP.MEDIAN.RELATIVE"], EMS[,"YFP.MEDIAN.RELATIVE"])$p.value
  }
}

TRANS[,"P.MEDIAN.EMS"] <- p.adjust(TRANS[,"P.MEDIAN.EMS"], method="fdr")

#"SourceData7.txt" can be found in SupplementaryFile11.zip.
write.table(TRANS,"SourceData7.txt",sep="\t",quote=FALSE,row.names=FALSE)


##################################################################
# 8 - Expression changes caused by linked mutations (Figure 2E). #
##################################################################

#Input files can be found in SupplementaryFile11.zip.
#These files were generated with R scripts above (sections 1, 2 and 6) from the experiment described in "TEMPLATE.SINGLE.SITE.MUTANTS.txt".
TRANS <- read.table("SourceData7.txt",header=TRUE)
FILTER.DATA <- read.table("SourceData8.txt",header=TRUE)

EMS.MUTANTS <- c("2P15D07","2P16H09","2P19F12","2P03B06","Y54")

LINK.TRANS <- subset(TRANS, EMS.MUTANT %in% EMS.MUTANTS | ID == 172)

LINK.TRANS[,"MUTATION"] <- factor(LINK.TRANS[,"MUTATION"],levels=c("WT","2P03B06","ATP23_G476A","COG6_G640A","2P15D07","CCC2_G2159A","EXG2_C379T","2P16H09","CIA2_G626A","RPF1_G127A","2P19F12","MOD5_G765A","DSE3_G249A","Y54","CHD1_G1277A_G2723A","CHD1_G1277A","CHD1_G2723A","SET1_TTATCGAGTACGTTGGTGA2890del"))

EXP.A <- subset(LINK.TRANS, EXPERIMENT == "A")
EXP.B <- subset(LINK.TRANS, EXPERIMENT == "B")

LINK.TRANS <- EXP.A
LINK.TRANS[1,] <- EXP.B[1,]

LINK.TRANS <- subset(LINK.TRANS, TYPE != "OFF.TARGET" & MUTATION != "CHD1_G2723A")
LINK.TRANS <- subset(LINK.TRANS, CLONE == 1 | MUTATION %in% c("DSE3_G249A","MOD5_G765A"))

LINK.TRANS <- droplevels(LINK.TRANS)
LINK.TRANS <- LINK.TRANS[order(LINK.TRANS[,"MUTATION"]),]

LINK.TRANS[,"POSITION"] <- c(1,3,4,5,7,8,9,11,12,13,15,16,17,19,20,21)

LINK.TRANS[,"UNIQUE.ID"] <- paste(LINK.TRANS$ID,LINK.TRANS$EXPERIMENT,LINK.TRANS$MUTATION,LINK.TRANS$CLONE,sep="_")
FILTER.DATA[,"UNIQUE.ID"] <- paste(FILTER.DATA$ID.1,FILTER.DATA$EXPERIMENT,FILTER.DATA$MUTATION,FILTER.DATA$CLONE,sep="_")

GOOD.DATA <- subset(FILTER.DATA, UNIQUE.ID %in% LINK.TRANS$UNIQUE.ID)

for (i in 1:nrow(GOOD.DATA))
{
  CUR.TRANS <- subset(LINK.TRANS, UNIQUE.ID == GOOD.DATA[i,"UNIQUE.ID"])
  
  GOOD.DATA[i,"PLOT.POSITION"] <- CUR.TRANS[1,"POSITION"]
  GOOD.DATA[i,"RAND.POSITION"] <- runif(1,GOOD.DATA[i,"PLOT.POSITION"]-0.25,GOOD.DATA[i,"PLOT.POSITION"]+0.25)
  
  if (GOOD.DATA[i,"MUTATION"] == "WT")
  {
    GOOD.DATA[i,"COLOR"] <- "#00000066"
  }
  if (GOOD.DATA[i,"TYPE"] == "EMS.MUTANT")
  {
    GOOD.DATA[i,"COLOR"] <- "#8D614299"
  }
  if (GOOD.DATA[i,"MUTATION"] %in% c("ATP23_G476A","CCC2_G2159A","CIA2_G626A","MOD5_G765A","CHD1_G1277A_G2723A"))
  {
    GOOD.DATA[i,"COLOR"] <- "#008080AA"
  }
  if (GOOD.DATA[i,"MUTATION"] %in% c("COG6_G640A","EXG2_C379T","RPF1_G127A","DSE3_G249A","CHD1_G1277A"))
  {
    GOOD.DATA[i,"COLOR"] <- "#80CDDBAA"
  }
}

pdf("Figure2E.pdf",width=7,height=6,useDingbats = F)
#windows(height=6,width=7)
plot(LINK.TRANS$POSITION,LINK.TRANS$YFP.MEDIAN.RELATIVE.MEAN,ylim=c(0.75,1.1),pch="-",cex=2,ylab="Median expression",xlab="",xaxt="n")
axis(1,at=c(1,3,4,5,7,8,9,11,12,13,15,16,17,19,20,21))
points(GOOD.DATA$PLOT.POSITION,GOOD.DATA$YFP.MEDIAN.RELATIVE,pch=16,col=GOOD.DATA$COLOR,cex=2)
abline(v=c(2,6,10,14,18),lty=1,col="#00000033")
abline(h=1,lty=2)
dev.off()


##########################################################################################
# 9 - Relationship between expression changes in single-site mutants and signed G-values #
#     of corresponding mutations (Figure 2F).                                            #
##########################################################################################

#Clear memory
rm(list=ls())
options(warn=-1)

###Load packages###
library(VariantAnnotation)
library(Deducer)
library(gtools)
library(zoo)
library(Hmisc)
library(ggplot2)
library(plotrix)
library(moments)
library(vcd)

###Set directory###
setwd("/Path.to.input.files")

#Load table describing all mutations that can be found in SuppplementaryFile11.zip.
ALL.MUT <- read.table("SourceData1.txt",header=TRUE)

#Load expression data for single site mutants generated with R scripts described above (sections 1, 2, 6 and 7).
#The template file used for this experiment was "TEMPLATE.SINGLE.SITE.MUTANTS.txt".
#These data files can be found in SupplementaryFile11.zip.
SS.DATA <- read.table("SourceData8.txt",header=TRUE)
TRANS <- read.table("SourceData7.txt",header=TRUE)

#Calculate number of mutations and effect size for each EMS mutant.
EFFECT.SIZE <- aggregate(cbind(MEDIAN.EXPR.MEAN.EMS.1, MEDIAN.EXPR.SD.EMS.1, MEDIAN.EXPR.N.EMS.1,N.CAUSAL) ~ STRAIN + MUT.RUN, data=ALL.MUT ,FUN=mean)
N.MUTATION <- aggregate(cbind(MEDIAN.EXPR.MEAN.EMS.1, MEDIAN.EXPR.SD.EMS.1, MEDIAN.EXPR.N.EMS.1) ~ STRAIN + MUT.RUN, data=ALL.MUT ,FUN=length)

N.MUTATION <- N.MUTATION[,3]

PLOT <- cbind(EFFECT.SIZE,N.MUTATION)
#PLOT <- subset(PLOT, N.MUTATION > 3)

for (i in 1:nrow(PLOT))
{
  if (PLOT[i,"MEDIAN.EXPR.MEAN.EMS.1"] > 1)
  {
    PLOT[i,"COLOR"] <- "red"
  } else {
    PLOT[i,"COLOR"] <- "blue"
  }
}

PLOT[,"EFFECT.SIZE"] <- PLOT[,"MEDIAN.EXPR.MEAN.EMS.1"]-1
PLOT[,"ABSOLUTE.EFFECT"] <- abs(1-PLOT[,"MEDIAN.EXPR.MEAN.EMS.1"])
PLOT[,"N.CAUSAL"] <- as.factor(PLOT[,"N.CAUSAL"])

ALL.MUTANT <- subset(PLOT, N.CAUSAL != 0)

PLOT <- TRANS

#Number of independent clones analyzed for each single-site mutant.
for(i in 1:nrow(PLOT))
{
  CUR <- subset(PLOT, EXPERIMENT == PLOT[i,"EXPERIMENT"] & MUTATION == PLOT[i,"MUTATION"] & REFERENCE == PLOT[i,"REFERENCE"] & MUTATION == PLOT[i,"MUTATION"] & TYPE == PLOT[i,"TYPE"])
  
  PLOT[i,"N.CLONES"] <- length(unique(CUR$CLONE))
}

#Focus on experiment A.
EXP.A <- subset(SS.DATA, EXPERIMENT == "A")

SS <- subset(EXP.A, TYPE %in% c("SINGLE.SITE","OFF.TARGET") & EMS.MUTANT != "NAN")
SS <- subset(SS, CAUSATIVE == "YES")

#Expression data for EMS mutants.
EMS <- cbind.data.frame(as.character(SS$EMS.MUTANT), SS$REFERENCE, SS$YFP.MEDIAN.RELATIVE.MEAN, SS$YFP.CV.RELATIVE.MEAN, SS$YFP.MEDIAN.RELATIVE.SD, SS$YFP.CV.RELATIVE.SD,SS$N)
colnames(EMS) <- c("EMS.MUTANT","REFERENCE","YFP.MEDIAN.RELATIVE.MEAN","YFP.CV.RELATIVE.MEAN","YFP.MEDIAN.RELATIVE.SD","YFP.CV.RELATIVE.SD","N")

LENGTH <- c()

for (i in 1:nrow(EMS))
{
  CUR.EMS <- subset(PLOT, TYPE == "EMS.MUTANT" & EMS.MUTANT == as.character(SS[i,"EMS.MUTANT"]) & EXPERIMENT == SS[i,"EXPERIMENT"])
  
  EMS[i,"REFERENCE"] <- CUR.EMS[1,"REFERENCE"]
  EMS[i,"YFP.MEDIAN.RELATIVE.MEAN"] <- CUR.EMS[1,"YFP.MEDIAN.RELATIVE.MEAN"]
  EMS[i,"YFP.CV.RELATIVE.MEAN"] <- CUR.EMS[1,"YFP.CV.RELATIVE.MEAN"]
  EMS[i,"YFP.MEDIAN.RELATIVE.SD"] <- CUR.EMS[1,"YFP.MEDIAN.RELATIVE.SD"]
  EMS[i,"YFP.CV.RELATIVE.SD"] <- CUR.EMS[1,"YFP.CV.RELATIVE.SD"]
  EMS[i,"N"] <- CUR.EMS[1,"N"]
  
  LENGTH[i] <- nrow(CUR.EMS)
}

#Retrieve information for EMS mutant Y96 from previous dataset.
EMS[47,"REFERENCE"] <- 170
EMS[47,"YFP.MEDIAN.RELATIVE.MEAN"] <- 0.935137547
EMS[47,"YFP.CV.RELATIVE.MEAN"] <- 0.98863486
EMS[47,"YFP.MEDIAN.RELATIVE.SD"] <- 0.02209943
EMS[47,"YFP.CV.RELATIVE.SD"] <- 0.06842578
EMS[47,"N"] <- 6

#Generate matching tables of EMS and single-site mutants carrying the same mutations.
ALL.MUTANT <- subset(ALL.MUTANT, STRAIN %nin% c("Y123","Y99"))
GOOD.STRAINS <- c(intersect(EMS[,"EMS.MUTANT"],ALL.MUTANT[,"STRAIN"]),"1P16A09")

GOOD.MUTANT <- subset(ALL.MUTANT, STRAIN %in% GOOD.STRAINS)
EMS <- subset(EMS, EMS.MUTANT %in% GOOD.STRAINS)
SS <-  subset(SS, EMS.MUTANT %in% GOOD.STRAINS)

#Add info about methods used to identify each mutation (Sanger and BSA-Seq).
for (i in 1:nrow(SS))
{
  CUR.MUT <- subset(ALL.MUT, STRAIN == as.character(SS[i,"EMS.MUTANT"]))
  
  SS[i,"SEQ.RUN"] <- CUR.MUT[1,"SEQ.RUN"]
  
  if (SS[i,"SEQ.RUN"] == "SANGER")
  {
    SS[i,"PCH"] <- 17
  } else {
    SS[i,"PCH"] <- 16
  }
}

#Transform expression values into absolute effect sizes.
SS[,"EFFECT.SIZE"] <- abs(1-SS[,"YFP.MEDIAN.RELATIVE.MEAN"])

#Add signed G-values of each mutation identified by BSA-Seq.
for (i in 1:nrow(SS))
{
  CUR.MUTATION <- subset(ALL.MUT, STRAIN == as.character(SS[i,"EMS.MUTANT"]) & GENE.FOCAL == as.character(SS[i,"GENE"]))
  
  SS[i,"G.value"] <- CUR.MUTATION[1,"G.VALUE"]
  
  if (nrow(CUR.MUTATION) != 0)
  {
    if (!is.na(CUR.MUTATION[,"G.VALUE"]))
    {
      if (CUR.MUTATION[1,"P.SCORE"] < 0)
      {
        SS[i,"G.Sign"] <- -1
      } else {
        SS[i,"G.Sign"] <- 1
      }
    }
  }
}

G.VAL <- SS[,"G.Sign"]*SS[,"G.value"]
EFFECT.SIZE <- SS[,"YFP.MEDIAN.RELATIVE.MEAN"] - 1

#Color of data points shown on Figure2F.
COLOR <- c()

for (i in 1:nrow(SS))
{
  if (SS[i,"P.MEDIAN.WT"] > 0.01)
  {
    COLOR[i] <- "red"
  } else if (G.VAL[i] > 0 & EFFECT.SIZE[i] < 0) {
    COLOR[i] <- "green"
  } else {
    COLOR[i] <- "black"
  }
}

cor.test(G.VAL, EFFECT.SIZE)

#Plotting Figure2F.
pdf("Figure2F.pdf",useDingbats=FALSE,height=10,width=10)
#windows(height=10,width=10)
plotCI(EFFECT.SIZE,G.VAL,ui=SS$YFP.MEDIAN.RELATIVE.MEAN-1+1.96*SS$YFP.MEDIAN.RELATIVE.SD/sqrt(SS$N),li=SS$YFP.MEDIAN.RELATIVE.MEAN-1-1.96*SS$YFP.MEDIAN.RELATIVE.SD/sqrt(SS$N),err="x",col=COLOR,xlab="Expression change in single site mutants",ylab="Signed G-value",pch=16,sfrac=0,xlim=c(-0.5,0.5),ylim=c(-250,250),scol="#00000066",gap=0,cex=1.5)
abline(v=0,lty=2)
abline(h=0,lty=2)
dev.off()


####################################################################################
# 10 - Permutation tests used to compare the expression of each single-site mutant #                 
#      to the expression of the EMS mutant carrying the same mutation.             #
####################################################################################


#Input files can be found in SupplementaryFile11.zip.

#This file was generated with R scripts above (sections 1, 2 and 6) from the experiment described in "TEMPLATE.SINGLE.SITE.MUTANTS.txt".
FILTER.DATA <- read.table("SourceData8.txt",header=TRUE)
FILTER.DATA <- subset(FILTER.DATA, YFP.OUTLIER == "NO")

#This file was generated with R scripts above (sections 1, 2 and 6) from flow data collected in Metzger et al, 2016 
#where the fluorescence of the WT strain was measured in four replicates at many plate positions.
BIG.DATA <- read.table("Expression_WT.txt",header=TRUE)
BIG.DATA <- subset(BIG.DATA, YFP.OUTLIER == "NO")

#Load EMS and single site mutants data
SS.DATA <- read.table("SourceData8.txt",header=TRUE)
TRANS <- read.table("SourceData7.txt",header=TRUE)


#The logic of the statistical tests below is to compare a) the difference of expression levels between a single site mutant
#and its corresponding EMS mutant to b) the difference of expression levels between random pairs of the WT strain at different plate positions.


#A-Compare variance among replicates for small and big screens.

#Create list of WT from small screen.
WT.SMALL <- subset(FILTER.DATA, ID.1 == 172)

WT.NAMES <- paste(WT.SMALL[,"EXPERIMENT"],"_",WT.SMALL[,"POSITION"],sep="")
N.WT <- length(unique(WT.NAMES))
WT.SMALL.LIST <- vector('list',N.WT)

for (j in 1:N.WT)
{
  CUR.WT <- subset(WT.SMALL, EXPERIMENT == strsplit(unique(WT.NAMES)[j],split="_")[[1]][1] & POSITION == strsplit(unique(WT.NAMES)[j],split="_")[[1]][2])
  WT.SMALL.LIST[[j]] <- CUR.WT$YFP.MEDIAN.RELATIVE
}

MEAN.SMALL <- c()
SD.SMALL <- c()

for (i in 1:length(WT.SMALL.LIST))
{
  MEAN.SMALL[i] <- mean(WT.SMALL.LIST[[i]])
  SD.SMALL[i] <- sd(WT.SMALL.LIST[[i]])
}

#Create list of WT from big screen.
WT.BIG <- subset(BIG.DATA, STATUS == "SHAM" & CLASS == "SHAM")
LOW.FSC <- median(WT.BIG$FSC.MEDIAN.CORRECT) - 4*mad(WT.BIG$FSC.MEDIAN.CORRECT)
HIGH.FSC <- median(WT.BIG$FSC.MEDIAN.CORRECT) + 4*mad(WT.BIG$FSC.MEDIAN.CORRECT)
LOW.FLUO <- median(WT.BIG$YFP.MEDIAN.CORRECT) - 4*mad(WT.BIG$YFP.MEDIAN.CORRECT)
HIGH.FLUO <- median(WT.BIG$YFP.MEDIAN.CORRECT) + 4*mad(WT.BIG$YFP.MEDIAN.CORRECT)
WT.BIG <- subset(WT.BIG, FSC.MEDIAN.CORRECT > LOW.FSC & FSC.MEDIAN.CORRECT < HIGH.FSC & YFP.MEDIAN.CORRECT > LOW.FLUO & YFP.MEDIAN.CORRECT < HIGH.FLUO)

WT.NAMES <- paste(WT.BIG[,"PLATE"],"_",WT.BIG[,"POSITION"],sep="")
N.WT <- length(unique(WT.NAMES))
WT.BIG.LIST <- vector('list',N.WT)

for (j in 1:N.WT)
{
  CUR.WT <- subset(WT.BIG, PLATE == strsplit(unique(WT.NAMES)[j],split="_")[[1]][1] & POSITION == strsplit(unique(WT.NAMES)[j],split="_")[[1]][2])
  WT.BIG.LIST[[j]] <- CUR.WT$YFP.MEDIAN.RELATIVE
}

MEAN.BIG <- c()
SD.BIG <- c()

for (i in 1:length(WT.BIG.LIST))
{
  MEAN.BIG[i] <- mean(WT.BIG.LIST[[i]])
  SD.BIG[i] <- sd(WT.BIG.LIST[[i]])
}

t.test(MEAN.SMALL,MEAN.BIG)
t.test(SD.SMALL,SD.BIG)
wilcox.test(MEAN.SMALL,MEAN.BIG)
wilcox.test(SD.SMALL,SD.BIG)


#B-Distribution of differences for WT from big screen.

WT.DIFF <- rep(0,length(WT.BIG.LIST)*(length(WT.BIG.LIST)-1)/2)
COUNT <- 1

for (i in 1:(length(WT.BIG.LIST)-1))
{
  for (j in (i+1):length(WT.BIG.LIST))
  {
    WT.1 <- mean(WT.BIG.LIST[[i]])
    WT.2 <- mean(WT.BIG.LIST[[j]])
    
    WT.DIFF[COUNT] <- abs(WT.1-WT.2)
    COUNT <- COUNT + 1
  }
}  

mean(WT.DIFF)
sd(WT.DIFF)
quantile(WT.DIFF,0.975)

#windows(height=8,width=8)
hist(WT.DIFF,breaks=20)
dev.off()

#C-Comparison of differences between 1) EMS and Single Site mutants and 2) Random pairs of WT positions.

PLOT <- TRANS

for(i in 1:nrow(PLOT))
{
  CUR <- subset(PLOT, EXPERIMENT == PLOT[i,"EXPERIMENT"] & MUTATION == PLOT[i,"MUTATION"] & REFERENCE == PLOT[i,"REFERENCE"] & MUTATION == PLOT[i,"MUTATION"] & TYPE == PLOT[i,"TYPE"])
  
  PLOT[i,"N.CLONES"] <- length(unique(CUR$CLONE))
}

#Experiment A

EXP.A <- subset(SS.DATA, EXPERIMENT == "A")

SS <- subset(EXP.A, TYPE %in% c("SINGLE.SITE","OFF.TARGET") & EMS.MUTANT != "NAN")
SS <- subset(SS, CAUSATIVE == "YES")

EMS <- cbind.data.frame(as.character(SS$EMS.MUTANT), SS$REFERENCE, SS$YFP.MEDIAN.RELATIVE.MEAN, SS$YFP.CV.RELATIVE.MEAN, SS$YFP.MEDIAN.RELATIVE.SD, SS$YFP.CV.RELATIVE.SD,SS$N)
colnames(EMS) <- c("EMS.MUTANT","REFERENCE","YFP.MEDIAN.RELATIVE.MEAN","YFP.CV.RELATIVE.MEAN","YFP.MEDIAN.RELATIVE.SD","YFP.CV.RELATIVE.SD","N")

LENGTH <- c()

for (i in 1:nrow(EMS))
{
  CUR.EMS <- subset(PLOT, TYPE == "EMS.MUTANT" & EMS.MUTANT == as.character(SS[i,"EMS.MUTANT"]) & EXPERIMENT == SS[i,"EXPERIMENT"])
  
  EMS[i,"REFERENCE"] <- CUR.EMS[1,"REFERENCE"]
  EMS[i,"YFP.MEDIAN.RELATIVE.MEAN"] <- CUR.EMS[1,"YFP.MEDIAN.RELATIVE.MEAN"]
  EMS[i,"YFP.CV.RELATIVE.MEAN"] <- CUR.EMS[1,"YFP.CV.RELATIVE.MEAN"]
  EMS[i,"YFP.MEDIAN.RELATIVE.SD"] <- CUR.EMS[1,"YFP.MEDIAN.RELATIVE.SD"]
  EMS[i,"YFP.CV.RELATIVE.SD"] <- CUR.EMS[1,"YFP.CV.RELATIVE.SD"]
  EMS[i,"N"] <- CUR.EMS[1,"N"]
  
  LENGTH[i] <- nrow(CUR.EMS)
}

#Retrieve information for EMS mutant Y96 from previous dataset.
EMS[47,"REFERENCE"] <- 170
EMS[47,"YFP.MEDIAN.RELATIVE.MEAN"] <- 0.935137547
EMS[47,"YFP.CV.RELATIVE.MEAN"] <- 0.98863486
EMS[47,"YFP.MEDIAN.RELATIVE.SD"] <- 0.02209943
EMS[47,"YFP.CV.RELATIVE.SD"] <- 0.06842578
EMS[47,"N"] <- 6

#Median expression
ALL.MUTANT <- subset(ALL.MUTANT, STRAIN %nin% c("Y123","Y99"))
GOOD.STRAINS <- intersect(EMS[,"EMS.MUTANT"],ALL.MUTANT[,"STRAIN"])

GOOD.MUTANT <- subset(ALL.MUTANT, STRAIN %in% GOOD.STRAINS)
EMS <- subset(EMS, EMS.MUTANT %in% GOOD.STRAINS)
SS <-  subset(SS, EMS.MUTANT %in% GOOD.STRAINS)

#Permutation tests
N.REP <- 1e5

for (i in 1:nrow(SS))
{
  CUR.EMS <- subset(FILTER.DATA, TYPE == "EMS.MUTANT" & EMS.MUTANT == as.character(SS[i,"EMS.MUTANT"]) & EXPERIMENT == SS[i,"EXPERIMENT"])
  CUR.SS <- subset(FILTER.DATA, TYPE != "EMS.MUTANT" & MUTATION == as.character(SS[i,"MUTATION"]) & EMS.MUTANT == as.character(SS[i,"EMS.MUTANT"]) & EXPERIMENT == SS[i,"EXPERIMENT"])
  
  CUR.DIFF <- abs(mean(CUR.EMS$YFP.MEDIAN.RELATIVE)-mean(CUR.SS$YFP.MEDIAN.RELATIVE))
  
  RANDOM.DIFF <- rep(0,N.REP)
  
  for (j in 1:N.REP)
  {
    WT.POS <- sample(1:length(WT.BIG.LIST),2,replace=FALSE)
    
    WT.1 <- WT.BIG.LIST[[WT.POS[1]]]
    WT.2 <- WT.BIG.LIST[[WT.POS[2]]]
    
    WT.DIFF <- abs(mean(WT.1)-mean(WT.2))
    
    OBS.DIFF <- CUR.DIFF - WT.DIFF
    
    #Adjust WT values
    MUTANT.MEAN <- mean(c(CUR.EMS$YFP.MEDIAN.RELATIVE,CUR.SS$YFP.MEDIAN.RELATIVE))
    WT.MEAN <- mean(c(WT.1,WT.2))
    WT.1 <- WT.1 - WT.MEAN + MUTANT.MEAN
    WT.2 <- WT.2 - WT.MEAN + MUTANT.MEAN
    
    #Shuffle
    COMBINED.1 <- c(CUR.EMS$YFP.MEDIAN.RELATIVE,WT.1)
    COMBINED.2 <- c(CUR.SS$YFP.MEDIAN.RELATIVE,WT.2)
    
    COMBINED.1 <- permute(COMBINED.1)
    COMBINED.2 <- permute(COMBINED.2)
    
    SHUFFLE.EMS <- COMBINED.1[1:nrow(CUR.EMS)]
    SHUFFLE.WT.1 <- COMBINED.1[(nrow(CUR.EMS)+1):length(COMBINED.1)]
    SHUFFLE.SS <- COMBINED.2[1:nrow(CUR.SS)]
    SHUFFLE.WT.2 <- COMBINED.2[(nrow(CUR.SS)+1):length(COMBINED.2)]
    
    SHUFFLE.MUTANT.DIFF <- abs(mean(SHUFFLE.EMS)-mean(SHUFFLE.SS))
    SHUFFLE.WT.DIFF <- abs(mean(SHUFFLE.WT.1)-mean(SHUFFLE.WT.2))
    
    SHUFFLE.DIFF <- SHUFFLE.MUTANT.DIFF - SHUFFLE.WT.DIFF
    
    RANDOM.DIFF[j] <- OBS.DIFF - SHUFFLE.DIFF
  }
  
  P.VAL <- (length(which(RANDOM.DIFF < 0))+1) / ((length(which(RANDOM.DIFF > 0)) + length(which(RANDOM.DIFF < 0))) + 1)
  SS[i,"P.MEDIAN.EMS.2"] <- P.VAL
}

P.VAL.ADJUST <- p.adjust(SS[,"P.MEDIAN.EMS.2"],method="fdr")
SS[,"P.MEDIAN.EMS.2"] <- P.VAL.ADJUST

#"SourceData9.txt" can be found in SupplementaryFile11.zip.
write.table(SS,"SourceData9.txt",sep="\t",quote=FALSE,row.names=FALSE)


#################################################################################################
# 11- Relationship between expression levels of single-site mutants and EMS mutants (Figure 2G). #
#################################################################################################

#SS and EMS variables were generated in section 9.
SS <- SS[-33,]
EMS <- EMS[-33,]

#Add info about methods used to identify each mutation (Sanger and BSA-Seq).
for (i in 1:nrow(SS))
{
  CUR.MUT <- subset(ALL.MUT, STRAIN == as.character(SS[i,"EMS.MUTANT"]))
  
  SS[i,"SEQ.RUN"] <- CUR.MUT[1,"SEQ.RUN"]
  
  if (SS[i,"SEQ.RUN"] == "SANGER")
  {
    SS[i,"PCH"] <- 17
  } else {
    SS[i,"PCH"] <- 16
  }
}

#Colors of dots on Figure 2G.
COLOR <- c()

for(i in 1:nrow(SS))
{
  if (SS[i,"P.MEDIAN.EMS.2"] < 0.01)
  {
    COLOR[i] <- "red"
  } else if (SS[i,"P.MEDIAN.EMS.2"] >= 0.01 & SS[i,"P.MEDIAN.EMS.2"] < 0.05) {
    COLOR[i] <- "blue"
  } else {
    COLOR[i] <- "black"
  }
}

#Correlation of expression between single site mutants and EMS mutants carrying the corresponding mutations.
cor.test(SS$YFP.MEDIAN.RELATIVE.MEAN, EMS$YFP.MEDIAN.RELATIVE.MEAN)
summary(lm(SS$YFP.MEDIAN.RELATIVE.MEAN ~ EMS$YFP.MEDIAN.RELATIVE.MEAN))

#Plotting Figure 2G.
pdf("Figure3.pdf",useDingbats=FALSE,width=8,height=8)
#windows(height=8,width=8)
plotCI(EMS$YFP.MEDIAN.RELATIVE.MEAN,SS$YFP.MEDIAN.RELATIVE.MEAN,ui=EMS$YFP.MEDIAN.RELATIVE.MEAN+1.96*EMS$YFP.MEDIAN.RELATIVE.SD/sqrt(EMS$N),li=EMS$YFP.MEDIAN.RELATIVE.MEAN-1.96*EMS$YFP.MEDIAN.RELATIVE.SD/sqrt(EMS$N),err="x",col=COLOR,xlab="Median Expression EMS mutant",ylab="Median Expression Single Site mutant",pch=SS[,"PCH"],sfrac=0,xlim=c(0.6,1.6),ylim=c(0.6,1.6),scol="#00000066",gap=0,main="Clones pooled",cex=1.5)
plotCI(EMS$YFP.MEDIAN.RELATIVE.MEAN,SS$YFP.MEDIAN.RELATIVE.MEAN,ui=SS$YFP.MEDIAN.RELATIVE.MEAN+1.96*SS$YFP.MEDIAN.RELATIVE.SD/sqrt(SS$N),li=SS$YFP.MEDIAN.RELATIVE.MEAN-1.96*SS$YFP.MEDIAN.RELATIVE.SD/sqrt(SS$N),err="y",col=COLOR,pch=SS[,"PCH"],sfrac=0,scol="#00000066",gap=0,add=TRUE,cex=1.5)
abline(h=1,lty=2,col="#00000099")
abline(v=1,lty=2,col="#00000099")
abline(a=0,b=1,col="#00000066")
abline(a=0.02607788 ,b=1)
abline(a=-0.02607788 ,b=1)
abline(a=0.03298485 ,b=1)
abline(a=-0.03298485 ,b=1)
abline(lm(SS$YFP.MEDIAN.RELATIVE.MEAN ~ EMS$YFP.MEDIAN.RELATIVE.MEAN),col="red")
legend("bottomright",pch=16,col=c("black","blue","red"),legend=c("P >= 0.05","0.01 <= P < 0.05","P < 0.01"))
dev.off()


################################################################################
# 12- Factors contributing to expression differences observed between EMS and  #
#     single-site mutants (Figure 2 - figure supplement 5).                    #
################################################################################

###A - Distribution of expression differences between single-site mutants and EMS mutants (Figure 2 - figure supplement 5A).###

#SS and EMS variables were generated in section 9.
SS <- SS[-33,]
EMS <- EMS[-33,]

#Calculate the absolute expression difference between each single site mutant and the EMS mutant carrying the same mutation.
SS[,"RESIDUALS"] <- abs(SS$YFP.MEDIAN.RELATIVE.MEAN - EMS$YFP.MEDIAN.RELATIVE.MEAN)

#Plot the distribution of expression differences between EMS and single-site mutants (Panel A of Figure 2 - figure supplement 5).
BREAKS <- seq(0,0.17,by=0.01)

POSITIONS <- seq(0.005,0.195,by=0.01)
EXPECTATION <- rep(0,length(POSITIONS))

#The variable WT.DIFF is generated in section 10.
for (i in 1:length(EXPECTATION))
{
  FREQ.WT <- length(which(WT.DIFF >= POSITIONS[i] - 0.005 & WT.DIFF < POSITIONS[i] + 0.005))/length(WT.DIFF)
  
  EXPECTATION[i] <- FREQ.WT*nrow(SS)
}

pdf("Figure 2 - figure supplement 5A.pdf",useDingbats = F,height=6,width=5)
#windows(height=6,width=5)
hist(SS$RESIDUALS,breaks=BREAKS,xlim=c(0,0.2),ylim=c(0,30),xlab="Absolute expression difference relative to reference")
points(POSITIONS,EXPECTATION,type="l",col="blue")
dev.off()


###B - Comparing expression differences between single site and EMS mutants to expression noise in EMS mutants (Figure 2 - figure supplement 5B).###

pdf("Figure 2 - figure supplement 5B.pdf",useDingbats=FALSE,width=8,height=8)
#windows(height=10,width=10)
plotCI(EMS$YFP.CV.RELATIVE.MEAN,SS$RESIDUALS,ui=EMS$YFP.CV.RELATIVE.MEAN+1.96*EMS$YFP.CV.RELATIVE.SD/sqrt(EMS$N),li=EMS$YFP.CV.RELATIVE.MEAN-1.96*EMS$YFP.CV.RELATIVE.SD/sqrt(EMS$N),err="x",col=COLOR,pch=SS[,"PCH"],sfrac=0,gap=0,cex=1.5)
abline(v=1,lty=2)
abline(lm(SS$RESIDUALS~EMS$YFP.CV.RELATIVE.MEAN))
dev.off()


###C - Expression differences between single-site and EMS mutants depending on the number of mutations associated with fluorescence in the EMS mutants (Figure 2 - figure supplement 5C).###

for (i in 1:nrow(SS))
{
  CUR.STRAIN <- SS[i,"EMS.MUTANT"]
  SS[i,"N.CAUSAL"] <- ALL.MUTANT[which(ALL.MUTANT[,"STRAIN"] == as.character(CUR.STRAIN)),"N.CAUSAL"]
  SS[i,"JITTER"] <- runif(1,as.integer(SS[i,"N.CAUSAL"])-0.1,as.integer(SS[i,"N.CAUSAL"])+0.1)
}

SS <- droplevels(SS)

pdf("Figure 2 - figure supplement 5C.pdf.pdf",useDingbats=FALSE,height=8,width=5)
#windows(height=8,width=5)
boxplot(SS$RESIDUALS ~ SS$N.CAUSAL,xlab="# candidate mutations",ylab="Absolute expression difference EMS vs single site mutants",varwidth=TRUE,notch=F,ylim=c(0,0.18))
points(SS$JITTER,SS$RESIDUALS,pch=16,col="#00000066",cex=1.3)
dev.off()

SS.1 <- subset(SS, N.CAUSAL == 1)
SS.2 <- subset(SS, N.CAUSAL == 2)
wilcox.test(SS.1$RESIDUALS,SS.2$RESIDUALS)
t.test(SS.1$RESIDUALS,SS.2$RESIDUALS)


###D - Expression differences between single-site and EMS mutants depending on the G-value of mutations closest but below significance threshold in BSA-Seq (Figure 2 - figure supplement 5D).###

#Retrieve G-values closest but below significance threshold for each EMS mutant.
for (i in 1:nrow(SS))
{
  CUR.MUTANT <- subset(ALL.MUT, STRAIN == as.character(SS[i,"EMS.MUTANT"]) & GENE.FOCAL != as.character(SS[i,"GENE"]) & LINKAGE == "NO" & G.VALUE < 20)
  
  GOOD.POS <- which(CUR.MUTANT$G.VALUE == max(CUR.MUTANT$G.VALUE))
  
  if (nrow(CUR.MUTANT) != 0)
  {
    SS[i,"G.value.2"] <- CUR.MUTANT[GOOD.POS,"G.VALUE"]
    if (!is.na(CUR.MUTANT[GOOD.POS,"G.VALUE"]))
    {
      if (CUR.MUTANT[GOOD.POS,"P.SCORE"] < 0)
      {
        SS[i,"G.Sign.2"] <- -1
      } else {
        SS[i,"G.Sign.2"] <- 1
      }
    }
  }
}

SS[,"DIFF"] <- EMS$YFP.MEDIAN.RELATIVE.MEAN - SS$YFP.MEDIAN.RELATIVE.MEAN
SS[,"G.VAL.2"] <- SS[,"G.Sign.2"]*SS[,"G.value.2"]

SS.2 <- subset(SS, G.VAL.2 > -20)

pdf("Figure 2 - figure supplement 5D.pdf",height=6,width=6,useDingbats=F)
#windows(height=6,width=6)
plot(SS[,"DIFF"],SS[,"G.VAL.2"],pch=16,ylim=c(-20,20),xlim=c(-0.2,0.2))
abline(v=0,h=0,lty=2)
MODEL <- lm(SS[,"G.VAL.2"]~SS[,"DIFF"])
abline(MODEL,col="red")
dev.off()

#Correlation between 1) Expression differences between single-site and EMS mutants and 2) the G-value closest but below 20 for each EMS mutant.
cor.test(SS[,"DIFF"],SS[,"G.VAL.2"])


###E - Expression differences between single-site and EMS mutants depending on expression difference between independent clones of the single-site mutants (Figure 2 - figure supplement 5E).###

#SourceData8.txt can be found in SupplementaryFile11.zip and was generated in section 6 of the script. 
CLONES <- read.table("SourceData8.txt",header=TRUE)

CLONES <- subset(CLONES, TYPE == "SINGLE.SITE")

for (i in 1:nrow(SS))
{
  CUR.CLONES <- subset(CLONES, MUTATION == as.character(SS[i,"MUTATION"]) & EXPERIMENT == as.character(SS[i,"EXPERIMENT"]))
  
  SS[i,"N.CLONES"] <- nrow(CUR.CLONES)
  
  if (nrow(CUR.CLONES) == 2)
  {
    SS[i,"DIFF.CLONES"] <- abs(CUR.CLONES$YFP.MEDIAN.RELATIVE.MEAN[1] - CUR.CLONES$YFP.MEDIAN.RELATIVE.MEAN[2])
  }
}

cor.test(SS$RESIDUALS,SS$DIFF.CLONES)
cor.test(SS$RESIDUALS,SS$P.MEDIAN.CLONES)
cor.test(SS$RESIDUALS,-log(SS$P.MEDIAN.CLONES),method="spearman")
cor.test(SS$P.MEDIAN.EMS,SS$P.MEDIAN.CLONES)

ONE.CLONE <- subset(SS, N.CLONES != 2)
TWO.CLONE <- subset(SS, N.CLONES == 2)

nrow(ONE.CLONE)
nrow(TWO.CLONE)
mean(ONE.CLONE$RESIDUALS)
mean(TWO.CLONE$RESIDUALS)

length(which(SS$P.MEDIAN.CLONES < 0.05))
length(which(SS$P.MEDIAN.CLONES > 0.05))

t.test(ONE.CLONE$RESIDUALS,TWO.CLONE$RESIDUALS)

pdf("Figure 2 - figure supplement 5E.pdf",height=6,width=6,useDingbats=F)
#windows(height=6,width=6)
plot(SS[,"RESIDUALS"],SS[,"DIFF.CLONES"],pch=16,xlim=c(0,0.2),ylim=c(0,0.2),xlab="Expression difference between EMS and single-site mutants",ylab="Expression difference between independent clones",col=COLOR)
abline(a=0,b=1,lty=1,col="#00000033")
MODEL <- lm(SS[,"DIFF.CLONES"]~SS[,"RESIDUALS"])
abline(MODEL,lty=2)
dev.off()


###F - Expression differences between independent clones of single-site mutants depending on the statistical
###    significance of expression differences between single-site and EMS mutants (Figure 2 - figure supplement 5F). ###

SS[,"COLOR"] <- as.factor(COLOR)
TWO.CLONE <- subset(SS, N.CLONES == 2)
TWO.CLONE <- droplevels(TWO.CLONE)

pdf("Figure 2 - figure supplement 5F.pdf",useDingbats=FALSE,height=8,width=5)
#windows(height=6,width=5)
boxplot(TWO.CLONE$DIFF.CLONES ~ TWO.CLONE$COLOR,xlab="Significance of expression difference",ylim=c(0,0.25),ylab="Expression difference between independent clones",varwidth=TRUE,notch=F)
dev.off()

SIG <- subset(TWO.CLONE, COLOR == "blue")
NO.SIG <- subset(TWO.CLONE, COLOR == "black")

wilcox.test(SIG$DIFF.CLONES,NO.SIG$DIF.CLONES)

#The two matching datasets with expression values of single-site mutants and EMS mutants can be found in SupplementaryFile11.zip.
write.table(SS,"SourceData11.txt",sep="\t",row.names=FALSE)
write.table(EMS,"SourceData12.txt",sep="\t",row.names=FALSE)


#################################################################################
# 13- Distribution of expression levels observed among RAP1 mutants (Figure 5E).#
#################################################################################

###Clear memory###
rm(list=ls())
options(warn=-1)

####Load packages###
##library(VariantAnnotation)
library(VariantAnnotation)
library(Deducer)
library(gtools)
library(zoo)
library(Hmisc)
library(ggplot2)
library(plotrix)
library(moments)
library(vcd)

###Set directory###
setwd("/Path.to.input.files")

#Expression data for RAP1 mutants generated using sections 1, 2, 6 and 7 of the script.
#The flow template of the experiment is included in the file TEMPLATE.RAP1.MUTANTS.txt in SupplementaryFile11.zip.
#SourceData14.txt is available in SupplementaryFile11.zip.
DATA <- read.table("SourceData14.txt",header=TRUE)

DATA <- subset(DATA, MUTATION == "RANDOM" & CLASS == "RANDOM")

#Filter out strains with whole genome duplication.
DATA <- subset(DATA, FSC.MEDIAN.RELATIVE.MEAN < 1.01)
DATA <- subset(DATA, YFP.MEDIAN.RELATIVE.SD < 0.15)

RANDOM <- subset(DATA, CLASS == "RANDOM")

#Apply correction for multiple testing.
RANDOM[,"P.FDR"] <- p.adjust(RANDOM$P.VAL.MEDIAN,method="fdr")
P.FDR <- p.adjust(RANDOM$P.VAL.MEDIAN,method="fdr")

SIG <- subset(RANDOM, P.FDR < 0.05 & (YFP.MEDIAN.RELATIVE.MEAN < 0.97 | YFP.MEDIAN.RELATIVE.MEAN > 1.03))

pdf("Figure6E_Distribution.pdf",useDingbats=F,height=8,width=13)
#windows(height=8,width=13)
LOW <- subset(RANDOM, P.FDR < 0.05 & YFP.MEDIAN.RELATIVE.MEAN < 1)
HIGH <- subset(RANDOM, P.FDR < 0.05 & YFP.MEDIAN.RELATIVE.MEAN > 1)
BREAKS <- seq(0,1.4,by=0.02)
hist(RANDOM$YFP.MEDIAN.RELATIVE.MEAN, breaks=BREAKS, main = "Random RAP1 mutants", xlab="Median Expression Relative to WT", ylab = "Frequency", col="gray",xlim=c(0,1.4),ylim=c(0,250),xaxt="n",freq=T)
hist(LOW$YFP.MEDIAN.RELATIVE.MEAN, breaks=BREAKS, main = "Random RAP1 mutants", xlab="Median Expression Relative to WT", ylab = "Frequency", col="red",xlim=c(0,1.4),ylim=c(0,250),xaxt="n",freq=T,add=T)
hist(HIGH$YFP.MEDIAN.RELATIVE.MEAN, breaks=BREAKS, main = "Random RAP1 mutants", xlab="Median Expression Relative to WT", ylab = "Frequency", col="blue",xlim=c(0,1.4),ylim=c(0,250),xaxt="n",freq=T,add=T)
axis(1,pos=0)
dev.off()


pdf("Figure6E_Pie_Chart.pdf",useDingbats=F,height=7,width=10)
#windows(height=7,width=10)
SLICES <- c(nrow(RANDOM)-nrow(LOW)-nrow(HIGH),nrow(LOW),nrow(HIGH))
LABELS <- c("No significant change","Decreased expression","Increased expression")
PCT <- round(SLICES/sum(SLICES)*100,1)
LABELS <- paste(LABELS, PCT)
LABELS <- paste(LABELS, "%", sep="")
pie(SLICES,labels=LABELS,col=c("#00000033","#FF0000CC","#0000FFCC"))
dev.off()


#################################################################################
# 14- Distribution of expression levels observed among GCR1 mutants (Figure 5F).#
#################################################################################

###Clear memory###
rm(list=ls())
options(warn=-1)

####Load packages###
##library(VariantAnnotation)
library(VariantAnnotation)
library(Deducer)
library(gtools)
library(zoo)
library(Hmisc)
library(ggplot2)
library(plotrix)
library(moments)
library(vcd)

###Set directory###
setwd("/Path.to.input.files")

#Expression data for GCR1 mutants generated using sections 1, 2, 6 and 7 of the script.
#The flow template of the experiment is included in the file TEMPLATE.GCR1.MUTANTS.txt in SupplementaryFile11.zip.
#SourceData15.txt is available in SupplementaryFile11.zip.
DATA <- read.table("SourceData15.txt",header=TRUE)

#Filter out strains with whole genome duplication.
PLOT <- subset(DATA,  (FSC.MEDIAN.RELATIVE.MEAN < 1 & YFP.MEDIAN.RELATIVE.SD < 0.10) | (P.VAL.FSC > 0.05 & YFP.MEDIAN.RELATIVE.SD < 0.10))

#Histogram random mutants

pdf("Figure6F_Distribution.pdf",useDingbats=F,height=8,width=11)
NOSIG <- subset(PLOT, P.VAL.MEDIAN > 0.05)
DEC <- subset(PLOT, YFP.MEDIAN.RELATIVE.MEAN < 1)
INC <- subset(PLOT, YFP.MEDIAN.RELATIVE.MEAN > 1)
BREAKS <- seq(0,1.4,by=0.02)
hist(DEC$YFP.MEDIAN.RELATIVE.MEAN, breaks=BREAKS, main = "Random GCR1 mutants", xlab="Median Expression Relative to WT", ylab = "Frequency", col="red",border="red", xlim=c(0,1.2), ylim=c(0,80), freq=T)
hist(INC$YFP.MEDIAN.RELATIVE.MEAN, breaks=BREAKS, main = "Random GCR1 mutants", xlab="Median Expression Relative to WT", ylab = "Frequency", col="blue", border="blue",add=TRUE, freq=T)
hist(NOSIG$YFP.MEDIAN.RELATIVE.MEAN, breaks=BREAKS, main = "Random GCR1 mutants", xlab="Median Expression Relative to WT", ylab = "Frequency", col="black",add=TRUE, freq=T)
dev.off()

#Pie chart random mutants

pdf("Figure6F_Pie_Chart.pdf",useDingbats=F,height=7,width=10)
#quartz(height=7,width=10)
NOSIG <- subset(PLOT, P.VAL.MEDIAN > 0.05)
SLICES <- c(nrow(NOSIG),nrow(subset(PLOT, YFP.MEDIAN.RELATIVE.MEAN < 1 & P.VAL.MEDIAN < 0.05)),nrow(subset(PLOT, YFP.MEDIAN.RELATIVE.MEAN > 1 & P.VAL.MEDIAN < 0.05)))
LABELS <- c("No significant change","Decreased expression","Increased expression")
PCT <- round(SLICES/sum(SLICES)*100,1)
LABELS <- paste(LABELS, PCT)
LABELS <- paste(LABELS, "%", sep="")
pie(SLICES,labels=LABELS,col=c("black","red","blue"))
dev.off()


###########################################
# 15- Fitness of GCR1 mutants in glucose. #
###########################################

#Fitness data were collected for cells grown either in glucose or in glycerol during the same experiment, but only the glucose data were used in Figure 5G.

###Clear memory##
rm(list=ls())
options(warn=-1)

###Necessary libraries##
library(flowCore)
library(flowClust)
library(mixtools)
library(mratios)
library(gplots)
library(fitdistrplus)
library(RColorBrewer)
library(pcaPP)
library(plotrix)


###Set working directory###
parent.dir <- "/Path.to.input.files"
setwd(parent.dir)

#The template file "TEMPLATE.GCR1.FITNESS.txt" can be found in SupplementaryFile11.zip.
Design.xp <- read.table("TEMPLATE.GCR1.FITNESS.txt",header=TRUE,as.is=TRUE)

###Create list of all fcs filenames###
FILENAMES <- list.files(".",pattern=".fcs",recursive=TRUE,include.dirs=TRUE)

###Create Output File###
Output.data <- Design.xp

RANDOM <- sample(1:6144,size=96,replace=FALSE)

#Open pdf for ploting flow data
pdf("Plot_YFP.GFP.pdf",width=14,height=12)

###Processing .fcs files###
for(i in 1:nrow(Design.xp))
{
  Merge.Frame <- read.FCS(as.character(FILENAMES[i]),transformation=FALSE,alter.names=TRUE)
  
  Output.data[i,"COUNTS.INITIAL"] <- nrow(exprs(Merge.Frame))
  
  
  if (is.na(Output.data[i,"COUNTS.INITIAL"]))
  {Output.data[i,"COUNTS.INITIAL"] <- 0}
  
  
  if (Output.data[i,"COUNTS.INITIAL"] >= 3000)
  {
    
    ###Log transformation of flow data###

    Data.Fluo <- exprs(Merge.Frame)
    Data.Fluo[Data.Fluo == 0] <- NA
    Data.Fluo <- Data.Fluo[complete.cases(Data.Fluo[,c("FSC.A","FSC.H","FL1.H","FL2.H")]),]
    Merge.Frame <- new("flowFrame",Data.Fluo)	
    
    logTrans <- logTransform(transformationId="log10-transformation",logbase=10,r=1,d=1)
    Merge.Frame <- transform(Merge.Frame,`logFSC.A`=logTrans(`FSC.A`))
    Merge.Frame <- transform(Merge.Frame,`logFSC.H`=logTrans(`FSC.H`))
    Merge.Frame <- transform(Merge.Frame,`logFL1.A`=logTrans(`FL1.A`))
    Merge.Frame <- transform(Merge.Frame,`logFL1.H`=logTrans(`FL1.H`))
    Merge.Frame <- transform(Merge.Frame,`logFL2.A`=logTrans(`FL2.A`))
    Merge.Frame <- transform(Merge.Frame,`logFL2.H`=logTrans(`FL2.H`))
    
    
    ###Calculate phenotypes of interest###

    Data.Fluo <- exprs(Merge.Frame)
    
    Phenotype1 <- Data.Fluo[,"logFL1.H"]^2/Data.Fluo[,"logFSC.A"]^3
    Phenotype1 <- as.matrix(Phenotype1)
    colnames(Phenotype1) <- "FL1^2/FSC^3"
    Merge.Frame <- cbind2(Merge.Frame, Phenotype1)
    
    Phenotype2 <- (Data.Fluo[,"logFL2.H"])^2/Data.Fluo[,"logFSC.A"]^3
    Phenotype2 <- as.matrix(Phenotype2)
    colnames(Phenotype2) <- "FL2^2/FSC^3"
    Merge.Frame <- cbind2(Merge.Frame, Phenotype2)
    
    Phenotype3 <- Phenotype2/Phenotype1
    Phenotype3 <- as.matrix(Phenotype3)
    colnames(Phenotype3) <- "FL2/FL1"
    Merge.Frame <- cbind2(Merge.Frame, Phenotype3)
    
    Phenotype4 <- (Data.Fluo[,"logFSC.A"]^8)/(Data.Fluo[,"logFSC.H"]^9)
    Phenotype4 <- as.matrix(Phenotype4)
    colnames(Phenotype4) <- "FSC.A/FSC.H"
    Merge.Frame <- cbind2(Merge.Frame, Phenotype4)
    
    PlotAll <- exprs(Merge.Frame)
    
    Output.data[i,"COUNTS.COMPLETE"] <- nrow(PlotAll)
    
    
    ###Gating YFP, GFP and YFP+GFP events###
    
    rectGate <- rectangleGate(filterId="Noise Removal","logFSC.A"=c(4.5,6.7),"FSC.A/FSC.H"=c(0.065,0.12), "logFL2.H"=c(1,5.9), "logFL1.H"=c(1,6.3))
    
    Hard.Gates <- Subset(Merge.Frame, rectGate)
    Hard.Gates.exp <- as.data.frame(exprs(Hard.Gates))
    
    #Removing doublets

    #FSC.A FILTERING
    
    DENSITY <- density(Hard.Gates.exp[,"logFSC.A"],adjust=3)
    
    MED <- median(Hard.Gates.exp[,"logFSC.A"])
    
    XY <- as.data.frame(cbind(DENSITY$x,DENSITY$y))
    colnames(XY) <- c("X","Y")
    
    LEFT <- subset(XY, X < MED)
    DIST <- abs(LEFT$Y - 0.1)
    LOW <- LEFT[DIST == min(DIST),"X"]
    
    RIGHT <- subset(XY, X > MED)
    DIST <- abs(RIGHT$Y - 0.1)
    HIGH <- RIGHT[DIST == min(DIST),"X"]
    
    Normal.Size <- subset(Hard.Gates.exp, logFSC.A > LOW & logFSC.A < HIGH)
    
    
    #FSC.A/FSC.H FILTERING
    
    DENSITY <- density(Normal.Size[,"FSC.A/FSC.H"],adjust=3)
    
    MED <- median(Hard.Gates.exp[,"FSC.A/FSC.H"])
    
    XY <- as.data.frame(cbind(DENSITY$x,DENSITY$y))
    colnames(XY) <- c("X","Y")
    
    LEFT <- subset(XY, X < MED)
    DIST <- abs(LEFT$Y - 5)
    LOW <- LEFT[DIST == min(DIST),"X"]
    
    RIGHT <- subset(XY, X > MED)
    DIST <- abs(RIGHT$Y - 5)
    HIGH <- RIGHT[DIST == min(DIST),"X"]
    
    Normal.Shape <- subset(Normal.Size, Normal.Size[,"FSC.A/FSC.H"] > LOW & Normal.Size[,"FSC.A/FSC.H"] < HIGH)
    
    
    #Select fluorescent clusters based on FL2/FL1

    #Remove correlation between FL1.H and FL2.H
    Fluo.Model <- PCAgrid(cbind(Normal.Shape[,"logFL1.H"], Normal.Shape[,"logFL2.H"]),k=2,scale="sd",method="sd",scores=TRUE,center="median")
    
    Scores <- Fluo.Model$scores
    
    # plot(Scores[,1],Scores[,2],pch=20,cex=0.3,col="#00000044")
    
    if (Fluo.Model$loadings[2,2] < 0)
    {
      Scores[,2] <- -1 * Scores[,2]
    }
    
    # Find Modes of Two Populations
    # HIST <- hist(Scores[,2],breaks=100)
    DENSITY <- density(Scores[,2])
    
    MODES <- c(0,0)
    COUNT <- 1
    
    for (j in 2:(length(DENSITY$y)-1))
    {
      if (DENSITY$y[j] > DENSITY$y[j-1] & DENSITY$y[j] > DENSITY$y[j+1] & DENSITY$y[j] > 0.3)
      {
        MODES[COUNT] <- DENSITY$x[j]
        COUNT <- 1 + COUNT
      }
    }	
    
    MODES <- MODES[c(1,length(MODES))]
    
    if (MODES[2] != 0)
    {
      THRESHOLD <- mean(MODES)
    } else if (median(Normal.Shape[,"FL2/FL1"]) < 0.7) {
      THRESHOLD <- 1000
    } else {
      THRESHOLD <- -1000
    }
    
    YFP.POSITIONS <- c(which(Scores[,2] > THRESHOLD), which(Scores[,2] < THRESHOLD & Normal.Shape[,"logFL1.H"] < 3.5 & Normal.Shape[,"logFL2.H"] < 3.3))
    GFP.POSITIONS <- which(Scores[,2] < THRESHOLD & (Normal.Shape[,"logFL1.H"] > 3.5 | Normal.Shape[,"logFL2.H"] > 3.3))
    
    #Select fluorescent clusters
    YFP.CELLS <- Normal.Shape[YFP.POSITIONS,]
    GFP.CELLS <- Normal.Shape[GFP.POSITIONS,]
    
    # plot(Scores[,1],Scores[,2],pch=20,cex=0.3,col="#00000044")
    # abline(h=GFP.LOW)
    # abline(h=GFP.HIGH)
    # abline(h=YFP.LOW)
    # abline(h=YFP.HIGH)
    # abline(v=LOW)
    
    
    ###Calculate number of YFP and GFP cells###

    N.YFP <- nrow(YFP.CELLS)
    N.GFP <- nrow(GFP.CELLS)
    
    
    ###Add data in output table###

    Output.data[i,"Filtered.events"] <- nrow(Normal.Shape)
    Output.data[i,"N.YFP"] <- N.YFP
    Output.data[i,"N.GFP"] <- N.GFP
    Output.data[i,"N.NoFP"] <- nrow(Normal.Shape)-(N.GFP+N.YFP)
    Output.data[i,"Freq.YFP"] <- N.YFP/(N.YFP+N.GFP)
    Output.data[i,"Freq.GFP"] <- N.GFP/(N.YFP+N.GFP)
    Output.data[i,"Freq.noFP"] <- (nrow(Normal.Shape)-(N.GFP+N.YFP))/nrow(Normal.Shape)
    
    
    ###Plot Final Gating in pdf###

    if (i %in% RANDOM)
    {
      #quartz(height=12,width=14)
      par(mfrow=c(2,2))
      
      plot(PlotAll[,"logFSC.A"],PlotAll[,"logFSC.H"],pch=20,cex=0.3,col="#00000044",main=paste(Output.data[i,"STRAIN"],"_",Output.data[i,"MUTATION"],"_","P",Output.data[i,"PLATE"],Output.data[i,"POSITION"],"_","T",Output.data[i,"TIME.POINT"],sep=""),cex.main=2,xlab="logFSC.A",ylab="logFSC.H")
      points(YFP.CELLS[,"logFSC.A"], YFP.CELLS[,"logFSC.H"],col="#FF990044",pch=20,cex=0.3)
      points(GFP.CELLS[,"logFSC.A"], GFP.CELLS[,"logFSC.H"],col="#22CC2244",pch=20,cex=0.3)
      abline(v=4.6)
      abline(v=6.7)
      
      plot(PlotAll[,"logFL2.H"],PlotAll[,"logFL1.H"],pch=20,cex=0.3,col="#00000044",xlab="logYFP.H",ylab="logGFP.H")
      points(YFP.CELLS[,"logFL2.H"], YFP.CELLS[,"logFL1.H"],col="#FF990044",pch=20,cex=0.3)
      points(GFP.CELLS[,"logFL2.H"], GFP.CELLS[,"logFL1.H"],col="#22CC2244",pch=20,cex=0.3)
      
      plot(Scores[,1],Scores[,2],pch=20,cex=0.3,col="#00000044",xlab="PC.1",ylab="PC.2")
      abline(h=THRESHOLD)
    }
  }
  
  cat(i," of ",nrow(Design.xp)," is done\n")
}

dev.off()

#write.table(Output.data,file="Experiment_Output.txt",row.names=FALSE,col.names=TRUE,sep="\t",quote=FALSE)
#Output.data <- read.table("Experiment_Output.txt",header=TRUE,as.is=TRUE)


###Look for mistakes in well separation or when saving data.###

Output.data <- subset(Output.data, ENVIRONMENT != "GLYCEROL" | PLATE != 2)

COUNTS <- rep(0,nrow(Output.data))

for (i in 1:nrow(Output.data))
{
  CUR <- subset(Output.data, COUNTS.INITIAL == Output.data[i,"COUNTS.INITIAL"] & COUNTS.COMPLETE == Output.data[i,"COUNTS.COMPLETE"])
  COUNTS[i] <- nrow(CUR)
}

POS <- which(COUNTS == 2)

length(unique(Output.data[,"COUNTS.INITIAL"]))

###Sort Data by Environment, Plate, plateposition and Time.point###

Sorted.data <- Output.data[order(Output.data[,"ENVIRONMENT"],Output.data[,"PLATE"],Output.data[,"POSITION"],Output.data[,"TIME.POINT"]),]

write.table(Sorted.data,file="Experiment_Output_Sorted.txt",row.names=FALSE,col.names=TRUE,sep="\t",quote=FALSE)


###Reformat table to remove redundancy from time points###

Data.preformat <- read.table("Experiment_Output_Sorted.txt",header=TRUE,as.is=TRUE)

#Remove bad samples (contaminations & wrong files)

Data.preformat <- subset(Data.preformat, PLATE != 4 | POSITION != "F10")
Data.preformat <- subset(Data.preformat, PLATE != 3 | POSITION != "H10" | ENVIRONMENT != "GLYCEROL")
Data.preformat <- subset(Data.preformat, PLATE != 4 | ROW != "H" | ENVIRONMENT != "GLUCOSE")

N.points <- 2

Data.format <- Data.preformat[,c("ID","STRAIN","COMPETITOR","POSITION","COLUMN","ROW","CLASS","ENVIRONMENT","PLATE","REP")] 

Raw.names <- rep(colnames(Data.preformat[,13:ncol(Data.preformat)]),each=N.points)
Numbers <- rep(c(1:N.points),length(Raw.names)/N.points)
Full.names <-c()

for (i in 1:length(Raw.names))
{
  Full.names[i] <- paste(Raw.names[i],Numbers[i],sep="_")
} 

# Data.format[1,1:ncol(Data.format)] <- rep(NA,ncol(Data.format))

for (i in Full.names)
{
  Data.format[,i] <- NA
}

Data.format <- Data.format[seq(1,nrow(Data.preformat),by=2),]

###Automatic filling of all columns

Data <- Data.preformat

for (j in 13:ncol(Data))
{
  Current.matrix <- matrix(Data[,j],ncol=N.points,nrow=nrow(Data)/N.points,byrow=TRUE)
  Data.format[,((13+((j-12)*N.points-(N.points-1)))-3):((13+((j-12)*N.points))-3)] <- as.data.frame(Current.matrix)
}


write.table(Data.format,file="Fitness/Experiment_Output_Formatted.txt",row.names=FALSE,col.names=TRUE,sep="\t",quote=FALSE)


###Compute relative fitness###

Data <- read.table("Fitness/Experiment_Output_Formatted.txt",header=TRUE,as.is=TRUE)

#Table with the time in minutes between the two flow cytometry runs for each plate. Can be found in SupplementaryFile11.zip.
Time <- read.table("Fitness/Time.Points.txt",header=TRUE,as.is=TRUE)

#Table with the dilution factors used to estimate the average number of generations during the fitness assay. Can be found in SupplementaryFile11.zip.
Dilution <- read.table("Fitness/Dilutions.txt",header=TRUE,as.is=TRUE)

#1-Compute number of generations for each sample
for (i in 1:nrow(Data))
{
  CUR.DILUTION <- subset(Dilution, ENVIRONMENT == Data[i,"ENVIRONMENT"] & PLATE == Data[i,"PLATE"])	
  START.DENSITY <- (Data[i,"DENSITY.YFP_1"]/CUR.DILUTION[1,"DILUTION.YFP"] + Data[i,"DENSITY.GFP_1"]/CUR.DILUTION[1,"DILUTION.GFP"] )/CUR.DILUTION[1,"DILUTION.MIX"]
  Data[i,"GENERATION"] <- log(Data[i,"DENSITY.MIX_2"]/START.DENSITY)/log(2)
}

#2-Compute number of generations for each plate
CONTROL <- subset(Data, CLASS == "CONTROL")

for (i in 1:nrow(Data))
{
  CUR.PLATE <- subset(CONTROL, ENVIRONMENT == Data[i,"ENVIRONMENT"] & PLATE == Data[i,"PLATE"])
  
  Data[i,"GEN.PLATE"] <- median(CUR.PLATE[,"GENERATION"])
}

#3-Calculate log.ratio YFP/GFP
Data.Mix <- subset(Data, COMPETITOR != "EMPTY")

Data.Mix[,"log.ratio_1"] <- log(Data.Mix[,"N.YFP_1"]/Data.Mix[,"N.GFP_1"])
Data.Mix[,"log.ratio_2"] <- log(Data.Mix[,"N.YFP_2"]/Data.Mix[,"N.GFP_2"])

#3-Calculate Fitness
for (i in 1:nrow(Data.Mix))
{
  log.ratio <- unname(unlist(Data.Mix[i,c("log.ratio_1","log.ratio_2")]))
  G <- c(0,unlist(Data.Mix[i,"GEN.PLATE"]))
  Generation <- c(G[1],sum(G[1:2]))
  DataFrame <- data.frame(cbind(log.ratio,Generation))
  colnames(DataFrame) <- c("log.ratio","Generation")
  Model <- lm(log.ratio~Generation,data=DataFrame,na.action=na.exclude)
  Data.Mix[i,"w.estimate"] <- exp(Model$coef[2])	
}

write.table(Data.Mix,file="Experiment_s.estimates.txt",row.names=FALSE,col.names=TRUE,sep="\t",quote=FALSE)


###Filter out outliers###

Data <- read.table("Experiment_s.estimates.txt",header=TRUE,as.is=TRUE)

#FLAG OUTLIERS FOR S.ESTIMATE
for (i in 1:nrow(Data))
{
  CUR.STRAIN <- subset(Data, ENVIRONMENT == Data[i,"ENVIRONMENT"] & STRAIN == Data[i,"STRAIN"])
  LOW <- median(CUR.STRAIN$w.estimate,na.rm=TRUE) - 4*mad(CUR.STRAIN$w.estimate,na.rm=TRUE)
  HIGH <- median(CUR.STRAIN$w.estimate,na.rm=TRUE) + 4*mad(CUR.STRAIN$w.estimate,na.rm=TRUE)
  
  if (is.na(Data[i,"w.estimate"]))
  {
    Data[i,"OUTLIER"] <- NA
  }	else if (Data[i,"w.estimate"] < LOW | Data[i,"w.estimate"] > HIGH) {
    Data[i,"OUTLIER"] <- "YES"
  } else {
    Data[i,"OUTLIER"] <- "NO"
  }
}

Data.filter <- subset(Data, Data[,"OUTLIER.2"] == "NO" | Data[,"OUTLIER"] == "NO")

write.table(Data.filter,file="Experiment_s.estimates_filtered.txt",row.names=FALSE,col.names=TRUE,sep="\t",quote=FALSE)


###Calculate Fitness Relative To WT###

for (i in 1:nrow(Data))
{
  WT <- subset(Data, ENVIRONMENT == Data[i,"ENVIRONMENT"] & PLATE == Data[i,"PLATE"] & CLASS == "CONTROL")
  Data[i,"Fitness"] <- Data[i,"w.estimate"]/mean(WT[,"w.estimate"])
}

write.table(Data,file="Experiment_fitness.txt",row.names=FALSE,col.names=TRUE,sep="\t",quote=FALSE)


###Make summary table with statistical power###

Data <- read.table("Experiment_fitness.txt",header=TRUE,as.is=TRUE)

Mean.table <- aggregate(Data[,c("w.estimate","Fitness")],by=list(Data$ID,Data$STRAIN,Data$POSITION,Data$CLASS,Data$ENVIRONMENT),FUN=mean)
colnames(Mean.table) <- c("ID","STRAIN","POSITION","CLASS","ENVIRONMENT","W_MEAN","FITNESS_MEAN")

SD.table <- aggregate(Data[,c("w.estimate","Fitness")],by=list(Data$ID,Data$STRAIN,Data$POSITION,Data$CLASS,Data$ENVIRONMENT),FUN=sd)
colnames(SD.table) <- c("ID","STRAIN","POSITION","CLASS","ENVIRONMENT","W_SD","FITNESS_SD")

N.table <- aggregate(Data[,c("w.estimate","Fitness")],by=list(Data$ID,Data$STRAIN,Data$POSITION,Data$CLASS,Data$ENVIRONMENT),FUN=length)
colnames(N.table) <- "N.REP"

Combined <- cbind(Mean.table,SD.table[,6:ncol(SD.table)],N.table[,ncol(N.table)])
colnames(Combined)[ncol(Combined)] <- "N.REP"

for (i in 1:nrow(Combined))
{
  s.power <- power.t.test(n=Combined[i,"N.REP"],sd=Combined[i,"FITNESS_SD"],sig.level=0.05,power=0.9,type="two.sample",alternative="two.sided")
  Combined[i,"POWER"] <- s.power$delta
}

write.table(Combined,file="Summary.data.txt",row.names=FALSE,col.names=TRUE,sep="\t",quote=FALSE)

GLYCEROL <- subset(Combined, ENVIRONMENT == "GLYCEROL" & CLASS %in% c("HIGH.EXPR","LOW.EXPR","WT.EXPR"))
GLUCOSE <- subset(Combined, ENVIRONMENT == "GLUCOSE" & CLASS %in% c("HIGH.EXPR","LOW.EXPR","WT.EXPR"))

#plot(GLUCOSE$FITNESS_MEAN,GLYCEROL$FITNESS_MEAN,pch=20,xlim=c(0.5,1.05),ylim=c(0.5,1.05))
cor.test(GLUCOSE$FITNESS_MEAN,GLYCEROL$FITNESS_MEAN)

#write.table(Combined,file="Summary.data.txt",row.names=FALSE,col.names=TRUE,sep="\t",quote=FALSE)
#Glucose.Fitness.txt can be found in SupplementaryFile11.zip.
write.table(GLUCOSE,file="Fitness/Glucose.Fitness.txt",row.names=FALSE,col.names=TRUE,sep="\t",quote=FALSE)
#write.table(GLYCEROL,file="Glycerol.data.txt",row.names=FALSE,col.names=TRUE,sep="\t",quote=FALSE)


####################################################################################
# 16- Relationship between fitness and YFP expression in GCR1 mutants (Figure 5G). #
####################################################################################

#All input files can be found in SupplementaryFile11.zip.

#Expression data averaged across replicates for each GCR1 mutant.
GLU <- read.table("Fitness/Summary.Expression.txt",header=TRUE)

#Expression data for each replicate population of GCR1 mutants.
GLU.DATA <- read.table("Fitness/All.Expression.txt",header=TRUE)

#Fitness data average across replicates for each GCR1 mutant.
FIT.GLU <- read.table("Fitness/Glucose.Fitness.txt",header=TRUE) 

GLU <- subset(GLU, CLASS %in% c("LOW.EXPR","HIGH.EXPR","WT.EXPR"))

FIT.GLU <- FIT.GLU[order(FIT.GLU[,"STRAIN"]),]

#Plotting relative fitness vs pTDH3-YFP expression for each GCR1 mutant.
pdf("Figure6G.pdf",useDingbats=FALSE,width=6,height=6)

#quartz(height=6,width=6)
par(mar=c(6,6,3,1)+0.1,mgp=c(4,1,0))
plotCI(x=GLU$YFP.MEDIAN.RELATIVE.MEAN,y=FIT.GLU$FITNESS_MEAN,uiw=1.96*GLU$YFP.MEDIAN.RELATIVE.SD/sqrt(GLU$N),liw=1.96*GLU$YFP.MEDIAN.RELATIVE.SD/sqrt(GLU$N),pch=20,cex=2,xlab="Median Expression in Glucose",ylab="Fitness in Glucose",pt.bg=par("bg"),gap=0,col="#00000000",err="x",font.axis=1,font.lab=2,cex.lab=1.2,cex.axis=1,sfrac=0,scol="#00000044",lwd=1,xaxt="n",yaxt="n")
plotCI(x=GLU$YFP.MEDIAN.RELATIVE.MEAN,y=FIT.GLU$FITNESS_MEAN,uiw=1.96*FIT.GLU$FITNESS_SD/sqrt(FIT.GLU$N.REP),liw=1.96*FIT.GLU$FITNESS_SD/sqrt(FIT.GLU$N.REP),pch=20,cex=2,xlab="Fitness in GLUCEROL",ylab="Fitness in GLUcerol",pt.bg=par("bg"),gap=0,col="#00000066",err="y",font.axis=1,font.lab=1,cex.lab=1.2,cex.axis=1,sfrac=0,scol="#00000044",lwd=1,add=TRUE)
axis(1,lwd=1,at=seq(0,1.2,by=0.2),font=2,cex.axis=1)
axis(2,lwd=1,at=seq(0.5,1.1,by=0.1),las=2,font=2,cex.axis=1)
abline(h=1,lty=2,lwd=1)
abline(v=1,lty=2,lwd=1)

MODEL.FIT <- loess(FIT.GLU$FITNESS_MEAN~GLU$YFP.MEDIAN.RELATIVE.MEAN,degree=2,span=0.66)

x.mid <- seq(min(GLU$YFP.MEDIAN.RELATIVE.MEAN),max(GLU$YFP.MEDIAN.RELATIVE.MEAN),by=0.001)
y.mid <- predict(MODEL.FIT, x.mid, se=TRUE)
points(x.mid,y.mid$fit,type="l",col="#00000099",lwd=2,lty=2)

y.err <- c(y.mid$fit + 2.58*y.mid$se.fit,rev(y.mid$fit - 2.58*y.mid$se.fit))

polygon(c(x.mid,rev(x.mid)),y.err,col="#00000033",border=NA)

dev.off()


##################################################################################
# 17- Effects of mutations in purine biosynthesis genes on YFP expression levels #
#     driven by different promoters (Figure 2 - figure supplement 4).            #
##################################################################################

#As described in the methods, flow cytometry data shown on Figure 2 - figure supplement 4 were collected on a different instrument and analyzed with a different script (below)
#than other cytometry data included in this study.

###Necessary libraries###
rm(list = ls())
library(flowCore)
library(flowWorkspace)
library(openCyto)
library(ggcyto)
library(flowAI)
library(gridExtra)
library(pcaPP)
library(MASS)
library(scales)
library(ggthemes)
library(ggsci)
library(tidyverse)

##Import files----
## Folders are imported.  Each folder is a batch.  With each .fcs file corresponds to a genotype.
myfiles071420 <- list.files(path="Experiments_071420/", pattern = ".fcs",
                            ignore.case = TRUE)
fs071420 <- read.flowSet(myfiles071420, path="Experiments_071420/", truncate_max_range = FALSE,
                         min.limit = 1)

myfiles071720 <- list.files(path="Experiments_071720/", pattern = ".fcs",
                            ignore.case = TRUE)
fs071720 <- read.flowSet(myfiles071720, path="Experiments_071720/", truncate_max_range = FALSE,
                         min.limit = 1)

myfiles072220 <- list.files(path="Experiments_072220/", pattern = ".fcs",
                            ignore.case = TRUE)
fs072220 <- read.flowSet(myfiles072220, path="Experiments_072220/", truncate_max_range = FALSE,
                         min.limit = 1)

#Channels found in the flow cytometry data
channels <- c("FSC-A", "FSC-H", "FSC-W", "SSC-A", "SSC-H", "SSC-W", "FL1-A", "FL1-H", "FL1-W")

###The file "ade_genotype_key.csv" included in SupplementaryFile11.zip contains a genotype key to convert strain number's to genotypes.
genotype_key <- read_csv("ade_genotype_key.csv") %>%
  mutate(Strain = as.character(Strain))

## Functions-----
## These are the functions that will process the data, including log transforming the data for drawing gates, correcting for  size
## and extracting summary statistics. 

rotmat <- function(fluo_model, fsc){
  Intercept <- c()
  Slope <- c()
  Theta <- c()
  ROT <- function(x,Rotation){ #Basic rotation function
    Result <- Rotation%*%x
    return(Result)
  }
  #2-Center of rotation
  x.center <- fluo_model$center[1]
  y.center <- fluo_model$center[2]
  #3-Initial Intercept and Slope
  Slope[1] <- fluo_model$loadings[2,1] / fluo_model$loadings[1,1]
  Intercept[1] <- fluo_model$center[2] - Slope[1]*fluo_model$center[1]
  
  #4-Calculate angle of rotation
  a <- c(x.center-0,y.center-Intercept[1]) #Vector from Intercept to Centroid
  b <- c(x.center-0,y.center-y.center) #Vector with slope 0 through Centroid
  Theta[1] <- acos(sum(a*b)/(sqrt(sum(a*a))*sqrt(sum(b*b)))) #Angle between 2 vectors
  if (Slope[1] < 0){
    Theta[1] <- -Theta[1]
  }		
  
  #5-Define rotation matrix
  Rotation <- matrix(c(cos(Theta[1]),-sin(Theta[1]),sin(Theta[1]),cos(Theta[1])),ncol=2,nrow=2)
  
  #6-Transform Data
  Coord <- t(as.matrix(fsc[,c("FSC.A","FL1.A")]))
  Coord[1,] <- Coord[1,] - x.center
  Coord[2,] <- Coord[2,] - y.center
  
  Result <- ROT(x=Coord,Rotation=Rotation)
  
  Result[1,] <- Result[1,] + x.center
  Result[2,] <- Result[2,] + y.center
  
  #7-Keep record of rotated values
  fsc[,"FSC.FINAL"] <- Result[1,]
  fsc[,"YFP.ROT"] <- Result[2,]
  
  return(fsc)
} #This takes the flow model from PCA grid and the original flow cytometry file 
##This will modify the flowSet file as well as create a list of dataframes for easier analysis
size_correction <- function(flowset){
  flowdata_adjusted <- c()
  flowdata_adjustedmatrix <- flowset
  for (i in 1:length(flowset)){
    singlet_gate <- flowCore::filter(flowset[[i]], rg2) #Establish the gate
    flowdata_adjustedmatrix[[i]] <- Subset(flowset[[i]], singlet_gate) #Keep only cells in that gate
    fs_data <- data.frame(exprs(flowdata_adjustedmatrix[[i]])) 
    fluo_model <- PCAgrid(cbind(fs_data[,"FSC.A"], fs_data[,"FL1.A"]),k=2,method="sd",scores=FALSE,center="median")
    fs_data_rot <- rotmat(fluo_model = fluo_model, fsc = fs_data) #Do size correction
    fs_data_rot <- fs_data_rot %>% mutate(Strain = flowdata_adjustedmatrix[[i]]@description$GUID.original)
    flowdata_adjusted[[i]] <- fs_data_rot #This produces the datamatrix, in case we want it for future
    flowdata_adjustedmatrix[[i]]@exprs <- data.matrix(fs_data_rot)  # This acts on original matrix
    
  }
  return(flowdata_adjusted)
}
#provide flowdata frame from size_correction & genotype key
make_flatdf <- function(flow_df, key_df){ 
  bind_rows(flow_df, .id = "column_label") %>%
    group_by(Strain) %>% mutate(across(2:13, rev_log)) %>% 
    mutate(Strain = substr(Strain, start = 1, stop = 4)) %>%
    full_join(genotype_key)
} #This produces a flat dataframe that plays nicely with ggplot
flow_stat_summary <- function(flat_df){
  flat_df %>% 
    group_by(Strain, Promoter, Mutation) %>%
    summarise(count = n(),
              med_YFP = median(YFP.ROT),
              med_YFP.I = median(FL1.A),
              med_FSC.F = median(FSC.FINAL),
              med_FSC.A = median(FSC.A),
              mad_YFP = mad(YFP.ROT),
              mad_YFP.I = mad(FL1.A),
              mad_FSC.F = mad(FSC.FINAL),
              mad_FSC.A = mad(FSC.A)
    )
} #Calculates median and  median adjusted deviation 

### These are functions from imported packages to do log-transformation
rev_log <- function(x){
  y <- 10^x
  return(y)
} # We will use this to de-log transform the data; matter of taste
logTrans <- logTransform(transformationId="defaultLogTransform", logbase=10, r=1, d=1)
trans <- transformList(channels,logTrans)

### Plotting functions
flow_density <- function(flat_df, xmin = 1e2, xmax = 1e6){
  ggplot(flat_df, aes(x = YFP.ROT, fill = Mutation)) +
    geom_density(alpha = 0.5) + 
    scale_x_log10(limits = c(xmin, xmax),   
                  breaks = scales::trans_breaks("log10", function(x) 10^x),
                  labels = scales::trans_format("log10", scales::math_format(10^.x))) +
    theme_linedraw() + scale_color_npg() + scale_fill_npg() +
    labs(x = "Fluorescence\n (a.u)") +
    theme(panel.border = element_rect(linetype = 1, fill = "NA"),
          plot.background = element_rect(color = NA),legend.position="top",
          axis.text=element_text(size=14), axis.title=element_text(size =14),
          panel.grid.minor.x = element_line(color = 'black', linetype = 3),
          panel.grid.major.y = element_line( color = 'gray30')) 
} #make flow density; facet-able if necessary 
flow_replicate_variation <- function(summary_df, xmin = 1e2, xmax = 1e6){
  ggplot(summary_df, aes(x = med_YFP, y = Replicate, col = Mutation)) +
    geom_point(size =2, alpha = 0.5, position = position_dodge(width = 0.5)) + 
    geom_errorbar(aes(xmin = med_YFP - mad_YFP, 
                      xmax = med_YFP + mad_YFP), 
                  position = position_dodge(width = 0.5), width = 0.2) +
    scale_x_log10(limits = c(xmin, xmax),   
                  breaks = scales::trans_breaks("log10", function(x) 10^x),
                  labels = scales::trans_format("log10", scales::math_format(10^.x))) +
    theme_linedraw() + scale_color_npg() + scale_fill_npg() +
    labs(x = "Fluorescence\n (a.u)") +
    theme(panel.border = element_rect(linetype = 1, fill = "NA"),
          plot.background = element_rect(color = NA),legend.position="top",
          axis.text=element_text(size=14), axis.title=element_text(size =14),
          panel.grid.minor.x = element_line(color = 'black', linetype = 3),
          panel.grid.major.y = element_line( color = 'gray30')) + 
    facet_grid(Promoter~.)
  
}

### Gating-----
#discretized for each replicate batch. Left in this form for ease of altering gates for replicates individually.
### The reported data uses the same gates for all genotypes and replicates for consistency; if adjusting rg2 (changing name), make sure
### to adjust it in the size_correction function. 

#Replicate 1
fs_071420trans <- transform(fs071420, trans) # log transform for easier gating
gs_071420<-GatingSet(fs_071420trans) #create the empty gating set
rg1_071420 <- rectangleGate("FSC-A"=c(4.5, Inf), filterId = "NoneDebris")
gs_pop_add(gs_071420, rg1_071420, parent="root")
rg2 <- rectangleGate("FSC-W"=c(2.39, 2.48),"FSC-H"=c(4.8, 5.5))
gs_pop_add(gs_071420, rg2, parent = "NoneDebris", name = "singlets1") 
recompute(gs_071420)  #Calculates cells left after gating; hierarchy can be checked withgs_get_pop_paths(gs_071420) 

#Replicate 2
fs_071720trans <- transform(fs071720, trans) 
gs_071720<-GatingSet(fs_071720trans) 
rg1_071720 <- rectangleGate("FSC-A"=c(4.5, Inf), filterId = "NoneDebris")
gs_pop_add(gs_071720, rg1_071720, parent="root")
rg2 <- rectangleGate("FSC-W"=c(2.39, 2.48),"FSC-H"=c(4.8, 5.5))
gs_pop_add(gs_071720, rg2, parent = "NoneDebris", name = "singlets1") 
recompute(gs_071720)  

#Replicate 3
fs_072220trans <- transform(fs072220, trans) 
gs_072220<-GatingSet(fs_072220trans) 
rg1_072220 <- rectangleGate("FSC-A"=c(4.5, Inf), filterId = "NoneDebris")
gs_pop_add(gs_072220, rg1_072220, parent="root")
rg2 <- rectangleGate("FSC-W"=c(2.39, 2.48),"FSC-H"=c(4.8, 5.5))
gs_pop_add(gs_072220, rg2, parent = "NoneDebris", name = "singlets1") 
recompute(gs_072220)  

#How the gates partition the data can be visualized with the following 
# ggcyto(fs_071720trans, aes(x = "FSC-W", y = 'FSC-H'))+ geom_hex(bins = 512) +
#   ggcyto_par_set(limits = list(x = c(2.2, 2.9), y = c(4.25,6.0))) +
#   geom_gate(gs_pop_get_gate(gs_072220, "singlets1")) #rep3 shown for example


## Data Processing ----
## The data for each replicate is normalized to account for the relationship between cell size and YFP fluoresence and 
## made into a dataframe for easier plotting/analysis 
fs_071420_corrected <- size_correction(fs_071420trans) 
fs_071420_corrected_df <- make_flatdf(fs_071420_corrected) %>% mutate(Replicate = "July-14")
fs_071420_summary <- flow_stat_summary(fs_071420_corrected_df)%>% mutate(Replicate = "July-14")

fs_071720_corrected <- size_correction(fs_071720trans) 
fs_071720_corrected_df <- make_flatdf(fs_071720_corrected) %>% mutate(Replicate = "July-17")
fs_071720_summary <- flow_stat_summary(fs_071720_corrected_df)%>% mutate(Replicate = "July-17")

fs_072220_corrected <- size_correction(fs_072220trans) 
fs_072220_corrected_df <- make_flatdf(fs_072220_corrected) %>% mutate(Replicate = "July-22")
fs_072220_summary <- flow_stat_summary(fs_072220_corrected_df)%>% mutate(Replicate = "July-22")

# Combine everything --These two files have the total and summary data.
fs_data_full <- full_join(fs_071420_corrected_df, fs_071720_corrected_df) %>% full_join(fs_072220_corrected_df)
fs_data_fullsummary <- full_join(fs_071420_summary, fs_071720_summary) %>% full_join(fs_072220_summary)

fs_data_fullsummary_repseparate <- fs_data_fullsummary %>% #data.frame with 95% CIs
  group_by(Strain, Promoter, Mutation) %>%
  summarise(Fluorescence = mean(med_YFP),
            CI_95 = 1.96*sd(med_YFP)/sqrt(3))

## Visualizations
summary_density <- flow_density(fs_data_full)+ #7.6 x 7 ratio looks nice
  facet_grid(Promoter~.)


fs_data_fullsummary_repseparate$Mutation <- fct_relevel(fs_data_fullsummary_repseparate$Mutation, c("None", "ADE2", "ADE5", "ADE6"))
ggplot(fs_data_fullsummary_repseparate, aes(x = Promoter, y = Fluorescence, col = Mutation)) +
  geom_point(size = 2, position = position_dodge(width = 0.5)) + 
  geom_errorbar(aes(ymin = Fluorescence - CI_95, 
                    ymax = Fluorescence + CI_95),
                position = position_dodge(width = 0.5), width = 0.2) +
  scale_y_log10(limits = c(1e2, 1e6 ),   
                breaks = scales::trans_breaks("log10", function(x) 10^x),
                labels = scales::trans_format("log10", scales::math_format(10^.x))) +
  theme_linedraw() + scale_color_npg() + scale_fill_npg() +
  labs(x = "", y = "Fluorescence\n (a.u)") + 
  theme(panel.border = element_rect(linetype = 1, fill = "NA"),
        plot.background = element_rect(color = NA),legend.position="top",
        axis.text=element_text(size=14), axis.title=element_text(size =14),
        panel.grid.minor.x = element_line(color = 'black', linetype = 3),
        panel.grid.major.y = element_line( color = 'gray30'))
```
