## Supplementary Files 1-19 for "Mutational sources of *trans-*regulatory variation affecting gene expression in *Saccharomyces cerevisiae*": SupplementaryFile15.sh

```
###############################################################
#Supplementary File 15: PBS script used to process FASTQ files#
###############################################################


########################################################################################################################################
# 1 - Template used to generate each PBS file (where placeholders were replaced to read the FASTQ files corresponding to each sample). #
########################################################################################################################################

#!/bin/sh
#### Begin PBS preamble
#PBS -N Sample_$LANE_$FILE_1_$FILE_2
#
#PBS -M
#PBS -m abe
#
#PBS -l nodes=1:ppn=4,pmem=4gb
#PBS -l walltime=03:00:00
#PBS -j oe
#
#PBS -V
#PBS -A lsa_flux
#PBS -l qos=flux
#PBS -q flux
#### End PBS preamble
#
#
################################################################################
#The following labels need to be replaced appropriately for each sample:
#Iterate $$ITER,LINE$$
#Iterate $$ITER,EXPR,","$$
#Iterate $$END$$
#Replace $ITER
#Replace $LANE with the lane number (e.g. 45300, 45301, 45302)
#Replace $FILE_1 with the low bulk sample (e.g. 01,03,05,07)
#Replace $FILE_2 with the high bulk sample (e.g. 02,04,06,08)
#Replace $BARCODE.1 with the low bulk barcode sequence (e.g. TAAGGCGA-CTCTCTAT)
#Replace $BARCODE.2 with the high bulk barcode sequence (e.g. CGTACTAG-CTCTCTAT)
#Replace $MACHINE with machine number (e.g. 1 or 2)
################################################################################
#
echo Start
date +"%T"
#
#################################################
###$LANE_$FILE_1 Quality control and alignment###
#################################################
(
cd /scratch/lsa_flux/fduveau/Run.$LANE/Sample_$LANE_$FILE_1
###Quality Control###
#SICKLE
#Remove low quality bases from either end
#
$$ITER_1,LINE$$
sickle pe -f $LANE_$BARCODE.1_S$FILE_1_L00$MACHINE_R1_00$ITER.fastq.gz -r $LANE_$BARCODE.1_S$FILE_1_L00$MACHINE_R2_00$ITER.fastq.gz -t sanger -o $LANE_$FILE_1_$ITER_F_CLIP.fastq -p $LANE_$FILE_1_$ITER_R_CLIP.fastq -s $LANE_$FILE_1_$ITER_S_CLIP.fastq
$$END$$
#
echo $LANE_$FILE_1 Sickle Finished
date +"%T"
#
#CUTADAPT
#Remove adapter sequences
#
$$ITER_1,LINE$$
cutadapt -g AATGATACGGCGACCACCGAGATCTACACNNNNNNNNTCGTCGGCAGCGTCAGATGTGTATAAGAGACAG -a CTGTCTCTTATACACATCTCCGAGCCCACGAGACNNNNNNNNATCTCGTATGCCGTCTTCTGCTTG     -e 0.2 -O 3 -m 15 -o $LANE_$FILE_1.tmp.1.fastq -p $LANE_$FILE_1.tmp.2.fastq $LANE_$FILE_1_$ITER_F_CLIP.fastq $LANE_$FILE_1_$ITER_R_CLIP.fastq
cutadapt -g CAAGCAGAAGACGGCATACGAGATNNNNNNNNGTCTCGTGGGCTCGGAGATGTGTATAAGAGACAG     -a AATGATACGGCGACCACCGAGATCTACACNNNNNNNNTCGTCGGCAGCGTCAGATGTGTATAAGAGACAG -e 0.2 -O 6 -m 15 -o $LANE_$FILE_1_$ITER_F_ADAP.fastq -p $LANE_$FILE_1_$ITER_R_ADAP.fastq $LANE_$FILE_1.tmp.2.fastq $LANE_$FILE_1.tmp.1.fastq
$$END$$
#
echo $LANE_$FILE_1 CutAdapt Finished
date +"%T"
#
###ALIGNMENT###
#BOWTIE2
#Align to reference genome
#
bowtie2 -I 0 -X 1000 -t --rg-id $LANE_$FILE_1 --rg SM:L -q --phred33 --very-sensitive --score-min L,-0.6,-0.6 -p 8 -x /scratch/lsa_flux/fduveau/Reference.Genome/S288c.mapping -1 $$ITER_1,EXPR,","$$$LANE_$FILE_1_$ITER_F_ADAP.fastq$$END$$ -2 $$ITER_1,EXPR,","$$$LANE_$FILE_1_$ITER_R_ADAP.fastq$$END$$ -S $LANE_$FILE_1.sam
#
echo $LANE_$FILE_1 Bowtie Finished
date +"%T"
#
#SAMTOOLS
#Convert sam to bam, sort and index
samtools view -hb -o $LANE_$FILE_1.bam $LANE_$FILE_1.sam
samtools sort -o $LANE_$FILE_1.SORT.bam -O bam -T $LANE_$FILE_1.temp1 $LANE_$FILE_1.bam
samtools index $LANE_$FILE_1.SORT.bam
#
echo $LANE_$FILE_1 Samtools Finished
date +"%T"
#
#CLIPOVERLAP
#Remove overlapping portions of paired end reads
bam clipOverlap --in $LANE_$FILE_1.SORT.bam --out $LANE_$FILE_1.OVER.bam --stats
samtools index $LANE_$FILE_1.OVER.bam
#
echo $LANE_$FILE_1 ClipOverlap Finished
date +"%T"
#
) &
#
#
#################################################
###$LANE_$FILE_2 Quality control and alignment###
#################################################
(
cd /scratch/lsa_flux/fduveau/Run.$LANE/Sample_$LANE_$FILE_2
###Quality Control
#SICKLE
#Remove low quality bases from either end
$$ITER_2,LINE$$
sickle pe -f $LANE_$BARCODE.2_S$FILE_2_L00$MACHINE_R1_00$ITER.fastq.gz -r $LANE_$BARCODE.2_S$FILE_2_L00$MACHINE_R2_00$ITER.fastq.gz -t sanger -o $LANE_$FILE_2_$ITER_F_CLIP.fastq -p $LANE_$FILE_2_$ITER_R_CLIP.fastq -s $LANE_$FILE_2_$ITER_S_CLIP.fastq
$$END$$
#
echo $LANE_$FILE_2 Sickle Finished
date +"%T"
#
#CUTADAPT
#Remove adapater sequences
$$ITER_2,LINE$$
cutadapt -g AATGATACGGCGACCACCGAGATCTACACNNNNNNNNTCGTCGGCAGCGTCAGATGTGTATAAGAGACAG -a CTGTCTCTTATACACATCTCCGAGCCCACGAGACNNNNNNNNATCTCGTATGCCGTCTTCTGCTTG     -e 0.2 -O 3 -m 15 -o $LANE_$FILE_2.tmp.1.fastq -p $LANE_$FILE_2.tmp.2.fastq $LANE_$FILE_2_$ITER_F_CLIP.fastq $LANE_$FILE_2_$ITER_R_CLIP.fastq
cutadapt -g CAAGCAGAAGACGGCATACGAGATNNNNNNNNGTCTCGTGGGCTCGGAGATGTGTATAAGAGACAG     -a AATGATACGGCGACCACCGAGATCTACACNNNNNNNNTCGTCGGCAGCGTCAGATGTGTATAAGAGACAG -e 0.2 -O 6 -m 15 -o $LANE_$FILE_2_$ITER_F_ADAP.fastq -p $LANE_$FILE_2_$ITER_R_ADAP.fastq $LANE_$FILE_2.tmp.2.fastq $LANE_$FILE_2.tmp.1.fastq
$$END$$
#
echo $LANE_$FILE_2 CutAdapt Finished
date +"%T"
#
###ALIGNMENT
#BOWTIE2
#Align to reference genome
bowtie2 -I 0 -X 1000 -t --rg-id $LANE_$FILE_2 --rg SM:H -q --phred33 --very-sensitive --score-min L,-0.6,-0.6 -p 8 -x /scratch/lsa_flux/fduveau/Reference.Genome/S288c.mapping -1 $$ITER_2,EXPR,","$$$LANE_$FILE_2_$ITER_F_ADAP.fastq$$END$$ -2 $$ITER_2,EXPR,","$$$LANE_$FILE_2_$ITER_R_ADAP.fastq$$END$$ -S $LANE_$FILE_2.sam
#
echo $LANE_$FILE_2 Bowtie Finished
date +"%T"
#
#SAMTOOLS
#Convert sam to bam, sort and index
samtools view -hb -o $LANE_$FILE_2.bam $LANE_$FILE_2.sam
samtools sort -o $LANE_$FILE_2.SORT.bam -O bam -T $LANE_$FILE_2.temp1 $LANE_$FILE_2.bam
samtools index $LANE_$FILE_2.SORT.bam
#
echo $LANE_$FILE_2 Samtools Finished
date +"%T"
#
#CLIPOVERLAP
#Remove overlapping portions of paired end reads
bam clipOverlap --in $LANE_$FILE_2.SORT.bam --out $LANE_$FILE_2.OVER.bam --stats
samtools index $LANE_$FILE_2.OVER.bam
#
echo $LANE_$FILE_2 ClipOverlap Finished
date +"%T"
#
) &
#
#
wait
#
echo Freebayes Start
date +"%T"
#
##################
###SNP CALLING####
##################
#FREEBAYES
#Call SNPs and get allele frequencies
cd /scratch/lsa_flux/fduveau/Run.$LANE
freebayes -f /scratch/lsa_flux/fduveau/Reference.Genome/S288c.mapping.fsa --use-best-n-alleles 2 --min-alternate-count 4 --min-alternate-fraction 0.1 --pooled-discrete --pooled-continuous /scratch/lsa_flux/fduveau/Run.$LANE/Sample_$LANE_$FILE_1/$LANE_$FILE_1.OVER.bam /scratch/lsa_flux/fduveau/Run.$LANE/Sample_$LANE_$FILE_2/$LANE_$FILE_2.OVER.bam > $LANE_$FILE_1_$FILE_2.vcf
#
echo Freebayes Finished
date +"%T"
#
#######################################
###$LANE_$FILE_1 Frequency and Stats###
#######################################
(
cd /scratch/lsa_flux/fduveau/Run.$LANE/Sample_$LANE_$FILE_1
###Frequency Estimates
#SAMTOOLS
#piluep file
samtools mpileup -B $LANE_$FILE_1.OVER.bam > $LANE_$FILE_1.mpileup
#
wait
#
echo $LANE_$FILE_1 mpileup Finished
date +"%T"
#
echo $LANE_$FILE_1 popoolation Finished
date +"%T"
#
###STATISTICS###
#SAMTOOLS
#Insert and Fragment sizes
samtools view -f66 $LANE_$FILE_1.bam | cut -f 9 | sed 's/^-//' > FragSize.$LANE_$FILE_1.txt
samtools view -f66 $LANE_$FILE_1.OVER.bam | cut -f 9 | sed 's/^-//' > InsertSize.$LANE_$FILE_1.txt
#
#BEDTOOLS
#Coverage at each site
bedtools genomecov -d -ibam $LANE_$FILE_1.OVER.bam > Coverage.$LANE_$FILE_1.txt
#
echo $LANE_$FILE_1 Stats Finished
date +"%T"
) &
#
#
#######################################
###$LANE_$FILE_2 Frequency and Stats###
#######################################
(
cd /scratch/lsa_flux/fduveau/Run.$LANE/Sample_$LANE_$FILE_2
###Frequency Estimates
#SAMTOOLS
#piluep file
samtools mpileup -B $LANE_$FILE_2.OVER.bam > $LANE_$FILE_2.mpileup
#
wait
#
echo $LANE_$FILE_2 mpileup Finished
date +"%T"
#
echo $LANE_$FILE_2 popoolation Finished
date +"%T"
#
###STATISTICS###
#SAMTOOLS
#Insert and Fragment sizes
samtools view -f66 $LANE_$FILE_2.bam | cut -f 9 | sed 's/^-//' > FragSize.$LANE_$FILE_2.txt
samtools view -f66 $LANE_$FILE_2.OVER.bam | cut -f 9 | sed 's/^-//' > InsertSize.$LANE_$FILE_2.txt
#
#BEDTOOLS
#Coverage at each site
bedtools genomecov -d -ibam $LANE_$FILE_2.OVER.bam > Coverage.$LANE_$FILE_2.txt
#
echo $LANE_$FILE_2 Stats Finished
date +"%T"
#
) &
#
wait
#
echo Stop
date +"%T"
#


##########################################################################################################################################
# 2 - Example: PBS script used to analyze libraries 01 and 02 in sequencing run 54374, corresponding to BSA-Seq data for mutant YPW2200. #
##########################################################################################################################################

#!/bin/sh
#### Begin PBS preamble
#PBS -N Sample_54374_01_02
#
#PBS -M
#PBS -m abe
#
#PBS -l nodes=1:ppn=4,pmem=4gb
#PBS -l walltime=03:00:00
#PBS -j oe
#
#PBS -V
#PBS -A lsa_flux
#PBS -l qos=flux
#PBS -q flux
#### End PBS preamble
#
#
echo Start
date +"%T"
#
#################################################
###54374_01 Quality control and alignment###
#################################################
(
cd /scratch/lsa_flux/fduveau/Run.54374/Sample_54374_01
###Quality Control###
#SICKLE
#Remove low quality bases from either end
#
sickle pe -f 54374_TAAGGCGA-AAGGAGTA_S01_L002_R1_001.fastq.gz -r 54374_TAAGGCGA-AAGGAGTA_S01_L002_R2_001.fastq.gz -t sanger -o 54374_01_1_F_CLIP.fastq -p 54374_01_1_R_CLIP.fastq -s 54374_01_1_S_CLIP.fastq
#
echo 54374_01 Sickle Finished
date +"%T"
#
#CUTADAPT
#Remove adapater sequences
#
cutadapt -g AATGATACGGCGACCACCGAGATCTACACNNNNNNNNTCGTCGGCAGCGTCAGATGTGTATAAGAGACAG -a CTGTCTCTTATACACATCTCCGAGCCCACGAGACNNNNNNNNATCTCGTATGCCGTCTTCTGCTTG     -e 0.2 -O 3 -m 15 -o 54374_01.tmp.1.fastq -p 54374_01.tmp.2.fastq 54374_01_1_F_CLIP.fastq 54374_01_1_R_CLIP.fastq
cutadapt -g CAAGCAGAAGACGGCATACGAGATNNNNNNNNGTCTCGTGGGCTCGGAGATGTGTATAAGAGACAG     -a AATGATACGGCGACCACCGAGATCTACACNNNNNNNNTCGTCGGCAGCGTCAGATGTGTATAAGAGACAG -e 0.2 -O 6 -m 15 -o 54374_01_1_F_ADAP.fastq -p 54374_01_1_R_ADAP.fastq 54374_01.tmp.2.fastq 54374_01.tmp.1.fastq
#
echo 54374_01 CutAdapt Finished
date +"%T"
#
###ALIGNMENT###
#BOWTIE2
#Align to reference genome
#
bowtie2 -I 0 -X 1000 -t --rg-id 54374_01 --rg SM:L -q --phred33 --very-sensitive --score-min L,-0.6,-0.6 -p 8 -x /scratch/lsa_flux/fduveau/Reference.Genome/S288c.mapping -1 54374_01_1_F_ADAP.fastq -2 54374_01_1_R_ADAP.fastq -S 54374_01.sam
#
echo 54374_01 Bowtie Finished
date +"%T"
#
#SAMTOOLS
#Convert sam to bam, sort and index
samtools view -hb -o 54374_01.bam 54374_01.sam
samtools sort -o 54374_01.SORT.bam -O bam -T 54374_01.temp1 54374_01.bam
samtools index 54374_01.SORT.bam
#
echo 54374_01 Samtools Finished
date +"%T"
#
#CLIPOVERLAP
#Remove overlapping portions of paired end reads
bam clipOverlap --in 54374_01.SORT.bam --out 54374_01.OVER.bam --stats
samtools index 54374_01.OVER.bam
#
echo 54374_01 ClipOverlap Finished
date +"%T"
#
) &
#
#
#################################################
###54374_02 Quality control and alignment###
#################################################
(
cd /scratch/lsa_flux/fduveau/Run.54374/Sample_54374_02
###Quality Control
#SICKLE
#Remove low quality bases from either end
sickle pe -f 54374_CGTACTAG-AAGGAGTA_S02_L002_R1_001.fastq.gz -r 54374_CGTACTAG-AAGGAGTA_S02_L002_R2_001.fastq.gz -t sanger -o 54374_02_1_F_CLIP.fastq -p 54374_02_1_R_CLIP.fastq -s 54374_02_1_S_CLIP.fastq
#
echo 54374_02 Sickle Finished
date +"%T"
#
#CUTADAPT
#Remove adapater sequences
cutadapt -g AATGATACGGCGACCACCGAGATCTACACNNNNNNNNTCGTCGGCAGCGTCAGATGTGTATAAGAGACAG -a CTGTCTCTTATACACATCTCCGAGCCCACGAGACNNNNNNNNATCTCGTATGCCGTCTTCTGCTTG     -e 0.2 -O 3 -m 15 -o 54374_02.tmp.1.fastq -p 54374_02.tmp.2.fastq 54374_02_1_F_CLIP.fastq 54374_02_1_R_CLIP.fastq
cutadapt -g CAAGCAGAAGACGGCATACGAGATNNNNNNNNGTCTCGTGGGCTCGGAGATGTGTATAAGAGACAG     -a AATGATACGGCGACCACCGAGATCTACACNNNNNNNNTCGTCGGCAGCGTCAGATGTGTATAAGAGACAG -e 0.2 -O 6 -m 15 -o 54374_02_1_F_ADAP.fastq -p 54374_02_1_R_ADAP.fastq 54374_02.tmp.2.fastq 54374_02.tmp.1.fastq
#
echo 54374_02 CutAdapt Finished
date +"%T"
#
###ALIGNMENT
#BOWTIE2
#Align to reference genome
bowtie2 -I 0 -X 1000 -t --rg-id 54374_02 --rg SM:H -q --phred33 --very-sensitive --score-min L,-0.6,-0.6 -p 8 -x /scratch/lsa_flux/fduveau/Reference.Genome/S288c.mapping -1 54374_02_1_F_ADAP.fastq -2 54374_02_1_R_ADAP.fastq -S 54374_02.sam
#
echo 54374_02 Bowtie Finished
date +"%T"
#
#SAMTOOLS
#Convert sam to bam, sort and index
samtools view -hb -o 54374_02.bam 54374_02.sam
samtools sort -o 54374_02.SORT.bam -O bam -T 54374_02.temp1 54374_02.bam
samtools index 54374_02.SORT.bam
#
echo 54374_02 Samtools Finished
date +"%T"
#
#CLIPOVERLAP
#Remove overlapping portions of paired end reads
bam clipOverlap --in 54374_02.SORT.bam --out 54374_02.OVER.bam --stats
samtools index 54374_02.OVER.bam
#
echo 54374_02 ClipOverlap Finished
date +"%T"
#
) &
#
#
wait
#
echo Freebayes Start
date +"%T"
#
##################
###SNP CALLING####
##################
#FREEBAYES
#Call SNPs and get allele frequencies
cd /scratch/lsa_flux/fduveau/Run.54374
freebayes -f /scratch/lsa_flux/fduveau/Reference.Genome/S288c.mapping.fsa --use-best-n-alleles 2 --min-alternate-count 4 --min-alternate-fraction 0.1 --pooled-discrete --pooled-continuous /scratch/lsa_flux/fduveau/Run.54374/Sample_54374_01/54374_01.OVER.bam /scratch/lsa_flux/fduveau/Run.54374/Sample_54374_02/54374_02.OVER.bam > 54374_01_02.vcf
#
echo Freebayes Finished
date +"%T"
#
#######################################
###54374_01 Frequency and Stats###
#######################################
(
cd /scratch/lsa_flux/fduveau/Run.54374/Sample_54374_01
###Frequency Estimates
#SAMTOOLS
#piluep file
samtools mpileup -B 54374_01.OVER.bam > 54374_01.mpileup
#
wait
#
echo 54374_01 mpileup Finished
date +"%T"
#
echo 54374_01 popoolation Finished
date +"%T"
#
###STATISTICS###
#SAMTOOLS
#Insert and Fragment sizes
samtools view -f66 54374_01.bam | cut -f 9 | sed 's/^-//' > FragSize.54374_01.txt
samtools view -f66 54374_01.OVER.bam | cut -f 9 | sed 's/^-//' > InsertSize.54374_01.txt
#
#BEDTOOLS
#Coverage at each site
bedtools genomecov -d -ibam 54374_01.OVER.bam > Coverage.54374_01.txt
#
echo 54374_01 Stats Finished
date +"%T"
) &
#
#
#######################################
###54374_02 Frequency and Stats###
#######################################
(
cd /scratch/lsa_flux/fduveau/Run.54374/Sample_54374_02
###Frequency Estimates
#SAMTOOLS
#piluep file
samtools mpileup -B 54374_02.OVER.bam > 54374_02.mpileup
#
wait
#
echo 54374_02 mpileup Finished
date +"%T"
#
echo 54374_02 popoolation Finished
date +"%T"
#
###STATISTICS###
#SAMTOOLS
#Insert and Fragment sizes
samtools view -f66 54374_02.bam | cut -f 9 | sed 's/^-//' > FragSize.54374_02.txt
samtools view -f66 54374_02.OVER.bam | cut -f 9 | sed 's/^-//' > InsertSize.54374_02.txt
#
#BEDTOOLS
#Coverage at each site
bedtools genomecov -d -ibam 54374_02.OVER.bam > Coverage.54374_02.txt
#
echo 54374_02 Stats Finished
date +"%T"
#
) &
#
wait
#
echo Stop
```
