## Supplementary Files 1-19 for "Mutational sources of *trans-*regulatory variation affecting gene expression in *Saccharomyces cerevisiae*": SupplementaryFile16.R

```
######################################################################################################################################
################################################### Supplementary File 16 ############################################################
######################################################################################################################################
# R scripts used for the analysis of BSA-Seq data and for comparing the properties of trans-regulatory and non-regulatory mutations. #
######################################################################################################################################

#A) Computing the average sequencing coverage for each chromosome in each sample.                                   ===> Line 36
#B) Filtering out false positive variant calls in vcf files to identify mutations in each mutant.                   ===> Line 266
#C) Likelihood-ratio tests (G-tests) to compare allele frequencies between low and high fluorescence bulks.         ===> Line 441
#D) Plotting the number of mutations identified per strain (Figure 2 - figure supplement 1).                        ===> Line 465
#E) Plotting expression changes in EMS mutants depending on the number of mutations associated with fluorescence    ===> Line 578
#   in BSA-Seq data (Figure 2 - figure supplement 2).
#F) Relationship between the total number of mutations and the absolute expression change                           ===> Line 640
#   among EMS mutants (Figure 2 - figure supplement 3). 
#G) Comparing the strengths of association with fluorescence (G-values) of genetically linked mutations             ===> Line 678
#   (Supplementary File 3).
#H) Types of trans-regulatory and non-regulatory mutations identified (Figure 3A).                                  ===> Line 961
#I) Genomic distributions of trans-regulatory and non-regulatory mutations (Figure 3B).                             ===> Line 1160
#J) Proportion of mutations observed on each chromosome (Figure 3 - figure supplement 1).                           ===> Line 1230
#K) Comparing the proportions of trans-regulatory and non regulatory mutations in coding, intergenic                ===> Line 1336
#   and intronic regions (Figure 3C).
#L) Comparing the proportions of trans-regulatory and non-regulatory mutations being synonymous,                    ===> Line 1400
#   nonsynonymous and nonsense mutations (Figure 3D).#
#M) Frequencies of amino acid changes caused by non-regulatory and trans-regulatory mutations                       ===> Line 1505
#   (Figure 3E, Figure 3- figure supplement 2)
#N) Testing for enrichment of trans-regulatory mutations in genes encoding transcription factors.                   ===> Line 1711
#O) Testing for enrichment of trans-regulatory mutations in a predicted TDH3 regulatory network (Figure 4).         ===> Line 1796
#P) Distributions of the number of mutations in GCR1 or RAP1 regions targeted for PCR mutagenesis (Figure 5C,D).    ===> Line 1930
#Q) Number of independent mutations identified in each gene (Figure 6A).                                            ===> Line 2089
#R) Enriched terms in gene ontology analysis (Figure 6B).                                                           ===> Line 2166
#S) Genomic overlap between trans-regulatory mutations and eQTL regions (Figure 7B).                                ===> Line 2192
#T) Proportions of non-regulatory and trans-regulatory mutations located in eQTL regions (Figure 7B).               ===> Line 2386
#U) Statistical tests for enrichment of trans-regulatory mutations in eQTL regions.                                 ===> Line 2571

##################################################################################
#A) Computing the average sequencing coverage for each chromosome in each sample.#
##################################################################################

###Clear memory###
rm(list=ls())
options(warn=-1)

###Load packages###
library(VariantAnnotation)
library(Deducer)
library(gtools)
library(zoo)
library(Hmisc)

###Set directory###
setwd("/Path.to.input.files")

###Input files###

#List of samples that can be found in SupplementaryFile11.zip
SAMPLES <- read.table("BSA_Samples.txt",header=TRUE) 

#Length of chromosomes analyzed that can be found in SupplementaryFile11.zip
CHROMOSOMES <- read.table("Chromosome_Length.txt",header=TRUE)

#All files generated with "SupplementaryFile15.sh" script must be placed in "/Path.to.input.files" in order to be listed below:
VCF.FILES <- list.files("VCF.2",pattern="LIBERAL",recursive=TRUE,full.names=TRUE)
COVERAGE.FILES <- list.files("Coverage",pattern="Coverage",recursive=TRUE,full.names=TRUE)
FRAGMENT.FILES <- list.files("FragSize",pattern="Frag",recursive=TRUE,full.names=TRUE)
SYNC.FILES <- list.files("Sync",pattern="sync",recursive=TRUE,full.names=TRUE)

#Variables used later.
N <- 1:(length(VCF.FILES))
UB <- numeric(length(N))
LB <- numeric(length(N))
OUTPUT <- list()
DATA.G <- list()
G.PRIME <- list()
DATA.G.PRIME <- list()
SEG <- list()
FIXED <- list()
FRAGMENT.LOW <- list()
FRAGMENT.HIGH <- list()
STATS <- list()


#Save basic coverage info for each sample.

SAMPLES[,"MEDIAN.COV.L"] <- 0
SAMPLES[,"MEDIAN.COV.H"] <- 0
SAMPLES[,"IQR.COV.L"] <- 0
SAMPLES[,"IQR.COV.H"] <- 0
SAMPLES[,"MAD.COV.L"] <- 0
SAMPLES[,"MAD.COV.H"] <- 0
SAMPLES[,"MEDIAN.COV.CHR.1.L"] <- 0
SAMPLES[,"MEDIAN.COV.CHR.1.H"] <- 0
SAMPLES[,"MEDIAN.COV.CHR.2.L"] <- 0
SAMPLES[,"MEDIAN.COV.CHR.2.H"] <- 0
SAMPLES[,"MEDIAN.COV.CHR.3.L"] <- 0
SAMPLES[,"MEDIAN.COV.CHR.3.H"] <- 0
SAMPLES[,"MEDIAN.COV.CHR.4.L"] <- 0
SAMPLES[,"MEDIAN.COV.CHR.4.H"] <- 0
SAMPLES[,"MEDIAN.COV.CHR.5.L"] <- 0
SAMPLES[,"MEDIAN.COV.CHR.5.H"] <- 0
SAMPLES[,"MEDIAN.COV.CHR.6.L"] <- 0
SAMPLES[,"MEDIAN.COV.CHR.6.H"] <- 0
SAMPLES[,"MEDIAN.COV.CHR.7.L"] <- 0
SAMPLES[,"MEDIAN.COV.CHR.7.H"] <- 0
SAMPLES[,"MEDIAN.COV.CHR.8.L"] <- 0
SAMPLES[,"MEDIAN.COV.CHR.8.H"] <- 0
SAMPLES[,"MEDIAN.COV.CHR.9.L"] <- 0
SAMPLES[,"MEDIAN.COV.CHR.9.H"] <- 0
SAMPLES[,"MEDIAN.COV.CHR.10.L"] <- 0
SAMPLES[,"MEDIAN.COV.CHR.10.H"] <- 0
SAMPLES[,"MEDIAN.COV.CHR.11.L"] <- 0
SAMPLES[,"MEDIAN.COV.CHR.11.H"] <- 0
SAMPLES[,"MEDIAN.COV.CHR.12.L"] <- 0
SAMPLES[,"MEDIAN.COV.CHR.12.H"] <- 0
SAMPLES[,"MEDIAN.COV.CHR.13.L"] <- 0
SAMPLES[,"MEDIAN.COV.CHR.13.H"] <- 0
SAMPLES[,"MEDIAN.COV.CHR.14.L"] <- 0
SAMPLES[,"MEDIAN.COV.CHR.14.H"] <- 0
SAMPLES[,"MEDIAN.COV.CHR.15.L"] <- 0
SAMPLES[,"MEDIAN.COV.CHR.15.H"] <- 0
SAMPLES[,"MEDIAN.COV.CHR.16.L"] <- 0
SAMPLES[,"MEDIAN.COV.CHR.16.H"] <- 0
SAMPLES[,"N.READS.CHR.1.L"] <- 0
SAMPLES[,"N.READS.CHR.1.H"] <- 0
SAMPLES[,"N.READS.CHR.2.L"] <- 0
SAMPLES[,"N.READS.CHR.2.H"] <- 0
SAMPLES[,"N.READS.CHR.3.L"] <- 0
SAMPLES[,"N.READS.CHR.3.H"] <- 0
SAMPLES[,"N.READSCHR.4.L"] <- 0
SAMPLES[,"N.READS.CHR.4.H"] <- 0
SAMPLES[,"N.READS.CHR.5.L"] <- 0
SAMPLES[,"N.READS.CHR.5.H"] <- 0
SAMPLES[,"N.READS.CHR.6.L"] <- 0
SAMPLES[,"N.READS.CHR.6.H"] <- 0
SAMPLES[,"N.READS.CHR.7.L"] <- 0
SAMPLES[,"N.READS.CHR.7.H"] <- 0
SAMPLES[,"N.READS.CHR.8.L"] <- 0
SAMPLES[,"N.READS.CHR.8.H"] <- 0
SAMPLES[,"N.READS.CHR.9.L"] <- 0
SAMPLES[,"N.READS.CHR.9.H"] <- 0
SAMPLES[,"N.READS.CHR.10.L"] <- 0
SAMPLES[,"N.READS.CHR.10.H"] <- 0
SAMPLES[,"N.READS.CHR.11.L"] <- 0
SAMPLES[,"N.READS.CHR.11.H"] <- 0
SAMPLES[,"N.READS.CHR.12.L"] <- 0
SAMPLES[,"N.READS.CHR.12.H"] <- 0
SAMPLES[,"N.READS.CHR.13.L"] <- 0
SAMPLES[,"N.READS.CHR.13.H"] <- 0
SAMPLES[,"N.READS.CHR.14.L"] <- 0
SAMPLES[,"N.READS.CHR.14.H"] <- 0
SAMPLES[,"N.READS.CHR.15.L"] <- 0
SAMPLES[,"N.READS.CHR.15.H"] <- 0
SAMPLES[,"N.READS.CHR.16.L"] <- 0
SAMPLES[,"N.READS.CHR.16.H"] <- 0
SAMPLES[,"N.READS.GENOME.L"] <- 0
SAMPLES[,"N.READS.GENOME.H"] <- 0

COVERAGE.FILES <-  c(COVERAGE.FILES,COVERAGE.FILES[37:45])
COVERAGE.FILES <-  COVERAGE.FILES[-c(37:45)]


for (i in 37:length(COVERAGE.FILES))
{
	CUR.COVERAGE  <- read.table(COVERAGE.FILES[i],header=FALSE)
	
	SAMPLES[ceiling(i/2),2-i%%2+3] <- median(CUR.COVERAGE[,3])
	SAMPLES[ceiling(i/2),2-i%%2+5] <- IQR(CUR.COVERAGE[,3])
	SAMPLES[ceiling(i/2),2-i%%2+7] <- mad(CUR.COVERAGE[,3])
	
	SAMPLES[ceiling(i/2),2-i%%2+9] <- median(CUR.COVERAGE[which(CUR.COVERAGE[,1] == "chr01"),3])
	SAMPLES[ceiling(i/2),2-i%%2+11] <- median(CUR.COVERAGE[which(CUR.COVERAGE[,1] == "chr02"),3])
	SAMPLES[ceiling(i/2),2-i%%2+13] <- median(CUR.COVERAGE[which(CUR.COVERAGE[,1] == "chr03"),3])
	SAMPLES[ceiling(i/2),2-i%%2+15] <- median(CUR.COVERAGE[which(CUR.COVERAGE[,1] == "chr04"),3])
	SAMPLES[ceiling(i/2),2-i%%2+17] <- median(CUR.COVERAGE[which(CUR.COVERAGE[,1] == "chr05"),3])
	SAMPLES[ceiling(i/2),2-i%%2+19] <- median(CUR.COVERAGE[which(CUR.COVERAGE[,1] == "chr06"),3])
	SAMPLES[ceiling(i/2),2-i%%2+21] <- median(CUR.COVERAGE[which(CUR.COVERAGE[,1] == "chr07"),3])
	SAMPLES[ceiling(i/2),2-i%%2+23] <- median(CUR.COVERAGE[which(CUR.COVERAGE[,1] == "chr08"),3])
	SAMPLES[ceiling(i/2),2-i%%2+25] <- median(CUR.COVERAGE[which(CUR.COVERAGE[,1] == "chr09"),3])
	SAMPLES[ceiling(i/2),2-i%%2+27] <- median(CUR.COVERAGE[which(CUR.COVERAGE[,1] == "chr10"),3])
	SAMPLES[ceiling(i/2),2-i%%2+29] <- median(CUR.COVERAGE[which(CUR.COVERAGE[,1] == "chr11"),3])
	SAMPLES[ceiling(i/2),2-i%%2+31] <- median(CUR.COVERAGE[which(CUR.COVERAGE[,1] == "chr12"),3])
	SAMPLES[ceiling(i/2),2-i%%2+33] <- median(CUR.COVERAGE[which(CUR.COVERAGE[,1] == "chr13"),3])
	SAMPLES[ceiling(i/2),2-i%%2+35] <- median(CUR.COVERAGE[which(CUR.COVERAGE[,1] == "chr14"),3])
	SAMPLES[ceiling(i/2),2-i%%2+37] <- median(CUR.COVERAGE[which(CUR.COVERAGE[,1] == "chr15"),3])
	SAMPLES[ceiling(i/2),2-i%%2+39] <- median(CUR.COVERAGE[which(CUR.COVERAGE[,1] == "chr16"),3])
	
	SAMPLES[ceiling(i/2),2-i%%2+41] <- sum(CUR.COVERAGE[which(CUR.COVERAGE[,1] == "chr01"),3])/300
	SAMPLES[ceiling(i/2),2-i%%2+43] <- sum(CUR.COVERAGE[which(CUR.COVERAGE[,1] == "chr02"),3])/300
	SAMPLES[ceiling(i/2),2-i%%2+45] <- sum(CUR.COVERAGE[which(CUR.COVERAGE[,1] == "chr03"),3])/300
	SAMPLES[ceiling(i/2),2-i%%2+47] <- sum(CUR.COVERAGE[which(CUR.COVERAGE[,1] == "chr04"),3])/300
	SAMPLES[ceiling(i/2),2-i%%2+49] <- sum(CUR.COVERAGE[which(CUR.COVERAGE[,1] == "chr05"),3])/300
	SAMPLES[ceiling(i/2),2-i%%2+51] <- sum(CUR.COVERAGE[which(CUR.COVERAGE[,1] == "chr06"),3])/300
	SAMPLES[ceiling(i/2),2-i%%2+53] <- sum(CUR.COVERAGE[which(CUR.COVERAGE[,1] == "chr07"),3])/300
	SAMPLES[ceiling(i/2),2-i%%2+55] <- sum(CUR.COVERAGE[which(CUR.COVERAGE[,1] == "chr08"),3])/300
	SAMPLES[ceiling(i/2),2-i%%2+57] <- sum(CUR.COVERAGE[which(CUR.COVERAGE[,1] == "chr09"),3])/300
	SAMPLES[ceiling(i/2),2-i%%2+59] <- sum(CUR.COVERAGE[which(CUR.COVERAGE[,1] == "chr10"),3])/300
	SAMPLES[ceiling(i/2),2-i%%2+61] <- sum(CUR.COVERAGE[which(CUR.COVERAGE[,1] == "chr11"),3])/300
	SAMPLES[ceiling(i/2),2-i%%2+63] <- sum(CUR.COVERAGE[which(CUR.COVERAGE[,1] == "chr12"),3])/300
	SAMPLES[ceiling(i/2),2-i%%2+65] <- sum(CUR.COVERAGE[which(CUR.COVERAGE[,1] == "chr13"),3])/300
	SAMPLES[ceiling(i/2),2-i%%2+67] <- sum(CUR.COVERAGE[which(CUR.COVERAGE[,1] == "chr14"),3])/300
	SAMPLES[ceiling(i/2),2-i%%2+69] <- sum(CUR.COVERAGE[which(CUR.COVERAGE[,1] == "chr15"),3])/300
	SAMPLES[ceiling(i/2),2-i%%2+71] <- sum(CUR.COVERAGE[which(CUR.COVERAGE[,1] == "chr16"),3])/300
	
	SAMPLES[ceiling(i/2),2-i%%2+73] <- sum(CUR.COVERAGE[,3])/300
}

SAMPLES[,42:ncol(SAMPLES)] <- round(SAMPLES[,42:ncol(SAMPLES)])

#Data contained in SAMPLES were used to make Table 1.
write.table(SAMPLES,"COVERAGE.SUMMARY.txt",row.names=FALSE,sep="\t",quote=FALSE)

#Computing G.values for the three aneuploidies by comparing sequencing coverage of extra chromosomes in low and high fluorescence bulks.
library(metaseqR)

G.MATRIX <- matrix(c(105388/2,71741/2,3778639/2,3373122/2),nrow=2,ncol=2)
G.TEST <- likelihood.test(G.MATRIX,conservative=TRUE)
G.TEST$statistic
pchisq(G.TEST$statistic,df=1,lower.tail=FALSE)
-log10(pchisq(G.TEST$statistic,df=1,lower.tail=FALSE))

G.MATRIX <- matrix(c(316588/2,207671/2,4237427/2,3781878/2),nrow=2,ncol=2)
G.TEST <- likelihood.test(G.MATRIX,conservative=TRUE)
G.TEST$statistic
pchisq(G.TEST$statistic,df=1,lower.tail=FALSE)
-log10(pchisq(G.TEST$statistic,df=1,lower.tail=FALSE))

G.MATRIX <- matrix(c(128040/2,105356/2,4739445/2,4872356/2),nrow=2,ncol=2)
G.TEST <- likelihood.test(G.MATRIX,conservative=TRUE)
G.TEST$statistic
pchisq(G.TEST$statistic,df=1,lower.tail=FALSE)
-log10(pchisq(G.TEST$statistic,df=1,lower.tail=FALSE))


###PLOT COVERAGE FOR EACH CHROMOSOME USING 1kb OVERLAPPING SLIDING WINDOWS###
for (n in N) {
  COVERAGE.LOW  <- read.table(COVERAGE.FILES[grep(as.character(SAMPLES$LOW[n]),COVERAGE.FILES)], header=FALSE)
  colnames(COVERAGE.LOW) <- c("CHR","POS","COV")
  COVERAGE.HIGH <- read.table(COVERAGE.FILES[grep(as.character(SAMPLES$HIGH[n]),COVERAGE.FILES)],header=FALSE)
  colnames(COVERAGE.HIGH) <- c("CHR","POS","COV")
  
  if (nrow(COVERAGE.LOW) == nrow(COVERAGE.HIGH))
  {
    
    CHRO <- unique(COVERAGE.HIGH$CHR)[1:16]
    
    pdf(paste(SAMPLES[n,1],".CHR_COVERAGE",".pdf",sep=""))
    
    for (i in 1:length(CHRO))
    {
      CUR.LOW <- subset(COVERAGE.LOW, CHR == as.character(CHRO[i])) 
      CUR.HIGH <- subset(COVERAGE.HIGH, CHR == as.character(CHRO[i]))
      POSITIONS <- rollapply(CUR.LOW$POS,width=1,FUN=median,by=5000,partial=TRUE)
      L <- rollapply(CUR.LOW$COV,width=10000,FUN=median,by=5000,partial=TRUE)
      H <- rollapply(CUR.HIGH$COV,width=10000,FUN=median,by=5000,partial=TRUE)
      
      plot(round(POSITIONS/1000),L,type="l",col="#FF0000FF",ylab="COVERAGE",xlab="POSITION",ylim=c(0,3*median(c(CUR.LOW$COV,CUR.HIGH$COV))),main=CHRO[i])
      points(round(POSITIONS/1000),H,type="l",col="#0000FFFF",ylab="COVERAGE",xlab="POSITION",ylim=c(0,3*median(c(CUR.LOW$COV,CUR.HIGH$COV))))
      abline(h=median(CUR.LOW$COV),col="#FF0000FF",lty=2)
      abline(h=median(CUR.HIGH$COV),col="#0000FFFF",lty=2)
    }
    
    dev.off()
  }
}

##################################################################################################
#B) Filtering out false positive variant calls in vcf files to identify mutations in each mutant.#
##################################################################################################

###Determine thresholds for SNP Filtering###
RANDOM <- readVcf(VCF.FILES[9],"S288c")

#List of mutations confirmed by sequencing and used to set up filtering parameters. Can be found in SupplementaryFile11.zip.
CONFIRMED <-  readVcf("ConfirmedSNP.vcf","S288c")

#List of SNP called in sample 1
RANDOM.1 <- RANDOM@fixed
RANDOM.2 <- RANDOM@info
RANDOM.SNP <- cbind(RANDOM.1,RANDOM.2)

RANDOM.FREQ <- RANDOM@assays$data

DEPTH <- as.data.frame(RANDOM.FREQ$DP)
colnames(DEPTH) <- c("DP.L","DP.H")

FREQ.R <- as.data.frame(RANDOM.FREQ$RO)
colnames(FREQ.R) <- c("RO.L","RO.H")

QUAL.R <- as.data.frame(RANDOM.FREQ$QR)
colnames(QUAL.R) <- c("QR.L","QR.H")

FREQ.A <- RANDOM.FREQ$AO
QUAL.A <- RANDOM.FREQ$QA

for (i in 1:nrow(FREQ.A))
{
	FREQ.A[[i,1]] <- sum(FREQ.A[[i,1]])
	FREQ.A[[i,2]] <- sum(FREQ.A[[i,2]])
	QUAL.A[[i,1]] <- sum(QUAL.A[[i,1]])
	QUAL.A[[i,2]] <- sum(QUAL.A[[i,2]])
}

FREQ.A <- as.data.frame(cbind(unlist(FREQ.A[,1]),unlist(FREQ.A[,2])))
colnames(FREQ.A) <- c("AO.L","AO.H")

QUAL.A <- as.data.frame(cbind(unlist(QUAL.A[,1]),unlist(QUAL.A[,2])))
colnames(QUAL.A) <- c("QA.L","QA.H")

CHROMOSOME <- unlist(strsplit(RANDOM@rowRanges@ranges@NAMES,":"))[seq(1,2*nrow(RANDOM.SNP),2)]
POSITION <- RANDOM@rowRanges@ranges@start

RANDOM.SNP <- cbind(CHROMOSOME,POSITION,RANDOM.SNP,DEPTH,FREQ.R,FREQ.A,QUAL.R,QUAL.A)

RANDOM.SNP[,"FREQ.REF.L"] <- RANDOM.SNP[,"RO.L"]/RANDOM.SNP[,"DP.L"]
RANDOM.SNP[,"FREQ.REF.H"] <- RANDOM.SNP[,"RO.H"]/RANDOM.SNP[,"DP.H"]
RANDOM.SNP[,"FREQ.REF"] <- (RANDOM.SNP[,"RO.L"] + RANDOM.SNP[,"RO.H"])/(RANDOM.SNP[,"DP.L"] + RANDOM.SNP[,"DP.H"])


#List of confirmed causative mutations
CONFIRMED.1 <- CONFIRMED@fixed
CONFIRMED.2 <- CONFIRMED@info
CONFIRMED.SNP <- cbind(CONFIRMED.1,CONFIRMED.2)

CONFIRMED.FREQ <- CONFIRMED@assays$data

DEPTH <- as.data.frame(CONFIRMED.FREQ$DP)
colnames(DEPTH) <- c("DP.L","DP.H")

FREQ.R <- as.data.frame(CONFIRMED.FREQ$RO)
colnames(FREQ.R) <- c("RO.L","RO.H")

QUAL.R <- as.data.frame(CONFIRMED.FREQ$QR)
colnames(QUAL.R) <- c("QR.L","QR.H")

FREQ.A <- CONFIRMED.FREQ$AO
QUAL.A <- CONFIRMED.FREQ$QA

for (i in 1:nrow(FREQ.A))
{
	FREQ.A[[i,1]] <- sum(FREQ.A[[i,1]])
	FREQ.A[[i,2]] <- sum(FREQ.A[[i,2]])
	QUAL.A[[i,1]] <- sum(QUAL.A[[i,1]])
	QUAL.A[[i,2]] <- sum(QUAL.A[[i,2]])
}

FREQ.A <- as.data.frame(cbind(unlist(FREQ.A[,1]),unlist(FREQ.A[,2])))
colnames(FREQ.A) <- c("AO.L","AO.H")

QUAL.A <- as.data.frame(cbind(unlist(QUAL.A[,1]),unlist(QUAL.A[,2])))
colnames(QUAL.A) <- c("QA.L","QA.H")

CHROMOSOME <- unlist(strsplit(CONFIRMED@rowRanges@ranges@NAMES,":"))[seq(1,2*nrow(CONFIRMED.SNP),2)]
POSITION <- CONFIRMED@rowRanges@ranges@start

CONFIRMED.SNP <- cbind(CHROMOSOME,POSITION,CONFIRMED.SNP,DEPTH,FREQ.R,FREQ.A,QUAL.R,QUAL.A)


ALL.SNP <- rbind(RANDOM.SNP,CONFIRMED.SNP)
ALL.SNP[,"Class"] <- c(rep("Random",nrow(RANDOM.SNP)),rep("Confirmed",nrow(CONFIRMED.SNP)))
ALL.SNP[,"Class"] <- as.factor(ALL.SNP[,"Class"])


#QUAL threshold
PLOT <- ALL.SNP$QUAL
boxplot(PLOT ~ ALL.SNP$Class, notch=T)
min(CONFIRMED.SNP$QUAL)

#QA threshold
PLOT <- sum(ALL.SNP$QA)
boxplot(PLOT ~ ALL.SNP$Class, notch=T)
min(sum(CONFIRMED.SNP$QA))

#MQM threshold
PLOT <- sum(ALL.SNP$MQM)
boxplot(PLOT ~ ALL.SNP$Class, notch=T)
min(sum(CONFIRMED.SNP$MQM))

#SAF threshold
PLOT <- sum(ALL.SNP$SAF)
boxplot(PLOT ~ ALL.SNP$Class, notch=T)
min(sum(CONFIRMED.SNP$SAF))
length(which(PLOT > 5))

#SAR threshold
PLOT2 <- sum(ALL.SNP$SAR)
boxplot(PLOT2 ~ ALL.SNP$Class, notch=T)
boxplot(PLOT/PLOT2 ~ ALL.SNP$Class, notch=T)

#RPL threshold
PLOT <- sum(ALL.SNP$RPL)
boxplot(PLOT ~ ALL.SNP$Class, notch=T)
min(sum(CONFIRMED.SNP$RPL))
length(which(PLOT > 3))

#RPR threshold
PLOT <- sum(ALL.SNP$RPR)
boxplot(PLOT ~ ALL.SNP$Class, notch=T)
min(sum(CONFIRMED.SNP$RPR))
length(which(PLOT > 5))

#PAIRED threshold
PLOT <- ALL.SNP$PAIRED

for (i in 1:length(PLOT))
{
	PLOT[[i]] <- PLOT[[i]][1]
}

PLOT <- unlist(PLOT)

boxplot(PLOT ~ ALL.SNP$Class, notch=T)
min(unlist(CONFIRMED.SNP$PAIRED))
length(which(PLOT > 0.99))

wilcox.test(unlist(RANDOM.SNP[,"QUAL"]),unlist(CONFIRMED.SNP[,"QUAL"]))
wilcox.test(unlist(RANDOM.SNP[,"SAP"]),unlist(CONFIRMED.SNP[,"SAP"]))
wilcox.test(unlist(RANDOM.SNP[,"SAR"])/(unlist(RANDOM.SNP[,"SAR"])+unlist(RANDOM.SNP[,"SAF"])),unlist(CONFIRMED.SNP[,"SAR"])/(unlist(RANDOM.SNP[,"SAR"])+unlist(RANDOM.SNP[,"SAF"])))
wilcox.test(unlist(RANDOM.SNP[,"DP.H"]),unlist(CONFIRMED.SNP[,"DP.H"]))
wilcox.test(unlist(RANDOM.SNP[,"PAIRED"]),unlist(CONFIRMED.SNP[,"PAIRED"]))

#Filter out variants with more than one alternative allele
ALLELES <- sum(RANDOM.SNP$LEN)
SNP.FILTER <- RANDOM.SNP[which(ALLELES == 1),]
SNP.FILTER[,"ALT"] <- paste(unlist(SNP.FILTER[,"ALT"]),sep="")

#Filter out bad quality variants
SNP.FILTER <- subset(SNP.FILTER, QUAL > 200)
SNP.FILTER <- subset(SNP.FILTER, DP > 20)
SNP.FILTER <- subset(SNP.FILTER, AO > 3)
SNP.FILTER <- subset(SNP.FILTER, RO > 3)
SNP.FILTER <- subset(SNP.FILTER, MQM > 27)
SNP.FILTER <- subset(SNP.FILTER, PAIRED > 0.8)
SNP.FILTER <- subset(SNP.FILTER, PAIREDR > 0.8)
SNP.FILTER <- subset(SNP.FILTER, EPP < 50)
SNP.FILTER <- subset(SNP.FILTER, RPP < 50)
SNP.FILTER <- subset(SNP.FILTER, SAP < 100)
SNP.FILTER <- subset(SNP.FILTER, FREQ.REF > 0.1)
SNP.FILTER <- subset(SNP.FILTER, SAP < 30 | EPP < 15 | RPP < 15)


############################################################################################################
#C) Likelihood-ratio tests (G-tests) to compare allele frequencies between low and high fluorescence bulks.#
############################################################################################################

#Compute G.values for each variant. The results can be found in SupplementaryFile2.txt.
for (i in 1:nrow(SNP.FILTER))
{
	G.MATRIX <- matrix(c(SNP.FILTER$RO.L[i],SNP.FILTER$RO.H[i],SNP.FILTER$AO.L[i],SNP.FILTER$AO.H[i]),nrow=2,ncol=2)
	G.TEST <- likelihood.test(G.MATRIX,conservative=TRUE)
	if (SNP.FILTER[i,"FREQ.REF.L"] >= SNP.FILTER[i,"FREQ.REF.H"])
	{
		SIGN <- 1
	} else {
		SIGN <- -1
	}
	SNP.FILTER[i,"G.VALUE"] <- G.TEST$statistic
	P.VAL <- pchisq(G.TEST$statistic,df=1,lower.tail=FALSE)
	SNP.FILTER[i,"P.VALUE"] <- P.VAL
	SNP.FILTER[i,"P.SCORE"] <- -log10(P.VAL) * SIGN
}

write.table(SNP.FILTER,"SNP.FILTER.txt",row.names=FALSE,sep="\t",quote=FALSE)


#############################################################################################
#D) Plotting the number of mutations identified per strain (Figure 2 - figure supplement 1).#
#############################################################################################

###Clear memory###
rm(list=ls())
options(warn=-1)

###Load packages###
library(VariantAnnotation)
library(Deducer)
library(gtools)
library(zoo)
library(Hmisc)
library(ggplot2)
library(plotrix)
library(moments)
library(vcd)

###Set directory###
setwd("/Path.to.input.files")

#Load table including all mutations identified by BSA-Seq and Sanger sequencing that can be found in SupplementaryFile11.zip.
ALL.MUT <- read.table("SourceData1.txt",header=TRUE)

RANDOM.MUT <- subset(ALL.MUT, TYPE == "RANDOM" & SEQ.RUN != "SANGER" & SEQ.RUN != 605)
length(unique(RANDOM.MUT$STRAIN))

LOW.MUT <- subset(ALL.MUT, TYPE == "LOW.TAIL" & SEQ.RUN != "SANGER" & SEQ.RUN != 605)
length(unique(LOW.MUT $STRAIN))

HIGH.MUT <- subset(ALL.MUT, TYPE == "HIGH.TAIL" & SEQ.RUN != "SANGER" & SEQ.RUN != 605)
length(unique(HIGH.MUT $STRAIN))

OLD.MUT <- subset(ALL.MUT, SEQ.RUN == 605)
length(unique(OLD.MUT $STRAIN))

MAP.MUT <- subset(ALL.MUT, SEQ.RUN != "SANGER")

#Calculate for each strain the number of mutations and the average sequencing depth.
EFFECT.MUT <- aggregate(cbind(MUTANT.DP.L, MUTANT.DP.H, MEDIAN.EXPR.MEAN.EMS.1, MEDIAN.EXPR.SD.EMS.1, MEDIAN.EXPR.N.EMS.1,N.CAUSAL) ~ STRAIN + MUT.RUN, data=MAP.MUT, FUN=mean)
EFFECT.MUT.N <- aggregate(cbind(MUTANT.DP.L, MUTANT.DP.H, MEDIAN.EXPR.MEAN.EMS.1, MEDIAN.EXPR.SD.EMS.1, MEDIAN.EXPR.N.EMS.1,N.CAUSAL) ~ STRAIN + MUT.RUN, data=MAP.MUT, FUN=length)
mean(c(EFFECT.MUT$MUTANT.DP.L,EFFECT.MUT$MUTANT.DP.H))
min(c(EFFECT.MUT$MUTANT.DP.L,EFFECT.MUT$MUTANT.DP.H))
max(c(EFFECT.MUT$MUTANT.DP.L,EFFECT.MUT$MUTANT.DP.H))


SEQ.MUT <- subset(ALL.MUT, SEQ.RUN != "SANGER" & SEQ.RUN != 605)
SEQ.MUT.N <- aggregate(cbind(MUTANT.DP.L, MUTANT.DP.H, MEDIAN.EXPR.MEAN.EMS.1, MEDIAN.EXPR.SD.EMS.1, MEDIAN.EXPR.N.EMS.1,N.CAUSAL) ~ STRAIN + MUT.RUN, data=SEQ.MUT, FUN=length)$N.CAUSAL
length(unique(SEQ.MUT$STRAIN))
SS.MUT <- subset(SEQ.MUT, TYPE.1 %nin% c("INDEL","DUPLICATION"))

SIGNIFICANT.MUT <- subset(SEQ.MUT, P.VALUE < 1e-3)

GA <- subset(SS.MUT, REF == "G" & ALT == "A")
CT <- subset(SS.MUT, REF == "C" & ALT == "T")

#Testing if number of mutations per strain are Poisson distributed using goodness of fit.
GF <- goodfit(SEQ.MUT.N, type = "poisson", method = "ML")
summary(GF)

#Permutation tests to determine if the dispersion of number of mutations among strains is different from expected under a Poisson distribution.
N.REP <- 1e5

VAR.OBS <- var(SEQ.MUT.N)
KUR.OBS <- kurtosis(SEQ.MUT.N)
MAD.OBS <- mad(SEQ.MUT.N)

VAR.RAND <- rep(0,N.REP)
KUR.RAND <- rep(0,N.REP)
MAD.RAND <- rep(0,N.REP)

for (i in 1:N.REP)
{
  CUR.SAMPLE <- rpois(length(SEQ.MUT.N),mean(SEQ.MUT.N))
  
  VAR.RAND[i] <- var(CUR.SAMPLE)
  KUR.RAND[i] <- kurtosis(CUR.SAMPLE)
  MAD.RAND[i] <- mad(CUR.SAMPLE)
}

LOW.VAR <- length(which(VAR.RAND < VAR.OBS))/length(VAR.RAND)
HIGH.VAR <- length(which(VAR.RAND > VAR.OBS))/length(VAR.RAND)
P.VAR <- 2*min(LOW.VAR,HIGH.VAR)

LOW.KUR <- length(which(KUR.RAND < KUR.OBS))/length(KUR.RAND)
HIGH.KUR <- length(which(KUR.RAND > KUR.OBS))/length(KUR.RAND)
P.KUR <- 2*min(LOW.KUR,HIGH.KUR)

LOW.MAD <- length(which(MAD.RAND < MAD.OBS))/length(MAD.RAND)
HIGH.MAD <- length(which(MAD.RAND > MAD.OBS))/length(MAD.RAND)
P.MAD <- 2*min(LOW.MAD,HIGH.MAD)

#Plotting the number of mutations per strain.
BREAKS <- seq(0,52,by=10/3)

POIS <- c()
POS <- c()

for (i in 2:length(BREAKS))
{
  POIS[i-1] <- abs(ppois(BREAKS[i],lambda=mean(SEQ.MUT.N))-ppois(BREAKS[i-1],lambda=mean(SEQ.MUT.N)))*length(SEQ.MUT.N)
  POS[i-1] <- mean(c(BREAKS[i],BREAKS[i-1]))
}

pdf("Figure 2 - figure supplement 1.pdf",useDingbats=F,height=6,width=7)
#windows(height=6,width=7)
hist(SEQ.MUT.N,breaks=BREAKS,ylim=c(0,20),xlab="# mutations per strain",ylab="# mutant strains")
points(POS,POIS,col="blue",pch=16,type="b")
abline(v=mean(SEQ.MUT.N),lty=2)
dev.off()


##################################################################################################################
#E) Plotting expression changes in EMS mutants depending on the number of mutations associated with fluorescence #
#   in BSA-Seq data (Figure 2 - figure supplement 2).                                                             #
##################################################################################################################

EFFECT.SIZE <- aggregate(cbind(MEDIAN.EXPR.MEAN.EMS.1, MEDIAN.EXPR.SD.EMS.1, MEDIAN.EXPR.N.EMS.1,N.CAUSAL) ~ STRAIN + MUT.RUN, data=ALL.MUT ,FUN=mean)
N.MUTATION <- aggregate(cbind(MEDIAN.EXPR.MEAN.EMS.1, MEDIAN.EXPR.SD.EMS.1, MEDIAN.EXPR.N.EMS.1) ~ STRAIN + MUT.RUN, data=ALL.MUT ,FUN=length)

N.MUTATION <- N.MUTATION[,3]

PLOT <- cbind(EFFECT.SIZE,N.MUTATION)

#Use different colors for mutants with increased fluorescence and decreased fluorescence relative to the progenitor strain.
for (i in 1:nrow(PLOT))
{
  if (PLOT[i,"MEDIAN.EXPR.MEAN.EMS.1"] > 1)
  {
    PLOT[i,"COLOR"] <- "red"
  } else {
    PLOT[i,"COLOR"] <- "blue"
  }
}

#Calculate absolute expression changes.
PLOT[,"EFFECT.SIZE"] <- PLOT[,"MEDIAN.EXPR.MEAN.EMS.1"]-1
PLOT[,"ABSOLUTE.EFFECT"] <- abs(1-PLOT[,"MEDIAN.EXPR.MEAN.EMS.1"])
PLOT[,"N.CAUSAL"] <- as.factor(PLOT[,"N.CAUSAL"])

PLOT <- subset(PLOT, N.MUTATION > 3)

PLOT[26,"N.CAUSAL"] <- 1

PLOT <- droplevels(PLOT)

#The table SourceData6.txt can be found in SupplementaryFile11.zip.
write.table(PLOT,"SourceData6.txt",sep="\t",row.names=FALSE)

for (i in 1:nrow(PLOT))
{
  if (PLOT[i,"EFFECT.SIZE"] > 0)
  {
    PLOT[i,"DIRECTION"] <- "INCREASE"
  }
  if (PLOT[i,"EFFECT.SIZE"] < 0)
  {
    PLOT[i,"DIRECTION"] <- "DECREASE"
  }
}

PLOT[,"DIRECTION"] <- as.factor(PLOT[,"DIRECTION"])

for (i in 1:nrow(PLOT))
{
  PLOT[i,"POSITION"] <- runif(1,as.integer(PLOT[i,"N.CAUSAL"])-0.15,as.integer(PLOT[i,"N.CAUSAL"])+0.15)
}

pdf("Figure 2 - figure supplement 2.pdf",useDingbats=FALSE,height=7,width=7)
boxplot(PLOT[,"ABSOLUTE.EFFECT"]~PLOT[,"N.CAUSAL"],xlab = "# associated mutations",ylab="Absolute fluorescence change in EMS mutant",notch=T,varwidth=T)
points(PLOT[,"POSITION"],PLOT[,"ABSOLUTE.EFFECT"],pch=16,col=PLOT[,"COLOR"],cex=1.3)
dev.off()


###########################################################################################
#F) Relationship between the total number of mutations and the absolute expression change #
#   among EMS mutants (Figure 2 - figure supplement 3).                                   #                                                        
###########################################################################################

MAP.INCREASE <- subset(PLOT, EFFECT.SIZE > 0 & N.CAUSAL != 0)
MAP.DECREASE <- subset(PLOT, EFFECT.SIZE < 0 & N.CAUSAL != 0)
UNMAP.INCREASE <- subset(PLOT, EFFECT.SIZE > 0 & N.CAUSAL == 0)
UNMAP.DECREASE <- subset(PLOT, EFFECT.SIZE < 0 & N.CAUSAL == 0)

MAP <- subset(PLOT, N.CAUSAL != 0)
UNMAP <- subset(PLOT, N.CAUSAL == 0)
INCREASE <- subset(PLOT, EFFECT.SIZE > 0)
DECREASE <- subset(PLOT, EFFECT.SIZE < 0)

MODEL.MAP <- lm(MAP[,"ABSOLUTE.EFFECT"] ~ MAP[,"N.MUTATION"])
MODEL.UNMAP <- lm(UNMAP[,"ABSOLUTE.EFFECT"] ~ UNMAP[,"N.MUTATION"])

pdf("Figure 2 - figure supplement 3.pdf",useDingbats=FALSE,height=7,width=7)
#windows(height=7,width=7)
plot(MAP.INCREASE[,"N.MUTATION"], MAP.INCREASE[,"ABSOLUTE.EFFECT"],pch=16,col="#00000066",xlab="Number of mutations",ylab="Expression change",xlim=c(0,60),ylim=c(0,0.6),cex=2)
points(MAP.DECREASE[,"N.MUTATION"], MAP.DECREASE[,"ABSOLUTE.EFFECT"],pch=16,col="#00000066",cex=2)
points(UNMAP.INCREASE[,"N.MUTATION"], UNMAP.INCREASE[,"ABSOLUTE.EFFECT"],pch=16,col="#00FF66BB",cex=2)
points(UNMAP.DECREASE[,"N.MUTATION"], UNMAP.DECREASE[,"ABSOLUTE.EFFECT"],pch=16,col="#00FF66BB",cex=2)

abline(lm(MAP[,"ABSOLUTE.EFFECT"] ~ MAP[,"N.MUTATION"]),col="#00000066",lty=1,lwd=1.5)
abline(lm(UNMAP[,"ABSOLUTE.EFFECT"] ~ UNMAP[,"N.MUTATION"]),col="#00FF66BB",lty=1,lwd=1.5)
dev.off()

summary(lm(PLOT[,"ABSOLUTE.EFFECT"] ~ PLOT[,"N.MUTATION"]))
summary(MODEL.MAP)
summary(MODEL.UNMAP)

cor.test(PLOT[,"N.MUTATION"],PLOT[,"ABSOLUTE.EFFECT"])
cor.test(MAP[,"N.MUTATION"],MAP[,"ABSOLUTE.EFFECT"])
cor.test(UNMAP[,"N.MUTATION"],UNMAP[,"ABSOLUTE.EFFECT"])


################################################################################################################################
#G) Comparing the strengths of association with fluorescence (G-values) of genetically linked mutations (Supplementary File 3).#
################################################################################################################################

###Clear memory###
rm(list=ls())
options(warn=-1)

###Load packages###
library(Deducer)
library(gtools)
library(zoo)
library(Hmisc)
library(ggplot2)
library(plotrix)
library(permute)

###Set directory###
setwd("/Path.to.input.files")

###Function to compute probability that lowest G-value is a causal mutation and highest G-value is not###

#D is the genetic distance in cM between mutations MUT1 and MUT2 (value given as example).
D <- 6.37692

#Allele counts for linked mutations MUT1 and MUT2 in low and high fluorescence bulks (values given as example).
N.REF.LOW.MUT1 <- 118
N.REF.LOW.MUT2 <- 99
N.REF.HIGH.MUT1 <- 63
N.REF.HIGH.MUT2 <- 50
N.ALT.LOW.MUT1 <- 13
N.ALT.LOW.MUT2 <- 18
N.ALT.HIGH.MUT1 <- 54
N.ALT.HIGH.MUT2 <- 40

#MUT2 is the mutation with lowest G-value and MUT1 the mutation with highest G-value.
P.CAUSAL <- function(D=D, N.REF.LOW.MUT1=N.REF.LOW.MUT1, N.REF.LOW.MUT2=N.REF.LOW.MUT2, N.REF.HIGH.MUT1=N.REF.HIGH.MUT1, N.REF.HIGH.MUT2=N.REF.HIGH.MUT2, N.ALT.LOW.MUT1=N.ALT.LOW.MUT1, N.ALT.LOW.MUT2=N.ALT.LOW.MUT2, N.ALT.HIGH.MUT1=N.ALT.HIGH.MUT1, N.ALT.HIGH.MUT2=N.ALT.HIGH.MUT2)
{
  N.REP <- 10^4
  G.SIM.MUT1 <- rep(0,N.REP) 
  
  for (i in 1:N.REP)
  {
    ###Pick alleles for site 1 in low bulk###
    
    #Take into account sampling error in the estimate of allele frequency for MUT2
    FREQ.REF.MUT2 <- rbinom(1,N.REF.LOW.MUT2+N.ALT.LOW.MUT2,N.REF.LOW.MUT2/(N.REF.LOW.MUT2+N.ALT.LOW.MUT2))/(N.REF.LOW.MUT2+N.ALT.LOW.MUT2)
    
    #Take into account sampling error in the estimate of allele frequency for MUT1
    N.MUT1 <- rbinom(1,N.REF.LOW.MUT1+N.ALT.LOW.MUT1,1-FREQ.REF.MUT2)
    STATES.MUT1 <- c(rep(-1,N.MUT1),rep(1,N.REF.LOW.MUT1+N.ALT.LOW.MUT1-N.MUT1))
    
    #Generate vector of recombination events
    P.REC <- (1-exp(-2*D/100))/2
    REC <- sample(c(1,-1),N.REF.LOW.MUT1+N.ALT.LOW.MUT1,replace=TRUE,prob=c(1-P.REC,P.REC))
    
    #Apply recombination to alleles at MUT1.
    STATES.MUT1 <- STATES.MUT1*REC
    
    N.SIM.REF.LOW.MUT1 <- length(which(STATES.MUT1 == 1))
    N.SIM.ALT.LOW.MUT1 <- length(which(STATES.MUT1 == -1))
    
    
    ###Pick alleles for site 2 in high bulk###
    
    #Take into account sampling error in the estimate of allele frequency for MUT2
    FREQ.REF.MUT2 <- rbinom(1,N.REF.HIGH.MUT2+N.ALT.HIGH.MUT2,N.REF.HIGH.MUT2/(N.REF.HIGH.MUT2+N.ALT.HIGH.MUT2))/(N.REF.HIGH.MUT2+N.ALT.HIGH.MUT2)
    
    #Take into account sampling error in the estimate of allele frequency for MUT1
    N.MUT1 <- rbinom(1,N.REF.HIGH.MUT1+N.ALT.HIGH.MUT1,1-FREQ.REF.MUT2)
    STATES.MUT1 <- c(rep(-1,N.MUT1),rep(1,N.REF.HIGH.MUT1+N.ALT.HIGH.MUT1-N.MUT1))
    
    #Generate vector of recombination events
    P.REC <- (1-exp(-2*D/100))/2
    REC <- sample(c(1,-1),N.REF.HIGH.MUT1+N.ALT.HIGH.MUT1,replace=TRUE,prob=c(1-P.REC,P.REC))
    
    #Apply recombination to alleles at MUT1.
    STATES.MUT1 <- STATES.MUT1*REC
    
    N.SIM.REF.HIGH.MUT1 <- length(which(STATES.MUT1 == 1))
    N.SIM.ALT.HIGH.MUT1 <- length(which(STATES.MUT1 == -1))
    
    
    ###Compute G.value for simulated allele frequencies at MUT1###
    
    G.MATRIX <- matrix(c(N.SIM.REF.LOW.MUT1,N.SIM.ALT.LOW.MUT1,N.SIM.REF.HIGH.MUT1,N.SIM.ALT.HIGH.MUT1),ncol=2)
    G.SIM.MUT1[i] <- likelihood.test(G.MATRIX,conservative=TRUE)$statistic
  }
  
  #Observed G.value at MUT1
  G.MATRIX <- matrix(c(N.REF.LOW.MUT1,N.ALT.LOW.MUT1,N.REF.HIGH.MUT1,N.ALT.HIGH.MUT1),ncol=2)
  G.MUT1 <- likelihood.test(G.MATRIX,conservative=TRUE)$statistic
  
  P.VAL <- length(which(G.SIM.MUT1 >= G.MUT1))/(2*length(G.SIM.MUT1))
  return(P.VAL)
}


###Function to compare observed lowest G-value to expected G-value based on linkage to highest G-value###

P.LINKED <- function(D=D, N.REF.LOW.MUT1=N.REF.LOW.MUT1, N.REF.LOW.MUT2=N.REF.LOW.MUT2, N.REF.HIGH.MUT1=N.REF.HIGH.MUT1, N.REF.HIGH.MUT2=N.REF.HIGH.MUT2, N.ALT.LOW.MUT1=N.ALT.LOW.MUT1, N.ALT.LOW.MUT2=N.ALT.LOW.MUT2, N.ALT.HIGH.MUT1=N.ALT.HIGH.MUT1, N.ALT.HIGH.MUT2=N.ALT.HIGH.MUT2)
{
  N.REP <- 10^4
  G.SIM.MUT2 <- rep(0,N.REP) 
  
  for (i in 1:N.REP)
  {
    
    ###Pick alleles for site 2 in low bulk###
    
    #Take into account sampling error in the estimate of allele frequency for MUT1
    FREQ.REF.MUT1 <- rbinom(1,N.REF.LOW.MUT1+N.ALT.LOW.MUT1,N.REF.LOW.MUT1/(N.REF.LOW.MUT1+N.ALT.LOW.MUT1))/(N.REF.LOW.MUT1+N.ALT.LOW.MUT1)
    
    #Take into account sampling error in the estimate of allele frequency for MUT2
    N.MUT2 <- rbinom(1,N.REF.LOW.MUT2+N.ALT.LOW.MUT2,1-FREQ.REF.MUT1)
    STATES.MUT2 <- c(rep(-1,N.MUT2),rep(1,N.REF.LOW.MUT2+N.ALT.LOW.MUT2-N.MUT2))
    
    #Generate vector of recombination events
    P.REC <- (1-exp(-2*D/100))/2
    REC <- sample(c(1,-1),N.REF.LOW.MUT2+N.ALT.LOW.MUT2,replace=TRUE,prob=c(1-P.REC,P.REC))
    
    #Apply recombination to alleles at MUT1.
    STATES.MUT2 <- STATES.MUT2*REC
    
    N.SIM.REF.LOW.MUT2 <- length(which(STATES.MUT2 == 1))
    N.SIM.ALT.LOW.MUT2 <- length(which(STATES.MUT2 == -1))
    
    
    ###Pick alleles for site 2 in high bulk###
    
    #Take into account sampling error in the estimate of allele frequency for MUT1
    FREQ.REF.MUT1 <- rbinom(1,N.REF.HIGH.MUT1+N.ALT.HIGH.MUT1,N.REF.HIGH.MUT1/(N.REF.HIGH.MUT1+N.ALT.HIGH.MUT1))/(N.REF.HIGH.MUT1+N.ALT.HIGH.MUT1)
    
    #Take into account sampling error in the estimate of allele frequency for MUT2
    N.MUT2 <- rbinom(1,N.REF.HIGH.MUT2+N.ALT.HIGH.MUT2,1-FREQ.REF.MUT1)
    STATES.MUT2 <- c(rep(-1,N.MUT2),rep(1,N.REF.HIGH.MUT2+N.ALT.HIGH.MUT2-N.MUT2))
    
    #Generate vector of recombination events
    P.REC <- (1-exp(-2*D/100))/2
    REC <- sample(c(1,-1),N.REF.HIGH.MUT2+N.ALT.HIGH.MUT2,replace=TRUE,prob=c(1-P.REC,P.REC))
    
    #Apply recombination to alleles at MUT1.
    STATES.MUT2 <- STATES.MUT2*REC
    
    N.SIM.REF.HIGH.MUT2 <- length(which(STATES.MUT2 == 1))
    N.SIM.ALT.HIGH.MUT2 <- length(which(STATES.MUT2 == -1))
    
    
    ###Compute G.value for simulated allele frequencies at MUT1###
    
    G.MATRIX <- matrix(c(N.SIM.REF.LOW.MUT2,N.SIM.ALT.LOW.MUT2,N.SIM.REF.HIGH.MUT2,N.SIM.ALT.HIGH.MUT2),ncol=2)
    G.SIM.MUT2[i] <- likelihood.test(G.MATRIX,conservative=TRUE)$statistic
    
  }
  
  #Observed G.value at MUT1
  G.MATRIX <- matrix(c(N.REF.LOW.MUT2,N.ALT.LOW.MUT2,N.REF.HIGH.MUT2,N.ALT.HIGH.MUT2),ncol=2)
  G.MUT2 <- likelihood.test(G.MATRIX,conservative=TRUE)$statistic
  
  P.VAL.LOW <- length(which(G.SIM.MUT2 >= G.MUT2))/length(G.SIM.MUT2)
  P.VAL.HIGH <- length(which(G.SIM.MUT2 <= G.MUT2))/length(G.SIM.MUT2)
  
  P.VAL <- min(P.VAL.LOW,P.VAL.HIGH)*2
  return(P.VAL)
}


###Function to calculate probability to observe highest G-value only by linkage to second highest G-value###

P.NOT.CAUSAL <- function(D=D, N.REF.LOW.MUT1=N.REF.LOW.MUT1, N.REF.LOW.MUT2=N.REF.LOW.MUT2, N.REF.HIGH.MUT1=N.REF.HIGH.MUT1, N.REF.HIGH.MUT2=N.REF.HIGH.MUT2, N.ALT.LOW.MUT1=N.ALT.LOW.MUT1, N.ALT.LOW.MUT2=N.ALT.LOW.MUT2, N.ALT.HIGH.MUT1=N.ALT.HIGH.MUT1, N.ALT.HIGH.MUT2=N.ALT.HIGH.MUT2)
{
  N.REP <- 10^4
  G.SIM.MUT2 <- rep(0,N.REP) 
  
  for (i in 1:N.REP)
  {
    
    ###Pick alleles for site 2 in low bulk###
    
    #Take into account sampling error in the estimate of allele frequency for MUT1
    FREQ.REF.MUT1 <- rbinom(1,N.REF.LOW.MUT1+N.ALT.LOW.MUT1,N.REF.LOW.MUT1/(N.REF.LOW.MUT1+N.ALT.LOW.MUT1))/(N.REF.LOW.MUT1+N.ALT.LOW.MUT1)
    
    #Take into account sampling error in the estimate of allele frequency for MUT2
    N.MUT2 <- rbinom(1,N.REF.LOW.MUT2+N.ALT.LOW.MUT2,1-FREQ.REF.MUT1)
    STATES.MUT2 <- c(rep(-1,N.MUT2),rep(1,N.REF.LOW.MUT2+N.ALT.LOW.MUT2-N.MUT2))
    
    #Generate vector of recombination events
    P.REC <- (1-exp(-2*D/100))/2
    REC <- sample(c(1,-1),N.REF.LOW.MUT2+N.ALT.LOW.MUT2,replace=TRUE,prob=c(1-P.REC,P.REC))
    
    #Apply recombination to alleles at MUT1.
    STATES.MUT2 <- STATES.MUT2*REC
    
    N.SIM.REF.LOW.MUT2 <- length(which(STATES.MUT2 == 1))
    N.SIM.ALT.LOW.MUT2 <- length(which(STATES.MUT2 == -1))
    
    
    ###Pick alleles for site 2 in high bulk###
    
    #Take into account sampling error in the estimate of allele frequency for MUT1
    FREQ.REF.MUT1 <- rbinom(1,N.REF.HIGH.MUT1+N.ALT.HIGH.MUT1,N.REF.HIGH.MUT1/(N.REF.HIGH.MUT1+N.ALT.HIGH.MUT1))/(N.REF.HIGH.MUT1+N.ALT.HIGH.MUT1)
    
    #Take into account sampling error in the estimate of allele frequency for MUT2
    N.MUT2 <- rbinom(1,N.REF.HIGH.MUT2+N.ALT.HIGH.MUT2,1-FREQ.REF.MUT1)
    STATES.MUT2 <- c(rep(-1,N.MUT2),rep(1,N.REF.HIGH.MUT2+N.ALT.HIGH.MUT2-N.MUT2))
    
    #Generate vector of recombination events
    P.REC <- (1-exp(-2*D/100))/2
    REC <- sample(c(1,-1),N.REF.HIGH.MUT2+N.ALT.HIGH.MUT2,replace=TRUE,prob=c(1-P.REC,P.REC))
    
    #Apply recombination to alleles at MUT1.
    STATES.MUT2 <- STATES.MUT2*REC
    
    N.SIM.REF.HIGH.MUT2 <- length(which(STATES.MUT2 == 1))
    N.SIM.ALT.HIGH.MUT2 <- length(which(STATES.MUT2 == -1))
    
    
    ###Compute G.value for simulated allele frequencies at MUT1###
    
    G.MATRIX <- matrix(c(N.SIM.REF.LOW.MUT2,N.SIM.ALT.LOW.MUT2,N.SIM.REF.HIGH.MUT2,N.SIM.ALT.HIGH.MUT2),ncol=2)
    G.SIM.MUT2[i] <- likelihood.test(G.MATRIX,conservative=TRUE)$statistic
    
  }
  
  #Observed G.value at MUT1
  G.MATRIX <- matrix(c(N.REF.LOW.MUT2,N.ALT.LOW.MUT2,N.REF.HIGH.MUT2,N.ALT.HIGH.MUT2),ncol=2)
  G.MUT2 <- likelihood.test(G.MATRIX,conservative=TRUE)$statistic
  
  P.VAL <- length(which(G.SIM.MUT2 >= G.MUT2))/length(G.SIM.MUT2)
  return(P.VAL)
}


#Input file with all linked mutations can be found in SupplementaryFile11.zip
LINKED.MUT <- read.table("Linked.Mutations.txt",header=TRUE)

for (i in 1:nrow(LINKED.MUT))
{
  if (LINKED.MUT[i,"GENETIC.DISTANCE"] == 0)
  {
    CUR.MUT <- subset(LINKED.MUT, STRAIN == LINKED.MUT[i,"STRAIN"])
    
    CUR.MUT <- CUR.MUT[order(CUR.MUT[,"G.VALUE"],decreasing = TRUE),]
    CUR.MUT1 <- CUR.MUT[2,]
    CUR.MUT2 <- LINKED.MUT[i,]
    
    D <- CUR.MUT1[1,"GENETIC.DISTANCE"]
    N.REF.LOW.MUT1 <- CUR.MUT1[1,"RO.L"]
    N.REF.LOW.MUT2 <- CUR.MUT2[1,"RO.L"]
    N.REF.HIGH.MUT1 <- CUR.MUT1[1,"RO.H"]
    N.REF.HIGH.MUT2 <- CUR.MUT2[1,"RO.H"]
    N.ALT.LOW.MUT1 <- CUR.MUT1[1,"AO.L"]
    N.ALT.LOW.MUT2 <- CUR.MUT2[1,"AO.L"]
    N.ALT.HIGH.MUT1 <- CUR.MUT1[1,"AO.H"]
    N.ALT.HIGH.MUT2 <- CUR.MUT2[1,"AO.H"]
    
    LINKED.MUT[i,"P.LINKED"] <- P.NOT.CAUSAL(D,N.REF.LOW.MUT1,N.REF.LOW.MUT2,N.REF.HIGH.MUT1,N.REF.HIGH.MUT2,N.ALT.LOW.MUT1,N.ALT.LOW.MUT2,N.ALT.HIGH.MUT1,N.ALT.HIGH.MUT2)
    LINKED.MUT[i,"P.CAUSAL"] <- NA
    
  } else {
    CUR.MUT1 <- subset(LINKED.MUT, STRAIN == LINKED.MUT[i,"STRAIN"] & GENETIC.DISTANCE == 0)
    CUR.MUT2 <- LINKED.MUT[i,]
    
    D <- CUR.MUT2[1,"GENETIC.DISTANCE"]
    N.REF.LOW.MUT1 <- CUR.MUT1[1,"RO.L"]
    N.REF.LOW.MUT2 <- CUR.MUT2[1,"RO.L"]
    N.REF.HIGH.MUT1 <- CUR.MUT1[1,"RO.H"]
    N.REF.HIGH.MUT2 <- CUR.MUT2[1,"RO.H"]
    N.ALT.LOW.MUT1 <- CUR.MUT1[1,"AO.L"]
    N.ALT.LOW.MUT2 <- CUR.MUT2[1,"AO.L"]
    N.ALT.HIGH.MUT1 <- CUR.MUT1[1,"AO.H"]
    N.ALT.HIGH.MUT2 <- CUR.MUT2[1,"AO.H"]
    
    #LINKED.MUT[i,"P.CAUSAL"] <- P.CAUSAL(D,N.REF.LOW.MUT1,N.REF.LOW.MUT2,N.REF.HIGH.MUT1,N.REF.HIGH.MUT2,N.ALT.LOW.MUT1,N.ALT.LOW.MUT2,N.ALT.HIGH.MUT1,N.ALT.HIGH.MUT2)
    #LINKED.MUT[i,"P.LINKED"] <- P.LINKED(D,N.REF.LOW.MUT1,N.REF.LOW.MUT2,N.REF.HIGH.MUT1,N.REF.HIGH.MUT2,N.ALT.LOW.MUT1,N.ALT.LOW.MUT2,N.ALT.HIGH.MUT1,N.ALT.HIGH.MUT2)
    
  }
}

#Output file used to make SupplementaryFile3.xls
write.table(LINKED.MUT,"Linked.Mutations.txt",sep="\t",row.names=FALSE)


####################################################################################
#H) Types of trans-regulatory and non-regulatory mutations identified (Figure 3A). #
####################################################################################

#Clear memory
rm(list=ls())
options(warn=-1)

###Load packages###
#library(VariantAnnotation)
library(Deducer)
library(gtools)
library(zoo)
library(Hmisc)
library(ggplot2)
library(plotrix)

###Set directory###
setwd("/Path.to.input.files")

###Open table with all mutations###
ALL.MUT <- read.table("SourceData8.txt",header=TRUE)
ALL.MUT <- subset(ALL.MUT, CHROMOSOME != "chrYFP")

###Mutation categories###

#CAUSAL is a dataframe including the 69 trans-regulatory mutations described in the manuscript.
CAUSAL <- subset(ALL.MUT, MUTATION == "CAUSAL") 

#RANDOM is a dataframe including only the trans-regulatory mutations that were identified in EMS mutants not enriched for large effects.
RANDOM <- subset(ALL.MUT, MUTATION == "CAUSAL" & TYPE == "RANDOM")

#RANDOM is a dataframe including only the trans-regulatory mutations that were identified in EMS mutants enriched for large effects.
TAIL <- subset(ALL.MUT, MUTATION == "CAUSAL" & TYPE %in% c("HIGH.TAIL","LOW.TAIL"))

#NEUTRAL is a dataframe including all non-regulatory mutations identified in EMS mutants.
NEUTRAL <- subset(ALL.MUT, MUTATION == "NEUTRAL") 

#SUBSTITUTION is a dataframe including only non-regulatory mutations corresponding to single nucleotide substitutions (excluding indels).
SUBSTITUTION <- NEUTRAL[-which(NEUTRAL[,"TYPE.1"] == 'INDEL'),]

#INDEL is a dataframe including only non-regulatory mutations corresponding to indels.
INDEL <- subset(ALL.MUT, MUTATION == "NEUTRAL" & TYPE.1 == "INDEL")

#Genes involved in iron homeostasis in which a trans-regulatory mutation was identified.
IRON.GENES <- c("YNL240C","YER145C","YLL029W","YDR270W","YHR122W","YLR136C")

#Genes involved in de novo purine biosynthesis in which a trans-regulatory mutation was identified.
ADE.GENES <- c("YMR300C","YOR128C","YGL234W","YGR061C")

#CONSERVATIVE is a dataframe including only trans-regulatory mutations that do not affect genes involved in iron homesotasis or purine biosynthesis.
CONSERVATIVE <- subset(RANDOM, GENE.FOCAL %nin% c(IRON.GENES,ADE.GENES))

SUB.RANDOM <- droplevels(rbind(NEUTRAL,RANDOM))
SUB.CAUSAL <- droplevels(rbind(NEUTRAL,CAUSAL))
SUB.TAIL <- droplevels(rbind(NEUTRAL,TAIL))
SUB.CONS <- droplevels(rbind(NEUTRAL,CONSERVATIVE))

MUT.TYPE <- c("NEUTRAL","CAUSAL","RANDOM","TAIL","CONSERVATIVE")
N.MUT <- c(nrow(NEUTRAL),nrow(CAUSAL),nrow(RANDOM),nrow(TAIL),nrow(CONSERVATIVE))

#SANGER is a dataframe including only trans-regulatory mutations that were identified by sequencing candidate genes.
SANGER <- subset(CAUSAL, SEQ.RUN == "SANGER")

#BSA.SEQ is a dataframe including only trans-regulatory mutations that were identified by BSA-Seq (genetic mapping).
BSA.SEQ <- subset(CAUSAL, SEQ.RUN != "SANGER")

MUT.TYPE.2 <- c(MUT.TYPE,"BSA.SEQ","SANGER")
N.MUT.2 <- c(N.MUT,nrow(BSA.SEQ),nrow(SANGER))


###Statistical tests used to compare the frequency of different types of mutations observed between trans-regulatory and non-regulatory mutations.
STAT <- data.frame(MUT.TYPE, N.MUT)

STAT[1,"N.G>A.C>T"] <- length(which((NEUTRAL[,"REF"] == "G" & NEUTRAL[,"ALT"] == "A")|(NEUTRAL[,"REF"] == "C" & NEUTRAL[,"ALT"] == "T")))
STAT[2,"N.G>A.C>T"] <- length(which((CAUSAL[,"REF"] == "G" & CAUSAL[,"ALT"] == "A")|(CAUSAL[,"REF"] == "C" & CAUSAL[,"ALT"] == "T")))
STAT[3,"N.G>A.C>T"] <- length(which((RANDOM[,"REF"] == "G" & RANDOM[,"ALT"] == "A")|(RANDOM[,"REF"] == "C" & RANDOM[,"ALT"] == "T")))
STAT[4,"N.G>A.C>T"] <- length(which((TAIL[,"REF"] == "G" & TAIL[,"ALT"] == "A")|(TAIL[,"REF"] == "C" & TAIL[,"ALT"] == "T")))
STAT[5,"N.G>A.C>T"] <- length(which((CONSERVATIVE[,"REF"] == "G" & CONSERVATIVE[,"ALT"] == "A")|(CONSERVATIVE[,"REF"] == "C" & CONSERVATIVE[,"ALT"] == "T")))

STAT[1,"N.NOT.G>A.C>T"] <- STAT[1,"N.MUT"] - STAT[1,"N.G>A.C>T"]
STAT[2,"N.NOT.G>A.C>T"] <- STAT[2,"N.MUT"] - STAT[2,"N.G>A.C>T"]
STAT[3,"N.NOT.G>A.C>T"] <- STAT[3,"N.MUT"] - STAT[3,"N.G>A.C>T"]
STAT[4,"N.NOT.G>A.C>T"] <- STAT[4,"N.MUT"] - STAT[4,"N.G>A.C>T"]
STAT[5,"N.NOT.G>A.C>T"] <- STAT[5,"N.MUT"] - STAT[5,"N.G>A.C>T"]

G.MATRIX.NEUTRAL <- matrix(c(unlist(STAT[1,c("N.G>A.C>T","N.NOT.G>A.C>T")]),unlist(STAT[1,c("N.G>A.C>T","N.NOT.G>A.C>T")])),nrow=2,byrow=TRUE)
G.MATRIX.CAUSAL <- matrix(c(unlist(STAT[1,c("N.G>A.C>T","N.NOT.G>A.C>T")]),unlist(STAT[2,c("N.G>A.C>T","N.NOT.G>A.C>T")])),nrow=2,byrow=TRUE)
G.MATRIX.RANDOM <- matrix(c(unlist(STAT[1,c("N.G>A.C>T","N.NOT.G>A.C>T")]),unlist(STAT[3,c("N.G>A.C>T","N.NOT.G>A.C>T")])),nrow=2,byrow=TRUE)
G.MATRIX.TAIL <- matrix(c(unlist(STAT[1,c("N.G>A.C>T","N.NOT.G>A.C>T")]),unlist(STAT[4,c("N.G>A.C>T","N.NOT.G>A.C>T")])),nrow=2,byrow=TRUE)
G.MATRIX.CONSERVATIVE <- matrix(c(unlist(STAT[1,c("N.G>A.C>T","N.NOT.G>A.C>T")]),unlist(STAT[5,c("N.G>A.C>T","N.NOT.G>A.C>T")])),nrow=2,byrow=TRUE)

STAT[,"G.STAT.TRANSITIONS"] <- c(likelihood.test(G.MATRIX.NEUTRAL,conservative=FALSE)$statistic,likelihood.test(G.MATRIX.CAUSAL,conservative=FALSE)$statistic,likelihood.test(G.MATRIX.RANDOM,conservative=FALSE)$statistic,likelihood.test(G.MATRIX.TAIL,conservative=FALSE)$statistic,likelihood.test(G.MATRIX.CONSERVATIVE,conservative=FALSE)$statistic)

STAT[,"P.VAL.TRANSITIONS"] <- c(pchisq(STAT[1,"G.STAT.TRANSITIONS"],df=1,lower.tail=FALSE),pchisq(STAT[2,"G.STAT.TRANSITIONS"],df=1,lower.tail=FALSE),pchisq(STAT[3,"G.STAT.TRANSITIONS"],df=1,lower.tail=FALSE),pchisq(STAT[4,"G.STAT.TRANSITIONS"],df=1,lower.tail=FALSE),pchisq(STAT[5,"G.STAT.TRANSITIONS"],df=1,lower.tail=FALSE))


STAT[1,"N.G>A"] <- length(which((NEUTRAL[,"REF"] == "G" & NEUTRAL[,"ALT"] == "A")))
STAT[2,"N.G>A"] <- length(which((CAUSAL[,"REF"] == "G" & CAUSAL[,"ALT"] == "A")))
STAT[3,"N.G>A"] <- length(which((RANDOM[,"REF"] == "G" & RANDOM[,"ALT"] == "A")))
STAT[4,"N.G>A"] <- length(which((TAIL[,"REF"] == "G" & TAIL[,"ALT"] == "A")))
STAT[5,"N.G>A"] <- length(which((CONSERVATIVE[,"REF"] == "G" & CONSERVATIVE[,"ALT"] == "A")))

STAT[1,"N.C>T"] <- length(which((NEUTRAL[,"REF"] == "C" & NEUTRAL[,"ALT"] == "T")))
STAT[2,"N.C>T"] <- length(which((CAUSAL[,"REF"] == "C" & CAUSAL[,"ALT"] == "T")))
STAT[3,"N.C>T"] <- length(which((RANDOM[,"REF"] == "C" & RANDOM[,"ALT"] == "T")))
STAT[4,"N.C>T"] <- length(which((TAIL[,"REF"] == "C" & TAIL[,"ALT"] == "T")))
STAT[5,"N.C>T"] <- length(which((CONSERVATIVE[,"REF"] == "C" & CONSERVATIVE[,"ALT"] == "T")))

G.MATRIX.NEUTRAL <- matrix(c(unlist(STAT[1,c("N.G>A","N.C>T")]),unlist(STAT[1,c("N.G>A","N.C>T")])),nrow=2,byrow=TRUE)
G.MATRIX.CAUSAL <- matrix(c(unlist(STAT[1,c("N.G>A","N.C>T")]),unlist(STAT[2,c("N.G>A","N.C>T")])),nrow=2,byrow=TRUE)
G.MATRIX.RANDOM <- matrix(c(unlist(STAT[1,c("N.G>A","N.C>T")]),unlist(STAT[3,c("N.G>A","N.C>T")])),nrow=2,byrow=TRUE)
G.MATRIX.TAIL <- matrix(c(unlist(STAT[1,c("N.G>A","N.C>T")]),unlist(STAT[4,c("N.G>A","N.C>T")])),nrow=2,byrow=TRUE)
G.MATRIX.CONSERVATIVE <- matrix(c(unlist(STAT[1,c("N.G>A","N.C>T")]),unlist(STAT[5,c("N.G>A","N.C>T")])),nrow=2,byrow=TRUE)

STAT[,"G.STAT.G>A.vs.C>T"] <- c(likelihood.test(G.MATRIX.NEUTRAL,conservative=FALSE)$statistic,likelihood.test(G.MATRIX.CAUSAL,conservative=FALSE)$statistic,likelihood.test(G.MATRIX.RANDOM,conservative=FALSE)$statistic,likelihood.test(G.MATRIX.TAIL,conservative=FALSE)$statistic,likelihood.test(G.MATRIX.CONSERVATIVE,conservative=FALSE)$statistic)

STAT[,"P.VAL.G>A.vs.C>T"] <- c(pchisq(STAT[1,"G.STAT.G>A.vs.C>T"],df=1,lower.tail=FALSE),pchisq(STAT[2,"G.STAT.G>A.vs.C>T"],df=1,lower.tail=FALSE),pchisq(STAT[3,"G.STAT.G>A.vs.C>T"],df=1,lower.tail=FALSE),pchisq(STAT[4,"G.STAT.G>A.vs.C>T"],df=1,lower.tail=FALSE),pchisq(STAT[5,"G.STAT.G>A.vs.C>T"],df=1,lower.tail=FALSE))


STAT[1,"N.INDELS"] <- length(which(nchar(as.character(NEUTRAL[,"REF"])) != nchar(as.character(NEUTRAL[,"ALT"]))))
STAT[2,"N.INDELS"] <- length(which(nchar(as.character(CAUSAL[,"REF"])) != nchar(as.character(CAUSAL[,"ALT"]))))
STAT[3,"N.INDELS"] <- length(which(nchar(as.character(RANDOM[,"REF"])) != nchar(as.character(RANDOM[,"ALT"]))))
STAT[4,"N.INDELS"] <- length(which(nchar(as.character(TAIL[,"REF"])) != nchar(as.character(TAIL[,"ALT"]))))
STAT[5,"N.INDELS"] <- length(which(nchar(as.character(CONSERVATIVE[,"REF"])) != nchar(as.character(CONSERVATIVE[,"ALT"]))))

STAT[1,"N.NOT.INDELS"] <- STAT[1,"N.MUT"] - STAT[1,"N.INDELS"]
STAT[2,"N.NOT.INDELS"] <- STAT[2,"N.MUT"] - STAT[2,"N.INDELS"]
STAT[3,"N.NOT.INDELS"] <- STAT[3,"N.MUT"] - STAT[3,"N.INDELS"]
STAT[4,"N.NOT.INDELS"] <- STAT[4,"N.MUT"] - STAT[4,"N.INDELS"]
STAT[5,"N.NOT.INDELS"] <- STAT[5,"N.MUT"] - STAT[5,"N.INDELS"]

G.MATRIX.NEUTRAL <- matrix(c(unlist(STAT[1,c("N.INDELS","N.NOT.INDELS")]),unlist(STAT[1,c("N.INDELS","N.NOT.INDELS")])),nrow=2,byrow=TRUE)
G.MATRIX.CAUSAL <- matrix(c(unlist(STAT[1,c("N.INDELS","N.NOT.INDELS")]),unlist(STAT[2,c("N.INDELS","N.NOT.INDELS")])),nrow=2,byrow=TRUE)
G.MATRIX.RANDOM <- matrix(c(unlist(STAT[1,c("N.INDELS","N.NOT.INDELS")]),unlist(STAT[3,c("N.INDELS","N.NOT.INDELS")])),nrow=2,byrow=TRUE)
G.MATRIX.TAIL <- matrix(c(unlist(STAT[1,c("N.INDELS","N.NOT.INDELS")]),unlist(STAT[4,c("N.INDELS","N.NOT.INDELS")])),nrow=2,byrow=TRUE)
G.MATRIX.CONSERVATIVE <- matrix(c(unlist(STAT[1,c("N.INDELS","N.NOT.INDELS")]),unlist(STAT[5,c("N.INDELS","N.NOT.INDELS")])),nrow=2,byrow=TRUE)

STAT[,"G.INDELS.vs.NOT.INDELS"] <- c(likelihood.test(G.MATRIX.NEUTRAL,conservative=FALSE)$statistic,likelihood.test(G.MATRIX.CAUSAL,conservative=FALSE)$statistic,likelihood.test(G.MATRIX.RANDOM,conservative=FALSE)$statistic,likelihood.test(G.MATRIX.TAIL,conservative=FALSE)$statistic,likelihood.test(G.MATRIX.CONSERVATIVE,conservative=FALSE)$statistic)

STAT[,"P.INDELS.vs.NOT.INDELS"] <- c(pchisq(STAT[1,"G.INDELS.vs.NOT.INDELS"],df=1,lower.tail=FALSE),pchisq(STAT[2,"G.INDELS.vs.NOT.INDELS"],df=1,lower.tail=FALSE),pchisq(STAT[3,"G.INDELS.vs.NOT.INDELS"],df=1,lower.tail=FALSE),pchisq(STAT[4,"G.INDELS.vs.NOT.INDELS"],df=1,lower.tail=FALSE),pchisq(STAT[5,"G.INDELS.vs.NOT.INDELS"],df=1,lower.tail=FALSE))


STAT[1,"N.ANEUPLOIDIES"] <- length(which(NEUTRAL$CLASS == "ANEUPLOIDY"))
STAT[2,"N.ANEUPLOIDIES"] <- length(which(CAUSAL$CLASS == "ANEUPLOIDY"))
STAT[3,"N.ANEUPLOIDIES"] <- length(which(RANDOM$CLASS == "ANEUPLOIDY"))
STAT[4,"N.ANEUPLOIDIES"] <- length(which(TAIL$CLASS == "ANEUPLOIDY"))
STAT[5,"N.ANEUPLOIDIES"] <- length(which(CONSERVATIVE$CLASS == "ANEUPLOIDY"))

STAT[1,"N.NOT.ANEUPLOIDIES"] <- STAT[1,"N.MUT"] - STAT[1,"N.ANEUPLOIDIES"]
STAT[2,"N.NOT.ANEUPLOIDIES"] <- STAT[2,"N.MUT"] - STAT[2,"N.ANEUPLOIDIES"]
STAT[3,"N.NOT.ANEUPLOIDIES"] <- STAT[3,"N.MUT"] - STAT[3,"N.ANEUPLOIDIES"]
STAT[4,"N.NOT.ANEUPLOIDIES"] <- STAT[4,"N.MUT"] - STAT[4,"N.ANEUPLOIDIES"]
STAT[5,"N.NOT.ANEUPLOIDIES"] <- STAT[5,"N.MUT"] - STAT[5,"N.ANEUPLOIDIES"]

G.MATRIX.NEUTRAL <- matrix(c(unlist(STAT[1,c("N.ANEUPLOIDIES","N.NOT.ANEUPLOIDIES")]),unlist(STAT[1,c("N.ANEUPLOIDIES","N.NOT.ANEUPLOIDIES")])),nrow=2,byrow=TRUE)
G.MATRIX.CAUSAL <- matrix(c(unlist(STAT[1,c("N.ANEUPLOIDIES","N.NOT.ANEUPLOIDIES")]),unlist(STAT[2,c("N.ANEUPLOIDIES","N.NOT.ANEUPLOIDIES")])),nrow=2,byrow=TRUE)
G.MATRIX.RANDOM <- matrix(c(unlist(STAT[1,c("N.ANEUPLOIDIES","N.NOT.ANEUPLOIDIES")]),unlist(STAT[3,c("N.ANEUPLOIDIES","N.NOT.ANEUPLOIDIES")])),nrow=2,byrow=TRUE)
G.MATRIX.TAIL <- matrix(c(unlist(STAT[1,c("N.ANEUPLOIDIES","N.NOT.ANEUPLOIDIES")]),unlist(STAT[4,c("N.ANEUPLOIDIES","N.NOT.ANEUPLOIDIES")])),nrow=2,byrow=TRUE)
G.MATRIX.CONSERVATIVE <- matrix(c(unlist(STAT[1,c("N.ANEUPLOIDIES","N.NOT.ANEUPLOIDIES")]),unlist(STAT[5,c("N.ANEUPLOIDIES","N.NOT.ANEUPLOIDIES")])),nrow=2,byrow=TRUE)

STAT[,"G.ANEUPLOIDIES"] <- c(likelihood.test(G.MATRIX.NEUTRAL,conservative=FALSE)$statistic,likelihood.test(G.MATRIX.CAUSAL,conservative=FALSE)$statistic,likelihood.test(G.MATRIX.RANDOM,conservative=FALSE)$statistic,likelihood.test(G.MATRIX.TAIL,conservative=FALSE)$statistic,likelihood.test(G.MATRIX.CONSERVATIVE,conservative=FALSE)$statistic)

STAT[,"P.ANEUPLOIDIES"] <- c(pchisq(STAT[1,"G.ANEUPLOIDIES"],df=1,lower.tail=FALSE),pchisq(STAT[2,"G.ANEUPLOIDIES"],df=1,lower.tail=FALSE),pchisq(STAT[3,"G.ANEUPLOIDIES"],df=1,lower.tail=FALSE),pchisq(STAT[4,"G.ANEUPLOIDIES"],df=1,lower.tail=FALSE),pchisq(STAT[5,"G.ANEUPLOIDIES"],df=1,lower.tail=FALSE))

write.table(STAT,"Statistics/STAT.MUTATION.TYPES.txt",sep="\t",row.names=FALSE)


###Barplot mutation types CAUSAL vs NEUTRAL (Figure 3A).###

#Create matrix with number of mutations of different types.
MUT.COUNTS <- matrix(rep(0,10),nrow=2)

rownames(MUT.COUNTS) <- c("NEUTRAL","CAUSAL")
#Types of mutations considered
colnames(MUT.COUNTS) <- c("C->T","G->A","Other substitutions","Indels","Aneuploidies")

MUT.COUNTS[1,1] <- length(which(NEUTRAL[,"REF"] == "C" & NEUTRAL[,"ALT"] == "T"))
MUT.COUNTS[1,2] <- length(which(NEUTRAL[,"REF"] == "G" & NEUTRAL[,"ALT"] == "A"))
MUT.COUNTS[1,4] <- length(which(NEUTRAL[,"TYPE.1"] == "INDEL"))
MUT.COUNTS[1,5] <- length(which(NEUTRAL[,"TYPE.1"] == "DUPLICATION"))
MUT.COUNTS[1,3] <- nrow(NEUTRAL) - sum(MUT.COUNTS[1,])

MUT.COUNTS[2,1] <- length(which(CAUSAL[,"REF"] == "C" & CAUSAL[,"ALT"] == "T"))
MUT.COUNTS[2,2] <- length(which(CAUSAL[,"REF"] == "G" & CAUSAL[,"ALT"] == "A"))
MUT.COUNTS[2,4] <- length(which(CAUSAL[,"TYPE.1"] == "INDEL"))
MUT.COUNTS[2,5] <- length(which(CAUSAL[,"TYPE.1"] == "DUPLICATION"))
MUT.COUNTS[2,3] <- nrow(CAUSAL) - sum(MUT.COUNTS[2,])

MUT.FREQ <- MUT.COUNTS
MUT.FREQ[1,] <- MUT.FREQ[1,]/sum(MUT.FREQ[1,])
MUT.FREQ[2,] <- MUT.FREQ[2,]/sum(MUT.FREQ[2,])

pdf("Figure 3A.pdf",useDingbats=F,height=7,width=12)
#windows(height=7,width=12)
barplot(MUT.FREQ, col=c("blue","red"), beside=TRUE, ylim=c(0,0.5),legend=rownames(MUT.FREQ))
dev.off()


########################################################################################
#I) Genomic distributions of trans-regulatory and non-regulatory mutations (Figure 3B).#
########################################################################################

#"S288c.genome.bed" is a table listing all nucleotides found at each position of each chromosome in the yeast genome.
#The first 10000 lines of this file ("S288c.genome.truncated.bed") can be found in SupplementaryFile11.zip.
#"S288c.genome.bed" was generated from the reference genome ("S288c.mapping.fsa") included in SupplementaryFile11.zip.
CHROM <- read.table("S288c.genome.bed",sep="\t",header=TRUE)

CHROM <- subset(CHROM, CHR %nin% c("chrMito","chrYFP","chrKan","chrNat"))
CHROM <- droplevels(CHROM)
UNIQUE.CHR <- unique(CHROM$CHR)

CHR.LENGTH <- rep(0,length(UNIQUE.CHR))
GENOME.POSITION <- rep(0,length(UNIQUE.CHR))

#Dataframe with the length of each yeast chromosome.
CHROM.POSITIONS <- data.frame(UNIQUE.CHR, CHR.LENGTH, GENOME.POSITION) 

for (i in 1:nrow(CHROM.POSITIONS))
{
  CHROM.POSITIONS[i,"CHR.LENGTH"] <- max(subset(CHROM, CHR == CHROM.POSITIONS[i,"UNIQUE.CHR"])[,"POSITION"],na.rm=T)
  CHROM.POSITIONS[i,"GENOME.POSITION"] <- sum(CHROM.POSITIONS[,"CHR.LENGTH"])
}

SUB.NEUTRAL <- subset(NEUTRAL, CHROMOSOME %nin% c("chrMito","chrYFP","chrKan","chrNat"))

BREAKS <- seq(0,max(ALL.MUT$GENOME.POSITION,na.rm=T)+100000,100000)

BREAKS <- seq(0,CHROM.POSITIONS[1,"GENOME.POSITION"],length.out=round(CHROM.POSITIONS[1,"CHR.LENGTH"]*100/12071326)+1)
for (i in 2:nrow(CHROM.POSITIONS))
{
  CUR.BREAKS <- seq(CHROM.POSITIONS[i-1,"GENOME.POSITION"],CHROM.POSITIONS[i,"GENOME.POSITION"],length.out=round(CHROM.POSITIONS[i,"CHR.LENGTH"]*100/12071326)+1)
  BREAKS <- c(BREAKS,CUR.BREAKS)
}


###Plotting the genomic distributions of mutations.###

pdf("Figure 3B.pdf",height=7,width=12,useDingbats=FALSE)
#windows(height=7,width=12)
hist(SUB.NEUTRAL[,"GENOME.POSITION"],breaks=BREAKS,col="blue",border=NA,freq=TRUE,xlab="Position in Genome",ylab="# Sites",main="Neutral vs All Causal Mutations",xaxt="n")
hist(CAUSAL[,"GENOME.POSITION"],breaks=BREAKS,col="green",border=NA,freq=TRUE,add=TRUE)
hist(RANDOM[,"GENOME.POSITION"],breaks=BREAKS,col="red",border=NA,freq=TRUE,add=TRUE)
abline(v=CHROM.POSITIONS[,"GENOME.POSITION"],lty=2,col="#00000066")
legend("topright",legend=c("Neutral","Causal"),box.lty=0,fill=c("blue","orange"))
axis(1,at=c(0,CHROM.POSITIONS[,"GENOME.POSITION"]))
dev.off()


###Statistical tests to compare the genomic distributions of trans-regulatory and non-regulatory mutations.###

STAT <- data.frame(MUT.TYPE.2, N.MUT.2)

STAT[,"MEAN.CHROMOSOME.POSITION"] <- c(mean(NEUTRAL[,"CHROMOSOME.POSITION"],na.rm=TRUE),mean(CAUSAL[,"CHROMOSOME.POSITION"],na.rm=TRUE),mean(RANDOM[,"CHROMOSOME.POSITION"],na.rm=TRUE),mean(TAIL[,"CHROMOSOME.POSITION"],na.rm=TRUE),mean(CONSERVATIVE[,"CHROMOSOME.POSITION"],na.rm=TRUE),mean(BSA.SEQ[,"CHROMOSOME.POSITION"],na.rm=TRUE),mean(SANGER[,"CHROMOSOME.POSITION"],na.rm=TRUE))
STAT[,"SD.CHROMOSOME.POSITION"] <- c(sd(NEUTRAL[,"CHROMOSOME.POSITION"],na.rm=TRUE),sd(CAUSAL[,"CHROMOSOME.POSITION"],na.rm=TRUE),sd(RANDOM[,"CHROMOSOME.POSITION"],na.rm=TRUE),sd(TAIL[,"CHROMOSOME.POSITION"],na.rm=TRUE),sd(CONSERVATIVE[,"CHROMOSOME.POSITION"],na.rm=TRUE),sd(BSA.SEQ[,"CHROMOSOME.POSITION"],na.rm=TRUE),sd(SANGER[,"CHROMOSOME.POSITION"],na.rm=TRUE))

STAT[,"WILCOX.MEAN.CHROMOSOME.POSITION"] <-  c(wilcox.test(SUBSTITUTION[,"CHROMOSOME.POSITION"],SUBSTITUTION[,"CHROMOSOME.POSITION"])$statistic,wilcox.test(SUBSTITUTION[,"CHROMOSOME.POSITION"],CAUSAL[,"CHROMOSOME.POSITION"])$statistic,wilcox.test(SUBSTITUTION[,"CHROMOSOME.POSITION"],RANDOM[,"CHROMOSOME.POSITION"])$statistic,wilcox.test(SUBSTITUTION[,"CHROMOSOME.POSITION"],TAIL[,"CHROMOSOME.POSITION"])$statistic,wilcox.test(SUBSTITUTION[,"CHROMOSOME.POSITION"],CONSERVATIVE[,"CHROMOSOME.POSITION"])$statistic,wilcox.test(SUBSTITUTION[,"CHROMOSOME.POSITION"],BSA.SEQ[,"CHROMOSOME.POSITION"])$statistic,wilcox.test(SUBSTITUTION[,"CHROMOSOME.POSITION"],SANGER[,"CHROMOSOME.POSITION"])$statistic)
STAT[,"P.MEAN.CHROMOSOME.POSITION"] <-  c(wilcox.test(SUBSTITUTION[,"CHROMOSOME.POSITION"],SUBSTITUTION[,"CHROMOSOME.POSITION"])$p.value,wilcox.test(SUBSTITUTION[,"CHROMOSOME.POSITION"],CAUSAL[,"CHROMOSOME.POSITION"])$p.value,wilcox.test(SUBSTITUTION[,"CHROMOSOME.POSITION"],RANDOM[,"CHROMOSOME.POSITION"])$p.value,wilcox.test(SUBSTITUTION[,"CHROMOSOME.POSITION"],TAIL[,"CHROMOSOME.POSITION"])$p.value,wilcox.test(SUBSTITUTION[,"CHROMOSOME.POSITION"],CONSERVATIVE[,"CHROMOSOME.POSITION"])$p.value,wilcox.test(SUBSTITUTION[,"CHROMOSOME.POSITION"],BSA.SEQ[,"CHROMOSOME.POSITION"])$p.value,wilcox.test(SUBSTITUTION[,"CHROMOSOME.POSITION"],SANGER[,"CHROMOSOME.POSITION"])$p.value)

STAT[,"LEVENE.CHROMOSOME.POSITION"] <- c(0,leveneTest(CHROMOSOME.POSITION ~ MUTATION, data = SUB.CAUSAL)$F[1],leveneTest(CHROMOSOME.POSITION ~ MUTATION, data = SUB.RANDOM)$F[1],leveneTest(CHROMOSOME.POSITION ~ MUTATION, data = SUB.TAIL)$F[1],leveneTest(CHROMOSOME.POSITION ~ MUTATION, data = SUB.CONS)$F[1],NA,NA)
STAT[,"P.SD.CHROMOSOME.POSITION"] <- c(1,leveneTest(CHROMOSOME.POSITION ~ MUTATION, data = SUB.CAUSAL)$P[1],leveneTest(CHROMOSOME.POSITION ~ MUTATION, data = SUB.RANDOM)$P[1],leveneTest(CHROMOSOME.POSITION ~ MUTATION, data = SUB.TAIL)$P[1],leveneTest(CHROMOSOME.POSITION ~ MUTATION, data = SUB.CONS)$P[1],NA,NA)

STAT[,"KS.CHROMOSOME.POSITION"] <- c(ks.test(NEUTRAL[,"CHROMOSOME.POSITION"],NEUTRAL[,"CHROMOSOME.POSITION"])$statistic,ks.test(NEUTRAL[,"CHROMOSOME.POSITION"],CAUSAL[,"CHROMOSOME.POSITION"])$statistic,ks.test(NEUTRAL[,"CHROMOSOME.POSITION"],RANDOM[,"CHROMOSOME.POSITION"])$statistic,ks.test(NEUTRAL[,"CHROMOSOME.POSITION"],TAIL[,"CHROMOSOME.POSITION"])$statistic,ks.test(NEUTRAL[,"CHROMOSOME.POSITION"],CONSERVATIVE[,"CHROMOSOME.POSITION"])$statistic,ks.test(NEUTRAL[,"CHROMOSOME.POSITION"],BSA.SEQ[,"CHROMOSOME.POSITION"])$statistic,ks.test(NEUTRAL[,"CHROMOSOME.POSITION"],SANGER[,"CHROMOSOME.POSITION"])$statistic)
STAT[,"P.KS.CHROMOSOME.POSITION"] <- c(ks.test(NEUTRAL[,"CHROMOSOME.POSITION"],NEUTRAL[,"CHROMOSOME.POSITION"])$p.value,ks.test(NEUTRAL[,"CHROMOSOME.POSITION"],CAUSAL[,"CHROMOSOME.POSITION"])$p.value,ks.test(NEUTRAL[,"CHROMOSOME.POSITION"],RANDOM[,"CHROMOSOME.POSITION"])$p.value,ks.test(NEUTRAL[,"CHROMOSOME.POSITION"],TAIL[,"CHROMOSOME.POSITION"])$p.value,ks.test(NEUTRAL[,"CHROMOSOME.POSITION"],CONSERVATIVE[,"CHROMOSOME.POSITION"])$p.value,ks.test(NEUTRAL[,"CHROMOSOME.POSITION"],BSA.SEQ[,"CHROMOSOME.POSITION"])$p.value,ks.test(NEUTRAL[,"CHROMOSOME.POSITION"],SANGER[,"CHROMOSOME.POSITION"])$p.value)

#write.table(STAT,"Statistics/STAT.CHROMOSOME.POSITION.txt",sep="\t",row.names=FALSE)


##########################################################################################
#J) Proportion of mutations observed on each chromosome (Figure 3 - figure supplement 1).#
##########################################################################################

CHR.COUNTS <- matrix(rep(0,64),nrow=4)

rownames(CHR.COUNTS) <- c("NEUTRAL","CAUSAL","RANDOM","TAIL")
colnames(CHR.COUNTS) <- c("chr. I","chr. II","chr. III","chr. IV","chr. V","chr. VI","chr. VII","chr. VIII","chr. IX","chr. X","chr. XI","chr. XII","chr. XIII","chr. XIV","chr. XV","chr. XVI")

CHR.NAMES <- CHROM.POSITIONS$UNIQUE.CHR

for (i in 1:length(CHR.NAMES))
{
  CHR.COUNTS[1,i] <- length(which(NEUTRAL$CHROMOSOME == as.character(CHR.NAMES[i])))
  CHR.COUNTS[2,i] <- length(which(CAUSAL$CHROMOSOME == as.character(CHR.NAMES[i])))
  CHR.COUNTS[3,i] <- length(which(RANDOM$CHROMOSOME == as.character(CHR.NAMES[i])))
  CHR.COUNTS[4,i] <- length(which(TAIL$CHROMOSOME == as.character(CHR.NAMES[i])))
}

CHR.FREQ <- CHR.COUNTS
CHR.FREQ[1,] <- CHR.COUNTS[1,]/sum(CHR.COUNTS[1,])
CHR.FREQ[2,] <- CHR.COUNTS[2,]/sum(CHR.COUNTS[2,])
CHR.FREQ[3,] <- CHR.COUNTS[3,]/sum(CHR.COUNTS[3,])
CHR.FREQ[4,] <- CHR.COUNTS[4,]/sum(CHR.COUNTS[4,])

pdf("Figure 3 - figure supplement 1.pdf",useDingbats=F,height=7,width=12)
#windows(height=7,width=12)
barplot(CHR.FREQ, col=c("blue","orange","red","green"), beside=TRUE, ylim=c(0,0.5),legend=rownames(CHR.FREQ))
dev.off()

###G-tests to compare the distributions of trans-regulatory and non-regulatory mutations across chromosomes.### 

STAT <- data.frame(MUT.TYPE.2, N.MUT.2)

CHR.NEUTRAL <- table(NEUTRAL[,"CHROMOSOME"])
CHR.CAUSAL <- table(CAUSAL[,"CHROMOSOME"])
CHR.RANDOM <- table(RANDOM[,"CHROMOSOME"])
CHR.TAIL <- table(TAIL[,"CHROMOSOME"])
CHR.CONS <- table(CONSERVATIVE[,"CHROMOSOME"])
CHR.BSA.SEQ <- table(BSA.SEQ[,"CHROMOSOME"])
CHR.SANGER <- table(SANGER[,"CHROMOSOME"])

CHR <- rbind(CHR.NEUTRAL, CHR.CAUSAL, CHR.RANDOM, CHR.TAIL, CHR.CONS, CHR.BSA.SEQ, CHR.SANGER)

STAT <- cbind(STAT,CHR)
rownames(STAT) <- c()

G.MATRIX.NEUTRAL <- matrix(c(unlist(STAT[1,c(3:18)]),unlist(STAT[1,c(3:18)])),nrow=2,byrow=TRUE)
G.MATRIX.CAUSAL <- matrix(c(unlist(STAT[1,c(3:18)]),unlist(STAT[2,c(3:18)])),nrow=2,byrow=TRUE)
G.MATRIX.RANDOM <- matrix(c(unlist(STAT[1,c(3:18)]),unlist(STAT[3,c(3:18)])),nrow=2,byrow=TRUE)
G.MATRIX.TAIL <- matrix(c(unlist(STAT[1,c(3:18)]),unlist(STAT[4,c(3:18)])),nrow=2,byrow=TRUE)
G.MATRIX.CONSERVATIVE <- matrix(c(unlist(STAT[1,c(3:18)]),unlist(STAT[5,c(3:18)])),nrow=2,byrow=TRUE)
G.MATRIX.BSA.SEQ <- matrix(c(unlist(STAT[1,c(3:18)]),unlist(STAT[6,c(3:18)])),nrow=2,byrow=TRUE)
G.MATRIX.SANGER <- matrix(c(unlist(STAT[1,c(3:18)]),unlist(STAT[7,c(3:18)])),nrow=2,byrow=TRUE)

STAT[,"G.STAT.CHROMOSOME"] <- c(likelihood.test(G.MATRIX.NEUTRAL,conservative=FALSE)$statistic,likelihood.test(G.MATRIX.CAUSAL,conservative=FALSE)$statistic,likelihood.test(G.MATRIX.RANDOM,conservative=FALSE)$statistic,likelihood.test(G.MATRIX.TAIL,conservative=FALSE)$statistic,likelihood.test(G.MATRIX.CONSERVATIVE,conservative=FALSE)$statistic,likelihood.test(G.MATRIX.BSA.SEQ,conservative=FALSE)$statistic,likelihood.test(G.MATRIX.SANGER,conservative=FALSE)$statistic)

STAT[,"P.VAL.CHROMOSOME"] <- c(pchisq(STAT[1,"G.STAT.CHROMOSOME"],df=15,lower.tail=FALSE),pchisq(STAT[2,"G.STAT.CHROMOSOME"],df=15,lower.tail=FALSE),pchisq(STAT[3,"G.STAT.CHROMOSOME"],df=15,lower.tail=FALSE),pchisq(STAT[4,"G.STAT.CHROMOSOME"],df=15,lower.tail=FALSE),pchisq(STAT[5,"G.STAT.CHROMOSOME"],df=15,lower.tail=FALSE),pchisq(STAT[6,"G.STAT.CHROMOSOME"],df=15,lower.tail=FALSE),pchisq(STAT[7,"G.STAT.CHROMOSOME"],df=15,lower.tail=FALSE))

#ENRICHMENT ON EACH CHR
for (i in 1:nrow(STAT))
{
  for (j in 3:20)
  {
    STAT[i,j+20] <- round(((STAT[i,j]/sum(STAT[i,3:20]) )- (STAT[1,j]/sum(STAT[1,3:20]))) / (STAT[1,j]/sum(STAT[1,3:20])),digits=2)
  }
}

colnames(STAT)[23:40] <- paste("Enr_",colnames(STAT[,c(3:20)]),sep="")

#write.table(STAT,"Statistics/STAT.MUTATION.CHROMOSOMES.txt",sep="\t",row.names=FALSE)

#Repeat analysis without mutations on chromosomes 7 and 13.
STAT.NO.7.13 <- STAT[,1:21]
STAT.NO.7.13 <- STAT.NO.7.13[,-c(9,15)]

for (i in 1:nrow(STAT.NO.7.13))
{
  STAT.NO.7.13[i,"N.MUT.2"] <- sum(STAT.NO.7.13[i,c(3:16)])
}

G.MATRIX.NEUTRAL <- matrix(c(unlist(STAT.NO.7.13[1,c(3:16)]),unlist(STAT.NO.7.13[1,c(3:16)])),nrow=2,byrow=TRUE)
G.MATRIX.CAUSAL <- matrix(c(unlist(STAT.NO.7.13[1,c(3:16)]),unlist(STAT.NO.7.13[2,c(3:16)])),nrow=2,byrow=TRUE)
G.MATRIX.RANDOM <- matrix(c(unlist(STAT.NO.7.13[1,c(3:16)]),unlist(STAT.NO.7.13[3,c(3:16)])),nrow=2,byrow=TRUE)
G.MATRIX.TAIL <- matrix(c(unlist(STAT.NO.7.13[1,c(3:16)]),unlist(STAT.NO.7.13[4,c(3:16)])),nrow=2,byrow=TRUE)
G.MATRIX.CONSERVATIVE <- matrix(c(unlist(STAT.NO.7.13[1,c(3:16)]),unlist(STAT.NO.7.13[5,c(3:16)])),nrow=2,byrow=TRUE)
G.MATRIX.BSA.SEQ <- matrix(c(unlist(STAT.NO.7.13[1,c(3:16)]),unlist(STAT.NO.7.13[6,c(3:16)])),nrow=2,byrow=TRUE)
G.MATRIX.SANGER <- matrix(c(unlist(STAT.NO.7.13[1,c(3:16)]),unlist(STAT.NO.7.13[7,c(3:16)])),nrow=2,byrow=TRUE)

STAT.NO.7.13[,"G.STAT.CHROMOSOME"] <- c(likelihood.test(G.MATRIX.NEUTRAL,conservative=FALSE)$statistic,likelihood.test(G.MATRIX.CAUSAL,conservative=FALSE)$statistic,likelihood.test(G.MATRIX.RANDOM,conservative=FALSE)$statistic,likelihood.test(G.MATRIX.TAIL,conservative=FALSE)$statistic,likelihood.test(G.MATRIX.CONSERVATIVE,conservative=FALSE)$statistic,likelihood.test(G.MATRIX.BSA.SEQ,conservative=FALSE)$statistic,likelihood.test(G.MATRIX.SANGER,conservative=FALSE)$statistic)

STAT.NO.7.13[,"P.VAL.CHROMOSOME"] <- c(pchisq(STAT.NO.7.13[1,"G.STAT.CHROMOSOME"],df=15,lower.tail=FALSE),pchisq(STAT.NO.7.13[2,"G.STAT.CHROMOSOME"],df=15,lower.tail=FALSE),pchisq(STAT.NO.7.13[3,"G.STAT.CHROMOSOME"],df=15,lower.tail=FALSE),pchisq(STAT.NO.7.13[4,"G.STAT.CHROMOSOME"],df=15,lower.tail=FALSE),pchisq(STAT.NO.7.13[5,"G.STAT.CHROMOSOME"],df=15,lower.tail=FALSE),pchisq(STAT.NO.7.13[6,"G.STAT.CHROMOSOME"],df=15,lower.tail=FALSE),pchisq(STAT.NO.7.13[7,"G.STAT.CHROMOSOME"],df=15,lower.tail=FALSE))

#Do G-tests to test enrichment of mutations on chromosomes 7 and 13.

G.MATRIX.7 <- matrix(c(164,17,1766-164,70-17),nrow=2,byrow=TRUE)
G.MATRIX.13 <- matrix(c(138,9,1766-138,70-9),nrow=2,byrow=TRUE)

G.7 <- likelihood.test(G.MATRIX.7,conservative=FALSE)$statistic
G.13 <- likelihood.test(G.MATRIX.13,conservative=FALSE)$statistic

P.7 <- pchisq(G.7,df=1,lower.tail=FALSE)
P.13 <- pchisq(G.13,df=1,lower.tail=FALSE)


#######################################################################################################################################
#K) Comparing the proportions of trans-regulatory and non regulatory mutations in coding, intergenic and intronic regions (Figure 3C).#
#######################################################################################################################################

#Exclude aneuploidies
CAUSAL <- subset(CAUSAL, CLASS != "ANEUPLOIDY")
RANDOM <- subset(RANDOM, CLASS != "ANEUPLOIDY")
CONSERVATIVE <- subset(CONSERVATIVE, CLASS != "ANEUPLOIDY")

#Statistical tests (G-tests)
N.MUT <- c(nrow(NEUTRAL),nrow(CAUSAL),nrow(RANDOM),nrow(TAIL),nrow(CONSERVATIVE))

N.EXONIC <- c(nrow(subset(NEUTRAL, CLASS == "EXONIC")),nrow(subset(CAUSAL, CLASS == "EXONIC")),nrow(subset(RANDOM, CLASS == "EXONIC")),nrow(subset(TAIL, CLASS == "EXONIC")),nrow(subset(CONSERVATIVE, CLASS == "EXONIC")))
N.INTRONIC <- c(nrow(subset(NEUTRAL, CLASS == "INTRONIC")),nrow(subset(CAUSAL, CLASS == "INTRONIC")),nrow(subset(RANDOM, CLASS == "INTRONIC")),nrow(subset(TAIL, CLASS == "INTRONIC")),nrow(subset(CONSERVATIVE, CLASS == "INTRONIC")))
N.INTERGENIC <- c(nrow(subset(NEUTRAL, CLASS == "INTERGENIC")),nrow(subset(CAUSAL, CLASS == "INTERGENIC")),nrow(subset(RANDOM, CLASS == "INTERGENIC")),nrow(subset(TAIL, CLASS == "INTERGENIC")),nrow(subset(CONSERVATIVE, CLASS == "INTERGENIC")))

STAT <- data.frame(MUT.TYPE, N.MUT, N.EXONIC, N.INTRONIC, N.INTERGENIC)

STAT[,"FREQ.EXONIC"] <- STAT[,"N.EXONIC"] / (STAT[,"N.EXONIC"] + STAT[,"N.INTRONIC"] + STAT[,"N.INTERGENIC"] )
STAT[,"FREQ.INTRONIC"] <- STAT[,"N.INTRONIC"] / (STAT[,"N.EXONIC"] + STAT[,"N.INTRONIC"] + STAT[,"N.INTERGENIC"] )
STAT[,"FREQ.INTERGENIC"] <- STAT[,"N.INTERGENIC"] / (STAT[,"N.EXONIC"] + STAT[,"N.INTRONIC"] + STAT[,"N.INTERGENIC"] )

G.MATRIX.NEUTRAL <- matrix(c(STAT[1,"N.EXONIC"],STAT[1,"N.INTERGENIC"],STAT[1,"N.EXONIC"],STAT[1,"N.INTERGENIC"]),nrow=2,ncol=2)
G.MATRIX.CAUSAL <- matrix(c(STAT[1,"N.EXONIC"],STAT[1,"N.INTERGENIC"],STAT[2,"N.EXONIC"],STAT[2,"N.INTERGENIC"]),nrow=2,ncol=2)
G.MATRIX.RANDOM <- matrix(c(STAT[1,"N.EXONIC"],STAT[1,"N.INTERGENIC"],STAT[3,"N.EXONIC"],STAT[3,"N.INTERGENIC"]),nrow=2,ncol=2)
G.MATRIX.TAIL <- matrix(c(STAT[1,"N.EXONIC"],STAT[1,"N.INTERGENIC"],STAT[4,"N.EXONIC"],STAT[4,"N.INTERGENIC"]),nrow=2,ncol=2)
G.MATRIX.CONSERVATIVE <- matrix(c(STAT[1,"N.EXONIC"],STAT[1,"N.INTERGENIC"],STAT[5,"N.EXONIC"],STAT[5,"N.INTERGENIC"]),nrow=2,ncol=2)

STAT[,"G.STAT.CODING"] <- c(likelihood.test(G.MATRIX.NEUTRAL,conservative=TRUE)$statistic,likelihood.test(G.MATRIX.CAUSAL,conservative=TRUE)$statistic,likelihood.test(G.MATRIX.RANDOM,conservative=TRUE)$statistic,likelihood.test(G.MATRIX.TAIL,conservative=TRUE)$statistic,likelihood.test(G.MATRIX.CONSERVATIVE,conservative=TRUE)$statistic)

STAT[,"P.VAL.CODING"] <- c(pchisq(STAT[1,"G.STAT.CODING"],df=1,lower.tail=FALSE),pchisq(STAT[2,"G.STAT.CODING"],df=1,lower.tail=FALSE),pchisq(STAT[3,"G.STAT.CODING"],df=1,lower.tail=FALSE),pchisq(STAT[4,"G.STAT.CODING"],df=1,lower.tail=FALSE),pchisq(STAT[5,"G.STAT.CODING"],df=1,lower.tail=FALSE))

G.MATRIX.NEUTRAL <- matrix(c(STAT[1,"N.INTRONIC"],STAT[1,"N.INTERGENIC"]+STAT[1,"N.EXONIC"],STAT[1,"N.INTRONIC"],STAT[1,"N.INTERGENIC"]+STAT[1,"N.EXONIC"]),nrow=2,ncol=2)
G.MATRIX.CAUSAL <- matrix(c(STAT[1,"N.INTRONIC"],STAT[1,"N.INTERGENIC"]+STAT[1,"N.EXONIC"],STAT[2,"N.INTRONIC"],STAT[2,"N.INTERGENIC"]+STAT[2,"N.EXONIC"]),nrow=2,ncol=2)
G.MATRIX.RANDOM <- matrix(c(STAT[1,"N.INTRONIC"],STAT[1,"N.INTERGENIC"]+STAT[1,"N.EXONIC"],STAT[3,"N.INTRONIC"],STAT[3,"N.INTERGENIC"]+STAT[3,"N.EXONIC"]),nrow=2,ncol=2)
G.MATRIX.TAIL <- matrix(c(STAT[1,"N.INTRONIC"],STAT[1,"N.INTERGENIC"]+STAT[1,"N.EXONIC"],STAT[4,"N.INTRONIC"],STAT[4,"N.INTERGENIC"]+STAT[4,"N.EXONIC"]),nrow=2,ncol=2)
G.MATRIX.CONSERVATIVE <- matrix(c(STAT[1,"N.INTRONIC"],STAT[1,"N.INTERGENIC"]+STAT[1,"N.EXONIC"],STAT[5,"N.INTRONIC"],STAT[5,"N.INTERGENIC"]+STAT[5,"N.EXONIC"]),nrow=2,ncol=2)

STAT[,"G.STAT.INTRONIC"] <- c(likelihood.test(G.MATRIX.NEUTRAL,conservative=TRUE)$statistic,likelihood.test(G.MATRIX.CAUSAL,conservative=TRUE)$statistic,likelihood.test(G.MATRIX.RANDOM,conservative=TRUE)$statistic,likelihood.test(G.MATRIX.TAIL,conservative=TRUE)$statistic,likelihood.test(G.MATRIX.CONSERVATIVE,conservative=TRUE)$statistic)

STAT[,"P.VAL.INTRONIC"] <- c(pchisq(STAT[1,"G.STAT.INTRONIC"],df=1,lower.tail=FALSE),pchisq(STAT[2,"G.STAT.INTRONIC"],df=1,lower.tail=FALSE),pchisq(STAT[3,"G.STAT.INTRONIC"],df=1,lower.tail=FALSE),pchisq(STAT[4,"G.STAT.INTRONIC"],df=1,lower.tail=FALSE),pchisq(STAT[5,"G.STAT.INTRONIC"],df=1,lower.tail=FALSE))

#write.table(STAT,"Statistics/STAT.CODING.vs.NON.CODING.txt",sep="\t",row.names=FALSE)

#Pie charts
PIE <- MUT.TYPE[6:7,]
PIE[,"REF"] <- as.character(PIE[,"REF"])
PIE[,"ALT"] <- as.character(PIE[,"ALT"])
PIE[3,"REF"] <- "Other"
PIE[3,"ALT"] <- "Other"
PIE[3,"N.Mut"] <- 1820-891-812
PIE[3,"Percent.Mut"] <- round((1820-891-812)/1820,digits=3)*100

pdf("Figure 3C Non-regulatory.pdf",width=6,height=6,useDingbats = F)
#windows(width=6,height=6)
pie(c(STAT[1,"N.EXONIC"],STAT[1,"N.INTRONIC"],STAT[1,"N.INTERGENIC"]), labels = c("Coding","Intron","Intergenic"), border="white", col=c("blue","black","gray"),init.angle=90)
dev.off()

pdf("Figure 3C Trans-regulatory.pdf",width=6,height=6,useDingbats = F)
#windows(width=6,height=6)
pie(c(STAT[2,"N.EXONIC"],STAT[2,"N.INTRONIC"],STAT[2,"N.INTERGENIC"]), labels = c("Coding","Intron","Intergenic"), border="white", col=c("blue","black","gray"),init.angle=90)
dev.off()


###################################################################################################################################################
#L) Comparing the proportions of trans-regulatory and non-regulatory mutations being synonymous, nonsynonymous and nonsense mutations (Figure 3D).#
###################################################################################################################################################

#Statistical tests (G-tests)
N.SYNONYMOUS <- c(nrow(subset(NEUTRAL, TYPE.1 == "SYNONYMOUS")),nrow(subset(CAUSAL, TYPE.1 == "SYNONYMOUS")),nrow(subset(RANDOM, TYPE.1 == "SYNONYMOUS")),nrow(subset(TAIL, TYPE.1 == "SYNONYMOUS")),nrow(subset(CONSERVATIVE, TYPE.1 == "SYNONYMOUS")))
N.NON.SYNONYMOUS <- c(nrow(subset(NEUTRAL, TYPE.1 == "NON.SYNONYMOUS")),nrow(subset(CAUSAL, TYPE.1 == "NON.SYNONYMOUS")),nrow(subset(RANDOM, TYPE.1 == "NON.SYNONYMOUS")),nrow(subset(TAIL, TYPE.1 == "NON.SYNONYMOUS")),nrow(subset(CONSERVATIVE, TYPE.1 == "NON.SYNONYMOUS")))
N.NON.SENSE <- c(nrow(subset(NEUTRAL, TYPE.1 == "NON.SENSE")),nrow(subset(CAUSAL, TYPE.1 == "NON.SENSE")),nrow(subset(RANDOM, TYPE.1 == "NON.SENSE")),nrow(subset(TAIL, TYPE.1 == "NON.SENSE")),nrow(subset(CONSERVATIVE, TYPE.1 == "NON.SENSE")))

STAT <- data.frame(MUT.TYPE, N.MUT, N.SYNONYMOUS, N.NON.SYNONYMOUS, N.NON.SENSE)

STAT[,"FREQ.SYNONYMOUS"] <- STAT[,"N.SYNONYMOUS"] / (STAT[,"N.SYNONYMOUS"] + STAT[,"N.NON.SYNONYMOUS"] + STAT[,"N.NON.SENSE"] )
STAT[,"FREQ.NON.SYNONYMOUS"] <- STAT[,"N.NON.SYNONYMOUS"] / (STAT[,"N.SYNONYMOUS"] + STAT[,"N.NON.SYNONYMOUS"] + STAT[,"N.NON.SENSE"] )
STAT[,"FREQ.NON.SENSE"] <- STAT[,"N.NON.SENSE"] / (STAT[,"N.SYNONYMOUS"] + STAT[,"N.NON.SYNONYMOUS"] + STAT[,"N.NON.SENSE"] )

G.MATRIX.NEUTRAL <- matrix(c(STAT[1,"N.SYNONYMOUS"],STAT[1,"N.NON.SYNONYMOUS"],STAT[1,"N.NON.SENSE"],STAT[1,"N.SYNONYMOUS"],STAT[1,"N.NON.SYNONYMOUS"],STAT[1,"N.NON.SENSE"]),nrow=2,ncol=3,byrow=TRUE)
G.MATRIX.CAUSAL <- matrix(c(STAT[1,"N.SYNONYMOUS"],STAT[1,"N.NON.SYNONYMOUS"],STAT[1,"N.NON.SENSE"],STAT[2,"N.SYNONYMOUS"],STAT[2,"N.NON.SYNONYMOUS"],STAT[2,"N.NON.SENSE"]),nrow=2,ncol=3,byrow=TRUE)
G.MATRIX.RANDOM <- matrix(c(STAT[1,"N.SYNONYMOUS"],STAT[1,"N.NON.SYNONYMOUS"],STAT[1,"N.NON.SENSE"],STAT[3,"N.SYNONYMOUS"],STAT[3,"N.NON.SYNONYMOUS"],STAT[3,"N.NON.SENSE"]),nrow=2,ncol=3,byrow=TRUE)
G.MATRIX.TAIL <- matrix(c(STAT[1,"N.SYNONYMOUS"],STAT[1,"N.NON.SYNONYMOUS"],STAT[1,"N.NON.SENSE"],STAT[4,"N.SYNONYMOUS"],STAT[4,"N.NON.SYNONYMOUS"],STAT[4,"N.NON.SENSE"]),nrow=2,ncol=3,byrow=TRUE)
G.MATRIX.CONSERVATIVE <- matrix(c(STAT[1,"N.SYNONYMOUS"],STAT[1,"N.NON.SYNONYMOUS"],STAT[1,"N.NON.SENSE"],STAT[5,"N.SYNONYMOUS"],STAT[5,"N.NON.SYNONYMOUS"],STAT[5,"N.NON.SENSE"]),nrow=2,ncol=3,byrow=TRUE)

STAT[,"G.STAT.TYPE"] <- c(likelihood.test(G.MATRIX.NEUTRAL,conservative=TRUE)$statistic,likelihood.test(G.MATRIX.CAUSAL,conservative=TRUE)$statistic,likelihood.test(G.MATRIX.RANDOM,conservative=TRUE)$statistic,likelihood.test(G.MATRIX.TAIL,conservative=TRUE)$statistic,likelihood.test(G.MATRIX.CONSERVATIVE,conservative=TRUE)$statistic)

STAT[,"P.VAL.TYPE"] <- c(pchisq(STAT[1,"G.STAT.TYPE"],df=2,lower.tail=FALSE),pchisq(STAT[2,"G.STAT.TYPE"],df=2,lower.tail=FALSE),pchisq(STAT[3,"G.STAT.TYPE"],df=2,lower.tail=FALSE),pchisq(STAT[4,"G.STAT.TYPE"],df=2,lower.tail=FALSE),pchisq(STAT[5,"G.STAT.TYPE"],df=2,lower.tail=FALSE))


STAT[,"FREQ.NON.SYN.NON.SENSE"] <- (STAT[,"N.NON.SYNONYMOUS"] + STAT[,"N.NON.SENSE"]) / (STAT[,"N.SYNONYMOUS"] + STAT[,"N.NON.SYNONYMOUS"] + STAT[,"N.NON.SENSE"] )

G.MATRIX.NEUTRAL <- matrix(c(STAT[1,"N.SYNONYMOUS"],STAT[1,"N.NON.SYNONYMOUS"]+STAT[1,"N.NON.SENSE"],STAT[1,"N.SYNONYMOUS"],STAT[1,"N.NON.SYNONYMOUS"]+STAT[1,"N.NON.SENSE"]),nrow=2,ncol=2,byrow=TRUE)
G.MATRIX.CAUSAL <- matrix(c(STAT[1,"N.SYNONYMOUS"],STAT[1,"N.NON.SYNONYMOUS"]+STAT[1,"N.NON.SENSE"],STAT[2,"N.SYNONYMOUS"],STAT[2,"N.NON.SYNONYMOUS"]+STAT[2,"N.NON.SENSE"]),nrow=2,ncol=2,byrow=TRUE)
G.MATRIX.RANDOM <- matrix(c(STAT[1,"N.SYNONYMOUS"],STAT[1,"N.NON.SYNONYMOUS"]+STAT[1,"N.NON.SENSE"],STAT[3,"N.SYNONYMOUS"],STAT[3,"N.NON.SYNONYMOUS"]+STAT[3,"N.NON.SENSE"]),nrow=2,ncol=2,byrow=TRUE)
G.MATRIX.TAIL <- matrix(c(STAT[1,"N.SYNONYMOUS"],STAT[1,"N.NON.SYNONYMOUS"]+STAT[1,"N.NON.SENSE"],STAT[4,"N.SYNONYMOUS"],STAT[4,"N.NON.SYNONYMOUS"]+STAT[4,"N.NON.SENSE"]),nrow=2,ncol=2,byrow=TRUE)
G.MATRIX.CONSERVATIVE <- matrix(c(STAT[1,"N.SYNONYMOUS"],STAT[1,"N.NON.SYNONYMOUS"]+STAT[1,"N.NON.SENSE"],STAT[5,"N.SYNONYMOUS"],STAT[5,"N.NON.SYNONYMOUS"]+STAT[5,"N.NON.SENSE"]),nrow=2,ncol=2,byrow=TRUE)

STAT[,"G.STAT.SYNONYMOUS"] <- c(likelihood.test(G.MATRIX.NEUTRAL,conservative=TRUE)$statistic,likelihood.test(G.MATRIX.CAUSAL,conservative=TRUE)$statistic,likelihood.test(G.MATRIX.RANDOM,conservative=TRUE)$statistic,likelihood.test(G.MATRIX.TAIL,conservative=TRUE)$statistic,likelihood.test(G.MATRIX.CONSERVATIVE,conservative=TRUE)$statistic)

STAT[,"P.VAL.SYNONYMOUS"] <- c(pchisq(STAT[1,"G.STAT.SYNONYMOUS"],df=2,lower.tail=FALSE),pchisq(STAT[2,"G.STAT.SYNONYMOUS"],df=2,lower.tail=FALSE),pchisq(STAT[3,"G.STAT.SYNONYMOUS"],df=2,lower.tail=FALSE),pchisq(STAT[4,"G.STAT.SYNONYMOUS"],df=2,lower.tail=FALSE),pchisq(STAT[5,"G.STAT.SYNONYMOUS"],df=2,lower.tail=FALSE))


STAT[,"FREQ.NON.SYNONYMOUS.2"] <- STAT[,"N.NON.SYNONYMOUS"]/ (STAT[,"N.NON.SYNONYMOUS"] + STAT[,"N.NON.SENSE"] )
STAT[,"FREQ.NON.SENSE.2"] <- (STAT[,"N.NON.SENSE"]) / (STAT[,"N.NON.SYNONYMOUS"] + STAT[,"N.NON.SENSE"] )

G.MATRIX.NEUTRAL <- matrix(c(STAT[1,"N.NON.SYNONYMOUS"],STAT[1,"N.NON.SENSE"],STAT[1,"N.NON.SYNONYMOUS"],STAT[1,"N.NON.SENSE"]),nrow=2,ncol=2,byrow=TRUE)
G.MATRIX.CAUSAL <- matrix(c(STAT[1,"N.NON.SYNONYMOUS"],STAT[1,"N.NON.SENSE"],STAT[2,"N.NON.SYNONYMOUS"],STAT[2,"N.NON.SENSE"]),nrow=2,ncol=2,byrow=TRUE)
G.MATRIX.RANDOM <- matrix(c(STAT[1,"N.NON.SYNONYMOUS"],STAT[1,"N.NON.SENSE"],STAT[3,"N.NON.SYNONYMOUS"],STAT[3,"N.NON.SENSE"]),nrow=2,ncol=2,byrow=TRUE)
G.MATRIX.TAIL <- matrix(c(STAT[1,"N.NON.SYNONYMOUS"],STAT[1,"N.NON.SENSE"],STAT[4,"N.NON.SYNONYMOUS"],STAT[4,"N.NON.SENSE"]),nrow=2,ncol=2,byrow=TRUE)
G.MATRIX.CONSERVATIVE <- matrix(c(STAT[1,"N.NON.SYNONYMOUS"],STAT[1,"N.NON.SENSE"],STAT[5,"N.NON.SYNONYMOUS"],STAT[5,"N.NON.SENSE"]),nrow=2,ncol=2,byrow=TRUE)

STAT[,"G.STAT.NON.SYN.NON.SENSE"] <- c(likelihood.test(G.MATRIX.NEUTRAL,conservative=TRUE)$statistic,likelihood.test(G.MATRIX.CAUSAL,conservative=TRUE)$statistic,likelihood.test(G.MATRIX.RANDOM,conservative=TRUE)$statistic,likelihood.test(G.MATRIX.TAIL,conservative=TRUE)$statistic,likelihood.test(G.MATRIX.CONSERVATIVE,conservative=TRUE)$statistic)

STAT[,"P.VAL.NON.SYN.NON.SENSE"] <- c(pchisq(STAT[1,"G.STAT.NON.SYN.NON.SENSE"],df=2,lower.tail=FALSE),pchisq(STAT[2,"G.STAT.NON.SYN.NON.SENSE"],df=2,lower.tail=FALSE),pchisq(STAT[3,"G.STAT.NON.SYN.NON.SENSE"],df=2,lower.tail=FALSE),pchisq(STAT[4,"G.STAT.NON.SYN.NON.SENSE"],df=2,lower.tail=FALSE),pchisq(STAT[5,"G.STAT.NON.SYN.NON.SENSE"],df=2,lower.tail=FALSE))

STAT[,"FREQ.NON.SYNONYMOUS.2"] <- STAT[,"N.NON.SYNONYMOUS"]/ (STAT[,"N.NON.SYNONYMOUS"] + STAT[,"N.NON.SENSE"] )
STAT[,"FREQ.NON.SENSE.2"] <- (STAT[,"N.NON.SENSE"]) / (STAT[,"N.NON.SYNONYMOUS"] + STAT[,"N.NON.SENSE"] )

G.MATRIX.NEUTRAL <- matrix(c(STAT[1,"N.NON.SYNONYMOUS"],STAT[1,"N.SYNONYMOUS"]+STAT[1,"N.NON.SENSE"],STAT[1,"N.NON.SYNONYMOUS"],STAT[1,"N.SYNONYMOUS"]+STAT[1,"N.NON.SENSE"]),nrow=2,ncol=2,byrow=TRUE)
G.MATRIX.CAUSAL <- matrix(c(STAT[1,"N.NON.SYNONYMOUS"],STAT[1,"N.SYNONYMOUS"]+STAT[1,"N.NON.SENSE"],STAT[2,"N.NON.SYNONYMOUS"],STAT[2,"N.SYNONYMOUS"]+STAT[2,"N.NON.SENSE"]),nrow=2,ncol=2,byrow=TRUE)
G.MATRIX.RANDOM <- matrix(c(STAT[1,"N.NON.SYNONYMOUS"],STAT[1,"N.SYNONYMOUS"]+STAT[1,"N.NON.SENSE"],STAT[3,"N.NON.SYNONYMOUS"],STAT[3,"N.SYNONYMOUS"]+STAT[3,"N.NON.SENSE"]),nrow=2,ncol=2,byrow=TRUE)
G.MATRIX.TAIL <- matrix(c(STAT[1,"N.NON.SYNONYMOUS"],STAT[1,"N.SYNONYMOUS"]+STAT[1,"N.NON.SENSE"],STAT[4,"N.NON.SYNONYMOUS"],STAT[4,"N.SYNONYMOUS"]+STAT[4,"N.NON.SENSE"]),nrow=2,ncol=2,byrow=TRUE)
G.MATRIX.CONSERVATIVE <- matrix(c(STAT[1,"N.NON.SYNONYMOUS"],STAT[1,"N.SYNONYMOUS"]+STAT[1,"N.NON.SENSE"],STAT[5,"N.NON.SYNONYMOUS"],STAT[5,"N.SYNONYMOUS"]+STAT[5,"N.NON.SENSE"]),nrow=2,ncol=2,byrow=TRUE)

STAT[,"G.STAT.NON.SYNONYMOUS"] <- c(likelihood.test(G.MATRIX.NEUTRAL,conservative=TRUE)$statistic,likelihood.test(G.MATRIX.CAUSAL,conservative=TRUE)$statistic,likelihood.test(G.MATRIX.RANDOM,conservative=TRUE)$statistic,likelihood.test(G.MATRIX.TAIL,conservative=TRUE)$statistic,likelihood.test(G.MATRIX.CONSERVATIVE,conservative=TRUE)$statistic)

STAT[,"P.VAL.NON.SYNONYMOUS"] <- c(pchisq(STAT[1,"G.STAT.NON.SYNONYMOUS"],df=2,lower.tail=FALSE),pchisq(STAT[2,"G.STAT.NON.SYNONYMOUS"],df=2,lower.tail=FALSE),pchisq(STAT[3,"G.STAT.NON.SYNONYMOUS"],df=2,lower.tail=FALSE),pchisq(STAT[4,"G.STAT.NON.SYNONYMOUS"],df=2,lower.tail=FALSE),pchisq(STAT[5,"G.STAT.NON.SYNONYMOUS"],df=2,lower.tail=FALSE))


G.MATRIX.NEUTRAL <- matrix(c(STAT[1,"N.NON.SENSE"],STAT[1,"N.NON.SYNONYMOUS"]+STAT[1,"N.SYNONYMOUS"],STAT[1,"N.NON.SENSE"],STAT[1,"N.NON.SYNONYMOUS"]+STAT[1,"N.SYNONYMOUS"]),nrow=2,ncol=2,byrow=TRUE)
G.MATRIX.CAUSAL <- matrix(c(STAT[1,"N.NON.SENSE"],STAT[1,"N.NON.SYNONYMOUS"]+STAT[1,"N.SYNONYMOUS"],STAT[2,"N.NON.SENSE"],STAT[2,"N.NON.SYNONYMOUS"]+STAT[2,"N.SYNONYMOUS"]),nrow=2,ncol=2,byrow=TRUE)
G.MATRIX.RANDOM <- matrix(c(STAT[1,"N.NON.SENSE"],STAT[1,"N.NON.SYNONYMOUS"]+STAT[1,"N.SYNONYMOUS"],STAT[3,"N.NON.SENSE"],STAT[3,"N.NON.SYNONYMOUS"]+STAT[3,"N.SYNONYMOUS"]),nrow=2,ncol=2,byrow=TRUE)
G.MATRIX.TAIL <- matrix(c(STAT[1,"N.NON.SENSE"],STAT[1,"N.NON.SYNONYMOUS"]+STAT[1,"N.SYNONYMOUS"],STAT[4,"N.NON.SENSE"],STAT[4,"N.NON.SYNONYMOUS"]+STAT[4,"N.SYNONYMOUS"]),nrow=2,ncol=2,byrow=TRUE)
G.MATRIX.CONSERVATIVE <- matrix(c(STAT[1,"N.NON.SENSE"],STAT[1,"N.NON.SYNONYMOUS"]+STAT[1,"N.SYNONYMOUS"],STAT[5,"N.NON.SENSE"],STAT[5,"N.NON.SYNONYMOUS"]+STAT[5,"N.SYNONYMOUS"]),nrow=2,ncol=2,byrow=TRUE)

STAT[,"G.STAT.NON.SENSE"] <- c(likelihood.test(G.MATRIX.NEUTRAL,conservative=TRUE)$statistic,likelihood.test(G.MATRIX.CAUSAL,conservative=TRUE)$statistic,likelihood.test(G.MATRIX.RANDOM,conservative=TRUE)$statistic,likelihood.test(G.MATRIX.TAIL,conservative=TRUE)$statistic,likelihood.test(G.MATRIX.CONSERVATIVE,conservative=TRUE)$statistic)

STAT[,"P.VAL.NON.SENSE"] <- c(pchisq(STAT[1,"G.STAT.NON.SENSE"],df=2,lower.tail=FALSE),pchisq(STAT[2,"G.STAT.NON.SENSE"],df=2,lower.tail=FALSE),pchisq(STAT[3,"G.STAT.NON.SENSE"],df=2,lower.tail=FALSE),pchisq(STAT[4,"G.STAT.NON.SENSE"],df=2,lower.tail=FALSE),pchisq(STAT[5,"G.STAT.NON.SENSE"],df=2,lower.tail=FALSE))


STAT[,"FREQ.NON.SYN.NON.SENSE"] <- (STAT[,"N.NON.SYNONYMOUS"] + STAT[,"N.NON.SENSE"]) / (STAT[,"N.SYNONYMOUS"] + STAT[,"N.NON.SYNONYMOUS"] + STAT[,"N.NON.SENSE"] )

G.MATRIX.NEUTRAL <- matrix(c(STAT[1,"N.SYNONYMOUS"],STAT[1,"N.NON.SYNONYMOUS"]+STAT[1,"N.NON.SENSE"],STAT[1,"N.SYNONYMOUS"],STAT[1,"N.NON.SYNONYMOUS"]+STAT[1,"N.NON.SENSE"]),nrow=2,ncol=2,byrow=TRUE)
G.MATRIX.CAUSAL <- matrix(c(STAT[1,"N.SYNONYMOUS"],STAT[1,"N.NON.SYNONYMOUS"]+STAT[1,"N.NON.SENSE"],STAT[2,"N.SYNONYMOUS"],STAT[2,"N.NON.SYNONYMOUS"]+STAT[2,"N.NON.SENSE"]),nrow=2,ncol=2,byrow=TRUE)
G.MATRIX.RANDOM <- matrix(c(STAT[1,"N.SYNONYMOUS"],STAT[1,"N.NON.SYNONYMOUS"]+STAT[1,"N.NON.SENSE"],STAT[3,"N.SYNONYMOUS"],STAT[3,"N.NON.SYNONYMOUS"]+STAT[3,"N.NON.SENSE"]),nrow=2,ncol=2,byrow=TRUE)
G.MATRIX.TAIL <- matrix(c(STAT[1,"N.SYNONYMOUS"],STAT[1,"N.NON.SYNONYMOUS"]+STAT[1,"N.NON.SENSE"],STAT[4,"N.SYNONYMOUS"],STAT[4,"N.NON.SYNONYMOUS"]+STAT[4,"N.NON.SENSE"]),nrow=2,ncol=2,byrow=TRUE)
G.MATRIX.CONSERVATIVE <- matrix(c(STAT[1,"N.SYNONYMOUS"],STAT[1,"N.NON.SYNONYMOUS"]+STAT[1,"N.NON.SENSE"],STAT[5,"N.SYNONYMOUS"],STAT[5,"N.NON.SYNONYMOUS"]+STAT[5,"N.NON.SENSE"]),nrow=2,ncol=2,byrow=TRUE)

STAT[,"G.STAT.SYNONYMOUS"] <- c(likelihood.test(G.MATRIX.NEUTRAL,conservative=TRUE)$statistic,likelihood.test(G.MATRIX.CAUSAL,conservative=TRUE)$statistic,likelihood.test(G.MATRIX.RANDOM,conservative=TRUE)$statistic,likelihood.test(G.MATRIX.TAIL,conservative=TRUE)$statistic,likelihood.test(G.MATRIX.CONSERVATIVE,conservative=TRUE)$statistic)

STAT[,"P.VAL.SYNONYMOUS"] <- c(pchisq(STAT[1,"G.STAT.SYNONYMOUS"],df=2,lower.tail=FALSE),pchisq(STAT[2,"G.STAT.SYNONYMOUS"],df=2,lower.tail=FALSE),pchisq(STAT[3,"G.STAT.SYNONYMOUS"],df=2,lower.tail=FALSE),pchisq(STAT[4,"G.STAT.SYNONYMOUS"],df=2,lower.tail=FALSE),pchisq(STAT[5,"G.STAT.SYNONYMOUS"],df=2,lower.tail=FALSE))

#write.table(STAT,"Statistics/STAT.MUTATION.TYPE.txt",sep="\t",row.names=FALSE)


#Pie charts

pdf("Figure 3F Non-regulatory.pdf",width=6,height=6,useDingbats = F)
#windows(width=6,height=6)
pie(c(STAT[1,"N.SYNONYMOUS"],STAT[1,"N.NON.SYNONYMOUS"],STAT[1,"N.NON.SENSE"]), labels = c("Synonymous","Non-synonymous","Non-sense"), border="white", col=c("black","blue","gray"),init.angle=90)
dev.off()

pdf("Figure 3F Trans-regulatory.pdf",width=6,height=6,useDingbats = F)
#windows(width=6,height=6)
pie(c(STAT[2,"N.SYNONYMOUS"],STAT[2,"N.NON.SYNONYMOUS"],STAT[2,"N.NON.SENSE"]), labels = c("Synonymous","Non-synonymous","Non-sense"), border="white", col=c("black","blue","gray"),init.angle=90)
dev.off()


##########################################################################################################################################
#M) Frequencies of amino acid changes caused by non-regulatory and trans-regulatory mutations (Figure 3E, Figure 3- figure supplement 2).#
##########################################################################################################################################

###1) All non-regulatory substitutions.###

AMINO.ACIDS <- c("Ala","Arg","Asn","Asp","Cys","Gln","Glu","Gly","His","Ile","Leu","Lys","Met","Phe","Pro","Ser","Thr","Trp","Tyr","Val","Stop")

#Create tables with the number and frequency of amino acid changes induced by non-regulatory mutations in coding sequences.
N.SUBSTITUTION <- matrix(0,nrow=22,ncol=22)
colnames(N.SUBSTITUTION) <- c(AMINO.ACIDS,"REF")
rownames(N.SUBSTITUTION) <- c(AMINO.ACIDS,"ALT")

for (i in 1:21)
{
  for (j in 1:21)
  {
    N.SUBSTITUTION[i,j] <- length(which(SUBSTITUTION[,"REF.AA"] == AMINO.ACIDS[i] & SUBSTITUTION[,"ALT.AA"] == AMINO.ACIDS[j]))
  }
}

N.SUBSTITUTION["ALT",] <- colSums(N.SUBSTITUTION)
N.SUBSTITUTION[,"REF"] <- rowSums(N.SUBSTITUTION)

FREQ.SUBSTITUTION <- round(100*N.SUBSTITUTION/N.SUBSTITUTION[22,22],1)

#The two tables can be found in SupplementaryFile11.zip.
write.table(N.SUBSTITUTION,"Amino.Acids/N.SUBSTITUTION.txt",sep="\t",row.names= TRUE)
write.table(FREQ.SUBSTITUTION,"Amino.Acids/FREQ.SUBSTITUTION.txt",sep="\t",row.names= TRUE)


###2) All causal substitutions.###

AMINO.ACIDS <- c("Ala","Arg","Asn","Asp","Cys","Gln","Glu","Gly","His","Ile","Leu","Lys","Met","Phe","Pro","Ser","Thr","Trp","Tyr","Val","Stop")

#Create tables with the number and frequency of amino acid changes induced by trans-regulatory mutations in coding sequences
#and one table with the enrichment of amino acids changes induced by trans-regulatory mutation relative to non-regulatory mutations.
N.CAUSAL <- matrix(0,nrow=22,ncol=22)
colnames(N.CAUSAL) <- c(AMINO.ACIDS,"REF")
rownames(N.CAUSAL) <- c(AMINO.ACIDS,"ALT")

for (i in 1:21)
{
  for (j in 1:21)
  {
    N.CAUSAL[i,j] <- length(which(CAUSAL[,"REF.AA"] == AMINO.ACIDS[i] & CAUSAL[,"ALT.AA"] == AMINO.ACIDS[j]))
  }
}

N.CAUSAL["ALT",] <- colSums(N.CAUSAL)
N.CAUSAL[,"REF"] <- rowSums(N.CAUSAL)

FREQ.CAUSAL <- round(100*N.CAUSAL/N.CAUSAL[22,22],1)
ENRICHMENT.CAUSAL <- round(log2(FREQ.CAUSAL/FREQ.SUBSTITUTION),2)
DIFF.CAUSAL <- FREQ.CAUSAL - FREQ.SUBSTITUTION

#Permutation tests to determine statistical significance of amino acid changes induced by trans-regulatory mutations.
N.REP <- 10000
NB.CAUSAL <- nrow(CAUSAL)
NB.SUBSTITUTION <- nrow(SUBSTITUTION)

COMBINED <- rbind(CAUSAL,SUBSTITUTION)

N.SAMPLE.CAUSAL <- matrix(0,nrow=22,ncol=22)
colnames(N.SAMPLE.CAUSAL) <- c(AMINO.ACIDS,"REF")
rownames(N.SAMPLE.CAUSAL) <- c(AMINO.ACIDS,"ALT")

N.SAMPLE.SUBSTITUTION <- matrix(0,nrow=22,ncol=22)
colnames(N.SAMPLE.SUBSTITUTION) <- c(AMINO.ACIDS,"REF")
rownames(N.SAMPLE.SUBSTITUTION) <- c(AMINO.ACIDS,"ALT")

COUNTS <- matrix(0,nrow=22,ncol=22)
colnames(COUNTS) <- c(AMINO.ACIDS,"REF")
rownames(COUNTS) <- c(AMINO.ACIDS,"ALT")

for (n in 1:N.REP)
{
  SHUFFLE <- COMBINED[sample(1:nrow(COMBINED),nrow(COMBINED)),]
  SAMPLE.CAUSAL <- SHUFFLE[1:NB.CAUSAL,]
  SAMPLE.SUBSTITUTION <- SHUFFLE[(1+NB.CAUSAL):nrow(SHUFFLE),]
  
  for (i in 1:21)
  {
    for (j in 1:21)
    {
      N.SAMPLE.CAUSAL[i,j] <- length(which(SAMPLE.CAUSAL[,"REF.AA"] == AMINO.ACIDS[i] & SAMPLE.CAUSAL[,"ALT.AA"] == AMINO.ACIDS[j]))
      N.SAMPLE.SUBSTITUTION[i,j] <- length(which(SAMPLE.SUBSTITUTION[,"REF.AA"] == AMINO.ACIDS[i] & SAMPLE.SUBSTITUTION[,"ALT.AA"] == AMINO.ACIDS[j]))
    }
  }
  
  N.SAMPLE.CAUSAL["ALT",] <- colSums(N.SAMPLE.CAUSAL[1:21,])
  N.SAMPLE.CAUSAL[,"REF"] <- rowSums(N.SAMPLE.CAUSAL[,1:21])
  N.SAMPLE.SUBSTITUTION["ALT",] <- colSums(N.SAMPLE.SUBSTITUTION[1:21,])
  N.SAMPLE.SUBSTITUTION[,"REF"] <- rowSums(N.SAMPLE.SUBSTITUTION[,1:21])
  
  FREQ.SAMPLE.CAUSAL <- round(100*N.SAMPLE.CAUSAL/N.SAMPLE.CAUSAL[22,22],1)
  FREQ.SAMPLE.SUBSTITUTION	<- round(100*N.SAMPLE.SUBSTITUTION/N.SAMPLE.SUBSTITUTION[22,22],1)
  DIFF.SAMPLE.CAUSAL <- FREQ.SAMPLE.CAUSAL - FREQ.SAMPLE.SUBSTITUTION
  
  TEST <- (abs(DIFF.SAMPLE.CAUSAL) >= abs(DIFF.CAUSAL))*1
  COUNTS <- COUNTS + TEST
}

P.VAL.CAUSAL <- ((COUNTS)/(N.REP))
LOG.P.VAL.CAUSAL <- round(-log10(((COUNTS+1)/(N.REP+1))),2)

#These tables can be found in SupplementaryFile11.zip.
write.table(N.CAUSAL,"Amino.Acids/N.CAUSAL.txt",sep="\t",row.names= TRUE)
write.table(FREQ.CAUSAL,"Amino.Acids/FREQ.CAUSAL.txt",sep="\t",row.names= TRUE)
write.table(ENRICHMENT.CAUSAL,"Amino.Acids/ENRICHMENT.CAUSAL.txt",sep="\t",row.names= TRUE)
write.table(DIFF.CAUSAL,"Amino.Acids/DIFF.CAUSAL.txt",sep="\t",row.names= TRUE) #Data used to plot Figure 3E.
write.table(P.VAL.CAUSAL,"Amino.Acids/P.VAL.CAUSAL.txt",sep="\t",row.names= TRUE)
write.table(LOG.P.VAL.CAUSAL,"Amino.Acids/LOG.P.VAL.CAUSAL.txt",sep="\t",row.names= TRUE) #Data used to plot Figure 3- figure supplement 2.


###3) Plot difference of frequency of amino acid changes induced by trans-regulatory mutations and non-regulatory mutations (Figure 3E).###

rotate <- function(x) t(apply(x,2,rev))

PLOT.CAUSAL <- rotate(round(DIFF.CAUSAL,0))

N.COL.POSITIVE <- 1000
MIN.CAUSAL <- min(PLOT.CAUSAL)
MAX.CAUSAL <- max(PLOT.CAUSAL)
N.COL.NEGATIVE <- abs(round(N.COL.POSITIVE*MIN.CAUSAL/MAX.CAUSAL))

COL.POSITIVE <- colorRampPalette(c("#DFFFDF","#009B00")) (N.COL.POSITIVE)[-1]
COL.NEGATIVE <- colorRampPalette(c("red","white")) (N.COL.NEGATIVE)
COL.RAMP <- c(COL.NEGATIVE,COL.POSITIVE)

pdf("Figure 3E.pdf",useDingbats=FALSE,height=10,width=14)

#windows(height=10,width=14)
par(mar=c(6,7,5,6))
image(PLOT.CAUSAL,col=COL.RAMP,xaxt="n",yaxt="n",lwd=2,ylab="From",xlab="To",cex.lab=1.5,font.lab=2,main="Frequency of amino acid change (%) for all causal mutations - neutral mutations",cex.main=1.5)
box(lwd=2)

BOTTOM <- 1/nrow(DIFF.CAUSAL)
abline(h=BOTTOM-BOTTOM/2,lty=2,lwd=2)

RIGHT <- 1/ncol(DIFF.CAUSAL)
abline(v=1-(RIGHT-RIGHT/2),lty=2,lwd=2)

#Add data values to the plot
X.POS <- (0:(ncol(DIFF.CAUSAL)-1))/(ncol(DIFF.CAUSAL)-1)
Y.POS <- (0:(nrow(DIFF.CAUSAL)-1))/(nrow(DIFF.CAUSAL)-1)

X.VAL <- rep(X.POS,nrow(DIFF.CAUSAL))
Y.VAL <- rep(Y.POS,each=ncol(DIFF.CAUSAL))

LABELS <- round(DIFF.CAUSAL,0) 

text(X.VAL,Y.VAL,labels=PLOT.CAUSAL)
axis(1,at=X.POS,labels=colnames(DIFF.CAUSAL))
axis(2,at=1-Y.POS,labels=rownames(DIFF.CAUSAL),las=2)
axis(4,at=1-Y.POS,labels=rownames(DIFF.CAUSAL),las=2)

dev.off()


###4) Plot P-values of statistical tests used to compare amino acid changes induced by trans-regulatory mutations and non-regulatory mutations (Figure 3 - figure supplement 2).###

rotate <- function(x) t(apply(x,2,rev))

PLOT.CAUSAL <- rotate(round(LOG.P.VAL.CAUSAL,2))

N.COL.POSITIVE <- 1000
MIN.CAUSAL <- min(PLOT.CAUSAL)
MAX.CAUSAL <- max(PLOT.CAUSAL)
N.COL.NEGATIVE <- abs(round(N.COL.POSITIVE*MIN.CAUSAL/MAX.CAUSAL))

COL.POSITIVE <- colorRampPalette(c("white","#009B00")) (N.COL.POSITIVE)[-1]
COL.NEGATIVE <- colorRampPalette(c("red","white")) (N.COL.NEGATIVE)
COL.RAMP <- c(COL.NEGATIVE,COL.POSITIVE)

pdf("Figure 3 - figure supplement 2.pdf",useDingbats=FALSE,height=10,width=14)

#windows(height=10,width=14)
par(mar=c(6,7,5,6))
image(PLOT.CAUSAL,col=COL.RAMP,xaxt="n",yaxt="n",lwd=2,ylab="From",xlab="To",cex.lab=1.5,font.lab=2,main="-log10(P-value) of permutations tests comparing frequencies of amino acid change for all trans-regulatory mutations vs non-regulatory mutations",cex.main=1.5)
box(lwd=2)

BOTTOM <- 1/nrow(LOG.P.VAL.CAUSAL)
abline(h=BOTTOM-BOTTOM/2,lty=2,lwd=2)

RIGHT <- 1/ncol(LOG.P.VAL.CAUSAL)
abline(v=1-(RIGHT-RIGHT/2),lty=2,lwd=2)

#Add data values to the plot
X.POS <- (0:(ncol(LOG.P.VAL.CAUSAL)-1))/(ncol(LOG.P.VAL.CAUSAL)-1)
Y.POS <- (0:(nrow(LOG.P.VAL.CAUSAL)-1))/(nrow(LOG.P.VAL.CAUSAL)-1)

X.VAL <- rep(X.POS,nrow(LOG.P.VAL.CAUSAL))
Y.VAL <- rep(Y.POS,each=ncol(LOG.P.VAL.CAUSAL))

LABELS <- round(LOG.P.VAL.CAUSAL,2) 

text(X.VAL,Y.VAL,labels=PLOT.CAUSAL)
axis(1,at=X.POS,labels=colnames(LOG.P.VAL.CAUSAL))
axis(2,at=1-Y.POS,labels=rownames(LOG.P.VAL.CAUSAL),las=2)
axis(4,at=1-Y.POS,labels=rownames(LOG.P.VAL.CAUSAL),las=2)

dev.off()


##################################################################################################
#N) Testing for enrichment of trans-regulatory mutations in genes encoding transcription factors.#
##################################################################################################

#The list of yeast genes considered to encode transcription factors was obtained from www.yeastract.com.
#This list can be found in SupplementaryFile11.zip.
ALL.TF <- read.table("Network/TF_Names.txt",header=TRUE)

ALL.TF[,"GENE"] <- as.character(ALL.TF[,"GENE"])

for (i in 1:nrow(ALL.TF))
{
  if (is.na(ALL.TF[i,"GENE"]))
  {
    GOOD.GENE <- subset(ALL.GENES, grepl(ALL.TF[i,"GENE.SHORT"],SHORT))
    
    if (nrow(GOOD.GENE) == 1)
    {
      ALL.TF[i,"GENE"] <- as.character(GOOD.GENE[1,"GENE"])
    }
  }
}

#write.table(ALL.TF,"Network/TF_Names_Fix.txt",sep="\t",row.names= TRUE)

#G-tests used to compare the frequency of trans-regulatory mutations and non-regulatory mutations located in transcription factor genes.
N.TF <- c(nrow(subset(NEUTRAL, CLASS == "EXONIC")),nrow(subset(CAUSAL, CLASS == "EXONIC")),nrow(subset(RANDOM, CLASS == "EXONIC")),nrow(subset(TAIL, CLASS == "EXONIC")),nrow(subset(CONSERVATIVE, CLASS == "EXONIC")))
N.NOT.TF <- c(nrow(subset(NEUTRAL, CLASS == "INTRONIC")),nrow(subset(CAUSAL, CLASS == "INTRONIC")),nrow(subset(RANDOM, CLASS == "INTRONIC")),nrow(subset(TAIL, CLASS == "INTRONIC")),nrow(subset(CONSERVATIVE, CLASS == "INTRONIC")))

N.TF.CODING <- c(nrow(subset(NEUTRAL, GENE.FOCAL %in% as.character(ALL.TF[,"GENE"]))),nrow(subset(CAUSAL, GENE.FOCAL %in% as.character(ALL.TF[,"GENE"]))),nrow(subset(RANDOM, GENE.FOCAL %in% as.character(ALL.TF[,"GENE"]))),nrow(subset(TAIL, GENE.FOCAL %in% as.character(ALL.TF[,"GENE"]))),nrow(subset(CONSERVATIVE, GENE.FOCAL %in% as.character(ALL.TF[,"GENE"]))))
N.NOT.TF.CODING <- c(nrow(subset(NEUTRAL, CLASS == "EXONIC" & GENE.FOCAL %nin% as.character(ALL.TF[,"GENE"]))),nrow(subset(CAUSAL, CLASS == "EXONIC" & GENE.FOCAL %nin% as.character(ALL.TF[,"GENE"]))),nrow(subset(RANDOM, CLASS == "EXONIC" & GENE.FOCAL %nin% as.character(ALL.TF[,"GENE"]))),nrow(subset(TAIL, CLASS == "EXONIC" & GENE.FOCAL %nin% as.character(ALL.TF[,"GENE"]))),nrow(subset(CONSERVATIVE, CLASS == "EXONIC" & GENE.FOCAL %nin% as.character(ALL.TF[,"GENE"]))))
N.TF.PROMOTER <- c(nrow(subset(NEUTRAL, (GENE.UPSTREAM %in% as.character(ALL.TF[,"GENE"]) & LOCATION.UPSTREAM == "PROMOTER") | (GENE.DOWNSTREAM %in% as.character(ALL.TF[,"GENE"]) & LOCATION.DOWNSTREAM == "PROMOTER"))),nrow(subset(CAUSAL, (GENE.UPSTREAM %in% as.character(ALL.TF[,"GENE"]) & LOCATION.UPSTREAM == "PROMOTER") | (GENE.DOWNSTREAM %in% as.character(ALL.TF[,"GENE"]) & LOCATION.DOWNSTREAM == "PROMOTER"))),nrow(subset(RANDOM, (GENE.UPSTREAM %in% as.character(ALL.TF[,"GENE"]) & LOCATION.UPSTREAM == "PROMOTER") | (GENE.DOWNSTREAM %in% as.character(ALL.TF[,"GENE"]) & LOCATION.DOWNSTREAM == "PROMOTER"))),nrow(subset(TAIL, (GENE.UPSTREAM %in% as.character(ALL.TF[,"GENE"]) & LOCATION.UPSTREAM == "PROMOTER") | (GENE.DOWNSTREAM %in% as.character(ALL.TF[,"GENE"]) & LOCATION.DOWNSTREAM == "PROMOTER"))),nrow(subset(CONSERVATIVE, (GENE.UPSTREAM %in% as.character(ALL.TF[,"GENE"]) & LOCATION.UPSTREAM == "PROMOTER") | (GENE.DOWNSTREAM %in% as.character(ALL.TF[,"GENE"]) & LOCATION.DOWNSTREAM == "PROMOTER"))))
N.NOT.TF.PROMOTER <- c(nrow(subset(NEUTRAL, (GENE.UPSTREAM %nin% as.character(ALL.TF[,"GENE"]) & LOCATION.UPSTREAM == "PROMOTER") | (GENE.DOWNSTREAM %nin% as.character(ALL.TF[,"GENE"]) & LOCATION.DOWNSTREAM == "PROMOTER"))),nrow(subset(CAUSAL, (GENE.UPSTREAM %nin% as.character(ALL.TF[,"GENE"]) & LOCATION.UPSTREAM == "PROMOTER") | (GENE.DOWNSTREAM %nin% as.character(ALL.TF[,"GENE"]) & LOCATION.DOWNSTREAM == "PROMOTER"))),nrow(subset(RANDOM, (GENE.UPSTREAM %nin% as.character(ALL.TF[,"GENE"]) & LOCATION.UPSTREAM == "PROMOTER") | (GENE.DOWNSTREAM %nin% as.character(ALL.TF[,"GENE"]) & LOCATION.DOWNSTREAM == "PROMOTER"))),nrow(subset(TAIL, (GENE.UPSTREAM %nin% as.character(ALL.TF[,"GENE"]) & LOCATION.UPSTREAM == "PROMOTER") | (GENE.DOWNSTREAM %nin% as.character(ALL.TF[,"GENE"]) & LOCATION.DOWNSTREAM == "PROMOTER"))),nrow(subset(CONSERVATIVE, (GENE.UPSTREAM %nin% as.character(ALL.TF[,"GENE"]) & LOCATION.UPSTREAM == "PROMOTER") | (GENE.DOWNSTREAM %nin% as.character(ALL.TF[,"GENE"]) & LOCATION.DOWNSTREAM == "PROMOTER"))))

STAT <- data.frame(MUT.TYPE, N.MUT, N.TF.CODING, N.NOT.TF.CODING,N.TF.PROMOTER,N.NOT.TF.PROMOTER)

STAT[,"FREQ.TF.CODING.vs.NOT.TF.CODING"] <- STAT[,"N.TF.CODING"]/(STAT[,"N.TF.CODING"] + STAT[,"N.NOT.TF.CODING"])
STAT[,"FREQ.TF.CODING.vs.NOT.TF.ALL"] <- STAT[,"N.TF.CODING"]/(STAT[,"N.MUT"])
STAT[,"FREQ.TF.vs.NOT.TF.ALL"] <- (STAT[,"N.TF.CODING"] + STAT[,"N.TF.PROMOTER"])/(STAT[,"N.MUT"])

G.MATRIX.NEUTRAL <- matrix(c(STAT[1,"N.TF.CODING"],STAT[1,"N.NOT.TF.CODING"],STAT[1,"N.TF.CODING"],STAT[1,"N.NOT.TF.CODING"]),nrow=2,ncol=2)
G.MATRIX.CAUSAL <- matrix(c(STAT[1,"N.TF.CODING"],STAT[1,"N.NOT.TF.CODING"],STAT[2,"N.TF.CODING"],STAT[2,"N.NOT.TF.CODING"]),nrow=2,ncol=2)
G.MATRIX.RANDOM <- matrix(c(STAT[1,"N.TF.CODING"],STAT[1,"N.NOT.TF.CODING"],STAT[3,"N.TF.CODING"],STAT[3,"N.NOT.TF.CODING"]),nrow=2,ncol=2)
G.MATRIX.TAIL <- matrix(c(STAT[1,"N.TF.CODING"],STAT[1,"N.NOT.TF.CODING"],STAT[4,"N.TF.CODING"],STAT[4,"N.NOT.TF.CODING"]),nrow=2,ncol=2)
G.MATRIX.CONSERVATIVE <- matrix(c(STAT[1,"N.TF.CODING"],STAT[1,"N.NOT.TF.CODING"],STAT[5,"N.TF.CODING"],STAT[5,"N.NOT.TF.CODING"]),nrow=2,ncol=2)

STAT[,"G.STAT.TF.CODING.vs.NOT.TF.CODING"] <- c(likelihood.test(G.MATRIX.NEUTRAL,conservative=TRUE)$statistic,likelihood.test(G.MATRIX.CAUSAL,conservative=TRUE)$statistic,likelihood.test(G.MATRIX.RANDOM,conservative=TRUE)$statistic,likelihood.test(G.MATRIX.TAIL,conservative=TRUE)$statistic,likelihood.test(G.MATRIX.CONSERVATIVE,conservative=TRUE)$statistic)

STAT[,"P.VAL.TF.CODING.vs.NOT.TF.CODING"] <- c(pchisq(STAT[1,"G.STAT.TF.CODING.vs.NOT.TF.CODING"],df=1,lower.tail=FALSE),pchisq(STAT[2,"G.STAT.TF.CODING.vs.NOT.TF.CODING"],df=1,lower.tail=FALSE),pchisq(STAT[3,"G.STAT.TF.CODING.vs.NOT.TF.CODING"],df=1,lower.tail=FALSE),pchisq(STAT[4,"G.STAT.TF.CODING.vs.NOT.TF.CODING"],df=1,lower.tail=FALSE),pchisq(STAT[5,"G.STAT.TF.CODING.vs.NOT.TF.CODING"],df=1,lower.tail=FALSE))

G.MATRIX.NEUTRAL <- matrix(c(STAT[1,"N.TF.CODING"],STAT[1,"N.MUT"] - STAT[1,"N.TF.CODING"],STAT[1,"N.TF.CODING"],STAT[1,"N.MUT"] - STAT[1,"N.TF.CODING"]),nrow=2,ncol=2)
G.MATRIX.CAUSAL <- matrix(c(STAT[1,"N.TF.CODING"],STAT[1,"N.MUT"] - STAT[1,"N.TF.CODING"],STAT[2,"N.TF.CODING"],STAT[2,"N.MUT"] - STAT[2,"N.TF.CODING"]),nrow=2,ncol=2)
G.MATRIX.RANDOM <- matrix(c(STAT[1,"N.TF.CODING"],STAT[1,"N.MUT"] - STAT[1,"N.TF.CODING"],STAT[3,"N.TF.CODING"],STAT[3,"N.MUT"] - STAT[3,"N.TF.CODING"]),nrow=2,ncol=2)
G.MATRIX.TAIL <- matrix(c(STAT[1,"N.TF.CODING"],STAT[1,"N.MUT"] - STAT[1,"N.TF.CODING"],STAT[4,"N.TF.CODING"],STAT[4,"N.MUT"] - STAT[4,"N.TF.CODING"]),nrow=2,ncol=2)
G.MATRIX.CONSERVATIVE <- matrix(c(STAT[1,"N.TF.CODING"],STAT[1,"N.MUT"] - STAT[1,"N.TF.CODING"],STAT[5,"N.TF.CODING"],STAT[5,"N.MUT"] - STAT[5,"N.TF.CODING"]),nrow=2,ncol=2)

STAT[,"G.STAT.TF.CODING.vs.NOT.TF.ALL"] <- c(likelihood.test(G.MATRIX.NEUTRAL,conservative=TRUE)$statistic,likelihood.test(G.MATRIX.CAUSAL,conservative=TRUE)$statistic,likelihood.test(G.MATRIX.RANDOM,conservative=TRUE)$statistic,likelihood.test(G.MATRIX.TAIL,conservative=TRUE)$statistic,likelihood.test(G.MATRIX.CONSERVATIVE,conservative=TRUE)$statistic)

STAT[,"P.VAL.TF.CODING.vs.NOT.TF.ALL"] <- c(pchisq(STAT[1,"G.STAT.TF.CODING.vs.NOT.TF.ALL"],df=1,lower.tail=FALSE),pchisq(STAT[2,"G.STAT.TF.CODING.vs.NOT.TF.ALL"],df=1,lower.tail=FALSE),pchisq(STAT[3,"G.STAT.TF.CODING.vs.NOT.TF.ALL"],df=1,lower.tail=FALSE),pchisq(STAT[4,"G.STAT.TF.CODING.vs.NOT.TF.ALL"],df=1,lower.tail=FALSE),pchisq(STAT[5,"G.STAT.TF.CODING.vs.NOT.TF.ALL"],df=1,lower.tail=FALSE))

G.MATRIX.NEUTRAL <- matrix(c(STAT[1,"N.TF.CODING"] + STAT[1,"N.TF.PROMOTER"],STAT[1,"N.MUT"] - STAT[1,"N.TF.CODING"] - STAT[1,"N.TF.PROMOTER"],STAT[1,"N.TF.CODING"] + STAT[1,"N.TF.PROMOTER"],STAT[1,"N.MUT"] - STAT[1,"N.TF.CODING"] - STAT[1,"N.TF.PROMOTER"]),nrow=2,ncol=2)
G.MATRIX.CAUSAL <- matrix(c(STAT[1,"N.TF.CODING"] + STAT[1,"N.TF.PROMOTER"],STAT[1,"N.MUT"] - STAT[1,"N.TF.CODING"] - STAT[1,"N.TF.PROMOTER"],STAT[2,"N.TF.CODING"] + STAT[2,"N.TF.PROMOTER"],STAT[2,"N.MUT"] - STAT[2,"N.TF.CODING"] - STAT[2,"N.TF.PROMOTER"]),nrow=2,ncol=2)
G.MATRIX.RANDOM <- matrix(c(STAT[1,"N.TF.CODING"] + STAT[1,"N.TF.PROMOTER"],STAT[1,"N.MUT"] - STAT[1,"N.TF.CODING"] - STAT[1,"N.TF.PROMOTER"],STAT[3,"N.TF.CODING"] + STAT[3,"N.TF.PROMOTER"],STAT[3,"N.MUT"] - STAT[3,"N.TF.CODING"] - STAT[3,"N.TF.PROMOTER"]),nrow=2,ncol=2)
G.MATRIX.TAIL <- matrix(c(STAT[1,"N.TF.CODING"] + STAT[1,"N.TF.PROMOTER"],STAT[1,"N.MUT"] - STAT[1,"N.TF.CODING"] - STAT[1,"N.TF.PROMOTER"],STAT[4,"N.TF.CODING"] + STAT[4,"N.TF.PROMOTER"],STAT[4,"N.MUT"] - STAT[4,"N.TF.CODING"] - STAT[4,"N.TF.PROMOTER"]),nrow=2,ncol=2)
G.MATRIX.CONSERVATIVE <- matrix(c(STAT[1,"N.TF.CODING"] + STAT[1,"N.TF.PROMOTER"],STAT[1,"N.MUT"] - STAT[1,"N.TF.CODING"] - STAT[1,"N.TF.PROMOTER"],STAT[5,"N.TF.CODING"] + STAT[5,"N.TF.PROMOTER"],STAT[5,"N.MUT"] - STAT[5,"N.TF.CODING"] - STAT[5,"N.TF.PROMOTER"]),nrow=2,ncol=2)

STAT[,"G.STAT.TF.vs.NOT.TF.ALL"] <- c(likelihood.test(G.MATRIX.NEUTRAL,conservative=TRUE)$statistic,likelihood.test(G.MATRIX.CAUSAL,conservative=TRUE)$statistic,likelihood.test(G.MATRIX.RANDOM,conservative=TRUE)$statistic,likelihood.test(G.MATRIX.TAIL,conservative=TRUE)$statistic,likelihood.test(G.MATRIX.CONSERVATIVE,conservative=TRUE)$statistic)

STAT[,"P.VAL.TF.vs.NOT.TF.ALL"] <- c(pchisq(STAT[1,"G.STAT.TF.vs.NOT.TF.ALL"],df=1,lower.tail=FALSE),pchisq(STAT[2,"G.STAT.TF.vs.NOT.TF.ALL"],df=1,lower.tail=FALSE),pchisq(STAT[3,"G.STAT.TF.vs.NOT.TF.ALL"],df=1,lower.tail=FALSE),pchisq(STAT[4,"G.STAT.TF.vs.NOT.TF.ALL"],df=1,lower.tail=FALSE),pchisq(STAT[5,"G.STAT.TF.vs.NOT.TF.ALL"],df=1,lower.tail=FALSE))


G.MATRIX.NEUTRAL <- matrix(c(STAT[1,"N.TF.CODING"]-9,STAT[1,"N.NOT.TF.CODING"],STAT[1,"N.TF.CODING"]-9,STAT[1,"N.NOT.TF.CODING"]),nrow=2,ncol=2)
G.MATRIX.CAUSAL <- matrix(c(STAT[1,"N.TF.CODING"]-9,STAT[1,"N.NOT.TF.CODING"],STAT[2,"N.TF.CODING"]-4,STAT[2,"N.NOT.TF.CODING"]),nrow=2,ncol=2)
G.MATRIX.RANDOM <- matrix(c(STAT[1,"N.TF.CODING"]-9,STAT[1,"N.NOT.TF.CODING"],STAT[3,"N.TF.CODING"]-2,STAT[3,"N.NOT.TF.CODING"]),nrow=2,ncol=2)
G.MATRIX.TAIL <- matrix(c(STAT[1,"N.TF.CODING"]-9,STAT[1,"N.NOT.TF.CODING"],STAT[4,"N.TF.CODING"]-2,STAT[4,"N.NOT.TF.CODING"]),nrow=2,ncol=2)
G.MATRIX.CONSERVATIVE <- matrix(c(STAT[1,"N.TF.CODING"]-9,STAT[1,"N.NOT.TF.CODING"],STAT[5,"N.TF.CODING"]-2,STAT[5,"N.NOT.TF.CODING"]),nrow=2,ncol=2)

STAT[,"G.STAT.TF.NOT.NETWORK"] <- c(likelihood.test(G.MATRIX.NEUTRAL,conservative=TRUE)$statistic,likelihood.test(G.MATRIX.CAUSAL,conservative=TRUE)$statistic,likelihood.test(G.MATRIX.RANDOM,conservative=TRUE)$statistic,likelihood.test(G.MATRIX.TAIL,conservative=TRUE)$statistic,likelihood.test(G.MATRIX.CONSERVATIVE,conservative=TRUE)$statistic)

STAT[,"P.VAL.TF.NOT.NETWORK"] <- c(pchisq(STAT[1,"G.STAT.TF.NOT.NETWORK"],df=1,lower.tail=FALSE),pchisq(STAT[2,"G.STAT.TF.NOT.NETWORK"],df=1,lower.tail=FALSE),pchisq(STAT[3,"G.STAT.TF.NOT.NETWORK"],df=1,lower.tail=FALSE),pchisq(STAT[4,"G.STAT.TF.NOT.NETWORK"],df=1,lower.tail=FALSE),pchisq(STAT[5,"G.STAT.TF.NOT.NETWORK"],df=1,lower.tail=FALSE))

#write.table(STAT,"STAT.TF.FREQUENCY.txt",sep="\t",row.names=FALSE)


############################################################################################################
#O) Testing for enrichment of trans-regulatory mutations in a predicted TDH3 regulatory network (Figure 4).#
############################################################################################################

#Matrices of regulatory interactions between each transcription factor and each yeast gene were obtained from www.yeastract.com.
#These matrices can be found in SupplementaryFile11.zip.
#BINDING is a matrix of pairwise interactions supported by evidence of biochemical interactions between a transcription factor and the promoter of the target gene.
BINDING <- read.table("Network/AllBindingInteractions.txt",header=TRUE)
#POSITIVE is a matrix of pairwise interactions supported by evidence of a transcription factor activating expression of the target gene.
POSITIVE <- read.table("Network/AllPositiveInteractions.txt",header=TRUE)
#NEGATIVE is a matrix of pairwise interactions supported by evidence of a transcription factor inhibiting expression of the target gene.
NEGATIVE <- read.table("Network/AllNegativeInteractions.txt",header=TRUE)

BINDING[,1] -> rownames(BINDING)
POSITIVE[,1] -> rownames(POSITIVE)
NEGATIVE[,1] -> rownames(NEGATIVE)

BINDING <- BINDING[,-1]
POSITIVE <- POSITIVE[,-1]
NEGATIVE <- NEGATIVE[,-1]

NEGATIVE <- NEGATIVE*-1

#Use systematic gene names as defined on www.yeastgenome.org
#Gene.List.txt can be found in SupplementaryFile11.zip
ALL.GENES <- read.table("Gene.List.txt",header=TRUE)
ALL.GENES[,"GENE"]

for (i in 1:nrow(BINDING))
{
  if (rownames(BINDING)[i] %nin% colnames(BINDING))
  {
    CUR.GENE <- subset(ALL.GENES, GENE == rownames(BINDING)[i])
    rownames(BINDING)[i] <- strsplit(as.character(CUR.GENE[1,"SHORT"]),";")[[1]][1]
  }
}

for (i in 1:nrow(POSITIVE))
{
  if (rownames(POSITIVE)[i] %nin% colnames(POSITIVE))
  {
    CUR.GENE <- subset(ALL.GENES, GENE == rownames(POSITIVE)[i])
    rownames(POSITIVE)[i] <- strsplit(as.character(CUR.GENE[1,"SHORT"]),";")[[1]][1]
  }
}

for (i in 1:nrow(NEGATIVE))
{
  if (rownames(NEGATIVE)[i] %nin% colnames(NEGATIVE))
  {
    CUR.GENE <- subset(ALL.GENES, GENE == rownames(NEGATIVE)[i])
    rownames(NEGATIVE)[i] <- strsplit(as.character(CUR.GENE[1,"SHORT"]),";")[[1]][1]
  }
}

rownames(BINDING)[70] <- "AFT1"
rownames(POSITIVE)[70] <- "AFT1"
rownames(NEGATIVE)[70] <- "AFT1"

rownames(BINDING)[105] <- "RSF2"
rownames(POSITIVE)[105] <- "RSF2"
rownames(NEGATIVE)[105] <- "RSF2"

rownames(BINDING)[219] <- "INM2"
rownames(POSITIVE)[219] <- "INM2"
rownames(NEGATIVE)[219] <- "INM2"

BINDING <- as.matrix(BINDING)
POSITIVE <- as.matrix(POSITIVE)
NEGATIVE <- as.matrix(NEGATIVE)

EXPRESSION <- 2*POSITIVE + NEGATIVE
INTERACTOME <- EXPRESSION*BINDING

#write.table(INTERACTOME,"Network/Interactome.txt",sep="\t",row.names=TRUE,col.names=TRUE)

#Tables with TDH3 direct regulators only
TDH3.LEVEL1 <- INTERACTOME[,"TDH3"]
TDH3.LEVEL1 <- TDH3.LEVEL1[which(TDH3.LEVEL1 != 0)]
DIRECT.REGULATORS <- names(TDH3.LEVEL1)

#Tables with TDH3 level1 and level2 regulators
TDH3.LEVEL2 <- INTERACTOME[,c("TDH3",DIRECT.REGULATORS)]

ROW.INFO <- c()

for (i in 1:nrow(TDH3.LEVEL2))
{
  ROW.INFO[i] <- length(which(TDH3.LEVEL2[i,] != 0))
}

TDH3.LEVEL2 <- TDH3.LEVEL2[which(ROW.INFO != 0),]

ALL.REGULATORS <- rownames(TDH3.LEVEL2)
TDH3.LEVEL1.2 <- INTERACTOME[,c("TDH3",ALL.REGULATORS)]

ROW.INFO <- c()

for (i in 1:nrow(TDH3.LEVEL1.2))
{
  ROW.INFO[i] <- length(which(TDH3.LEVEL1.2[i,] != 0))
}

TDH3.LEVEL1.2 <- TDH3.LEVEL1.2[which(ROW.INFO != 0),]
TDH3.LEVEL1.2 <- TDH3.LEVEL1.2[which(rownames(TDH3.LEVEL1.2) %in% colnames(TDH3.LEVEL1.2)),]

write.table(TDH3.LEVEL1.2,"Network/TDH3.Network.txt",sep="\t",row.names=TRUE,col.names=TRUE)


#TDH3.NETWORK is a list of all genes that are in the TDH3 regulatory network.
TDH3.NETWORK <- read.table("Network/TDH3.Network.Genes.txt",header=TRUE)

N.NETWORK <- c(nrow(subset(NEUTRAL, GENE.FOCAL %in% as.character(TDH3.NETWORK[,"GENE"]))),nrow(subset(CAUSAL, GENE.FOCAL %in% as.character(TDH3.NETWORK[,"GENE"]))),nrow(subset(RANDOM, GENE.FOCAL %in% as.character(TDH3.NETWORK[,"GENE"]))),nrow(subset(TAIL, GENE.FOCAL %in% as.character(TDH3.NETWORK[,"GENE"]))),nrow(subset(CONSERVATIVE, GENE.FOCAL %in% as.character(TDH3.NETWORK[,"GENE"]))))
N.NOT.NETWORK <- c(nrow(subset(NEUTRAL, GENE.FOCAL %nin% as.character(TDH3.NETWORK[,"GENE"]))),nrow(subset(CAUSAL, GENE.FOCAL %nin% as.character(TDH3.NETWORK[,"GENE"]))),nrow(subset(RANDOM, GENE.FOCAL %nin% as.character(TDH3.NETWORK[,"GENE"]))),nrow(subset(TAIL, GENE.FOCAL %nin% as.character(TDH3.NETWORK[,"GENE"]))),nrow(subset(CONSERVATIVE, GENE.FOCAL %nin% as.character(TDH3.NETWORK[,"GENE"]))))

STAT <- data.frame(MUT.TYPE, N.MUT, N.NETWORK, N.NOT.NETWORK)

STAT[,"FREQ.NETWORK"] <- STAT[,"N.NETWORK"]/(STAT[,"N.NETWORK"] + STAT[,"N.NOT.NETWORK"])
STAT[,"FREQ.NOT.NETWORK"] <- STAT[,"N.NOT.NETWORK"]/(STAT[,"N.NETWORK"] + STAT[,"N.NOT.NETWORK"])

G.MATRIX.NEUTRAL <- matrix(c(STAT[1,"N.NETWORK"],STAT[1,"N.NOT.NETWORK"],STAT[1,"N.NETWORK"],STAT[1,"N.NOT.NETWORK"]),nrow=2,ncol=2)
G.MATRIX.CAUSAL <- matrix(c(STAT[1,"N.NETWORK"],STAT[1,"N.NOT.NETWORK"],STAT[2,"N.NETWORK"],STAT[2,"N.NOT.NETWORK"]),nrow=2,ncol=2)
G.MATRIX.RANDOM <- matrix(c(STAT[1,"N.NETWORK"],STAT[1,"N.NOT.NETWORK"],STAT[3,"N.NETWORK"],STAT[3,"N.NOT.NETWORK"]),nrow=2,ncol=2)
G.MATRIX.TAIL <- matrix(c(STAT[1,"N.NETWORK"],STAT[1,"N.NOT.NETWORK"],STAT[4,"N.NETWORK"],STAT[4,"N.NOT.NETWORK"]),nrow=2,ncol=2)
G.MATRIX.CONSERVATIVE <- matrix(c(STAT[1,"N.NETWORK"],STAT[1,"N.NOT.NETWORK"],STAT[5,"N.NETWORK"],STAT[5,"N.NOT.NETWORK"]),nrow=2,ncol=2)

STAT[,"G.STAT.NETWORK"] <- c(likelihood.test(G.MATRIX.NEUTRAL,conservative=TRUE)$statistic,likelihood.test(G.MATRIX.CAUSAL,conservative=TRUE)$statistic,likelihood.test(G.MATRIX.RANDOM,conservative=TRUE)$statistic,likelihood.test(G.MATRIX.TAIL,conservative=TRUE)$statistic,likelihood.test(G.MATRIX.CONSERVATIVE,conservative=TRUE)$statistic)

STAT[,"P.VAL.NETWORK"] <- c(pchisq(STAT[1,"G.STAT.NETWORK"],df=1,lower.tail=FALSE),pchisq(STAT[2,"G.STAT.NETWORK"],df=1,lower.tail=FALSE),pchisq(STAT[3,"G.STAT.NETWORK"],df=1,lower.tail=FALSE),pchisq(STAT[4,"G.STAT.NETWORK"],df=1,lower.tail=FALSE),pchisq(STAT[5,"G.STAT.NETWORK"],df=1,lower.tail=FALSE))

#write.table(STAT,"Statistics/STAT.TDH3.NETWORK.txt",sep="\t",row.names=FALSE)


#################################################################################################################
#P) Distributions of the number of mutations in GCR1 or RAP1 regions targeted for PCR mutagenesis (Figure 5C,D).#
#################################################################################################################

###Clear memory###
rm(list=ls())
options(warn=-1)

####Load packages###
##library(VariantAnnotation)
library(VariantAnnotation)
library(Deducer)
library(gtools)
library(zoo)
library(Hmisc)
library(ggplot2)
library(plotrix)
library(moments)
library(vcd)

###Set directory###
setwd("/Path.to.input.files")

#SourceData13.txt is a table containing the number of RAP1 and GCR1 mutations identified by Sanger sequencing in independent clones after PCR mutagenesis.
#SourceData13.txt can be found in SupplementaryFile11.zip.
REG.MUT <- read.table("SourceData13.txt",header=TRUE)


###RAP1 MUTANTS (Figure 5C)###

RAP1.RANDOM <- subset(REG.MUT, Gene == "RAP1" & Class == "Random")
RAP1.TAIL <- subset(REG.MUT, Gene == "RAP1" & Class == "Large.Effect")

#Testing if number of mutations per strain are Poisson distributed.
SEQ.MUT.N <- RAP1.RANDOM[,"N.Mutations"]

GF <- goodfit(SEQ.MUT.N, type = "poisson", method = "MinChisq")
summary(GF)

N.REP <- 1e5

VAR.OBS <- var(SEQ.MUT.N)
KUR.OBS <- kurtosis(SEQ.MUT.N)
MAD.OBS <- mad(SEQ.MUT.N)

VAR.RAND <- rep(0,N.REP)
KUR.RAND <- rep(0,N.REP)
MAD.RAND <- rep(0,N.REP)

for (i in 1:N.REP)
{
  CUR.SAMPLE <- rpois(length(SEQ.MUT.N),mean(SEQ.MUT.N))
  
  VAR.RAND[i] <- var(CUR.SAMPLE)
  KUR.RAND[i] <- kurtosis(CUR.SAMPLE)
  MAD.RAND[i] <- mad(CUR.SAMPLE)
}

LOW.VAR <- length(which(VAR.RAND < VAR.OBS))/length(VAR.RAND)
HIGH.VAR <- length(which(VAR.RAND > VAR.OBS))/length(VAR.RAND)
P.VAR <- 2*min(LOW.VAR,HIGH.VAR)

LOW.KUR <- length(which(KUR.RAND < KUR.OBS))/length(KUR.RAND)
HIGH.KUR <- length(which(KUR.RAND > KUR.OBS))/length(KUR.RAND)
P.KUR <- 2*min(LOW.KUR,HIGH.KUR)

LOW.MAD <- length(which(MAD.RAND < MAD.OBS))/length(MAD.RAND)
HIGH.MAD <- length(which(MAD.RAND > MAD.OBS))/length(MAD.RAND)
P.MAD <- 2*min(LOW.MAD,HIGH.MAD)

BREAKS <- seq(-0.5,7.5,by=1)

POIS <- c()
POS <- c()

for (i in 2:length(BREAKS))
{
  POIS[i-1] <- abs(ppois(BREAKS[i],lambda=mean(SEQ.MUT.N))-ppois(BREAKS[i-1],lambda=mean(SEQ.MUT.N)))*length(SEQ.MUT.N)
  POS[i-1] <- mean(c(BREAKS[i],BREAKS[i-1]))
}

pdf("Figure 5C.pdf",useDingbats=F,height=6,width=7)
#windows(height=6,width=7)
hist(SEQ.MUT.N,breaks=BREAKS,ylim=c(0,0.5),xlab="Number of mutations in RAP1",ylab="Proportion of strains",xaxt="n",freq=F)
axis(1,at=0:7)
points(POS,POIS/sum(POIS),col="blue",pch=16,type="b")
abline(v=mean(SEQ.MUT.N),lty=2)
dev.off()

#Comparing random and large effect mutants
t.test(RAP1.RANDOM$N.Mutations,RAP1.TAIL$N.Mutations)


###GCR1 MUTANTS (Figure 5D)###

GCR1.RANDOM <- subset(REG.MUT, Gene == "GCR1" & Class == "Random")
GCR1.TAIL <- subset(REG.MUT, Gene == "GCR1" & Class == "Large.Effect")

#Testing if number of mutations per strain are Poisson distributed.
SEQ.MUT.N <- GCR1.RANDOM[,"N.Mutations"]

GF <- goodfit(SEQ.MUT.N, type = "poisson", method = "MinChisq")
summary(GF)

N.REP <- 1e5

VAR.OBS <- var(SEQ.MUT.N)
KUR.OBS <- kurtosis(SEQ.MUT.N)
MAD.OBS <- mad(SEQ.MUT.N)

VAR.RAND <- rep(0,N.REP)
KUR.RAND <- rep(0,N.REP)
MAD.RAND <- rep(0,N.REP)

for (i in 1:N.REP)
{
  CUR.SAMPLE <- rpois(length(SEQ.MUT.N),mean(SEQ.MUT.N))
  
  VAR.RAND[i] <- var(CUR.SAMPLE)
  KUR.RAND[i] <- kurtosis(CUR.SAMPLE)
  MAD.RAND[i] <- mad(CUR.SAMPLE)
}

LOW.VAR <- length(which(VAR.RAND < VAR.OBS))/length(VAR.RAND)
HIGH.VAR <- length(which(VAR.RAND > VAR.OBS))/length(VAR.RAND)
P.VAR <- 2*min(LOW.VAR,HIGH.VAR)

LOW.KUR <- length(which(KUR.RAND < KUR.OBS))/length(KUR.RAND)
HIGH.KUR <- length(which(KUR.RAND > KUR.OBS))/length(KUR.RAND)
P.KUR <- 2*min(LOW.KUR,HIGH.KUR)

LOW.MAD <- length(which(MAD.RAND < MAD.OBS))/length(MAD.RAND)
HIGH.MAD <- length(which(MAD.RAND > MAD.OBS))/length(MAD.RAND)
P.MAD <- 2*min(LOW.MAD,HIGH.MAD)

BREAKS <- seq(-0.5,7.5,by=1)

POIS <- c()
POS <- c()

for (i in 2:length(BREAKS))
{
  POIS[i-1] <- abs(ppois(BREAKS[i],lambda=mean(SEQ.MUT.N))-ppois(BREAKS[i-1],lambda=mean(SEQ.MUT.N)))*length(SEQ.MUT.N)
  POS[i-1] <- mean(c(BREAKS[i],BREAKS[i-1]))
}

pdf("Figure 5D.pdf",useDingbats=F,height=6,width=7)
#windows(height=6,width=7)
hist(SEQ.MUT.N,breaks=BREAKS,ylim=c(0,0.5),xlab="Number of mutations in GCR1",ylab="Proportion of strains",xaxt="n",freq=F)
axis(1,at=0:7)
points(POS,POIS/sum(POIS),col="blue",pch=16,type="b")
abline(v=mean(SEQ.MUT.N),lty=2)
dev.off()

#Comparing random and large effect mutants
t.test(GCR1.RANDOM$N.Mutations,GCR1.TAIL$N.Mutations)


#########################################################################
#Q) Number of independent mutations identified in each gene (Figure 6A).#
#########################################################################

#The variables CAUSAL and SUBSITUTION are defined in section H of the script.

CODING <- subset(CAUSAL, CLASS == "EXONIC") 

CODING <- droplevels(CODING)

MUT.GENES <- table(CODING[,"GENE.FOCAL"])
length(MUT.GENES)

OBS.GENES <- length(which(MUT.GENES > 1))

CODING.NEUTRAL <- subset(SUBSTITUTION, CLASS != "INTERGENIC")

#Resampling tests
N.REP <- 1e5
RESAMPLE.GENES <- rep(0,N.REP)

for (i in 1:N.REP)
{
  RESAMPLE <- sample(CODING.NEUTRAL[,"GENE.FOCAL"],size=nrow(CODING),replace=TRUE)
  N.GENES <- table(as.character(RESAMPLE))
  
  RESAMPLE.GENES[i] <- length(which(N.GENES > 1))
}

PROP <- length(which(RESAMPLE.GENES >= OBS.GENES))/length(RESAMPLE.GENES)
P.VAL <- 1-2*abs(0.5-PROP)

#Barplots
N.REP <- 1e3
GENE.MATRIX <- matrix(0,nrow=N.REP,ncol=8)
colnames(GENE.MATRIX) <- 1:8

for (i in 1:N.REP)
{
  RESAMPLE <- sample(CODING.NEUTRAL[,"GENE.FOCAL"],size=nrow(CODING),replace=TRUE)
  N.GENES <- table(as.character(RESAMPLE))
  
  for (j in 1:ncol(GENE.MATRIX))
  {
    GENE.MATRIX[i,j] <- length(which(N.GENES == j))
  }
}

PROP.RANDOM <- rep(0,8)

for (i in 1:length(PROP.RANDOM))
{
  PROP.RANDOM[i] <- mean(GENE.MATRIX[,i])
}

PROP.RANDOM <- PROP.RANDOM/sum(PROP.RANDOM)

PROP.CAUSAL <- rep(0,8)

for (i in 1:length(PROP.CAUSAL))
{
  PROP.CAUSAL[i] <- length(which(MUT.GENES == i))
}

PROP.CAUSAL <- PROP.CAUSAL/sum(PROP.CAUSAL)


PLOT <- as.matrix(rbind(PROP.CAUSAL,PROP.RANDOM))
colnames(PLOT) <- 1:8

pdf("Figure 6a.pdf",height=6,width=7,useDingbats=FALSE)
#windows(height=6,width=7)
barplot(PLOT, xlab = "Number of mutations identified in the same gene", ylab = "Frequency", col=c("orange","blue"),legend=c("65 causative mutations","65 neutral mutations"),ylim=c(0,1),beside=TRUE)
dev.off()


##########################################################
#R) Enriched terms in gene ontology analysis (Figure 6B).#
##########################################################


###Set directory###
setwd("/Path.to.input.files")

#SourceData16.txt is the result of GO term analysis performed on http://www.pantherdb.org/ and it can be found in SupplementaryFile11.zip.
GO <- read.table("SourceData16.txt",header=TRUE)

pdf("Figure 6b_Enrichment.pdf",useDingbats=F,height=10,width=9)
#windows(height=10,width=9)
par(mar=c(5,25,5,2))
barplot(rev(GO[,"Enrichment"]),horiz=T,names=rev(GO[,"Description"]),las=1,col="#2D69B3",xlim=c(0,50),xlab="Enrichment")
axis(3,at=seq(0,50,by=10))
dev.off()

pdf("Figure 6b_P.score.pdf",useDingbats=F,height=10,width=9)
#windows(height=10,width=9)
par(mar=c(5,25,5,2))
barplot(rev(GO[,"P.score"]),horiz=T,names=rev(GO[,"Description"]),las=1,col="#2D69B3",xlim=c(0,5),xlab="-log10(P-value)")
axis(3,at=seq(0,5,by=1))
dev.off()


#####################################################################################
#S) Genomic overlap between trans-regulatory mutations and eQTL regions (Figure 7B).#
#####################################################################################

#"S288c.genome.bed" is a table listing all nucleotides found at each position of each chromosome in the yeast genome.
#The first 10000 lines of this file ("S288c.genome.truncated.bed") can be found in SupplementaryFile11.zip.
#"S288c.genome.bed" was generated from the reference genome ("S288c.mapping.fsa") included in SupplementaryFile11.zip.
CHROM <- read.table("S288c.genome.bed",sep="\t",header=TRUE)

#QTL.Positions.txt is a dataset from Metzger and Wittkopp (2019) describing the genomic positions of all eQTL regions identified in 3 crosses.
#QTL.Positions.txt can be found in SupplementaryFile11.zip.
QTL.POSITIONS <- read.table("QTL.Positions.txt",header=TRUE)


CHROM <- subset(CHROM, CHR %nin% c("chrMito","chrYFP","chrKan","chrNat"))
CHROM <- droplevels(CHROM)
UNIQUE.CHR <- unique(CHROM$CHR)

CHR.LENGTH <- rep(0,length(UNIQUE.CHR))
GENOME.POSITION <- rep(0,length(UNIQUE.CHR))

CHROM.POSITIONS <- data.frame(UNIQUE.CHR, CHR.LENGTH, GENOME.POSITION) 
CHROM.POSITIONS[,"CHR"] <- 1:16

for (i in 1:nrow(CHROM.POSITIONS))
{
  CHROM.POSITIONS[i,"CHR.LENGTH"] <- max(subset(CHROM, CHR == CHROM.POSITIONS[i,"UNIQUE.CHR"])[,"POSITION"],na.rm=T)
  CHROM.POSITIONS[i,"GENOME.POSITION"] <- sum(CHROM.POSITIONS[,"CHR.LENGTH"])
}

#Add positions on genomic coordinates instead of chromosomal coordinates

for (i in 1:nrow(QTL.POSITIONS))
{
  if (QTL.POSITIONS[i,"Chromosome"] == 1)
  {
    QTL.POSITIONS[i,"Lower_Genomic"] <- QTL.POSITIONS[i,"Lower_Bound"]
    QTL.POSITIONS[i,"Higher_Genomic"] <- QTL.POSITIONS[i,"Higher_Bound"]
  } else {
    CUR.CHR <- CHROM.POSITIONS[which(CHROM.POSITIONS[,"CHR"] == QTL.POSITIONS[i,"Chromosome"]-1),"GENOME.POSITION"]
    
    QTL.POSITIONS[i,"Lower_Genomic"] <- QTL.POSITIONS[i,"Lower_Bound"] + CUR.CHR
    QTL.POSITIONS[i,"Higher_Genomic"] <- QTL.POSITIONS[i,"Higher_Bound"] + CUR.CHR
  }
}

SK1 <- subset(QTL.POSITIONS, Cross == "SK1xBY")
M22 <- subset(QTL.POSITIONS, Cross == "M22xBY")
YPS1000 <- subset(QTL.POSITIONS, Cross == "YPS1000xBY")

#Add color to each mutation corresponding to QTL overlap
for (i in 1:nrow(RANDOM))
{
  QTL.SK1 <- length(which(SK1[,"Lower_Genomic"] <= RANDOM[i,"GENOME.POSITION"] & SK1[,"Higher_Genomic"] >= RANDOM[i,"GENOME.POSITION"]))
  QTL.M22 <- length(which(M22[,"Lower_Genomic"] <= RANDOM[i,"GENOME.POSITION"] & M22[,"Higher_Genomic"] >= RANDOM[i,"GENOME.POSITION"]))
  QTL.YPS1000 <- length(which(YPS1000[,"Lower_Genomic"] <= RANDOM[i,"GENOME.POSITION"] & YPS1000[,"Higher_Genomic"] >= RANDOM[i,"GENOME.POSITION"]))
  
  if (QTL.SK1 == 0 & QTL.M22 == 0 & QTL.YPS1000 == 0)
  {
    RANDOM[i,"eQTL"] <- "None"
    RANDOM[i,"eQTL_Color"] <- "black"
  }
  
  if (QTL.SK1 != 0 & QTL.M22 == 0 & QTL.YPS1000 == 0)
  {
    RANDOM[i,"eQTL"] <- "SK1"
    RANDOM[i,"eQTL_Color"] <- "blue"
  }
  
  if (QTL.SK1 == 0 & QTL.M22 == 0 & QTL.YPS1000 != 0)
  {
    RANDOM[i,"eQTL"] <- "YPS1000"
    RANDOM[i,"eQTL_Color"] <- "yellow"
  }
  
  if (QTL.SK1 == 0 & QTL.M22 != 0 & QTL.YPS1000 == 0)
  {
    RANDOM[i,"eQTL"] <- "M22"
    RANDOM[i,"eQTL_Color"] <- "red"
  }
  
  if (QTL.SK1 != 0 & QTL.M22 != 0 & QTL.YPS1000 == 0)
  {
    RANDOM[i,"eQTL"] <- "SK1_M22"
    RANDOM[i,"eQTL_Color"] <- "purple"
  }
  
  if (QTL.SK1 != 0 & QTL.M22 == 0 & QTL.YPS1000 != 0)
  {
    RANDOM[i,"eQTL"] <- "SK1_YPS1000"
    RANDOM[i,"eQTL_Color"] <- "green"
  }
  
  if (QTL.SK1 == 0 & QTL.M22 != 0 & QTL.YPS1000 != 0)
  {
    RANDOM[i,"eQTL"] <- "M22_YPS1000"
    RANDOM[i,"eQTL_Color"] <- "orange"
  }
  
  if (QTL.SK1 != 0 & QTL.M22 != 0 & QTL.YPS1000 != 0)
  {
    RANDOM[i,"eQTL"] <- "SK1_M22_YPS1000"
    RANDOM[i,"eQTL_Color"] <- "brown"
  }
  
}

for (i in 1:nrow(TAIL))
{
  QTL.SK1 <- length(which(SK1[,"Lower_Genomic"] <= TAIL[i,"GENOME.POSITION"] & SK1[,"Higher_Genomic"] >= TAIL[i,"GENOME.POSITION"]))
  QTL.M22 <- length(which(M22[,"Lower_Genomic"] <= TAIL[i,"GENOME.POSITION"] & M22[,"Higher_Genomic"] >= TAIL[i,"GENOME.POSITION"]))
  QTL.YPS1000 <- length(which(YPS1000[,"Lower_Genomic"] <= TAIL[i,"GENOME.POSITION"] & YPS1000[,"Higher_Genomic"] >= TAIL[i,"GENOME.POSITION"]))
  
  if (QTL.SK1 == 0 & QTL.M22 == 0 & QTL.YPS1000 == 0)
  {
    TAIL[i,"eQTL"] <- "None"
    TAIL[i,"eQTL_Color"] <- "black"
  }
  
  if (QTL.SK1 != 0 & QTL.M22 == 0 & QTL.YPS1000 == 0)
  {
    TAIL[i,"eQTL"] <- "SK1"
    TAIL[i,"eQTL_Color"] <- "blue"
  }
  
  if (QTL.SK1 == 0 & QTL.M22 == 0 & QTL.YPS1000 != 0)
  {
    TAIL[i,"eQTL"] <- "YPS1000"
    TAIL[i,"eQTL_Color"] <- "yellow"
  }
  
  if (QTL.SK1 == 0 & QTL.M22 != 0 & QTL.YPS1000 == 0)
  {
    TAIL[i,"eQTL"] <- "M22"
    TAIL[i,"eQTL_Color"] <- "red"
  }
  
  if (QTL.SK1 != 0 & QTL.M22 != 0 & QTL.YPS1000 == 0)
  {
    TAIL[i,"eQTL"] <- "SK1_M22"
    TAIL[i,"eQTL_Color"] <- "purple"
  }
  
  if (QTL.SK1 != 0 & QTL.M22 == 0 & QTL.YPS1000 != 0)
  {
    TAIL[i,"eQTL"] <- "SK1_YPS1000"
    TAIL[i,"eQTL_Color"] <- "green"
  }
  
  if (QTL.SK1 == 0 & QTL.M22 != 0 & QTL.YPS1000 != 0)
  {
    TAIL[i,"eQTL"] <- "M22_YPS1000"
    TAIL[i,"eQTL_Color"] <- "orange"
  }
  
  if (QTL.SK1 != 0 & QTL.M22 != 0 & QTL.YPS1000 != 0)
  {
    TAIL[i,"eQTL"] <- "SK1_M22_YPS1000"
    TAIL[i,"eQTL_Color"] <- "brown"
  }
  
}


pdf("Figure 7A.pdf",height=5,width=14,useDingbats=FALSE)

#windows(height=5,width=14)

plot(0,0,type="n",xlim=c(0,max(CHROM.POSITIONS[,"GENOME.POSITION"])),xaxt="n",yaxt="n",xlab="Genomic position (Mb)",ylim=c(0,4),ylab="")
axis(1,at=c(0,CHROM.POSITIONS[,"GENOME.POSITION"]))
abline(v=CHROM.POSITIONS[,"GENOME.POSITION"],lty=2,col="#00000066")

for (i in 1:nrow(SK1))
{
  rect(xleft=SK1[i,"Lower_Genomic"],ybottom=0,xright=SK1[i,"Higher_Genomic"],ytop=1,col="blue",border=NA)
}

for (i in 1:nrow(YPS1000))
{
  rect(xleft=YPS1000[i,"Lower_Genomic"],ybottom=1,xright=YPS1000[i,"Higher_Genomic"],ytop=2,col="yellow",border=NA)
}

for (i in 1:nrow(M22))
{
  rect(xleft=M22[i,"Lower_Genomic"],ybottom=2,xright=M22[i,"Higher_Genomic"],ytop=3,col="red",border=NA)
}

points(RANDOM[,"GENOME.POSITION"],rep(3.3,nrow(RANDOM)),col=RANDOM[,"eQTL_Color"],pch=6,cex=2)
points(TAIL[,"GENOME.POSITION"],rep(3.7,nrow(TAIL)),col=TAIL[,"eQTL_Color"],pch=6,cex=2)

dev.off()


######################################################################################################
#T) Proportions of non-regulatory and trans-regulatory mutations located in eQTL regions (Figure 7B).#
######################################################################################################

SUB.NEUTRAL <- subset(NEUTRAL, CHROMOSOME %nin% c("chrMito","chrYFP","chrKan","chrNat"))

for (i in 1:nrow(SUB.NEUTRAL))
{
  QTL.SK1 <- length(which(SK1[,"Lower_Genomic"] <= SUB.NEUTRAL[i,"GENOME.POSITION"] & SK1[,"Higher_Genomic"] >= SUB.NEUTRAL[i,"GENOME.POSITION"]))
  QTL.M22 <- length(which(M22[,"Lower_Genomic"] <= SUB.NEUTRAL[i,"GENOME.POSITION"] & M22[,"Higher_Genomic"] >= SUB.NEUTRAL[i,"GENOME.POSITION"]))
  QTL.YPS1000 <- length(which(YPS1000[,"Lower_Genomic"] <= SUB.NEUTRAL[i,"GENOME.POSITION"] & YPS1000[,"Higher_Genomic"] >= SUB.NEUTRAL[i,"GENOME.POSITION"]))
  
  if (QTL.SK1 == 0 & QTL.M22 == 0 & QTL.YPS1000 == 0)
  {
    SUB.NEUTRAL[i,"eQTL"] <- "None"
    SUB.NEUTRAL[i,"eQTL_Color"] <- "black"
  }
  
  if (QTL.SK1 != 0 & QTL.M22 == 0 & QTL.YPS1000 == 0)
  {
    SUB.NEUTRAL[i,"eQTL"] <- "SK1"
    SUB.NEUTRAL[i,"eQTL_Color"] <- "blue"
  }
  
  if (QTL.SK1 == 0 & QTL.M22 == 0 & QTL.YPS1000 != 0)
  {
    SUB.NEUTRAL[i,"eQTL"] <- "YPS1000"
    SUB.NEUTRAL[i,"eQTL_Color"] <- "yellow"
  }
  
  if (QTL.SK1 == 0 & QTL.M22 != 0 & QTL.YPS1000 == 0)
  {
    SUB.NEUTRAL[i,"eQTL"] <- "M22"
    SUB.NEUTRAL[i,"eQTL_Color"] <- "red"
  }
  
  if (QTL.SK1 != 0 & QTL.M22 != 0 & QTL.YPS1000 == 0)
  {
    SUB.NEUTRAL[i,"eQTL"] <- "SK1_M22"
    SUB.NEUTRAL[i,"eQTL_Color"] <- "purple"
  }
  
  if (QTL.SK1 != 0 & QTL.M22 == 0 & QTL.YPS1000 != 0)
  {
    SUB.NEUTRAL[i,"eQTL"] <- "SK1_YPS1000"
    SUB.NEUTRAL[i,"eQTL_Color"] <- "green"
  }
  
  if (QTL.SK1 == 0 & QTL.M22 != 0 & QTL.YPS1000 != 0)
  {
    SUB.NEUTRAL[i,"eQTL"] <- "M22_YPS1000"
    SUB.NEUTRAL[i,"eQTL_Color"] <- "orange"
  }
  
  if (QTL.SK1 != 0 & QTL.M22 != 0 & QTL.YPS1000 != 0)
  {
    SUB.NEUTRAL[i,"eQTL"] <- "SK1_M22_YPS1000"
    SUB.NEUTRAL[i,"eQTL_Color"] <- "brown"
  }
  
}

#eQTL fraction whole genome

GENOME <- SUB.NEUTRAL[,c("eQTL","eQTL_Color")]

colnames(GENOME) <- c("eQTL","eQTL_Color") 
GENOME[,"eQTL"] <- as.character(GENOME[,"eQTL"])
GENOME[,"eQTL_Color"] <- as.character(GENOME[,"eQTL_Color"])

eQTL <- rep("SK1",max(CHROM.POSITIONS$GENOME.POSITION))
eQTL_Color <- rep("black",max(CHROM.POSITIONS$GENOME.POSITION))


for(i in 1:max(CHROM.POSITIONS$GENOME.POSITION))
{
  QTL.SK1 <- length(which(SK1[,"Lower_Genomic"] <= i & SK1[,"Higher_Genomic"] >= i))
  QTL.YPS1000 <- length(which(YPS1000[,"Lower_Genomic"] <= i & YPS1000[,"Higher_Genomic"] >= i))
  QTL.M22 <- length(which(M22[,"Lower_Genomic"] <= i & M22[,"Higher_Genomic"] >= i))
  
  if (QTL.SK1 == 0 & QTL.M22 == 0 & QTL.YPS1000 == 0)
  {
    eQTL[i] <- "None"
    eQTL_Color[i] <- "black"
  }
  
  if (QTL.SK1 != 0 & QTL.M22 == 0 & QTL.YPS1000 == 0)
  {
    eQTL[i] <- "SK1"
    eQTL_Color[i] <- "blue"
  }
  
  if (QTL.SK1 == 0 & QTL.M22 == 0 & QTL.YPS1000 != 0)
  {
    eQTL[i] <- "YPS1000"
    eQTL_Color[i] <- "yellow"
  }
  
  if (QTL.SK1 == 0 & QTL.M22 != 0 & QTL.YPS1000 == 0)
  {
    eQTL[i] <- "M22"
    eQTL_Color[i] <- "red"
  }
  
  if (QTL.SK1 != 0 & QTL.M22 != 0 & QTL.YPS1000 == 0)
  {
    eQTL[i] <- "SK1_M22"
    eQTL_Color[i] <- "purple"
  }
  
  if (QTL.SK1 != 0 & QTL.M22 == 0 & QTL.YPS1000 != 0)
  {
    eQTL[i] <- "SK1_YPS1000"
    eQTL_Color[i] <- "green"
  }
  
  if (QTL.SK1 == 0 & QTL.M22 != 0 & QTL.YPS1000 != 0)
  {
    eQTL[i] <- "M22_YPS1000"
    eQTL_Color[i] <- "orange"
  }
  
  if (QTL.SK1 != 0 & QTL.M22 != 0 & QTL.YPS1000 != 0)
  {
    eQTL[i] <- "SK1_M22_YPS1000"
    eQTL_Color[i] <- "brown"
  }
  
}

GENOME <- data.frame(1:max(CHROM.POSITIONS$GENOME.POSITION),eQTL,eQTL_Color)

write.table(GENOME,"eQTL_Genome.txt",sep="\t",row.names=FALSE)

#Make matrix with frequencies to plot
GENOME <- read.table(file = "eQTL_Genome.txt", header = TRUE)

FREQ.QTL <- matrix(0,nrow=5,ncol=4)
colnames(FREQ.QTL) <- c("Any","SK1","YPS1000","M22")
rownames(FREQ.QTL) <- c("Genome","Neutral","Causative","Random","Tail")

ALL.eQTL <- c("SK1","YPS1000","M22","SK1_M22","SK1_YPS1000","SK1_M22_YPS1000","M22_YPS1000")
SK1.eQTL <- c("SK1","SK1_M22","SK1_YPS1000","SK1_M22_YPS1000")
YPS1000.eQTL <- c("YPS1000","M22_YPS1000","SK1_YPS1000","SK1_M22_YPS1000")
M22.eQTL <- c("M22","SK1_M22","M22_YPS1000","SK1_M22_YPS1000")

eQTL.REGIONS <- list(ALL.eQTL,SK1.eQTL,YPS1000.eQTL,M22.eQTL)

SUB.NEUTRAL <- subset(SUB.NEUTRAL, GENOME.POSITION > 0)
RANDOM <- subset(RANDOM, GENOME.POSITION > 0)
TAIL <- subset(TAIL, GENOME.POSITION > 0)


for (i in 1:4)
{
  FREQ.QTL[1,i] <- length(which(GENOME[,"eQTL"] %in% eQTL.REGIONS[[i]]))/nrow(GENOME)
  FREQ.QTL[2,i] <- length(which(SUB.NEUTRAL[,"eQTL"] %in% eQTL.REGIONS[[i]]))/nrow(SUB.NEUTRAL)
  FREQ.QTL[3,i] <- (length(which(RANDOM[,"eQTL"] %in% eQTL.REGIONS[[i]])) + length(which(TAIL[,"eQTL"] %in% eQTL.REGIONS[[i]])))/(nrow(RANDOM)+nrow(TAIL))
  FREQ.QTL[4,i] <- length(which(RANDOM[,"eQTL"] %in% eQTL.REGIONS[[i]]))/nrow(RANDOM)
  FREQ.QTL[5,i] <- length(which(TAIL[,"eQTL"] %in% eQTL.REGIONS[[i]]))/nrow(TAIL)
}

N.QTL <- matrix(0,nrow=5,ncol=4)
colnames(N.QTL) <- c("Any","SK1","YPS1000","M22")
rownames(N.QTL) <- c("Genome","Neutral","Causative","Random","Tail")

for (i in 1:4)
{
  N.QTL[1,i] <- length(which(GENOME[,"eQTL"] %in% eQTL.REGIONS[[i]]))
  N.QTL[2,i] <- length(which(SUB.NEUTRAL[,"eQTL"] %in% eQTL.REGIONS[[i]]))
  N.QTL[3,i] <- length(which(RANDOM[,"eQTL"] %in% eQTL.REGIONS[[i]])) + length(which(TAIL[,"eQTL"] %in% eQTL.REGIONS[[i]]))
  N.QTL[4,i] <- length(which(RANDOM[,"eQTL"] %in% eQTL.REGIONS[[i]]))
  N.QTL[5,i] <- length(which(TAIL[,"eQTL"] %in% eQTL.REGIONS[[i]]))
}


pdf("Figure 7B.pdf",height=5,width=13,useDingbats=FALSE)

#windows(height=5,width=13)
barplot(FREQ.QTL,xlab="",ylab="Frequency",col=c("black","blue","orange","red","green"),ylim=c(0,1),legend=rownames(FREQ.QTL),beside=TRUE)

dev.off()


####################################################################################
#U) Statistical tests for enrichment of trans-regulatory mutations in eQTL regions.#
####################################################################################


###a) Analyses including all eQTL regions from the 3 crosses.###

#QTL.Positions.txt is a dataset from Metzger and Wittkopp (2019) describing the genomic positions of all eQTL regions identified in 3 crosses.
#QTL.Positions.txt can be found in SupplementaryFile11.zip.
QTL.POSITIONS <- read.table("QTL.Positions.txt",header=TRUE)

#Tables indicating for different types of mutations whether each mutation is located in one or more eQTL regions.
CHR.NEUTRAL <- SUB.NEUTRAL

for (i in 1:nrow(CHR.NEUTRAL))
{
  CUR.CHR <- as.integer(strsplit(as.character(CHR.NEUTRAL[i,"CHROMOSOME"]),"chr")[[1]][2])
  
  CUR.QTL <- subset(QTL.POSITIONS, Chromosome == CUR.CHR & Lower_Bound <= CHR.NEUTRAL[i,"POSITION"] & Higher_Bound >= CHR.NEUTRAL[i,"POSITION"])
  CHR.NEUTRAL[i,"N.eQTL"] <- nrow(CUR.QTL)
}

CHR.CAUSAL <- rbind(RANDOM,TAIL)

for (i in 1:nrow(CHR.CAUSAL))
{
  CUR.CHR <- as.integer(strsplit(as.character(CHR.CAUSAL[i,"CHROMOSOME"]),"chr")[[1]][2])
  
  CUR.QTL <- subset(QTL.POSITIONS, Chromosome == CUR.CHR & Lower_Bound <= CHR.CAUSAL[i,"POSITION"] & Higher_Bound >= CHR.CAUSAL[i,"POSITION"])
  CHR.CAUSAL[i,"N.eQTL"] <- nrow(CUR.QTL)
}

CHR.RANDOM <- subset(RANDOM, CHROMOSOME %nin% c("chrMito","chrYFP"))

for (i in 1:nrow(CHR.RANDOM))
{
  CUR.CHR <- as.integer(strsplit(as.character(CHR.RANDOM[i,"CHROMOSOME"]),"chr")[[1]][2])
  
  CUR.QTL <- subset(QTL.POSITIONS, Chromosome == CUR.CHR & Lower_Bound <= CHR.RANDOM[i,"POSITION"] & Higher_Bound >= CHR.RANDOM[i,"POSITION"])
  CHR.RANDOM[i,"N.eQTL"] <- nrow(CUR.QTL)
}

CHR.TAIL <- subset(TAIL, CHROMOSOME %nin% c("chrMito","chrYFP"))

for (i in 1:nrow(CHR.TAIL))
{
  CUR.CHR <- as.integer(strsplit(as.character(CHR.TAIL[i,"CHROMOSOME"]),"chr")[[1]][2])
  
  CUR.QTL <- subset(QTL.POSITIONS, Chromosome == CUR.CHR & Lower_Bound <= CHR.TAIL[i,"POSITION"] & Higher_Bound >= CHR.TAIL[i,"POSITION"])
  CHR.TAIL[i,"N.eQTL"] <- nrow(CUR.QTL)
}

CHR.CONSERVATIVE <- subset(CONSERVATIVE, CHROMOSOME %nin% c("chrMito","chrYFP"))

for (i in 1:nrow(CHR.CONSERVATIVE))
{
  CUR.CHR <- as.integer(strsplit(as.character(CHR.CONSERVATIVE[i,"CHROMOSOME"]),"chr")[[1]][2])
  
  CUR.QTL <- subset(QTL.POSITIONS, Chromosome == CUR.CHR & Lower_Bound <= CHR.CONSERVATIVE[i,"POSITION"] & Higher_Bound >= CHR.CONSERVATIVE[i,"POSITION"])
  CHR.CONSERVATIVE[i,"N.eQTL"] <- nrow(CUR.QTL)
}

CHR.COD.NEUTRAL <- subset(SUBSTITUTION, CHROMOSOME %nin% c("chrMito","chrYFP") & CLASS == "EXONIC")

for (i in 1:nrow(CHR.COD.NEUTRAL))
{
  CUR.CHR <- as.integer(strsplit(as.character(CHR.COD.NEUTRAL[i,"CHROMOSOME"]),"chr")[[1]][2])
  
  CUR.QTL <- subset(QTL.POSITIONS, Chromosome == CUR.CHR & Lower_Bound <= CHR.COD.NEUTRAL[i,"POSITION"] & Higher_Bound >= CHR.COD.NEUTRAL[i,"POSITION"])
  CHR.COD.NEUTRAL[i,"N.eQTL"] <- nrow(CUR.QTL)
}

CHR.COD.CAUSAL <- subset(CAUSAL, CHROMOSOME %nin% c("chrMito","chrYFP") & CLASS == "EXONIC")

for (i in 1:nrow(CHR.COD.CAUSAL))
{
  CUR.CHR <- as.integer(strsplit(as.character(CHR.COD.CAUSAL[i,"CHROMOSOME"]),"chr")[[1]][2])
  
  CUR.QTL <- subset(QTL.POSITIONS, Chromosome == CUR.CHR & Lower_Bound <= CHR.COD.CAUSAL[i,"POSITION"] & Higher_Bound >= CHR.COD.CAUSAL[i,"POSITION"])
  CHR.COD.CAUSAL[i,"N.eQTL"] <- nrow(CUR.QTL)
}

CHR.COD.RANDOM <- subset(RANDOM, CHROMOSOME %nin% c("chrMito","chrYFP") & CLASS == "EXONIC")

for (i in 1:nrow(CHR.COD.RANDOM))
{
  CUR.CHR <- as.integer(strsplit(as.character(CHR.COD.RANDOM[i,"CHROMOSOME"]),"chr")[[1]][2])
  
  CUR.QTL <- subset(QTL.POSITIONS, Chromosome == CUR.CHR & Lower_Bound <= CHR.COD.RANDOM[i,"POSITION"] & Higher_Bound >= CHR.COD.RANDOM[i,"POSITION"])
  CHR.COD.RANDOM[i,"N.eQTL"] <- nrow(CUR.QTL)
}

CHR.COD.TAIL <- subset(TAIL, CHROMOSOME %nin% c("chrMito","chrYFP") & CLASS == "EXONIC")

for (i in 1:nrow(CHR.COD.TAIL))
{
  CUR.CHR <- as.integer(strsplit(as.character(CHR.COD.TAIL[i,"CHROMOSOME"]),"chr")[[1]][2])
  
  CUR.QTL <- subset(QTL.POSITIONS, Chromosome == CUR.CHR & Lower_Bound <= CHR.COD.TAIL[i,"POSITION"] & Higher_Bound >= CHR.COD.TAIL[i,"POSITION"])
  CHR.COD.TAIL[i,"N.eQTL"] <- nrow(CUR.QTL)
}

CHR.COD.CONSERVATIVE <- subset(CONSERVATIVE, CHROMOSOME %nin% c("chrMito","chrYFP") & CLASS == "EXONIC")

for (i in 1:nrow(CHR.COD.TAIL))
{
  CUR.CHR <- as.integer(strsplit(as.character(CHR.COD.CONSERVATIVE[i,"CHROMOSOME"]),"chr")[[1]][2])
  
  CUR.QTL <- subset(QTL.POSITIONS, Chromosome == CUR.CHR & Lower_Bound <= CHR.COD.CONSERVATIVE[i,"POSITION"] & Higher_Bound >= CHR.COD.CONSERVATIVE[i,"POSITION"])
  CHR.COD.CONSERVATIVE[i,"N.eQTL"] <- nrow(CUR.QTL)
}

#G-tests used to compare the frequency of non-regulatory mutations and other types of mutations located in eQTL regions.
N.MUT <- c(nrow(CHR.NEUTRAL),nrow(CHR.CAUSAL),nrow(CHR.RANDOM),nrow(CHR.TAIL),nrow(CHR.CONSERVATIVE))
N.NOT.QTL <- c(length(which(CHR.NEUTRAL$N.eQTL == 0)),length(which(CHR.CAUSAL$N.eQTL == 0)),length(which(CHR.RANDOM$N.eQTL == 0)),length(which(CHR.TAIL$N.eQTL == 0)),length(which(CHR.CONSERVATIVE$N.eQTL == 0)))
N.QTL <- N.MUT - N.NOT.QTL
N.CODING <- c(nrow(CHR.COD.NEUTRAL),nrow(CHR.COD.CAUSAL),nrow(CHR.COD.RANDOM),nrow(CHR.COD.TAIL),nrow(CHR.COD.CONSERVATIVE))
N.CODING.NOT.QTL <- c(length(which(CHR.COD.NEUTRAL$N.eQTL == 0)),length(which(CHR.COD.CAUSAL$N.eQTL == 0)),length(which(CHR.COD.RANDOM$N.eQTL == 0)),length(which(CHR.COD.TAIL$N.eQTL == 0)),length(which(CHR.COD.CONSERVATIVE$N.eQTL == 0)))
N.CODING.QTL <- N.CODING - N.CODING.NOT.QTL

FREQ.QTL <- round(N.QTL/N.MUT,3)
FREQ.NOT.QTL <- round(N.NOT.QTL/N.MUT,3)
FREQ.CODING.QTL <- round(N.CODING.QTL/N.CODING,3)
FREQ.CODING.NOT.QTL <- round(N.CODING.NOT.QTL/N.CODING,3)

STAT <- data.frame(MUT.TYPE, N.MUT, N.QTL, N.NOT.QTL, N.CODING, N.CODING.QTL, N.CODING.NOT.QTL, FREQ.QTL, FREQ.NOT.QTL, FREQ.CODING.QTL, FREQ.CODING.NOT.QTL)

G.MATRIX.NEUTRAL <- matrix(c(STAT[1,"N.QTL"],STAT[1,"N.NOT.QTL"],STAT[1,"N.QTL"],STAT[1,"N.NOT.QTL"]),nrow=2,ncol=2)
G.MATRIX.CAUSAL <- matrix(c(STAT[1,"N.QTL"],STAT[1,"N.NOT.QTL"],STAT[2,"N.QTL"],STAT[2,"N.NOT.QTL"]),nrow=2,ncol=2)
G.MATRIX.RANDOM <- matrix(c(STAT[1,"N.QTL"],STAT[1,"N.NOT.QTL"],STAT[3,"N.QTL"],STAT[3,"N.NOT.QTL"]),nrow=2,ncol=2)
G.MATRIX.TAIL <- matrix(c(STAT[1,"N.QTL"],STAT[1,"N.NOT.QTL"],STAT[4,"N.QTL"],STAT[4,"N.NOT.QTL"]),nrow=2,ncol=2)
G.MATRIX.CONSERVATIVE <- matrix(c(STAT[1,"N.QTL"],STAT[1,"N.NOT.QTL"],STAT[5,"N.QTL"],STAT[5,"N.NOT.QTL"]),nrow=2,ncol=2)

STAT[,"G.STAT.QTL"] <- c(likelihood.test(G.MATRIX.NEUTRAL,conservative=TRUE)$statistic,likelihood.test(G.MATRIX.CAUSAL,conservative=TRUE)$statistic,likelihood.test(G.MATRIX.RANDOM,conservative=TRUE)$statistic,likelihood.test(G.MATRIX.TAIL,conservative=TRUE)$statistic,likelihood.test(G.MATRIX.CONSERVATIVE,conservative=TRUE)$statistic)

STAT[,"P.VAL.QTL"] <- c(pchisq(STAT[1,"G.STAT.QTL"],df=1,lower.tail=FALSE),pchisq(STAT[2,"G.STAT.QTL"],df=1,lower.tail=FALSE),pchisq(STAT[3,"G.STAT.QTL"],df=1,lower.tail=FALSE),pchisq(STAT[4,"G.STAT.QTL"],df=1,lower.tail=FALSE),pchisq(STAT[5,"G.STAT.QTL"],df=1,lower.tail=FALSE))


G.MATRIX.NEUTRAL <- matrix(c(STAT[1,"N.CODING.QTL"],STAT[1,"N.CODING.NOT.QTL"],STAT[1,"N.CODING.QTL"],STAT[1,"N.CODING.NOT.QTL"]),nrow=2,ncol=2)
G.MATRIX.CAUSAL <- matrix(c(STAT[1,"N.CODING.QTL"],STAT[1,"N.CODING.NOT.QTL"],STAT[2,"N.CODING.QTL"],STAT[2,"N.CODING.NOT.QTL"]),nrow=2,ncol=2)
G.MATRIX.RANDOM <- matrix(c(STAT[1,"N.CODING.QTL"],STAT[1,"N.CODING.NOT.QTL"],STAT[3,"N.CODING.QTL"],STAT[3,"N.CODING.NOT.QTL"]),nrow=2,ncol=2)
G.MATRIX.TAIL <- matrix(c(STAT[1,"N.CODING.QTL"],STAT[1,"N.CODING.NOT.QTL"],STAT[4,"N.CODING.QTL"],STAT[4,"N.CODING.NOT.QTL"]),nrow=2,ncol=2)
G.MATRIX.CONSERVATIVE <- matrix(c(STAT[1,"N.CODING.QTL"],STAT[1,"N.CODING.NOT.QTL"],STAT[5,"N.CODING.QTL"],STAT[5,"N.CODING.NOT.QTL"]),nrow=2,ncol=2)

STAT[,"G.STAT.CODING.QTL"] <- c(likelihood.test(G.MATRIX.NEUTRAL,conservative=TRUE)$statistic,likelihood.test(G.MATRIX.CAUSAL,conservative=TRUE)$statistic,likelihood.test(G.MATRIX.RANDOM,conservative=TRUE)$statistic,likelihood.test(G.MATRIX.TAIL,conservative=TRUE)$statistic,likelihood.test(G.MATRIX.CONSERVATIVE,conservative=TRUE)$statistic)

STAT[,"P.VAL.CODING.QTL"] <- c(pchisq(STAT[1,"G.STAT.CODING.QTL"],df=1,lower.tail=FALSE),pchisq(STAT[2,"G.STAT.CODING.QTL"],df=1,lower.tail=FALSE),pchisq(STAT[3,"G.STAT.CODING.QTL"],df=1,lower.tail=FALSE),pchisq(STAT[4,"G.STAT.CODING.QTL"],df=1,lower.tail=FALSE),pchisq(STAT[5,"G.STAT.CODING.QTL"],df=1,lower.tail=FALSE))


###b) Analyses including only eQTL regions from YPS1000xBY cross.###

#QTL.Positions.txt is a dataset from Metzger and Wittkopp (2019) describing the genomic positions of all eQTL regions identified in 3 crosses.
#QTL.Positions.txt can be found in SupplementaryFile11.zip.
QTL.POSITIONS <- read.table("QTL.Positions.txt",header=TRUE)

QTL.POSITIONS <- subset(QTL.POSITIONS, Cross == "YPS1000xBY")

CHR.NEUTRAL <- SUB.NEUTRAL

for (i in 1:nrow(CHR.NEUTRAL))
{
  CUR.CHR <- as.integer(strsplit(as.character(CHR.NEUTRAL[i,"CHROMOSOME"]),"chr")[[1]][2])
  
  CUR.QTL <- subset(QTL.POSITIONS, Chromosome == CUR.CHR & Lower_Bound <= CHR.NEUTRAL[i,"POSITION"] & Higher_Bound >= CHR.NEUTRAL[i,"POSITION"])
  CHR.NEUTRAL[i,"N.eQTL"] <- nrow(CUR.QTL)
}

CHR.CAUSAL <- rbind(RANDOM,TAIL)

for (i in 1:nrow(CHR.CAUSAL))
{
  CUR.CHR <- as.integer(strsplit(as.character(CHR.CAUSAL[i,"CHROMOSOME"]),"chr")[[1]][2])
  
  CUR.QTL <- subset(QTL.POSITIONS, Chromosome == CUR.CHR & Lower_Bound <= CHR.CAUSAL[i,"POSITION"] & Higher_Bound >= CHR.CAUSAL[i,"POSITION"])
  CHR.CAUSAL[i,"N.eQTL"] <- nrow(CUR.QTL)
}

CHR.RANDOM <- subset(RANDOM, CHROMOSOME %nin% c("chrMito","chrYFP"))

for (i in 1:nrow(CHR.RANDOM))
{
  CUR.CHR <- as.integer(strsplit(as.character(CHR.RANDOM[i,"CHROMOSOME"]),"chr")[[1]][2])
  
  CUR.QTL <- subset(QTL.POSITIONS, Chromosome == CUR.CHR & Lower_Bound <= CHR.RANDOM[i,"POSITION"] & Higher_Bound >= CHR.RANDOM[i,"POSITION"])
  CHR.RANDOM[i,"N.eQTL"] <- nrow(CUR.QTL)
}

CHR.TAIL <- subset(TAIL, CHROMOSOME %nin% c("chrMito","chrYFP"))

for (i in 1:nrow(CHR.TAIL))
{
  CUR.CHR <- as.integer(strsplit(as.character(CHR.TAIL[i,"CHROMOSOME"]),"chr")[[1]][2])
  
  CUR.QTL <- subset(QTL.POSITIONS, Chromosome == CUR.CHR & Lower_Bound <= CHR.TAIL[i,"POSITION"] & Higher_Bound >= CHR.TAIL[i,"POSITION"])
  CHR.TAIL[i,"N.eQTL"] <- nrow(CUR.QTL)
}

CHR.CONSERVATIVE <- subset(CONSERVATIVE, CHROMOSOME %nin% c("chrMito","chrYFP"))

for (i in 1:nrow(CHR.CONSERVATIVE))
{
  CUR.CHR <- as.integer(strsplit(as.character(CHR.CONSERVATIVE[i,"CHROMOSOME"]),"chr")[[1]][2])
  
  CUR.QTL <- subset(QTL.POSITIONS, Chromosome == CUR.CHR & Lower_Bound <= CHR.CONSERVATIVE[i,"POSITION"] & Higher_Bound >= CHR.CONSERVATIVE[i,"POSITION"])
  CHR.CONSERVATIVE[i,"N.eQTL"] <- nrow(CUR.QTL)
}

CHR.COD.NEUTRAL <- subset(SUBSTITUTION, CHROMOSOME %nin% c("chrMito","chrYFP") & CLASS == "EXONIC")

for (i in 1:nrow(CHR.COD.NEUTRAL))
{
  CUR.CHR <- as.integer(strsplit(as.character(CHR.COD.NEUTRAL[i,"CHROMOSOME"]),"chr")[[1]][2])
  
  CUR.QTL <- subset(QTL.POSITIONS, Chromosome == CUR.CHR & Lower_Bound <= CHR.COD.NEUTRAL[i,"POSITION"] & Higher_Bound >= CHR.COD.NEUTRAL[i,"POSITION"])
  CHR.COD.NEUTRAL[i,"N.eQTL"] <- nrow(CUR.QTL)
}

CHR.COD.CAUSAL <- subset(CAUSAL, CHROMOSOME %nin% c("chrMito","chrYFP") & CLASS == "EXONIC")

for (i in 1:nrow(CHR.COD.CAUSAL))
{
  CUR.CHR <- as.integer(strsplit(as.character(CHR.COD.CAUSAL[i,"CHROMOSOME"]),"chr")[[1]][2])
  
  CUR.QTL <- subset(QTL.POSITIONS, Chromosome == CUR.CHR & Lower_Bound <= CHR.COD.CAUSAL[i,"POSITION"] & Higher_Bound >= CHR.COD.CAUSAL[i,"POSITION"])
  CHR.COD.CAUSAL[i,"N.eQTL"] <- nrow(CUR.QTL)
}

CHR.COD.RANDOM <- subset(RANDOM, CHROMOSOME %nin% c("chrMito","chrYFP") & CLASS == "EXONIC")

for (i in 1:nrow(CHR.COD.RANDOM))
{
  CUR.CHR <- as.integer(strsplit(as.character(CHR.COD.RANDOM[i,"CHROMOSOME"]),"chr")[[1]][2])
  
  CUR.QTL <- subset(QTL.POSITIONS, Chromosome == CUR.CHR & Lower_Bound <= CHR.COD.RANDOM[i,"POSITION"] & Higher_Bound >= CHR.COD.RANDOM[i,"POSITION"])
  CHR.COD.RANDOM[i,"N.eQTL"] <- nrow(CUR.QTL)
}

CHR.COD.TAIL <- subset(TAIL, CHROMOSOME %nin% c("chrMito","chrYFP") & CLASS == "EXONIC")

for (i in 1:nrow(CHR.COD.TAIL))
{
  CUR.CHR <- as.integer(strsplit(as.character(CHR.COD.TAIL[i,"CHROMOSOME"]),"chr")[[1]][2])
  
  CUR.QTL <- subset(QTL.POSITIONS, Chromosome == CUR.CHR & Lower_Bound <= CHR.COD.TAIL[i,"POSITION"] & Higher_Bound >= CHR.COD.TAIL[i,"POSITION"])
  CHR.COD.TAIL[i,"N.eQTL"] <- nrow(CUR.QTL)
}

CHR.COD.CONSERVATIVE <- subset(CONSERVATIVE, CHROMOSOME %nin% c("chrMito","chrYFP") & CLASS == "EXONIC")

for (i in 1:nrow(CHR.COD.TAIL))
{
  CUR.CHR <- as.integer(strsplit(as.character(CHR.COD.CONSERVATIVE[i,"CHROMOSOME"]),"chr")[[1]][2])
  
  CUR.QTL <- subset(QTL.POSITIONS, Chromosome == CUR.CHR & Lower_Bound <= CHR.COD.CONSERVATIVE[i,"POSITION"] & Higher_Bound >= CHR.COD.CONSERVATIVE[i,"POSITION"])
  CHR.COD.CONSERVATIVE[i,"N.eQTL"] <- nrow(CUR.QTL)
}


N.MUT <- c(nrow(CHR.NEUTRAL),nrow(CHR.CAUSAL),nrow(CHR.RANDOM),nrow(CHR.TAIL),nrow(CHR.CONSERVATIVE))
N.NOT.QTL <- c(length(which(CHR.NEUTRAL$N.eQTL == 0)),length(which(CHR.CAUSAL$N.eQTL == 0)),length(which(CHR.RANDOM$N.eQTL == 0)),length(which(CHR.TAIL$N.eQTL == 0)),length(which(CHR.CONSERVATIVE$N.eQTL == 0)))
N.QTL <- N.MUT - N.NOT.QTL
N.CODING <- c(nrow(CHR.COD.NEUTRAL),nrow(CHR.COD.CAUSAL),nrow(CHR.COD.RANDOM),nrow(CHR.COD.TAIL),nrow(CHR.COD.CONSERVATIVE))
N.CODING.NOT.QTL <- c(length(which(CHR.COD.NEUTRAL$N.eQTL == 0)),length(which(CHR.COD.CAUSAL$N.eQTL == 0)),length(which(CHR.COD.RANDOM$N.eQTL == 0)),length(which(CHR.COD.TAIL$N.eQTL == 0)),length(which(CHR.COD.CONSERVATIVE$N.eQTL == 0)))
N.CODING.QTL <- N.CODING - N.CODING.NOT.QTL

FREQ.QTL <- round(N.QTL/N.MUT,3)
FREQ.NOT.QTL <- round(N.NOT.QTL/N.MUT,3)
FREQ.CODING.QTL <- round(N.CODING.QTL/N.CODING,3)
FREQ.CODING.NOT.QTL <- round(N.CODING.NOT.QTL/N.CODING,3)

STAT <- data.frame(MUT.TYPE, N.MUT, N.QTL, N.NOT.QTL, N.CODING, N.CODING.QTL, N.CODING.NOT.QTL, FREQ.QTL, FREQ.NOT.QTL, FREQ.CODING.QTL, FREQ.CODING.NOT.QTL)

G.MATRIX.NEUTRAL <- matrix(c(STAT[1,"N.QTL"],STAT[1,"N.NOT.QTL"],STAT[1,"N.QTL"],STAT[1,"N.NOT.QTL"]),nrow=2,ncol=2)
G.MATRIX.CAUSAL <- matrix(c(STAT[1,"N.QTL"],STAT[1,"N.NOT.QTL"],STAT[2,"N.QTL"],STAT[2,"N.NOT.QTL"]),nrow=2,ncol=2)
G.MATRIX.RANDOM <- matrix(c(STAT[1,"N.QTL"],STAT[1,"N.NOT.QTL"],STAT[3,"N.QTL"],STAT[3,"N.NOT.QTL"]),nrow=2,ncol=2)
G.MATRIX.TAIL <- matrix(c(STAT[1,"N.QTL"],STAT[1,"N.NOT.QTL"],STAT[4,"N.QTL"],STAT[4,"N.NOT.QTL"]),nrow=2,ncol=2)
G.MATRIX.CONSERVATIVE <- matrix(c(STAT[1,"N.QTL"],STAT[1,"N.NOT.QTL"],STAT[5,"N.QTL"],STAT[5,"N.NOT.QTL"]),nrow=2,ncol=2)

STAT[,"G.STAT.QTL"] <- c(likelihood.test(G.MATRIX.NEUTRAL,conservative=TRUE)$statistic,likelihood.test(G.MATRIX.CAUSAL,conservative=TRUE)$statistic,likelihood.test(G.MATRIX.RANDOM,conservative=TRUE)$statistic,likelihood.test(G.MATRIX.TAIL,conservative=TRUE)$statistic,likelihood.test(G.MATRIX.CONSERVATIVE,conservative=TRUE)$statistic)

STAT[,"P.VAL.QTL"] <- c(pchisq(STAT[1,"G.STAT.QTL"],df=1,lower.tail=FALSE),pchisq(STAT[2,"G.STAT.QTL"],df=1,lower.tail=FALSE),pchisq(STAT[3,"G.STAT.QTL"],df=1,lower.tail=FALSE),pchisq(STAT[4,"G.STAT.QTL"],df=1,lower.tail=FALSE),pchisq(STAT[5,"G.STAT.QTL"],df=1,lower.tail=FALSE))


G.MATRIX.NEUTRAL <- matrix(c(STAT[1,"N.CODING.QTL"],STAT[1,"N.CODING.NOT.QTL"],STAT[1,"N.CODING.QTL"],STAT[1,"N.CODING.NOT.QTL"]),nrow=2,ncol=2)
G.MATRIX.CAUSAL <- matrix(c(STAT[1,"N.CODING.QTL"],STAT[1,"N.CODING.NOT.QTL"],STAT[2,"N.CODING.QTL"],STAT[2,"N.CODING.NOT.QTL"]),nrow=2,ncol=2)
G.MATRIX.RANDOM <- matrix(c(STAT[1,"N.CODING.QTL"],STAT[1,"N.CODING.NOT.QTL"],STAT[3,"N.CODING.QTL"],STAT[3,"N.CODING.NOT.QTL"]),nrow=2,ncol=2)
G.MATRIX.TAIL <- matrix(c(STAT[1,"N.CODING.QTL"],STAT[1,"N.CODING.NOT.QTL"],STAT[4,"N.CODING.QTL"],STAT[4,"N.CODING.NOT.QTL"]),nrow=2,ncol=2)
G.MATRIX.CONSERVATIVE <- matrix(c(STAT[1,"N.CODING.QTL"],STAT[1,"N.CODING.NOT.QTL"],STAT[5,"N.CODING.QTL"],STAT[5,"N.CODING.NOT.QTL"]),nrow=2,ncol=2)

STAT[,"G.STAT.CODING.QTL"] <- c(likelihood.test(G.MATRIX.NEUTRAL,conservative=TRUE)$statistic,likelihood.test(G.MATRIX.CAUSAL,conservative=TRUE)$statistic,likelihood.test(G.MATRIX.RANDOM,conservative=TRUE)$statistic,likelihood.test(G.MATRIX.TAIL,conservative=TRUE)$statistic,likelihood.test(G.MATRIX.CONSERVATIVE,conservative=TRUE)$statistic)

STAT[,"P.VAL.CODING.QTL"] <- c(pchisq(STAT[1,"G.STAT.CODING.QTL"],df=1,lower.tail=FALSE),pchisq(STAT[2,"G.STAT.CODING.QTL"],df=1,lower.tail=FALSE),pchisq(STAT[3,"G.STAT.CODING.QTL"],df=1,lower.tail=FALSE),pchisq(STAT[4,"G.STAT.CODING.QTL"],df=1,lower.tail=FALSE),pchisq(STAT[5,"G.STAT.CODING.QTL"],df=1,lower.tail=FALSE))

write.table(STAT,"Statistics/eQTL/STAT.YPS1000.QTL.txt",sep="\t",row.names=FALSE)


###c) Analyses including only eQTL regions from SK1xBY cross.###

#QTL.Positions.txt is a dataset from Metzger and Wittkopp (2019) describing the genomic positions of all eQTL regions identified in 3 crosses.
#QTL.Positions.txt can be found in SupplementaryFile11.zip.
QTL.POSITIONS <- read.table("QTL.Positions.txt",header=TRUE)

QTL.POSITIONS <- subset(QTL.POSITIONS, Cross == "SK1xBY")

CHR.NEUTRAL <- subset(SUB.NEUTRAL, CHROMOSOME %nin% c("chrMito","chrYFP"))

for (i in 1:nrow(CHR.NEUTRAL))
{
  CUR.CHR <- as.integer(strsplit(as.character(CHR.NEUTRAL[i,"CHROMOSOME"]),"chr")[[1]][2])
  
  CUR.QTL <- subset(QTL.POSITIONS, Chromosome == CUR.CHR & Lower_Bound <= CHR.NEUTRAL[i,"POSITION"] & Higher_Bound >= CHR.NEUTRAL[i,"POSITION"])
  CHR.NEUTRAL[i,"N.eQTL"] <- nrow(CUR.QTL)
}

CHR.CAUSAL <- rbind(RANDOM,TAIL)

for (i in 1:nrow(CHR.CAUSAL))
{
  CUR.CHR <- as.integer(strsplit(as.character(CHR.CAUSAL[i,"CHROMOSOME"]),"chr")[[1]][2])
  
  CUR.QTL <- subset(QTL.POSITIONS, Chromosome == CUR.CHR & Lower_Bound <= CHR.CAUSAL[i,"POSITION"] & Higher_Bound >= CHR.CAUSAL[i,"POSITION"])
  CHR.CAUSAL[i,"N.eQTL"] <- nrow(CUR.QTL)
}

CHR.RANDOM <- subset(RANDOM, CHROMOSOME %nin% c("chrMito","chrYFP"))

for (i in 1:nrow(CHR.RANDOM))
{
  CUR.CHR <- as.integer(strsplit(as.character(CHR.RANDOM[i,"CHROMOSOME"]),"chr")[[1]][2])
  
  CUR.QTL <- subset(QTL.POSITIONS, Chromosome == CUR.CHR & Lower_Bound <= CHR.RANDOM[i,"POSITION"] & Higher_Bound >= CHR.RANDOM[i,"POSITION"])
  CHR.RANDOM[i,"N.eQTL"] <- nrow(CUR.QTL)
}

CHR.TAIL <- subset(TAIL, CHROMOSOME %nin% c("chrMito","chrYFP"))

for (i in 1:nrow(CHR.TAIL))
{
  CUR.CHR <- as.integer(strsplit(as.character(CHR.TAIL[i,"CHROMOSOME"]),"chr")[[1]][2])
  
  CUR.QTL <- subset(QTL.POSITIONS, Chromosome == CUR.CHR & Lower_Bound <= CHR.TAIL[i,"POSITION"] & Higher_Bound >= CHR.TAIL[i,"POSITION"])
  CHR.TAIL[i,"N.eQTL"] <- nrow(CUR.QTL)
}

CHR.CONSERVATIVE <- subset(CONSERVATIVE, CHROMOSOME %nin% c("chrMito","chrYFP"))

for (i in 1:nrow(CHR.CONSERVATIVE))
{
  CUR.CHR <- as.integer(strsplit(as.character(CHR.CONSERVATIVE[i,"CHROMOSOME"]),"chr")[[1]][2])
  
  CUR.QTL <- subset(QTL.POSITIONS, Chromosome == CUR.CHR & Lower_Bound <= CHR.CONSERVATIVE[i,"POSITION"] & Higher_Bound >= CHR.CONSERVATIVE[i,"POSITION"])
  CHR.CONSERVATIVE[i,"N.eQTL"] <- nrow(CUR.QTL)
}

CHR.COD.NEUTRAL <- subset(SUBSTITUTION, CHROMOSOME %nin% c("chrMito","chrYFP") & CLASS == "EXONIC")

for (i in 1:nrow(CHR.COD.NEUTRAL))
{
  CUR.CHR <- as.integer(strsplit(as.character(CHR.COD.NEUTRAL[i,"CHROMOSOME"]),"chr")[[1]][2])
  
  CUR.QTL <- subset(QTL.POSITIONS, Chromosome == CUR.CHR & Lower_Bound <= CHR.COD.NEUTRAL[i,"POSITION"] & Higher_Bound >= CHR.COD.NEUTRAL[i,"POSITION"])
  CHR.COD.NEUTRAL[i,"N.eQTL"] <- nrow(CUR.QTL)
}

CHR.COD.CAUSAL <- subset(CAUSAL, CHROMOSOME %nin% c("chrMito","chrYFP") & CLASS == "EXONIC")

for (i in 1:nrow(CHR.COD.CAUSAL))
{
  CUR.CHR <- as.integer(strsplit(as.character(CHR.COD.CAUSAL[i,"CHROMOSOME"]),"chr")[[1]][2])
  
  CUR.QTL <- subset(QTL.POSITIONS, Chromosome == CUR.CHR & Lower_Bound <= CHR.COD.CAUSAL[i,"POSITION"] & Higher_Bound >= CHR.COD.CAUSAL[i,"POSITION"])
  CHR.COD.CAUSAL[i,"N.eQTL"] <- nrow(CUR.QTL)
}

CHR.COD.RANDOM <- subset(RANDOM, CHROMOSOME %nin% c("chrMito","chrYFP") & CLASS == "EXONIC")

for (i in 1:nrow(CHR.COD.RANDOM))
{
  CUR.CHR <- as.integer(strsplit(as.character(CHR.COD.RANDOM[i,"CHROMOSOME"]),"chr")[[1]][2])
  
  CUR.QTL <- subset(QTL.POSITIONS, Chromosome == CUR.CHR & Lower_Bound <= CHR.COD.RANDOM[i,"POSITION"] & Higher_Bound >= CHR.COD.RANDOM[i,"POSITION"])
  CHR.COD.RANDOM[i,"N.eQTL"] <- nrow(CUR.QTL)
}

CHR.COD.TAIL <- subset(TAIL, CHROMOSOME %nin% c("chrMito","chrYFP") & CLASS == "EXONIC")

for (i in 1:nrow(CHR.COD.TAIL))
{
  CUR.CHR <- as.integer(strsplit(as.character(CHR.COD.TAIL[i,"CHROMOSOME"]),"chr")[[1]][2])
  
  CUR.QTL <- subset(QTL.POSITIONS, Chromosome == CUR.CHR & Lower_Bound <= CHR.COD.TAIL[i,"POSITION"] & Higher_Bound >= CHR.COD.TAIL[i,"POSITION"])
  CHR.COD.TAIL[i,"N.eQTL"] <- nrow(CUR.QTL)
}

CHR.COD.CONSERVATIVE <- subset(CONSERVATIVE, CHROMOSOME %nin% c("chrMito","chrYFP") & CLASS == "EXONIC")

for (i in 1:nrow(CHR.COD.TAIL))
{
  CUR.CHR <- as.integer(strsplit(as.character(CHR.COD.CONSERVATIVE[i,"CHROMOSOME"]),"chr")[[1]][2])
  
  CUR.QTL <- subset(QTL.POSITIONS, Chromosome == CUR.CHR & Lower_Bound <= CHR.COD.CONSERVATIVE[i,"POSITION"] & Higher_Bound >= CHR.COD.CONSERVATIVE[i,"POSITION"])
  CHR.COD.CONSERVATIVE[i,"N.eQTL"] <- nrow(CUR.QTL)
}


N.MUT <- c(nrow(CHR.NEUTRAL),nrow(CHR.CAUSAL),nrow(CHR.RANDOM),nrow(CHR.TAIL),nrow(CHR.CONSERVATIVE))
N.NOT.QTL <- c(length(which(CHR.NEUTRAL$N.eQTL == 0)),length(which(CHR.CAUSAL$N.eQTL == 0)),length(which(CHR.RANDOM$N.eQTL == 0)),length(which(CHR.TAIL$N.eQTL == 0)),length(which(CHR.CONSERVATIVE$N.eQTL == 0)))
N.QTL <- N.MUT - N.NOT.QTL
N.CODING <- c(nrow(CHR.COD.NEUTRAL),nrow(CHR.COD.CAUSAL),nrow(CHR.COD.RANDOM),nrow(CHR.COD.TAIL),nrow(CHR.COD.CONSERVATIVE))
N.CODING.NOT.QTL <- c(length(which(CHR.COD.NEUTRAL$N.eQTL == 0)),length(which(CHR.COD.CAUSAL$N.eQTL == 0)),length(which(CHR.COD.RANDOM$N.eQTL == 0)),length(which(CHR.COD.TAIL$N.eQTL == 0)),length(which(CHR.COD.CONSERVATIVE$N.eQTL == 0)))
N.CODING.QTL <- N.CODING - N.CODING.NOT.QTL

FREQ.QTL <- round(N.QTL/N.MUT,3)
FREQ.NOT.QTL <- round(N.NOT.QTL/N.MUT,3)
FREQ.CODING.QTL <- round(N.CODING.QTL/N.CODING,3)
FREQ.CODING.NOT.QTL <- round(N.CODING.NOT.QTL/N.CODING,3)

STAT <- data.frame(MUT.TYPE, N.MUT, N.QTL, N.NOT.QTL, N.CODING, N.CODING.QTL, N.CODING.NOT.QTL, FREQ.QTL, FREQ.NOT.QTL, FREQ.CODING.QTL, FREQ.CODING.NOT.QTL)

G.MATRIX.NEUTRAL <- matrix(c(STAT[1,"N.QTL"],STAT[1,"N.NOT.QTL"],STAT[1,"N.QTL"],STAT[1,"N.NOT.QTL"]),nrow=2,ncol=2)
G.MATRIX.CAUSAL <- matrix(c(STAT[1,"N.QTL"],STAT[1,"N.NOT.QTL"],STAT[2,"N.QTL"],STAT[2,"N.NOT.QTL"]),nrow=2,ncol=2)
G.MATRIX.RANDOM <- matrix(c(STAT[1,"N.QTL"],STAT[1,"N.NOT.QTL"],STAT[3,"N.QTL"],STAT[3,"N.NOT.QTL"]),nrow=2,ncol=2)
G.MATRIX.TAIL <- matrix(c(STAT[1,"N.QTL"],STAT[1,"N.NOT.QTL"],STAT[4,"N.QTL"],STAT[4,"N.NOT.QTL"]),nrow=2,ncol=2)
G.MATRIX.CONSERVATIVE <- matrix(c(STAT[1,"N.QTL"],STAT[1,"N.NOT.QTL"],STAT[5,"N.QTL"],STAT[5,"N.NOT.QTL"]),nrow=2,ncol=2)

STAT[,"G.STAT.QTL"] <- c(likelihood.test(G.MATRIX.NEUTRAL,conservative=TRUE)$statistic,likelihood.test(G.MATRIX.CAUSAL,conservative=TRUE)$statistic,likelihood.test(G.MATRIX.RANDOM,conservative=TRUE)$statistic,likelihood.test(G.MATRIX.TAIL,conservative=TRUE)$statistic,likelihood.test(G.MATRIX.CONSERVATIVE,conservative=TRUE)$statistic)

STAT[,"P.VAL.QTL"] <- c(pchisq(STAT[1,"G.STAT.QTL"],df=1,lower.tail=FALSE),pchisq(STAT[2,"G.STAT.QTL"],df=1,lower.tail=FALSE),pchisq(STAT[3,"G.STAT.QTL"],df=1,lower.tail=FALSE),pchisq(STAT[4,"G.STAT.QTL"],df=1,lower.tail=FALSE),pchisq(STAT[5,"G.STAT.QTL"],df=1,lower.tail=FALSE))


G.MATRIX.NEUTRAL <- matrix(c(STAT[1,"N.CODING.QTL"],STAT[1,"N.CODING.NOT.QTL"],STAT[1,"N.CODING.QTL"],STAT[1,"N.CODING.NOT.QTL"]),nrow=2,ncol=2)
G.MATRIX.CAUSAL <- matrix(c(STAT[1,"N.CODING.QTL"],STAT[1,"N.CODING.NOT.QTL"],STAT[2,"N.CODING.QTL"],STAT[2,"N.CODING.NOT.QTL"]),nrow=2,ncol=2)
G.MATRIX.RANDOM <- matrix(c(STAT[1,"N.CODING.QTL"],STAT[1,"N.CODING.NOT.QTL"],STAT[3,"N.CODING.QTL"],STAT[3,"N.CODING.NOT.QTL"]),nrow=2,ncol=2)
G.MATRIX.TAIL <- matrix(c(STAT[1,"N.CODING.QTL"],STAT[1,"N.CODING.NOT.QTL"],STAT[4,"N.CODING.QTL"],STAT[4,"N.CODING.NOT.QTL"]),nrow=2,ncol=2)
G.MATRIX.CONSERVATIVE <- matrix(c(STAT[1,"N.CODING.QTL"],STAT[1,"N.CODING.NOT.QTL"],STAT[5,"N.CODING.QTL"],STAT[5,"N.CODING.NOT.QTL"]),nrow=2,ncol=2)

STAT[,"G.STAT.CODING.QTL"] <- c(likelihood.test(G.MATRIX.NEUTRAL,conservative=TRUE)$statistic,likelihood.test(G.MATRIX.CAUSAL,conservative=TRUE)$statistic,likelihood.test(G.MATRIX.RANDOM,conservative=TRUE)$statistic,likelihood.test(G.MATRIX.TAIL,conservative=TRUE)$statistic,likelihood.test(G.MATRIX.CONSERVATIVE,conservative=TRUE)$statistic)

STAT[,"P.VAL.CODING.QTL"] <- c(pchisq(STAT[1,"G.STAT.CODING.QTL"],df=1,lower.tail=FALSE),pchisq(STAT[2,"G.STAT.CODING.QTL"],df=1,lower.tail=FALSE),pchisq(STAT[3,"G.STAT.CODING.QTL"],df=1,lower.tail=FALSE),pchisq(STAT[4,"G.STAT.CODING.QTL"],df=1,lower.tail=FALSE),pchisq(STAT[5,"G.STAT.CODING.QTL"],df=1,lower.tail=FALSE))

write.table(STAT,"Statistics/eQTL/STAT.SK1.QTL.txt",sep="\t",row.names=FALSE)


###d) Analyses including only eQTL regions from M22xBY cross.###

#QTL.Positions.txt is a dataset from Metzger and Wittkopp (2019) describing the genomic positions of all eQTL regions identified in 3 crosses.
#QTL.Positions.txt can be found in SupplementaryFile11.zip.
QTL.POSITIONS <- read.table("QTL.Positions.txt",header=TRUE)

QTL.POSITIONS <- subset(QTL.POSITIONS, Cross == "M22xBY")

CHR.NEUTRAL <- subset(SUB.NEUTRAL, CHROMOSOME %nin% c("chrMito","chrYFP"))

for (i in 1:nrow(CHR.NEUTRAL))
{
  CUR.CHR <- as.integer(strsplit(as.character(CHR.NEUTRAL[i,"CHROMOSOME"]),"chr")[[1]][2])
  
  CUR.QTL <- subset(QTL.POSITIONS, Chromosome == CUR.CHR & Lower_Bound <= CHR.NEUTRAL[i,"POSITION"] & Higher_Bound >= CHR.NEUTRAL[i,"POSITION"])
  CHR.NEUTRAL[i,"N.eQTL"] <- nrow(CUR.QTL)
}

CHR.CAUSAL <- rbind(RANDOM,TAIL)

for (i in 1:nrow(CHR.CAUSAL))
{
  CUR.CHR <- as.integer(strsplit(as.character(CHR.CAUSAL[i,"CHROMOSOME"]),"chr")[[1]][2])
  
  CUR.QTL <- subset(QTL.POSITIONS, Chromosome == CUR.CHR & Lower_Bound <= CHR.CAUSAL[i,"POSITION"] & Higher_Bound >= CHR.CAUSAL[i,"POSITION"])
  CHR.CAUSAL[i,"N.eQTL"] <- nrow(CUR.QTL)
}

CHR.RANDOM <- subset(RANDOM, CHROMOSOME %nin% c("chrMito","chrYFP"))

for (i in 1:nrow(CHR.RANDOM))
{
  CUR.CHR <- as.integer(strsplit(as.character(CHR.RANDOM[i,"CHROMOSOME"]),"chr")[[1]][2])
  
  CUR.QTL <- subset(QTL.POSITIONS, Chromosome == CUR.CHR & Lower_Bound <= CHR.RANDOM[i,"POSITION"] & Higher_Bound >= CHR.RANDOM[i,"POSITION"])
  CHR.RANDOM[i,"N.eQTL"] <- nrow(CUR.QTL)
}

CHR.TAIL <- subset(TAIL, CHROMOSOME %nin% c("chrMito","chrYFP"))

for (i in 1:nrow(CHR.TAIL))
{
  CUR.CHR <- as.integer(strsplit(as.character(CHR.TAIL[i,"CHROMOSOME"]),"chr")[[1]][2])
  
  CUR.QTL <- subset(QTL.POSITIONS, Chromosome == CUR.CHR & Lower_Bound <= CHR.TAIL[i,"POSITION"] & Higher_Bound >= CHR.TAIL[i,"POSITION"])
  CHR.TAIL[i,"N.eQTL"] <- nrow(CUR.QTL)
}

CHR.CONSERVATIVE <- subset(CONSERVATIVE, CHROMOSOME %nin% c("chrMito","chrYFP"))

for (i in 1:nrow(CHR.CONSERVATIVE))
{
  CUR.CHR <- as.integer(strsplit(as.character(CHR.CONSERVATIVE[i,"CHROMOSOME"]),"chr")[[1]][2])
  
  CUR.QTL <- subset(QTL.POSITIONS, Chromosome == CUR.CHR & Lower_Bound <= CHR.CONSERVATIVE[i,"POSITION"] & Higher_Bound >= CHR.CONSERVATIVE[i,"POSITION"])
  CHR.CONSERVATIVE[i,"N.eQTL"] <- nrow(CUR.QTL)
}

CHR.COD.NEUTRAL <- subset(SUBSTITUTION, CHROMOSOME %nin% c("chrMito","chrYFP") & CLASS == "EXONIC")

for (i in 1:nrow(CHR.COD.NEUTRAL))
{
  CUR.CHR <- as.integer(strsplit(as.character(CHR.COD.NEUTRAL[i,"CHROMOSOME"]),"chr")[[1]][2])
  
  CUR.QTL <- subset(QTL.POSITIONS, Chromosome == CUR.CHR & Lower_Bound <= CHR.COD.NEUTRAL[i,"POSITION"] & Higher_Bound >= CHR.COD.NEUTRAL[i,"POSITION"])
  CHR.COD.NEUTRAL[i,"N.eQTL"] <- nrow(CUR.QTL)
}

CHR.COD.CAUSAL <- subset(CAUSAL, CHROMOSOME %nin% c("chrMito","chrYFP") & CLASS == "EXONIC")

for (i in 1:nrow(CHR.COD.CAUSAL))
{
  CUR.CHR <- as.integer(strsplit(as.character(CHR.COD.CAUSAL[i,"CHROMOSOME"]),"chr")[[1]][2])
  
  CUR.QTL <- subset(QTL.POSITIONS, Chromosome == CUR.CHR & Lower_Bound <= CHR.COD.CAUSAL[i,"POSITION"] & Higher_Bound >= CHR.COD.CAUSAL[i,"POSITION"])
  CHR.COD.CAUSAL[i,"N.eQTL"] <- nrow(CUR.QTL)
}

CHR.COD.RANDOM <- subset(RANDOM, CHROMOSOME %nin% c("chrMito","chrYFP") & CLASS == "EXONIC")

for (i in 1:nrow(CHR.COD.RANDOM))
{
  CUR.CHR <- as.integer(strsplit(as.character(CHR.COD.RANDOM[i,"CHROMOSOME"]),"chr")[[1]][2])
  
  CUR.QTL <- subset(QTL.POSITIONS, Chromosome == CUR.CHR & Lower_Bound <= CHR.COD.RANDOM[i,"POSITION"] & Higher_Bound >= CHR.COD.RANDOM[i,"POSITION"])
  CHR.COD.RANDOM[i,"N.eQTL"] <- nrow(CUR.QTL)
}

CHR.COD.TAIL <- subset(TAIL, CHROMOSOME %nin% c("chrMito","chrYFP") & CLASS == "EXONIC")

for (i in 1:nrow(CHR.COD.TAIL))
{
  CUR.CHR <- as.integer(strsplit(as.character(CHR.COD.TAIL[i,"CHROMOSOME"]),"chr")[[1]][2])
  
  CUR.QTL <- subset(QTL.POSITIONS, Chromosome == CUR.CHR & Lower_Bound <= CHR.COD.TAIL[i,"POSITION"] & Higher_Bound >= CHR.COD.TAIL[i,"POSITION"])
  CHR.COD.TAIL[i,"N.eQTL"] <- nrow(CUR.QTL)
}

CHR.COD.CONSERVATIVE <- subset(CONSERVATIVE, CHROMOSOME %nin% c("chrMito","chrYFP") & CLASS == "EXONIC")

for (i in 1:nrow(CHR.COD.TAIL))
{
  CUR.CHR <- as.integer(strsplit(as.character(CHR.COD.CONSERVATIVE[i,"CHROMOSOME"]),"chr")[[1]][2])
  
  CUR.QTL <- subset(QTL.POSITIONS, Chromosome == CUR.CHR & Lower_Bound <= CHR.COD.CONSERVATIVE[i,"POSITION"] & Higher_Bound >= CHR.COD.CONSERVATIVE[i,"POSITION"])
  CHR.COD.CONSERVATIVE[i,"N.eQTL"] <- nrow(CUR.QTL)
}


N.MUT <- c(nrow(CHR.NEUTRAL),nrow(CHR.CAUSAL),nrow(CHR.RANDOM),nrow(CHR.TAIL),nrow(CHR.CONSERVATIVE))
N.NOT.QTL <- c(length(which(CHR.NEUTRAL$N.eQTL == 0)),length(which(CHR.CAUSAL$N.eQTL == 0)),length(which(CHR.RANDOM$N.eQTL == 0)),length(which(CHR.TAIL$N.eQTL == 0)),length(which(CHR.CONSERVATIVE$N.eQTL == 0)))
N.QTL <- N.MUT - N.NOT.QTL
N.CODING <- c(nrow(CHR.COD.NEUTRAL),nrow(CHR.COD.CAUSAL),nrow(CHR.COD.RANDOM),nrow(CHR.COD.TAIL),nrow(CHR.COD.CONSERVATIVE))
N.CODING.NOT.QTL <- c(length(which(CHR.COD.NEUTRAL$N.eQTL == 0)),length(which(CHR.COD.CAUSAL$N.eQTL == 0)),length(which(CHR.COD.RANDOM$N.eQTL == 0)),length(which(CHR.COD.TAIL$N.eQTL == 0)),length(which(CHR.COD.CONSERVATIVE$N.eQTL == 0)))
N.CODING.QTL <- N.CODING - N.CODING.NOT.QTL

FREQ.QTL <- round(N.QTL/N.MUT,3)
FREQ.NOT.QTL <- round(N.NOT.QTL/N.MUT,3)
FREQ.CODING.QTL <- round(N.CODING.QTL/N.CODING,3)
FREQ.CODING.NOT.QTL <- round(N.CODING.NOT.QTL/N.CODING,3)

STAT <- data.frame(MUT.TYPE, N.MUT, N.QTL, N.NOT.QTL, N.CODING, N.CODING.QTL, N.CODING.NOT.QTL, FREQ.QTL, FREQ.NOT.QTL, FREQ.CODING.QTL, FREQ.CODING.NOT.QTL)

G.MATRIX.NEUTRAL <- matrix(c(STAT[1,"N.QTL"],STAT[1,"N.NOT.QTL"],STAT[1,"N.QTL"],STAT[1,"N.NOT.QTL"]),nrow=2,ncol=2)
G.MATRIX.CAUSAL <- matrix(c(STAT[1,"N.QTL"],STAT[1,"N.NOT.QTL"],STAT[2,"N.QTL"],STAT[2,"N.NOT.QTL"]),nrow=2,ncol=2)
G.MATRIX.RANDOM <- matrix(c(STAT[1,"N.QTL"],STAT[1,"N.NOT.QTL"],STAT[3,"N.QTL"],STAT[3,"N.NOT.QTL"]),nrow=2,ncol=2)
G.MATRIX.TAIL <- matrix(c(STAT[1,"N.QTL"],STAT[1,"N.NOT.QTL"],STAT[4,"N.QTL"],STAT[4,"N.NOT.QTL"]),nrow=2,ncol=2)
G.MATRIX.CONSERVATIVE <- matrix(c(STAT[1,"N.QTL"],STAT[1,"N.NOT.QTL"],STAT[5,"N.QTL"],STAT[5,"N.NOT.QTL"]),nrow=2,ncol=2)

STAT[,"G.STAT.QTL"] <- c(likelihood.test(G.MATRIX.NEUTRAL,conservative=TRUE)$statistic,likelihood.test(G.MATRIX.CAUSAL,conservative=TRUE)$statistic,likelihood.test(G.MATRIX.RANDOM,conservative=TRUE)$statistic,likelihood.test(G.MATRIX.TAIL,conservative=TRUE)$statistic,likelihood.test(G.MATRIX.CONSERVATIVE,conservative=TRUE)$statistic)

STAT[,"P.VAL.QTL"] <- c(pchisq(STAT[1,"G.STAT.QTL"],df=1,lower.tail=FALSE),pchisq(STAT[2,"G.STAT.QTL"],df=1,lower.tail=FALSE),pchisq(STAT[3,"G.STAT.QTL"],df=1,lower.tail=FALSE),pchisq(STAT[4,"G.STAT.QTL"],df=1,lower.tail=FALSE),pchisq(STAT[5,"G.STAT.QTL"],df=1,lower.tail=FALSE))


G.MATRIX.NEUTRAL <- matrix(c(STAT[1,"N.CODING.QTL"],STAT[1,"N.CODING.NOT.QTL"],STAT[1,"N.CODING.QTL"],STAT[1,"N.CODING.NOT.QTL"]),nrow=2,ncol=2)
G.MATRIX.CAUSAL <- matrix(c(STAT[1,"N.CODING.QTL"],STAT[1,"N.CODING.NOT.QTL"],STAT[2,"N.CODING.QTL"],STAT[2,"N.CODING.NOT.QTL"]),nrow=2,ncol=2)
G.MATRIX.RANDOM <- matrix(c(STAT[1,"N.CODING.QTL"],STAT[1,"N.CODING.NOT.QTL"],STAT[3,"N.CODING.QTL"],STAT[3,"N.CODING.NOT.QTL"]),nrow=2,ncol=2)
G.MATRIX.TAIL <- matrix(c(STAT[1,"N.CODING.QTL"],STAT[1,"N.CODING.NOT.QTL"],STAT[4,"N.CODING.QTL"],STAT[4,"N.CODING.NOT.QTL"]),nrow=2,ncol=2)
G.MATRIX.CONSERVATIVE <- matrix(c(STAT[1,"N.CODING.QTL"],STAT[1,"N.CODING.NOT.QTL"],STAT[5,"N.CODING.QTL"],STAT[5,"N.CODING.NOT.QTL"]),nrow=2,ncol=2)

STAT[,"G.STAT.CODING.QTL"] <- c(likelihood.test(G.MATRIX.NEUTRAL,conservative=TRUE)$statistic,likelihood.test(G.MATRIX.CAUSAL,conservative=TRUE)$statistic,likelihood.test(G.MATRIX.RANDOM,conservative=TRUE)$statistic,likelihood.test(G.MATRIX.TAIL,conservative=TRUE)$statistic,likelihood.test(G.MATRIX.CONSERVATIVE,conservative=TRUE)$statistic)

STAT[,"P.VAL.CODING.QTL"] <- c(pchisq(STAT[1,"G.STAT.CODING.QTL"],df=1,lower.tail=FALSE),pchisq(STAT[2,"G.STAT.CODING.QTL"],df=1,lower.tail=FALSE),pchisq(STAT[3,"G.STAT.CODING.QTL"],df=1,lower.tail=FALSE),pchisq(STAT[4,"G.STAT.CODING.QTL"],df=1,lower.tail=FALSE),pchisq(STAT[5,"G.STAT.CODING.QTL"],df=1,lower.tail=FALSE))

write.table(STAT,"Statistics/eQTL/STAT.M22.QTL.txt",sep="\t",row.names=FALSE)
```
