## Supplementary Files 1-19 for "Mutational sources of *trans-*regulatory variation affecting gene expression in *Saccharomyces cerevisiae*": SupplementaryFile17.R

```
##############################################################################################
################################## Supplementary File 17 #####################################
##############################################################################################
#             R script used to annotate variants identified in BSA-Seq data.                 #
##############################################################################################

#Annotations of all mutations identified in BSA-Seq data generated with this script are included in SupplementaryFile2.txt.

###Clean up memory###
rm(list=ls())

###Set directory###
setwd("/Path.to.input.files")

###Load libraries###
library(data.table)

###Function returning complementary bases###
REV.COMP <- function(x) {
		
	if (x == "A") {
		RC <- "T"
	} else if (x == "C") {
		RC <- "G"
	} else if (x == "G") {
		RC <- "C"
	} else if (x == "T") {
		RC <- "A"
	}
	
	return(RC)
}

#################################################################################
###OPEN GENOME SEQUENCE, GENOME ANNOTATIONS, GENETIC CODE AND INTRON POSITIONS###
#################################################################################


#"S288c.genome.bed" is a table listing all nucleotides found at each position of each chromosome in the yeast genome.
#The first 10000 lines of this file ("S288c.genome.truncated.bed") can be found in SupplementaryFile11.zip.
#"S288c.genome.bed" was generated from the reference genome ("S288c.mapping.fsa") included in SupplementaryFile11.zip.
SEQ <- read.table("S288c.genome.bed",header=TRUE,colClasses=c("factor","integer","character"))
SEQ <- as.data.table(SEQ)

#"Gene.List.txt" is a table indicating the location of 5743 ORF in the yeast genome. 
#It was made from "orf_coding.fasta.gz" file available on https://www.yeastgenome.org/ (S288c genome release R64-1-1). 
#"Gene.List.txt" can be found in SupplementaryFile11.zip.
GENES <- read.table("Gene.List.txt",header=TRUE)
GENES <- GENES[,1:10]

#"Genetic.Code.txt" is a table containing the genetic code and it can be found in SupplementaryFile11.zip.
CODE <- read.table("Genetic.Code.txt",header=TRUE)

#"Introns.txt" is a table containing the location of well-characterized introns in S288c genome.
INTRONS <- read.table("Introns.txt",header=TRUE)

#List of file(s) containing the variants to be annotated. These files are generated with the R script in SupplementaryFile16.R. 
VAR.PATH <- c("SNP.FILTER.txt")

######################
#REFORMAT INTRON FILE#
######################

INTRONS[,"LENGTH"] <- nchar(as.character(INTRONS[,"SEQ"]))
INTRONS[,"DIFF"] <- INTRONS[,"END"] - INTRONS[,"START"]
INTRONS[,"NAME"] <- as.character(INTRONS[,"NAME"])
INTRONS.NAME <- strsplit(INTRONS[,"NAME"],"_")

for (i in 1:length(INTRONS.NAME))
{
	INTRONS[i,"NAME.LENGTH"] <- abs(as.integer(INTRONS.NAME[[i]][3])-as.integer(INTRONS.NAME[[i]][4]))
	INTRONS[i,"START.NAME"] <- min(as.integer(INTRONS.NAME[[i]][3:4]))
	INTRONS[i,"END.NAME"] <- max(as.integer(INTRONS.NAME[[i]][3:4]))
}

##########################
###CALL VARIANT CLASSES###
##########################

for (i in 1:length(VAR.PATH)) {
	
	VARIANTS <- read.table(VAR.PATH[i],header=TRUE)
#	VARIANTS <- VARIANTS[,c(1:10,14)]
	VARIANTS <- VARIANTS[complete.cases(VARIANTS),]
	VARIANTS[,"REF"] <- as.character(VARIANTS[,"REF"])
	VARIANTS[,"ALT"] <- as.character(VARIANTS[,"ALT"])

	VARIANTS[,"CLASS"] <- NA
	VARIANTS[,"TYPE"] <- NA
	VARIANTS[,"ALERT"] <- NA
	VARIANTS[,"GENE.UPSTREAM"] <- NA
	VARIANTS[,"GENE.DOWNSTREAM"] <- NA
	VARIANTS[,"GENE.SHORT.UP"] <- NA
	VARIANTS[,"GENE.SHORT.DOWN"] <- NA
	VARIANTS[,"LOCATION.UPSTREAM"] <- NA	
	VARIANTS[,"LOCATION.DOWNSTREAM"] <- NA
	VARIANTS[,"GENE.FOCAL"] <- NA
	VARIANTS[,"GENE.SHORT"] <- NA
	VARIANTS[,"ORF.LENGTH"] <- NA
	VARIANTS[,"ORF.POSITION"] <- NA
	VARIANTS[,"AA.POSITION"] <- NA
	VARIANTS[,"PROT.LENGTH"] <- NA
	VARIANTS[,"REF.CODON"] <- NA
	VARIANTS[,"ALT.CODON"] <- NA
	VARIANTS[,"REF.AA"] <- NA
	VARIANTS[,"ALT.AA"] <- NA


	for (j in 1:nrow(VARIANTS)) {
		
		##########################################################################
		#DETERMINE CLASS OF VARIANTS (INTERGENIC, INTRONIC, EXONIC OR MULTIGENIC)#
		##########################################################################

		CUR.CHR <- VARIANTS[j,"CHROMOSOME"]
		CUR.POS <- VARIANTS[j,"POSITION"]	

		GENE.CHR <- subset(GENES, CHR == as.character(CUR.CHR))	
		
		GOOD.GENE <- GENE.CHR[which(GENE.CHR$START < CUR.POS & GENE.CHR$STOP > CUR.POS),]
		VARIANTS[j,"ALERT"] <- "OK"		

		if (nrow(GOOD.GENE) == 0) {
			VARIANTS[j,"CLASS"] <- "INTERGENIC"
		} else if (nrow(GOOD.GENE) == 1) {
			CUR.INTRON <- which(INTRONS$GENE == as.character(GOOD.GENE$GENE[1]))

			if (length(CUR.INTRON) == 0) {
				VARIANTS[j,"CLASS"] <- "EXONIC" 
			} else {		
				if (length(CUR.INTRON)>1) {
					VARIANTS[j,"CLASS"] <- "CHECK"
					VARIANTS[j,"ALERT"] <- "MULTIPLE.INTRONS"
				} else if (VARIANTS[j,"POSITION"] > INTRONS[CUR.INTRON,"START.NAME"] & VARIANTS[j,"POSITION"] < INTRONS[CUR.INTRON,"END.NAME"]) {
					VARIANTS[j,"CLASS"] <- "INTRONIC"
				} else {
					VARIANTS[j,"CLASS"] <- "EXONIC"
					VARIANTS[j,"ALERT"] <- "INTRON"
				}
			}
		} else if (nrow(GOOD.GENE) > 1) {
			VARIANTS[j,"CLASS"] <- "MULTIGENIC"	
		}

		##################################
		#CHARACTERIZE INTERGENIC VARIANTS#
		##################################

		if (VARIANTS[j,"CLASS"] == "INTERGENIC") {
			DIST <- VARIANTS[j,"POSITION"] - GENE.CHR$STOP
			GOOD.DIST <- min(DIST[which(DIST > 0)])
			UPSTREAM <- GENE.CHR[which(DIST == GOOD.DIST),]

			DIST <- GENE.CHR$START - VARIANTS[j,"POSITION"]
			GOOD.DIST <- min(DIST[which(DIST > 0)])
			DOWNSTREAM <- GENE.CHR[which(DIST == GOOD.DIST),]

			VARIANTS[j,"GENE.DOWNSTREAM"] <- as.character(DOWNSTREAM[1,"GENE"])
			VARIANTS[j,"GENE.UPSTREAM"] <- as.character(UPSTREAM[1,"GENE"])
			VARIANTS[j,"GENE.SHORT.DOWN"] <- as.character(DOWNSTREAM[1,"SHORT"])
			VARIANTS[j,"GENE.SHORT.UP"] <- as.character(UPSTREAM[1,"SHORT"])
			
			if (!is.na(UPSTREAM[1,"DIRECTION"]) & UPSTREAM[1,"DIRECTION"] == "+") {
				VARIANTS[j,"LOCATION.UPSTREAM"] <- "TERMINATOR"
			} else if (!is.na(UPSTREAM[1,"DIRECTION"])) {
				VARIANTS[j,"LOCATION.UPSTREAM"] <- "PROMOTER"
			} else {VARIANTS[j,"LOCATION.UPSTREAM"] <- "TELOMERE"}
	
			if (!is.na(DOWNSTREAM[1,"DIRECTION"]) & DOWNSTREAM[1,"DIRECTION"] == "-") {
				VARIANTS[j,"LOCATION.DOWNSTREAM"] <- "TERMINATOR"
			} else if (!is.na(DOWNSTREAM[1,"DIRECTION"])) {
				VARIANTS[j,"LOCATION.DOWNSTREAM"] <- "PROMOTER"
			} else {VARIANTS[j,"LOCATION.DOWNSTREAM"] <- "TELOMERE"}
		}
	
		###########################################
		#CHARACTERIZE EXONIC AND INTRONIC VARIANTS#
		###########################################

		if (nrow(GOOD.GENE) == 1) {
			VARIANTS[j,"GENE.FOCAL"] <- as.character(GOOD.GENE[1,"GENE"])
			VARIANTS[j,"GENE.SHORT"] <- as.character(GOOD.GENE[1,"SHORT"])
		}
			
		##################
		#CODING MUTATIONS#
		##################
			
		if (nchar(VARIANTS[j,"REF"]) > 1 | nchar(VARIANTS[j,"ALT"]) > 1) {
			VARIANTS[j,"TYPE"] <- "INDEL"
		}
		
		###A) CODING MUTATION IN GENES WITH 0 OR 1 INTRON###

		if (nchar(VARIANTS[j,"REF"]) == 1 & nchar(VARIANTS[j,"ALT"]) == 1 & VARIANTS[j,"CLASS"] == "EXONIC") {
		
			VARIANTS[j,"ORF.LENGTH"] <- GOOD.GENE[1,"LENGTH.CDS"]

			####A.1-DETERMINE REFERENCE AND ALTERNATIVE CODON####			
			
			####A.1.1-CASE WHERE GENE IS IN DIRECT ORIENTATION####

			if (GOOD.GENE[1,"DIRECTION"] == "+") {
				START <- GOOD.GENE[1,"START"]

				####GENE WITHOUT INTRON#################

				if (VARIANTS[j,"ALERT"] != "INTRON") {
					SNP.POS <- VARIANTS[j,"POSITION"] - START + 1
				}

				####GENE WITH INTRON####################

				if (VARIANTS[j,"ALERT"] == "INTRON") {
					GOOD.INTRON <- INTRONS[which(INTRONS$GENE == VARIANTS[j,"GENE.FOCAL"]),]

					if (VARIANTS[j,"POSITION"] < GOOD.INTRON[1,"START.NAME"])
					{
						SNP.POS <- VARIANTS[j,"POSITION"] - START + 1
					} else {
						SNP.POS.RNA <- VARIANTS[j,"POSITION"] - START + 1	
						INTRON.LENGTH <- GOOD.INTRON[1,"END.NAME"] - GOOD.INTRON[1,"START.NAME"] + 1
						SNP.POS <- SNP.POS.RNA - INTRON.LENGTH
					}
						
				}

				CODON.POS <- (SNP.POS-1)%%3 + 1

				if (CODON.POS == 1)
				{
					NUC.1 <- VARIANTS[j,"REF"]
					NUC.2 <- SEQ[CHR == as.character(VARIANTS[j,"CHROMOSOME"]) & POSITION == (VARIANTS[j,"POSITION"]+1)]
					NUC.2 <- as.data.frame(NUC.2)[1,"NUCLEOTIDE"]
					NUC.3 <- SEQ[CHR == as.character(VARIANTS[j,"CHROMOSOME"]) & POSITION == (VARIANTS[j,"POSITION"]+2)]
					NUC.3 <- as.data.frame(NUC.3)[1,"NUCLEOTIDE"]
		
					REF.CODON <- paste(NUC.1,NUC.2,NUC.3,sep="")
					VARIANTS[j,"REF.CODON"] <- REF.CODON
				
					NUC.1 <- VARIANTS[j,"ALT"]
					ALT.CODON <- paste(NUC.1,NUC.2,NUC.3,sep="")
					VARIANTS[j,"ALT.CODON"] <- ALT.CODON 
				}		
		
				if (CODON.POS == 2)
				{
					NUC.2 <- VARIANTS[j,"REF"]
					NUC.1 <- SEQ[CHR == as.character(VARIANTS[j,"CHROMOSOME"]) & POSITION == (VARIANTS[j,"POSITION"]-1)]
					NUC.1 <- as.data.frame(NUC.1)[1,"NUCLEOTIDE"]
					NUC.3 <- SEQ[CHR == as.character(VARIANTS[j,"CHROMOSOME"]) & POSITION == (VARIANTS[j,"POSITION"]+1)]
					NUC.3 <- as.data.frame(NUC.3)[1,"NUCLEOTIDE"]
		
					REF.CODON <- paste(NUC.1,NUC.2,NUC.3,sep="")
					VARIANTS[j,"REF.CODON"] <- REF.CODON
				
					NUC.2 <- VARIANTS[j,"ALT"]
					ALT.CODON <- paste(NUC.1,NUC.2,NUC.3,sep="")
					VARIANTS[j,"ALT.CODON"] <- ALT.CODON 
				}		


				if (CODON.POS == 3)
				{
					NUC.3 <- VARIANTS[j,"REF"]
					NUC.1 <- SEQ[CHR == as.character(VARIANTS[j,"CHROMOSOME"]) & POSITION == (VARIANTS[j,"POSITION"]-2)]
					NUC.1 <- as.data.frame(NUC.1)[1,"NUCLEOTIDE"]
					NUC.2 <- SEQ[CHR == as.character(VARIANTS[j,"CHROMOSOME"]) & POSITION == (VARIANTS[j,"POSITION"]-1)]
					NUC.2 <- as.data.frame(NUC.2)[1,"NUCLEOTIDE"]
		
					REF.CODON <- paste(NUC.1,NUC.2,NUC.3,sep="")
					VARIANTS[j,"REF.CODON"] <- REF.CODON
				
					NUC.3 <- VARIANTS[j,"ALT"]
					ALT.CODON <- paste(NUC.1,NUC.2,NUC.3,sep="")
					VARIANTS[j,"ALT.CODON"] <- ALT.CODON 
				}		

				VARIANTS[j,"ORF.POSITION"] <- SNP.POS
				VARIANTS[j,"AA.POSITION"] <-  ceiling(SNP.POS/3)
				VARIANTS[j,"PROT.LENGTH"] <- GOOD.GENE[1,"LENGTH.CDS"]/3

				VARIANTS[j,"REF.AA"] <- as.character(CODE[which(CODE$CODON == REF.CODON),"AMINO.ACID"])
				VARIANTS[j,"ALT.AA"] <- as.character(CODE[which(CODE$CODON == ALT.CODON),"AMINO.ACID"])
				
				if (VARIANTS[j,"REF.AA"] != VARIANTS[j,"ALT.AA"] & VARIANTS[j,"ALT.AA"] == "Stop") 
				{
					VARIANTS[j,"TYPE"] <- "NON.SENSE"
				} else if (VARIANTS[j,"REF.AA"] != VARIANTS[j,"ALT.AA"] & VARIANTS[j,"ALT.AA"] != "Stop") {
					VARIANTS[j,"TYPE"] <- "NON.SYNONYMOUS"
				} else {VARIANTS[j,"TYPE"] <- "SYNONYMOUS"}

			####A.1.2-CASE WHERE GENE IS IN REVERSE ORIENTATION####

			} else if (GOOD.GENE[1,"DIRECTION"] == "-") {
				START <- GOOD.GENE[1,"STOP"]
				SNP.POS <- START - VARIANTS[j,"POSITION"] + 1
				CODON.POS <- (SNP.POS-1)%%3 + 1

				if (CODON.POS == 1)
				{
					NUC.1 <- REV.COMP(VARIANTS[j,"REF"])
					NUC.2 <- SEQ[CHR == as.character(VARIANTS[j,"CHROMOSOME"]) & POSITION == (VARIANTS[j,"POSITION"]-1)]
					NUC.2 <- REV.COMP(as.data.frame(NUC.2)[1,"NUCLEOTIDE"])
					NUC.3 <- SEQ[CHR == as.character(VARIANTS[j,"CHROMOSOME"]) & POSITION == (VARIANTS[j,"POSITION"]-2)]
					NUC.3 <- REV.COMP(as.data.frame(NUC.3)[1,"NUCLEOTIDE"])
		
					REF.CODON <- paste(NUC.1,NUC.2,NUC.3,sep="")
					VARIANTS[j,"REF.CODON"] <- REF.CODON
				
					NUC.1 <- REV.COMP(VARIANTS[j,"ALT"])
					ALT.CODON <- paste(NUC.1,NUC.2,NUC.3,sep="")
					VARIANTS[j,"ALT.CODON"] <- ALT.CODON 
				}		
				
				if (CODON.POS == 2)
				{
					NUC.2 <- REV.COMP(VARIANTS[j,"REF"])
					NUC.1 <- SEQ[CHR == as.character(VARIANTS[j,"CHROMOSOME"]) & POSITION == (VARIANTS[j,"POSITION"]+1)]
					NUC.1 <- REV.COMP(as.data.frame(NUC.1)[1,"NUCLEOTIDE"])
					NUC.3 <- SEQ[CHR == as.character(VARIANTS[j,"CHROMOSOME"]) & POSITION == (VARIANTS[j,"POSITION"]-1)]
					NUC.3 <- REV.COMP(as.data.frame(NUC.3)[1,"NUCLEOTIDE"])
		
					REF.CODON <- paste(NUC.1,NUC.2,NUC.3,sep="")
					VARIANTS[j,"REF.CODON"] <- REF.CODON
				
					NUC.2 <- REV.COMP(VARIANTS[j,"ALT"])
					ALT.CODON <- paste(NUC.1,NUC.2,NUC.3,sep="")
					VARIANTS[j,"ALT.CODON"] <- ALT.CODON 
				}	
	
				if (CODON.POS == 3)
				{
					NUC.3 <- REV.COMP(VARIANTS[j,"REF"])
					NUC.1 <- SEQ[CHR == as.character(VARIANTS[j,"CHROMOSOME"]) & POSITION == (VARIANTS[j,"POSITION"]+2)]
					NUC.1 <- REV.COMP(as.data.frame(NUC.1)[1,"NUCLEOTIDE"])
					NUC.2 <- SEQ[CHR == as.character(VARIANTS[j,"CHROMOSOME"]) & POSITION == (VARIANTS[j,"POSITION"]+1)]
					NUC.2 <- REV.COMP(as.data.frame(NUC.2)[1,"NUCLEOTIDE"])
		
					REF.CODON <- paste(NUC.1,NUC.2,NUC.3,sep="")
					VARIANTS[j,"REF.CODON"] <- REF.CODON
				
					NUC.3 <- REV.COMP(VARIANTS[j,"ALT"])
					ALT.CODON <- paste(NUC.1,NUC.2,NUC.3,sep="")
					VARIANTS[j,"ALT.CODON"] <- ALT.CODON 
				}		
				
				VARIANTS[j,"ORF.POSITION"] <- SNP.POS
				VARIANTS[j,"AA.POSITION"] <-  ceiling(SNP.POS/3)
				VARIANTS[j,"PROT.LENGTH"] <- GOOD.GENE[1,"LENGTH.CDS"]/3
	
				VARIANTS[j,"REF.AA"] <- as.character(CODE[which(CODE$CODON == REF.CODON),"AMINO.ACID"])
				VARIANTS[j,"ALT.AA"] <- as.character(CODE[which(CODE$CODON == ALT.CODON),"AMINO.ACID"])
				
				if (VARIANTS[j,"REF.AA"] != VARIANTS[j,"ALT.AA"] & VARIANTS[j,"ALT.AA"] == "Stop") 
				{
					VARIANTS[j,"TYPE"] <- "NON.SENSE"
				} else if (VARIANTS[j,"REF.AA"] != VARIANTS[j,"ALT.AA"] & VARIANTS[j,"ALT.AA"] != "Stop") {
					VARIANTS[j,"TYPE"] <- "NON.SYNONYMOUS"
				} else {VARIANTS[j,"TYPE"] <- "SYNONYMOUS"}
				

			}		
			
	 			
		}
		
	}

	FILENAME <- paste(unlist(strsplit(VAR.PATH[i],".txt")),"_Annotated.txt",sep="")

	write.table(VARIANTS,FILENAME,sep="\t",row.names=FALSE,quote=FALSE)
	print(i)
}
```
