## Supplementary Table 1 for "Mutational sources of *trans-*regulatory variation affecting gene expression in *Saccharomyces cerevisiae*"

| **Supplementary Table 1. Statistical associations between aneuploidies and fluorescence level.** | | | | | | | | | | | | | | |
| --- | --- | --- | --- | --- | --- | --- | --- | --- | --- | --- | --- | --- | --- | --- |
| **Aneuploidy** | | | **Extra chromosome coverage** | | | | | **Whole-genome coverage** | | | | | **Statistics** | |
| *EMS Mutant* | *Extra chrom.* | *Fluo. level* | *Coverage low bulk* | *Coverage high bulk* | *N reads low bulk* | *N reads high bulk* | *Ratio low / high* | *Coverage low bulk* | *Coverage high bulk* | *N reads low bulk* | *N reads high bulk* | *Ratio low / high* | *G-value* | *P-value* |
| YPW2109 | 1 | 0.894 | 141x | 94x | 105388 | 71741 | 1.50 | 87x | 77x | 3778639 | 3373122 | 1.13 | 1552.4 | < 2.2E-16 |
| YPW2223 | 5 | 0.858 | 171x | 112x | 316588 | 207671 | 1.53 | 96x | 88x | 4237427 | 3781878 | 1.09 | 5682.4 | < 2.2E-16 |
| YPW2197 | 1 | 0.881 | 172x | 142x | 128040 | 105356 | 1.21 | 109x | 114x | 4739445 | 4872356 | 0.96 | 1406.0 | < 2.2E-16 |
